# A shared threat-anticipation circuit is dynamically engaged at different moments by certain and uncertain threat

**DOI:** 10.1101/2024.07.10.602972

**Authors:** Brian R. Cornwell, Paige R. Didier, Shannon E. Grogans, Allegra S. Anderson, Samiha Islam, Hyung Cho Kim, Manuel Kuhn, Rachael M. Tillman, Juyoen Hur, Zachary S. Scott, Andrew S. Fox, Kathryn A. DeYoung, Jason F. Smith, Alexander J. Shackman

**Affiliations:** Department of Psychological & Brain Sciences, George Washington University, Washington, DC 20006 USA; Department of Psychology, University of Maryland, College Park, MD 20742 USA; Department of Neuroscience and Cognitive Science Program, University of Maryland, College Park, MD 20742 USA; Maryland Neuroimaging Center, University of Maryland, College Park, MD 20742 USA; Department of Psychiatry and Human Behavior, Brown University, Providence, RI 02912 USA; Department of Psychology, University of Pennsylvania, Philadelphia, PA USA; Department of Child and Adolescent Psychiatry and Behavioral Sciences, Children’s Hospital of Philadelphia, Philadelphia, PA 19139 USA; Center for Depression, Anxiety and Stress Research, McLean Hospital, Harvard Medical School, Belmont, MA 02478 USA; McGill Neuropsychology, Bethesda, MD 20814 USA; Department of Psychology, Yonsei University, Seoul 03722, Republic of Korea; Department of Psychology, University of California, Davis, CA 95616 USA; California National Primate Research Center, University of California, Davis, CA 95616 USA

**Keywords:** human affective neuroscience, bed nucleus of the stria terminalis (BST/BNST), central extended amygdala (EAc), fear and anxiety, fMRI, Research Domain Criteria (RDoC)

## Abstract

Temporal dynamics play a central role in models of emotion: *“fear”* is widely conceptualized as a phasic response to certain-and-imminent danger, whereas *“anxiety”* is a sustained response to uncertain-or-distal harm. Yet the underlying neurobiology remains contentious. Leveraging a translationally relevant fMRI paradigm and theory-driven modeling approach in 220 adult humans, we demonstrate that certain- and uncertain-threat anticipation recruit a shared circuit that encompasses the central extended amygdala (EAc), periaqueductal gray, midcingulate, and anterior insula. This circuit exhibits persistently elevated activation when threat is uncertain and distal, and transient bursts of activation just before certain encounters with threat. Although there is agreement that the EAc plays a critical role in orchestrating responses to threat, confusion persists about the respective contributions of its major subdivisions, the bed nucleus of the stria terminalis (BST) and central nucleus of the amygdala (Ce). Here we used anatomical regions-of-interest to demonstrate that the BST and Ce exhibit statistically indistinguishable threat dynamics. Both regions exhibited activation dynamics that run counter to popular models, with the Ce showing sustained responses to uncertain-and-distal threat and the BST showing phasic responses to certain-and-imminent threat. For many scientists, feelings are the hallmark of fear and anxiety. Here we used an independently validated multivoxel brain ‘signature’ to covertly probe the moment-by-moment dynamics of anticipatory distress for the first time. Results mirrored the dynamics of neural activation. These observations provide fresh insights into the neurobiology of threat-elicited emotions and set the stage for more ambitious clinical and mechanistic research.

**SIGNIFICANCE STATEMENT:** *“Fear”* is widely viewed as a phasic response to certain-and-imminent danger, whereas *“anxiety”* is a sustained response to uncertain-or-distal harm. Prior work has begun to reveal the neural systems recruited by certain and uncertain anticipated threats, but has yet to rigorously plumb the moment-by-moment dynamics anticipated by theory. Here we used a novel combination of neuroimaging techniques to demonstrate that certain and uncertain threat recruit a common threat-anticipation circuit. Activity in this circuit and covert measures of distress showed similar patterns of context-dependent dynamics, exhibiting persistent increases when anticipating uncertain-threat encounters and transient surges just before certain encounters. These observations provide fresh insights into the neurobiology of fear and anxiety, laying the groundwork for more ambitious clinical and mechanistic research.

## INTRODUCTION

Fear and anxiety are evolutionarily conserved features of mammalian life that help protect us from harm (<u>Grogans et al., 2023</u>). But when expressed too strongly or pervasively, they can be crippling (<u>Salomon et</u> <u>al., 2015</u>). Anxiety-related disorders impose a staggering burden on global public health, afflicting ∼360 million individuals annually (<u>GBD, 2024</u>). In the U.S., ∼1 in 3 individuals will experience a lifetime disorder, service utilization is surging, and annual healthcare costs exceed $40B, drawing the attention of top policymakers (<u>WHO, 2022</u>; <u>Grogans et al., 2023</u>; <u>SAMHSA, 2023</u>; <u>White House, 2023</u>). Existing treatments are far from curative for many, underscoring the need to clarify the underlying neurobiology (<u>Fox and</u> <u>Shackman, 2024</u>).

Temporal dynamics play a central role in most models of fear and anxiety. Many theorists and clinicians conceptualize *“fear”* as a phasic response to certain-and-imminent danger and *“anxiety”* as a sustained response to uncertain-or-distal harm (<u>Barlow, 2000</u>; <u>Davis et al., 2010</u>; <u>Grupe and Nitschke, 2013</u>; <u>Tovote</u> <u>et al., 2015</u>; <u>Mobbs et al., 2020</u>; <u>Penninx et al., 2021</u>; <u>Moscarello and Penzo, 2022</u>; <u>Roelofs and Dayan, 2022</u>; <u>Grogans et al., 2023</u>). Work harnessing the millisecond resolution of psychophysiological measures supports this view, showing that human defensive behaviors exhibit specific temporal patterns across different threat contexts (<u>Grillon et al., 1993a</u>; <u>Grillon et al., 1993b</u>; <u>Löw et al., 2015</u>; <u>Moberg et al., 2017</u>; <u>Abend et al., 2022</u>). When the timing of threat encounters is uncertain, a sustained state of heightened reactivity is evident. In contrast, when encounters are certain and imminent, a phasic burst of heightened defensive responding is triggered. Both effects are consistent with evidence gleaned from animal research (<u>Millenson and Hendry, 1967</u>; <u>Blanchard et al., 1986</u>; <u>Chu et al., 2024</u>). Among humans, long-duration certain-threat cues, with explicit ‘count-down’ signals, elicit a mixture of weak sustained and robust phasic signals, resulting in a quadratic signal (<u>e.g., Grillon et al., 1993b</u>).

Neuroimaging studies have begun to reveal the regions recruited by certain- and uncertain-threat anticipation—including the central extended amygdala (EAc), midcingulate, and anterior insula—but have yet to systematically plumb the moment-by-moment neural dynamics anticipated by theory and psychophysiological research (<u>Chavanne and Robinson, 2021</u>; <u>Grogans et al., 2024</u>). Most studies have relied on simplified ‘boxcar’ modeling approaches that assume static, time-invariant neural responses to anticipated threat encounters. As critics have noted, this effectively reduces threat-related activation to a single average response, precluding inferences about more nuanced activation dynamics (<u>Wang et al.,</u> <u>2024</u>). Several groups have explored finer-grained models, but have yet to leverage them for rigorous hypothesis testing (<u>Hur et al., 2020b</u>; <u>Murty et al., 2023</u>). Consequently, it remains unclear whether phasic (*“fear”*) and sustained (*“anxiety”*) neural responses to threat are segregated into dissociable anatomical systems, as some have posited (<u>NIMH, 2011</u>; <u>Avery et al., 2016</u>; <u>LeDoux and Pine, 2016</u>; <u>NIMH, 2020b</u>, <u>a</u>), or are co-localized to a singular system that shows distinctive activation dynamics in response to certain- and uncertain-threat anticipation, as others have hypothesized (<u>Fox and Shackman, 2019</u>; <u>Hur et al.,</u> <u>2020b</u>; <u>Shackman et al., 2024</u>).

To help adjudicate this debate, we used a novel combination of fMRI techniques—including two complementary theory-driven hemodynamic models; focused region-of-interest (ROI) analyses of the EAc, a key player in many models of fear and anxiety; and multivoxel brain-signature analyses—to interrogate the moment-by-moment dynamics of threat-elicited neural activity and subjective distress in 220 racially diverse adults. Data were acquired using the Maryland Threat Countdown (MTC), a well-established paradigm for manipulating the temporal certainty of threat encounters (**Figure 1**) (<u>Hur et al., 2020b</u>). The MTC is an fMRI-optimized variant of assays that have been pharmacologically and psychophysiologically validated in rodents and humans, maximizing translational relevance (<u>Hur et al., 2020b</u>). Prior work in this sample and others demonstrates that the MTC robustly amplifies subjective symptoms of distress and objective signs of arousal (skin conductance), reinforcing its validity as an experimental probe of human fear and anxiety (<u>Kim et al., 2023</u>; Grogans et al., 2024).

**Figure 1.**
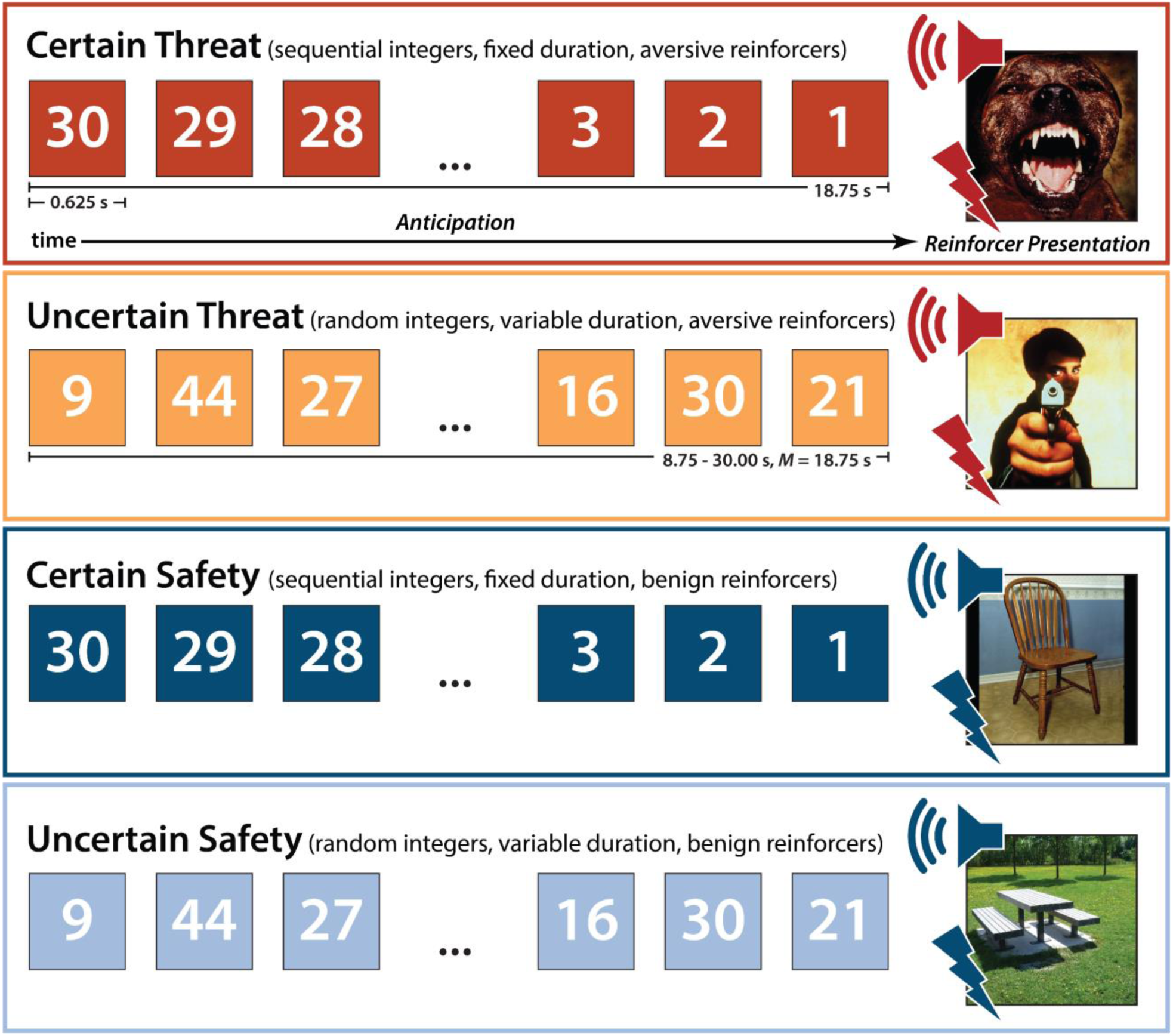
Threat-Anticipation paradigm. The Maryland Threat Countdown paradigm takes the form of a 2 (*Valence:* Threat/Safety) × 2 (*Temporal Certainty:* Certain/Uncertain) randomized event-related design. Participants were completely informed about the task design and contingencies prior to scanning. On certain-threat trials, participants saw a descending stream of integers (‘count-down’) for 18.75 s. To ensure robust emotion induction, the anticipation epoch always terminated with the presentation of a noxious electric shock, unpleasant photograph, and thematically related audio clip (e.g., scream). Uncertain-threat trials were similar, but the integer stream was randomized and presented for an uncertain and variable duration (8.75-30.00 s; *M*=18.75 s). Participants knew that something aversive was going to occur, but they had no way of knowing precisely *when*. Safety trials were similar but terminated with the delivery of emotionally neutral reinforcers (e.g., just-perceptible electrical stimulation). Abbreviation—s, seconds.

## MATERIALS AND METHODS

### Study Overview and Recruitment

As part of a recently completed prospective-longitudinal study focused on individuals at risk for the development of anxiety disorders and depression (R01-MH107444), we used well-established psychometric measures of Neuroticism/Negative Emotionality (N/NE) to screen 6,594 first-year university students (57.1% female; 59.0% White, 19.0% Asian, 9.9% African American, 6.3% Hispanic, 5.8% Multiracial/Other; *M*=19.2 years, *SD*=1.1 years) (Grogans et al., 2024). Screening data were stratified into quartiles (top quartile, middle quartiles, bottom quartile), separately for males and females. Individuals who met preliminary inclusion criteria were independently and randomly recruited via email from each of the resulting six strata. Because of the parent project’s focus on internalizing risk, approximately half the participants were recruited from the top quartile, with the remainder evenly split between the middle and bottom quartiles. This enabled us to sample a broad spectrum of psychiatric risk without gaps or discontinuities—in contrast to prior work focused on convenience samples—while balancing biological sex, consistent with recent recommendations (<u>Charpentier et al., 2021</u>; <u>Kang et al.,</u> <u>2024</u>). Simulations show that this over-sampling (‘enrichment’) approach does not bias statistical tests to a degree that would compromise their validity (<u>Hauner et al., 2014</u>). All participants had normal or corrected-to-normal color vision, and reported the absence of lifetime neurological symptoms, pervasive developmental disorder, very premature birth, medical conditions that would contraindicate MRI, and prior experience with noxious electrical stimulation. All participants were free from a lifetime history of psychotic and bipolar disorders; a current diagnosis of a mood, anxiety, or trauma disorder (past 2 months); severe substance abuse; active suicidality; and on-going psychiatric treatment as determined by an experienced masters-level diagnostician using the Structured Clinical Interview for DSM-5 (<u>First et al.,</u> <u>2015</u>). Participants provided informed written consent and all procedures were approved by the Institutional Review Board at the University of Maryland, College Park (Protocol #659385).

Data from this study were featured in prior work focused on validation of the threat-anticipation paradigm (<u>Hur et al., 2020b</u>; <u>Shackman et al., 2024</u>), neuroanatomical correlates of childhood anxiety (<u>Bas-</u> <u>Hoogendam et al., 2022</u>), threat-related neural activity and negative affect (<u>Hur et al., 2022</u>; <u>Grogans et al.,</u> <u>2024</u>), personality and internalizing symptoms (<u>Conway et al., 2024</u>), and social anxiety and negative affect (<u>Hur et al., 2020a</u>), but have never been used to address the present aims.

### Threat-Anticipation Paradigm

#### Paradigm Structure and Design Considerations

The Maryland Threat Countdown paradigm is a well-established, fMRI-optimized variant of temporally uncertain-threat assays that have been validated using fear-potentiated startle and acute anxiolytic administration (e.g., benzodiazepine) in mice, rats, and humans (<u>Miles et al., 2011</u>; <u>Hefner et al., 2013</u>; <u>Daldrup et al., 2015</u>; <u>Lange et al., 2017</u>; <u>Moberg et al., 2017</u>).

The paradigm has been successfully deployed in independent samples of samples of university students and community volunteers (<u>Kim et al., 2023</u>; <u>Grogans et al., 2024</u>).

As shown schematically in **Figure 1**, the paradigm takes the form of a 2 (*Valence:* Threat, Safety) × 2 (*Temporal Certainty:* Uncertain, Certain) randomized, event-related, repeated-measures design (3 scans; 6 trials/condition/scan). Participants were completely informed about the task design and contingencies prior to scanning. Simulations were used to optimize the detection and deconvolution of task-related hemodynamic signals. Stimulus presentation was controlled using Presentation software (version 19.0, Neurobehavioral Systems, Berkeley, CA).

On certain-threat trials, participants saw a descending stream of integers (‘count-down;’ e.g., 30, 29, 28…3, 2, 1) for 18.75 s. To ensure robust fear and anxiety, this anticipation epoch culminated with the presentation of a noxious electric shock, unpleasant photograph (e.g., mutilated body), and thematically related audio clip (e.g., scream). Uncertain-threat trials were similar, but the integer stream was randomized and presented for an uncertain and variable duration (8.75-30.00 s; *M*=18.75 s). Participants knew that something aversive was going to occur, but they had no way of knowing precisely when. Consistent with methodological recommendations (<u>Shackman and Fox, 2016</u>), the average duration of the anticipation epoch was identical across conditions, ensuring an equal number of measurements (TRs/condition). The specific mean duration was chosen to enhance detection of task-related differences in the blood oxygen level-dependent (BOLD) signal (‘activation’) (<u>Henson, 2007</u>) and to allow sufficient time for sustained responses to become evident. Certain- and uncertain-safety trials were similar but terminated with the presentation of benign reinforcers (see below). Valence was continuously signaled by the background color of the display. Temporal certainty was signaled by the nature of the integer stream. Certain trials always began with the presentation of the number 30. On uncertain trials, integers were randomly drawn from a near-uniform distribution ranging from 1 to 45 to reinforce the impression that they could be much shorter or longer than certain trials and to minimize incidental temporal learning (‘time-keeping’). To demonstrate the variable duration of Uncertain trials, during scanning, the first three uncertain trials featured short (8.75 s), medium (15.00 s), and long (28.75 s) anticipation epochs. To mitigate potential confusion and eliminate mnemonic demands, a lower-case ‘c’ or ‘u’ was presented at the lower edge of the display throughout the anticipatory epoch. White-noise visual masks (3.2 s) were presented between trials to minimize the persistence of visual reinforcers in iconic memory. Anticipatory distress ratings and skin conductance were also acquired, as detailed previously (<u>Grogans et al., 2024</u>).

#### Procedures

Prior to scanning, participants practiced an abbreviated version of the paradigm (without electrical stimulation) until they indicated and staff confirmed understanding. Benign and aversive electrical stimulation levels were individually titrated. *Benign Stimulation.* Participants were asked whether they could “reliably detect” a 20 V stimulus and whether it was “at all unpleasant.” If the participant could not detect the stimulus, the voltage was increased by 4 V and the process repeated. If the participant indicated that the stimulus was unpleasant, the voltage was reduced by 4 V and the process was repeated. The final level chosen served as the benign electrical stimulation during the imaging assessment (*M*=21.06 V, *SD*=5.06). *Aversive Stimulation.* Participants received a 100 V stimulus and were asked whether it was “as unpleasant as you are willing to tolerate”—an instruction specifically chosen to maximize anticipatory distress and arousal. If the participant indicated that they were willing to tolerate more intense stimulation, the voltage was increased by 10 V and the process repeated. If the participant indicated that the stimulus was too intense, the voltage was reduced by 5 V and the process repeated. The final level chosen served as the aversive electrical stimulation during the imaging assessment (*M*=117.85 V, *SD*=26.10). Following each scan, staff re-assessed whether stimulation was sufficiently intense and increased the level as necessary.

#### Electrical Stimuli

Electrical stimuli (100 ms; 2 ms pulses every 10 ms) were generated using an MRI-compatible constant-voltage stimulator system (STMEPM-MRI; Biopac Systems, Inc., Goleta, CA) and delivered using MRI-compatible, disposable carbon electrodes (Biopac) attached to the fourth and fifth digits of the non-dominant hand.

#### Visual Stimuli

A total of 72 aversive and benign photographs (1.8 s) were selected from the International Affective Picture System (<u>Hur et al., 2020b</u>). Visual stimuli were digitally back-projected (Powerlite Pro G5550, Epson America, Inc., Long Beach, CA) onto a semi-opaque screen mounted at the head-end of the scanner bore and viewed using a mirror mounted on the head-coil.

#### Auditory Stimuli

Seventy-two aversive and benign auditory stimuli (0.8 s) were adapted from open-access online sources and delivered using an amplifier (PA-1 Whirlwind) with in-line noise-reducing filters and ear buds (S14; Sensimetrics, Gloucester, MA) fitted with noise-reducing ear plugs (Hearing Components, Inc., St. Paul, MN).

### MRI Data Acquisition

MRI data were acquired using a Siemens Magnetom TIM Trio 3 Tesla scanner (32-channel head-coil). Foam inserts were used to immobilize the participant’s head within the head-coil and mitigate potential motion artifact. Participants were continuously monitored using an MRI-compatible eye-tracker (Eyelink 1000; SR Research, Ottawa, Ontario, Canada) and the AFNI real-time motion plugin (<u>Cox, 1996</u>). Sagittal T1-weighted anatomical images were acquired using a magnetization prepared rapid acquisition gradient echo sequence (TR=2,400 ms; TE=2.01 ms; inversion time=1,060 ms; flip=8°; slice thickness=0.8 mm; in-plane=0.8 × 0.8 mm; matrix=300 × 320; field-of-view=240 × 256). A T2-weighted image was collected co-planar to the T1-weighted image (TR=3,200 ms; TE=564 ms; flip angle=120°). To enhance resolution, a multi-band sequence was used to collect oblique-axial EPI volumes (multiband acceleration=6; TR=1,250 ms; TE=39.4 ms; flip=36.4°; slice thickness=2.2 mm, number of slices=60; in-plane resolution=2.1875 × 2.1875 mm; matrix=96 × 96). Images were collected in the oblique-axial plane (approximately −20° relative to the AC-PC plane) to minimize potential susceptibility artifacts. For the threat-anticipation task, three 478-volume EPI scans were acquired. The scanner automatically discarded 7 volumes prior to the first recorded volume. To enable fieldmap correction, two oblique-axial spin echo (SE) images were collected in opposing phase-encoding directions (rostral-to-caudal and caudal-to-rostral) at the same location and resolution as the functional volumes (i.e., co-planar; TR=7,220 ms; TE=73 ms). Measures of respiration and pulse were continuously acquired during scanning using a respiration belt and photo- plethysmograph affixed to the first digit of the non-dominant hand. Following the last scan, participants were removed from the scanner, debriefed, compensated, and discharged.

### MRI Pipeline

Methods were optimized to minimize spatial normalization error and other potential sources of noise and are similar to other recent work by our group (<u>e.g., Grogans et al., 2024</u>). Data were visually inspected before and after processing for quality assurance.

#### Anatomical Data Processing

T1- and T2-weighted images were inhomogeneity corrected using *N4* (<u>Tustison et al., 2010</u>) and denoised using *ANTS* (<u>Avants et al., 2011</u>). The brain was then extracted using *BEaST* (<u>Eskildsen et al., 2012</u>) and brain-extracted and normalized reference brains from *IXI* (<u>BIAC, 2022</u>).

Brain-extracted T1 images were normalized to a version of the brain-extracted 1-mm T1-weighted MNI152 template (<u>non-linear 6th-generation symmetric average; Grabner et al., 2006</u>) modified to remove extracerebral tissue. Normalization was performed using the diffeomorphic approach implemented in *SyN* (version 2.3.4) (<u>Avants et al., 2011</u>). T2-weighted images were rigidly co-registered with the corresponding T1 prior to normalization. The brain extraction mask from the T1 was then applied. Tissue priors were unwarped to native space using the inverse of the diffeomorphic transformation (<u>Lorio et al., 2016</u>). Brain-extracted T1 and T2 images were segmented using native-space priors generated in *FAST* (version 6.0.4) (<u>Jenkinson et al., 2012</u>) for subsequent use in T1-EPI co-registration (see below).

#### Fieldmap Data Processing

SE images and *topup* were used to create fieldmaps. Fieldmaps were converted to radians, median-filtered, and smoothed (2-mm). The average of the distortion-corrected SE images was inhomogeneity corrected using *N4* and masked to remove extracerebral voxels using *3dSkullStrip* (*AFNI* version 23.1.10). The resulting mask was minimally eroded to further exclude extracerebral voxels.

#### Functional Data Processing

EPI files were de-spiked (*3dDespike*), slice-time corrected to the TR-center using *3dTshift*, and motion-corrected to the first volume and inhomogeneity corrected using *ANTS* (12-parameter affine). Transformations were saved in ITK-compatible format for subsequent processing (<u>McCormick et al., 2014</u>). The first volume was extracted for EPI-T1 co-registration. The reference EPI volume was simultaneously co-registered with the corresponding T1-weighted image in native space and corrected for geometric distortions using boundary-based registration (<u>Jenkinson et al., 2012</u>). This step incorporated the previously created fieldmap, undistorted SE, T1, white matter (WM) image, and masks. The spatial transformations necessary to transform each EPI volume from native space to the reference EPI, from the reference EPI to the T1, and from the T1 to the template were concatenated and applied to the processed EPI data in a single step to minimize incidental spatial blurring. Normalized EPI data were resampled (2 mm^3^) using fifth-order b-splines. Voxelwise analyses employed data that were spatially smoothed (4-mm) using *3DblurInMask*. To minimize signal mixing, smoothing was confined to the gray-matter compartment, defined using a variant of the Harvard-Oxford cortical and subcortical atlases that was expanded to include the bed nucleus of the stria terminalis (BST) and periaqueductal gray (PAG) (<u>Frazier et al., 2005</u>; <u>Desikan et al., 2006</u>; <u>Makris et al., 2006</u>; <u>Edlow et al., 2012</u>; <u>Theiss et al., 2017</u>). Focal analyses of the EAc leveraged spatially unsmoothed data and anatomically defined regions of interest (see below), as in prior work (<u>Tillman et al., 2018</u>; <u>Kim et al., 2023</u>; <u>Grogans et al., 2024</u>; <u>Hur et al., *in press*</u>).

### fMRI Data Exclusions and Hemodynamic Modeling

#### Data Exclusions

Volume-to-volume displacement (>0.5 mm) was used to assess residual motion artifact. Scans with excessively frequent residual artifacts (>2 *SD*) were discarded. Participants with insufficient usable fMRI data (<2 scans) were excluded from analyses (see above).

#### Overview of First-Level (Single-Subject) fMRI Modeling

For each participant, first-level modeling was performed using general linear models (GLMs) implemented in *3dREMLfit* (ARMA_1,1_; 4^th^-order Legendre high-pass filter). Regressors were convolved with the *SPM12* canonical hemodynamic-response function (HRF). Epochs corresponding to the presentation of the four types of reinforcers, white-noise visual masks, and rating prompts were simultaneously modeled using the same approach. As in our prior work, nuisance variates included volume-to-volume displacement and its first derivative, 6 motion parameters and their first derivatives, cerebrospinal fluid (CSF) signal, instantaneous pulse and respiration rates, and nuisance signals (e.g., brain edge, CSF edge, global motion, white matter, extracerebral soft tissue) (<u>Anderson et al.,</u> <u>2011</u>; <u>Pruim et al., 2015</u>). Volumes with excessive volume-to-volume displacement (>0.75 mm) and those during and immediately following reinforcer delivery were censored. EPI volumes acquired before the first trial and following the final trial were unmodeled and contributed to the baseline estimate.

#### Conventional ‘Boxcar’ Model

The present sample of 220 datasets represents a superset of the 99 featured in an earlier report from our group that employed a conventional ‘boxcar’ fMRI modeling approach and an older data-processing pipeline (<u>Hur et al., 2020b</u>). As a precursor to hypothesis testing, we used a conventional first-level model to confirm that the larger, reprocessed dataset broadly reproduced our published observations. Hemodynamic reactivity to the threat-anticipation paradigm was modeled using variable-duration rectangular (‘boxcar’) regressors that spanned the entirety of the anticipation (‘countdown’) epoch for uncertain-threat, certain-threat, and uncertain-safety trials (8.75-30.00 s; **Figure 1**). To maximize design efficiency, certain-safety anticipation served as the first-level reference condition and contributed to the baseline estimate (<u>Poline et al., 2007</u>).

#### Onset-Sustained-Phasic (OSP) Model

Neuroimaging research by our group and others has relied on simplified ‘boxcar’ modeling approaches that reduce the neural dynamics anticipated by theory and psychophysiological research to a single average response (<u>Wang et al., 2024</u>). Here we used two complementary hemodynamic models to characterize the time-varying signals elicited by certain- and uncertain-threat anticipation. The OSP model used a multiple-regression framework to identify the variance in threat-anticipation signals that was uniquely associated, in the partial-correlation sense, with temporally overlapping Onset, Sustained, and Phasic regressors (**Figure 2a**). The first-level design matrix incorporated a punctate event or ‘impulse’ time-locked to the onset of the anticipation epoch, a variable-duration rectangular function that spanned the entirety of the anticipation epoch (to capture sustained increases in activation), and a rectangular-function time-locked to the offset of the anticipation epoch (to capture phasic surges in activation just prior to threat encounters). Paralleling the ‘boxcar’ model, the Sustained regressor for certain-safety served as the first-level reference condition and contributed to the baseline estimate. The duration of the Phasic regressor (6.15 s) was chosen based on a combination of theory and simulations aimed at minimizing regressor co-linearity. The mean condition-wise variance inflation factor was <1.93 for the full task-related model (excluding nuisance regressors). As detailed in **Figure 2**, the Sustained regressor captures variance in the hemodynamic signal associated with a particular trial type (e.g., uncertain-threat anticipation) above-and-beyond that captured by the Onset and Phasic regressors. Unlike conventional boxcar models, this provides an estimate of sustained activation that is unconfounded by non-specific orienting or salience responses reflexively triggered by the dramatic change in sensory stimulation associated with trial onset (<u>Sokolov et al., 2002</u>; <u>Menon, 2015</u>). Likewise, the Phasic regressor captures variance in the hemodynamic signal above-and-beyond that captured by the Onset and Sustained regressors, with positive coefficients indicating an increase in activation in the final moments of the anticipation epoch relative to that associated with the Sustained and Onset regressors. In sum, the OSP model casts the overall magnitude of the hemodynamic signal as a linear combination of the Onset, Sustained, and Phasic regressors; nuisance regressors; and error (**Figure 2**).

**Figure 2.**
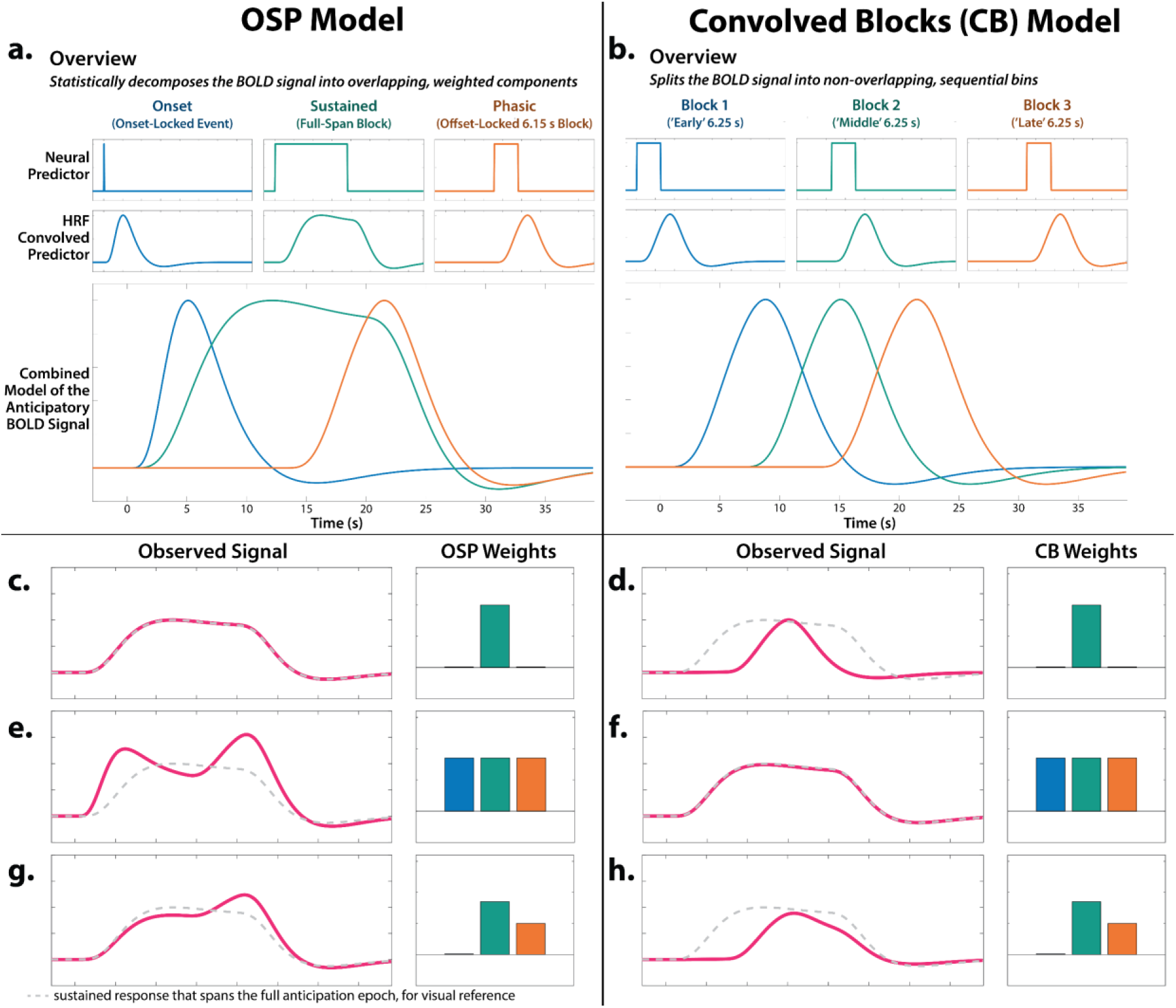
Theory-driven hemodynamic modeling. We used two modeling approaches to investigate time-varying responses to certain- and uncertain-threat anticipation. The relative merits of the two models are described in the main text (‘Complementary Strengths and Limitations of the Two Models’). Because these models are uncommon, we provide a general overview in panels *a*-*b* and, for readers interested in greater detail, a description of their nuances in panels *c*-*f*. **a. *Overview of the OSP Model.*** The OSP model used multiple-regression to partition the variance in threat-anticipation signals uniquely associated with temporally overlapping Onset, Sustained, and Phasic regressors (see main text for details). The OSP is ideal for examining between *within*-moment contrasts (e.g., *Phasic regressor:* certain vs. uncertain threat). **b. *Overview of the Convolved Blocks (CB) Model.*** The CB model splits the anticipation epoch into a sequence of short (6.25 s), non-overlapping rectangular functions or ‘blocks,’ each convolved with a canonical HRF. The CB model is ideal for examining *between*-moment contrasts and temporal trends (e.g., quadratic effects), and it enabled a rigorous test of surges in activation in the moments just prior to certain-threat encounters (*Certain-Threat:* late vs. middle block)—something not permitted by the OSP model. **c.** Consider a voxel that shows a sustained level of heightened activation during certain-threat anticipation (*red line*). In the OSP model, this is captured by a strong loading or ‘weight’ on the Sustained regressor and nil loadings on the Onset and Phasic regressors. **d.** In the CB model, this pattern of weights is instead associated with a transient increase in the hemodynamic signal in the middle of the anticipation epoch. **f.** In the CB model, sustained activation is captured by uniformly strong loadings across the early, middle, and late regressors. **e.** In the OSP model, this pattern of weights is instead associated with transient onset and phasic responses, superimposed on a strong sustained response. In effect, the Onset and Phasic regressors serve to modulate the leading and trailing edges of a sustained wave of activation. It merits comment that, although the three weights are equally strong (*barplot: O* ≈ *S* ≈ *P*), the moment-by-moment height of the hemodynamic signal is not (*red line: O* > *S* < *P*). This reflects the fact that the OSP model casts the threat-anticipation signal as the linear combination of 3 temporally overlapping regressors. **g.** In the OSP model, the relative height of activation (*red line: S* < *P*) can be reversed from the rank order of the weights (*barplot: S* > *P*). **h.** In contrast, the CB model provides a one-to-one mapping between the moment-by-moment height of the hemodynamic signal and the early, middle, and late regression weights. Abbreviations—BOLD, blood-oxygenation-level-dependent; CB, Convolved Blocks model; HRF, hemodynamic response function; fMRI, functional magnetic resonance imaging; OSP, Onset-Sustained-Phasic; s, seconds.

#### Convolved Blocks Model

To clarify interpretation (see next section), we employed a piecewise approach that arbitrarily splits the anticipation epoch into a sequence of 2-5 short (6.25 s), non-overlapping rectangular functions or ‘blocks’, each convolved with a canonical HRF (**Figure 2b**). Here the second certain-safety block (6.25-12.5 s) served as the first-level reference condition and contributed to the baseline estimate. Regressor co-linearity was acceptable (<u>Mumford et al., 2015</u>). The mean condition-wise variance inflation factor was <1.91 for the full task-related model.

#### Complementary Strengths and Limitations of the Two Models

The OSP and Convolved Blocks models have complementary strengths and limitations (**Figure 2**). The OSP Phasic regressor is time-locked to the *offset* of the anticipation epoch, ensuring that it always indexes neural activation in the seconds just before threat is encountered—regardless of its temporal certainty or the overall duration of the anticipation (‘countdown’) epoch. Because it is effectively a partial correlation, the OSP Phasic regressor captures variance in activation above-and-beyond that captured by the temporally overlapping Sustained regressor, providing a more ‘pure’ or conservative estimate of phasic surges in activation (**Figures 2c, 2e, 2g**). In short, the OSP model provides a unified, non-arbitrary way to address the variable duration of different trial types, making it ideal for examining *within*-moment contrasts (e.g., *Phasic regressor:* certain vs. uncertain threat). Despite these strengths, the partial correlations yielded by OSP model do not permit straightforward assessments of dynamic changes in activation. In contrast, the Convolved Blocks model, while arbitrary in timing, yields activation estimates that are statistically independent and directly comparable across moments of time (**Figures 2d**, **2f**, **2h**). As such, the Convolved Blocks model is ideal for examining *between*-moment contrasts and overall temporal trends in anticipatory activation (e.g., quadratic effects). In particular, the Convolved Blocks model provides a natural way to rigorously test hypothesized surges in activation in the seconds just prior to temporally certain encounters with threat (*Certain-Threat:* late vs. middle block).

#### EAc ROIs

The central extended amygdala (EAc) occupies center stage in most neurobiological models of fear and anxiety, including RDoC (<u>e.g., NIMH, 2011</u>; <u>e.g., Tovote et al., 2015</u>; <u>LeDoux and Pine, 2016</u>; <u>Fox</u> <u>and Shackman, 2019</u>; <u>Mobbs et al., 2019</u>; <u>NIMH, 2020b</u>, <u>a</u>; <u>Tseng et al., 2023</u>; <u>Akiki et al., 2025</u>). The EAc is a functional macrocircuit encompassing the central nucleus of the amygdala (Ce) and the neighboring bed nucleus of the stria terminalis (BST) (<u>Fox et al., 2015a</u>; <u>Shackman and Fox, 2016</u>; <u>Fox and Shackman, 2019</u>). As in prior work by our group (<u>e.g., Grogans et al., 2024</u>), EAc activation was quantified using anatomically defined probabilistic ROIs (<u>Theiss et al., 2017</u>; <u>Tillman et al., 2018</u>). The BST ROI was modified to remove voxels encroaching upon neighboring regions of the striatum, thalamus, and ventricles. It mostly encompasses the supra-commissural BST, given the difficulty of reliably discriminating the borders of regions below the anterior commissure in T1-weighted images (<u>Walter et al., 1991</u>; <u>Kruger et al., 2015</u>).

Bilateral ROIs were decimated to the 2-mm resolution of the fMRI data.

### Experimental Design and Statistical Analysis

#### Participants

A total of 241 participants were recruited and scanned. Of these, 6 withdrew from the study due to excess distress during the imaging session and 1 withdrew for undisclosed reasons following the imaging session. Another 14 participants were excluded from fMRI analyses due to incidental neurological findings (*n*=4), technical problems (*n*=2), or insufficient usable data (*n*=8; see below), yielding a racially diverse sample of 220 participants (49.5% female; 61.4% White, 18.2% Asian, 8.6% African American, 4.1% Hispanic, 7.3% Multiracial/Other; *M*=18.8 years, *SD*=0.4 years).

#### Power Analysis

To enable readers to better interpret in our results, we performed a post hoc power analysis. G-power (version 3.1.9.2) indicated that the final sample of 220 usable fMRI datasets provides 80% power to detect a benchmark (‘generic’) mean difference as small as Cohen’s *d*=0.19 (α=0.05, two-tailed) (<u>Cohen, 1988</u>; <u>Faul et al., 2007</u>). The study was not preregistered.

#### Analytics Overview

Analyses were performed using a combination of *SPM12* (<u>Wellcome Centre for Human</u> <u>Neuroimaging, 2022</u>), *SPSS* (version 27.0.1), and *JASP* (version 0.16.4.0) (<u>Love et al., 2019</u>). Diagnostic procedures and data visualizations were used to confirm that test assumptions were satisfied (<u>Tukey,</u> <u>1977</u>). Some figures were created using *R* (version 4.0.2), *Rstudio* (version 1.2.1335), *tidyverse* (version 2.0), *ggplot2* (version 3.3.6) *ggpubr* (version 0.6.0), *plotrix* (version 3.8-4), and *MRIcron (*version 1.0.20190902) (<u>Lemon, 2006</u>; <u>Wickham, 2016</u>; <u>Rorden, 2019</u>; <u>Wickham et al., 2019</u>; <u>R Core Team, 2022</u>; <u>RStudio Team, 2022</u>; <u>Kassambara, 2023</u>). Clusters and peaks were labeled using the Harvard–Oxford atlas, supplemented by the Mai, Allen, Jülich (version 3.1), and other atlases and resources (<u>Frazier et al., 2005</u>; <u>Desikan et al., 2006</u>; <u>Makris et al., 2006</u>; <u>Mai et al., 2015</u>; <u>Ding et al., 2016</u>; <u>Theiss et al., 2017</u>; <u>ten Donkelaar</u> <u>et al., 2018</u>; <u>Tillman et al., 2018</u>; <u>Amunts et al., 2021</u>).

#### Resource Sharing

Raw data are available at the National Institute of Mental Health Data Archive (https://nda.nih.gov/edit_collection.html?id=2447). Neuroimaging maps are available at NeuroVault (https://neurovault.org/collections/15274). Task materials, statistical code, de-identified processed data, and neuroimaging cluster tables are available at OSF (https://osf.io/e2ngf). The negative affect brain signature is available at Github (https://github.com/canlab/Neuroimaging_Pattern_Masks/tree/master/Multivariate_signature_patterns/2021_Ceko_MPA2_multiaversive).

#### Conventional ‘Boxcar’ Model

The present sample of 220 datasets represents a superset of the 99 featured in an earlier report that employed a conventional ‘boxcar’ hemodynamic-modeling approach, older data-processing pipeline, and larger spatial-smoothing kernel (6-mm) (<u>Hur et al., 2020b</u>). Here we used standard voxelwise GLMs to confirm that conventional modeling of the larger, reprocessed dataset broadly reproduced our published results (FDR *q*<0.05, whole-brain corrected). As in prior work by our group (<u>Hur</u> <u>et al., 2020b</u>; <u>Kim et al., 2023</u>), a minimum-conjunction analysis (Logical ‘AND’) was used to identify regions showing significant activation during the anticipation of certain *and* uncertain threat relative to their respective second-level reference conditions (<u>Nichols et al., 2005</u>).

#### Voxelwise Analyses of Sustained and Phasic Activation Dynamics

A major goal of the present study was to identify regions showing sustained levels of heighted activation during uncertain-threat anticipation and phasic surges in activation during the final moments of certain-threat anticipation. Standard voxelwise GLMs were used to compare each kind of anticipated threat (e.g., uncertain threat) to its second-level reference condition (e.g., uncertain safety) and to one another (e.g., certain threat; FDR *q*<0.05, whole-brain corrected). Hypothesis testing focused on the OSP Sustained and Phasic regressors. A minimum-conjunction analysis was used to identify regions showing significant sustained activation during uncertain-threat anticipation *and* significant phasic activation during the terminal portion of certain-threat anticipation, that is, regions showing evidence of neuroanatomical co-localization (<u>Fox and Shackman,</u> <u>2019</u>; <u>Hur et al., 2020b</u>; <u>Shackman and Fox, 2021</u>; <u>Shackman et al., 2024</u>). While not a focus of the present study, exploratory analyses of the OSP Onset regressor, which mainly captures reflexive orienting responses, are also briefly summarized.

#### Focused Tests of Phasic Activation

While the OSP model is well-suited for *within*-moment contrasts (e.g., *Phasic regressor:* certain vs. uncertain threat), the resulting partial-regression coefficients do not permit straightforward interpretation of *between*-moment contrasts (for details, see above and **Figure 2**). To more fully test phasic effects, we used activation estimates from the Convolved Blocks model and a standard voxelwise GLM to identify regions showing significant increases in activation during the final (12.5-18.75 s) relative to the middle (6.25-12.5 s) third of the certain-threat anticipation epoch (FDR *q*<0.05, whole-brain corrected).

We used activation estimates derived from the Convolved Blocks model and a standard voxelwise GLM to identify regions that differed in their responses to the final block of certain threat (an indicator of phasic surges in activation) compared to the second block of uncertain threat (an indicator of sustained activation), each relative to their respective reference condition (FDR *q*<0.05, whole-brain corrected).

#### EAc ROI Analyses

ROI analyses used activation estimates (i.e., standardized regression coefficients) for convolved blocks 1-3, extracted and averaged for each contrast (e.g., uncertain-threat vs. uncertain-safety anticipation), region, and participant (enabling us to examine overall temporal trends). This approach enabled us to span the mean duration of the anticipation epoch (0-18.75 s). Unlike conventional whole-brain voxelwise analyses—which screen thousands of voxels for statistical significance and yield optimistically biased associations—anatomically defined ROIs ‘fix’ the measurements-of-interest *a priori*, providing statistically unbiased effect-size estimates (<u>Poldrack et al., 2017</u>). As a precursor to hypothesis testing, we used one-sample Student’s *t*-tests to confirm that the Ce and BST ROIs are engaged by anticipated threat (*p*<0.05, uncorrected). For hypothesis testing, we used a standard 2 (*Region:* BST, Ce) × 2 (*Threat*-*Certainty:* Certain, Uncertain) × 3 (*Convolved Block:* 1, 2, 3) repeated-measures GLM (Huynh– Feldt correction) to interrogate potential regional differences in activation dynamics. Paralleling the voxelwise analyses, hypothesis testing focused on the second block (6.25-12.5 s) of uncertain-threat anticipation (a proxy for sustained activation) and the final block (12.5-18.75 s) of certain-threat anticipation (a proxy for phasic surges in activation). Significant GLM effects were interrogated using linear and quadratic polynomial contrasts, as in prior psychophysiological research (<u>Grillon et al., 1993a</u>; <u>Grillon</u> <u>et al., 1993b</u>; <u>Löw et al., 2015</u>). Polynomial-trend analyses enabled us to evaluate whether the time-course of activation differed as a function of Region, Threat-Certainty, or their interaction. In particular, we sought to test whether certain-and-imminent threat is associated with phasic surges in activation in the final third of the anticipation epoch (relative to the middle third), whether this hypothesized quadratic trend is stronger than that evinced during the identical moments of uncertain-threat anticipation, and whether these temporal dynamics differ between the BST and Ce.

Frequentist (Cohen’s *d*) and Bayesian (*BF_10_*) effect sizes were used to clarify non-significant regional differences. Cohen’s *d* was interpreted using established benchmarks (<u>Cohen, 1988</u>; <u>Cohen, 1994</u>; <u>Schimmack, 2019</u>), ranging from *small* (*d*=0.20) to *nil* (*d*≤0.10). *BF_10_* quantifies the relative performance of the null hypothesis (*H_0_*; e.g., the absence of a credible mean difference) and the alternative hypothesis (*H_1_*; e.g., the presence of a credible mean difference) on a 0 to ∞ scale. A key advantage of the Bayesian approach is that it can be used to formally quantify the relative strength of the evidence for H_0_ (‘test the null’), in contrast to standard null-hypothesis significance tests (<u>Wagenmakers et al., 2018</u>; <u>Bo et al., 2024</u>). It also does not require the data analyst to decide what constitutes a trivial difference, unlike traditional equivalence tests (<u>Hur et al., 2020b</u>). *BF_10_* was interpreted using established benchmarks (<u>van Doorn et al.,</u> <u>2021</u>). Values <1 were interpreted as evidence of statistical equivalence (i.e., support for the null hypothesis), ranging from s*trong* (*BF_10_*≤0.10), to *moderate* (*BF_10_*=0.10-0.33), to *weak* (*BF_10_*=0.33-1). The reciprocal of *BF_10_* represents the relative likelihood of the null hypothesis (e.g., *BF_10_*=0.10, *H_0_* is 10 times more likely than *H_1_*).

At the behest of a reviewer, we also explored potential differences in BST and Ce function using activation estimates derived from the OSP model. Because the OSP model does not permit meaningful between-moment contrasts, we used paired Student’s *t*-tests to compare (a) sustained responses to uncertain-threat and (b) phasic responses to certain-threat anticipation, each relative to their second-level reference conditions (e.g., certain threat vs. certain safety).

#### Brain Signature Analyses

Sandard fMRI analyses cannot address the momentary dynamics of threat-evoked emotions (<u>Poldrack, 2011</u>; <u>Grogans et al., 2023</u>). Here we used an independently trained and validated multivoxel pattern or ‘signature’ of negative affect to covertly probe moment-by-moment fluctuations in anticipatory distress during the ‘countdown’ period—something that would otherwise entail the imposition of a secondary rating task (e.g., using a continuous dial or randomized prompts), with unknown consequences for on-going emotional experience. Čeko, Wager, and colleagues used machine-learning to develop a pattern of voxelwise weights predictive of the intensity of negative affect in unseen data (<u>Čeko et al., 2022</u>). They demonstrated that the signature is a sensitive indicator of distress elicited by a range of noxious experiences—including thermal and mechanical pain, unpleasant photographs, and aversive auditory stimuli—but unrelated to the intensity of feelings triggered by positive stimuli, indicating specificity. We computed the dot-product between the negative-affect signature and activation estimates derived for the present sample using the Convolved Blocks model, enabling us to generate signature responses (a probabilistic estimate of negative affect intensity) for every combination of threat certainty (certain, uncertain), block (1, 2, 3), and participant. We used one-sample Student’s *t*-tests to confirm that the signature, which was trained using activation estimates time-locked to the presentation of aversive stimuli, is sensitive to the anticipation of threat encounters (*p*<0.05, uncorrected). For hypothesis testing, we used a standard 2 (*Threat Certainty:* Certain, Uncertain) × 3 (*Convolved Block:* 1, 2, 3) repeated-measures GLM (Huynh–Feldt correction) to assess dynamic fluctuations in signature-estimated distress across threat contexts. Significant GLM effects were decomposed using polynomial-trend analyses, as detailed in the prior section.

## RESULTS

### Overview

Temporal dynamics play a prominent role in models of emotion and emotional illness: *“anxiety”* is often conceptualized as a sustained response to uncertain-or-distal harm, whereas *“fear”* is a phasic response to certain-and-imminent danger (<u>Clark et al., 2017</u>; <u>Grogans et al., 2023</u>). Yet the underlying neurobiology remains incompletely understood. The Results are organized into three general sections. In the first section, we use two complementary hemodynamic models—one optimized for *within*-moment contrasts, the other for *between*-moment contrasts (**Figure 2**)—to characterize the neural systems showing sustained and phasic responses to different kinds of anticipated threat. In the second section, we use anatomical regions-of-interest to probe the functional architecture of the EAc, a central player in neurobiological models of fear and anxiety. For many scientists, feelings are the hallmark of fear and anxiety. In the third section, we use an independently validated multivoxel brain ‘signature’ to covertly decode the moment-by-moment dynamics of threat-elicited subjective distress.

### Conventional ‘boxcar’ modeling reveals a shared threat-anticipation circuit

The present sample of 220 datasets represents a superset of the 99 featured in an earlier report that relied on a conventional ‘boxcar’ modeling approach, older data-processing pipeline, and coarser spatial-smoothing kernel (6-mm) (<u>Hur et al., 2020b</u>). As a precursor to hypothesis testing, we used standard voxelwise GLMs to confirm that conventional boxcar modeling of the larger and reprocessed dataset broadly reproduced our previously published results. As expected, results revealed significant activation during periods of uncertain-threat anticipation, both in subcortical regions implicated in rodent models of fear and anxiety—such as the periaqueductal gray (PAG), BST, and dorsal amygdala—and in frontocortical regions that are especially well-developed in primates—including the midcingulate cortex (MCC), anterior insula/frontal operculum (AI/FrO), and rostral dorsolateral prefrontal cortex (dlPFC; FDR *q*<0.05, whole-brain corrected; **Figure 3** and **Supplementary Tables S1-S5,** https://osf.io/e2ngf). The same pattern was evident during certain-threat anticipation, with overlapping voxels evident for both kinds of threat in each of these key regions. In short, when viewed through the macroscopic lens of conventional fMRI modeling (<u>Fox and Shackman, 2024</u>), uncertain- and certain-threat anticipation engage co-localized neural circuits, suggesting a common neural substrate in humans.

**Figure 3.**
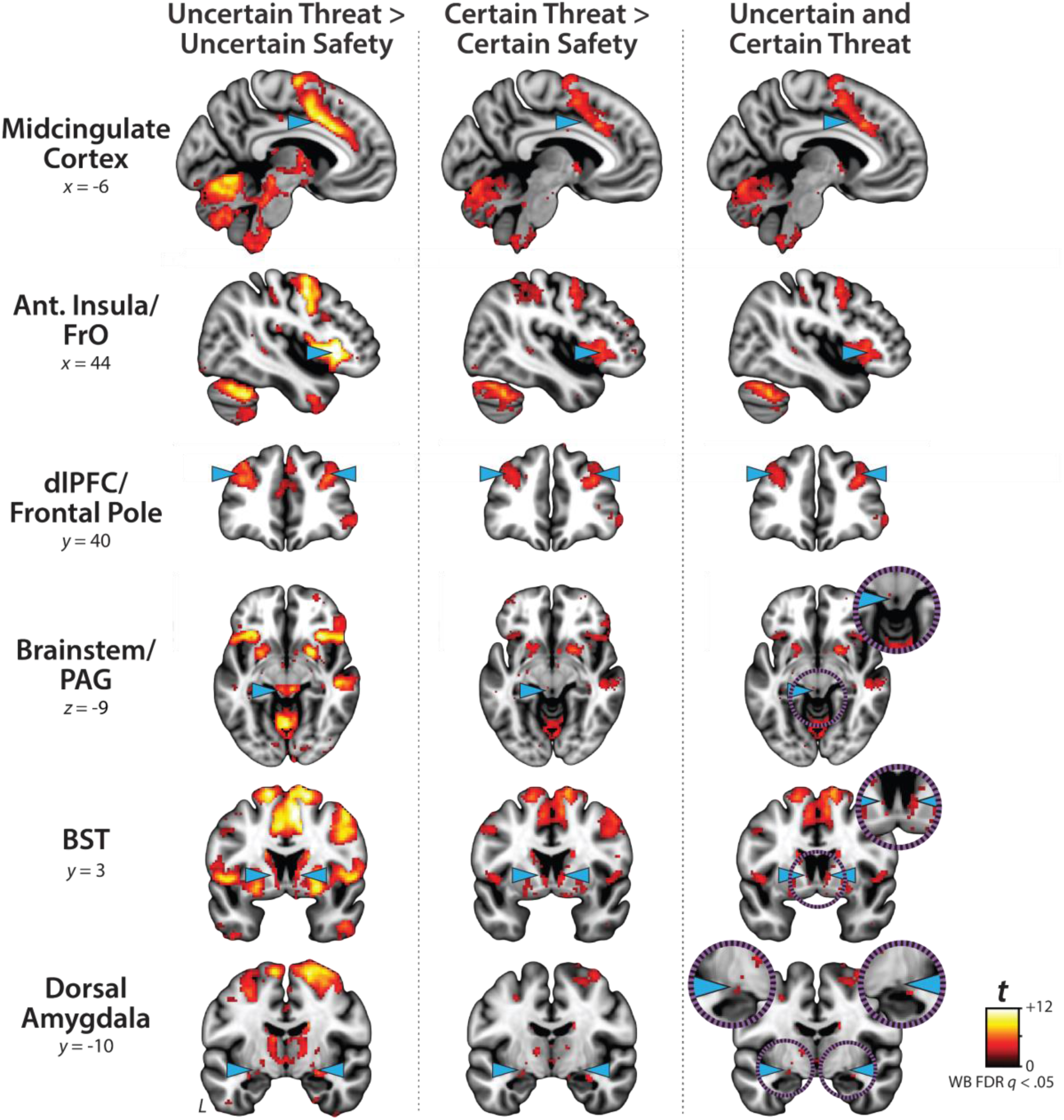
Uncertain- and certain-threat anticipation recruit a shared cortico-subcortical circuit. To facilitate comparison with prior work, we computed a conventional ‘boxcar’ analysis, which models the anticipation epoch (‘countdown’) as a single average response. As shown in the *left column*, uncertain-threat anticipation was associated with significant activation across a widely distributed network of regions previously implicated in the expression and regulation of human fear and anxiety (FDR *q*<.05, whole-brain corrected) (<u>Shackman and Fox, 2021</u>). As shown in the *middle column*, similar results were evident for certain-threat anticipation. In fact, as shown in the right column, a minimum-conjunction (logical ‘AND’) analysis of the two thresholded contrasts confirmed voxelwise co-localization in every key region. These observations replicate prior work in university and community samples, confirm that the MTC paradigm robustly engages the canonical threat-anticipation circuit, and set the stage for more granular analyses of neural dynamics (<u>Hur et al., 2020b</u>; <u>Kim et al., 2023</u>; <u>Grogans et al., 2024</u>; <u>Shackman et al., 2024</u>). Note. To enhance resolution, these analyses leveraged a smaller spatial-smoothing kernel (4-mm) than prior work by our group (6-mm). Abbreviations—Ant., anterior; BST, bed nucleus of the stria terminalis; dlPFC, dorsolateral prefrontal cortex; FDR, false discovery rate; FrO, frontal operculum; L, left; PAG, periaqueductal gray; t, Student’s *t*-test; WB, whole-brain corrected.

### Sustained activation is evident during both uncertain- and certain-threat anticipation

While useful, conventional hemodynamic-modeling approaches cannot resolve time-varying neural responses to anticipated threat encounters. To address this, we used a multiple-regression framework to transform the measured hemodynamic signal into a weighted linear combination of Onset, Sustained, and Phasic responses (**Figure 2a***, OSP Model*). Standard voxelwise GLMs were then used to identify regions showing sustained activation during the anticipation of uncertain and/or certain threat (FDR *q*<0.05, whole-brain corrected). Results closely resembled those yielded by conventional ‘boxcar’ analyses (**Figure 4**), with sustained activation evident throughout the canonical threat-anticipation circuit (<u>Shackman and</u> <u>Fox, 2021</u>)—including the dorsal amygdala—during the anticipation of both kinds of threat (**Figure 4**, *third column***; Supplementary Tables S6-S12,** https://osf.io/e2ngf). Despite this qualitative similarity, direct comparison of the two threats indicated that sustained responses were significantly stronger in canonical threat-related regions when the timing of threat encounters was uncertain (**Figure 4**, *fourth column;* FDR *q*<0.05, whole-brain corrected). While not a focus of our study, ancillary analyses indicated that uncertain-threat anticipation was associated with diminished sustained responses relative to uncertain safety (‘de-activation’) in a set of midline regions that broadly resembled the default mode network, including the frontal pole, rostral gyrus, pre- and post-central gyri, and precuneus (**Supplementary Table S10,** https://osf.io/e2ngf). The same pattern was evident in rostromedial and ventromedial sectors of the amygdala—including portions of the basal, cortical, and medial nuclei, and amygdalohippocampal transition area—consistent with prior work and with the known functional heterogeneity of this complex structure (<u>Hur et al., 2020b</u>; <u>Murty et al., 2022</u>; <u>Murty et al., 2023</u>; <u>Fox and</u> <u>Shackman, 2024</u>).

**Figure 4.**
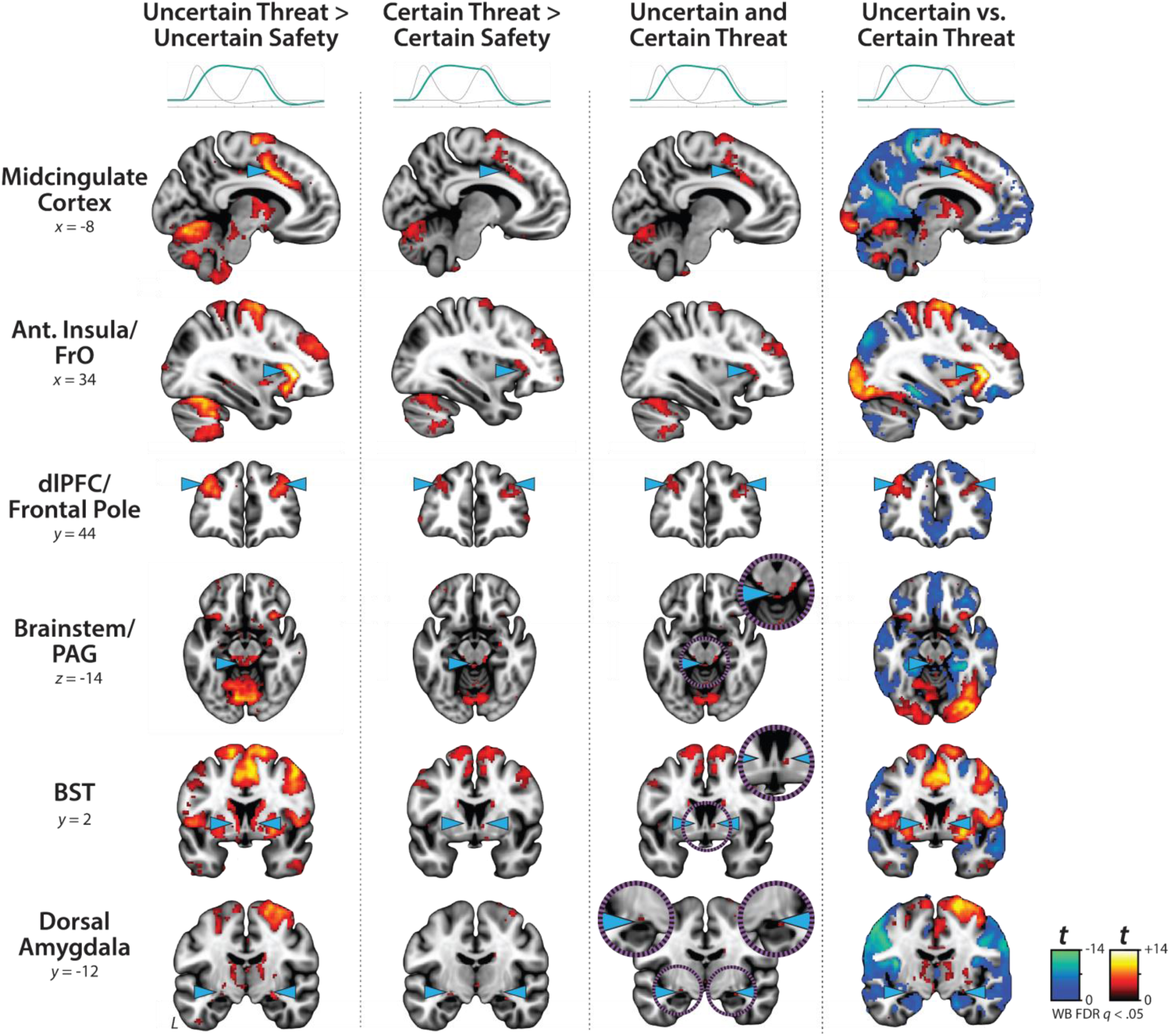
Sustained activation is evident during both uncertain- and certain-threat anticipation. Key regions showing evidence of sustained hemodynamic activity during the anticipation of temporally uncertain threat (*first column*) and certain threat (*second column*) relative to their respective control conditions (FDR *q*<0.05, whole-brain corrected). A minimum-conjunction of the two contrasts revealed colocalization throughout the threat-anticipation circuit (*third column*). Direct contrast of the two threat conditions showed that sustained signals are more pronounced in canonical threat-related regions during uncertain-threat anticipation (*fourth column*). Note: 4-mm smoothing kernel. Abbreviations—Ant., anterior; BST, bed nucleus of the stria terminalis; dlPFC, dorsolateral prefrontal cortex; FDR, false discovery rate; FrO, frontal operculum; L, left; PAG, periaqueductal gray; t, Student’s *t*-test; vs., versus; WB, whole-brain corrected.

### Phasic responses to certain-and-imminent threat are evident in the same regions that show sustained responses to uncertain threat, indicating a shared threat-anticipation circuit

Emotion theory and psychophysiological research both suggest that defensive responses surge in the moments just before certain threat encounters, but the underlying human neurobiology has remained unclear. Here we used a voxelwise GLM focused on the Phasic component of the OSP model—which is time-locked to the *offset* of the anticipation epoch, regardless of its duration—to identify regions showing significantly increased activation to certain-and-imminent threat (FDR *q*<0.05, whole-brain corrected; **Supplementary Tables S13-S19,** https://osf.io/e2ngf). Results revealed robust phasic responses during the terminal portion of certain-threat anticipation in every key region, including the BST (**Figure 5**, *left column*). ‘De-activation’ relative to certain-safety was minimal and largely confined to superficial cortical regions (e.g., superior parietal lobule; **Supplementary Table S17**). As expected, phasic responses were notably weaker (e.g., midcingulate) or nonsignificant (e.g., BST) during the corresponding moments of uncertain-threat anticipation, when the temporal imminence of threat is unknown to participants (**Figure 5**, *middle columns*). Inspection of these results suggests that the regions showing phasic responses to certain-and-imminent threat recapitulate those showing sustained responses during uncertain-threat anticipation (**Figure 4**). Consistent with this impression, a minimum-conjunction of the two thresholded contrasts revealed voxelwise overlap in all key regions (**Figure 5**, *right column*). The noteworthy degree of co-localization indicates that both kinds of threat recruit a shared threat-anticipation circuit that exhibits context-specific dynamics: sustained levels of heightened activation when threat encounters are uncertain and distal, and phasic surges in activation when threat encounters are certain and imminent. Importantly, because both conditions ultimately culminate in threat encounters (**Figure 1**), the absence of robust phasic responses during *uncertain*-threat indicates that phasic recruitment of the threat-anticipation circuit during *certain*-threat anticipation is not an artifact of reinforcer delivery (e.g., shock).

**Figure 5.**
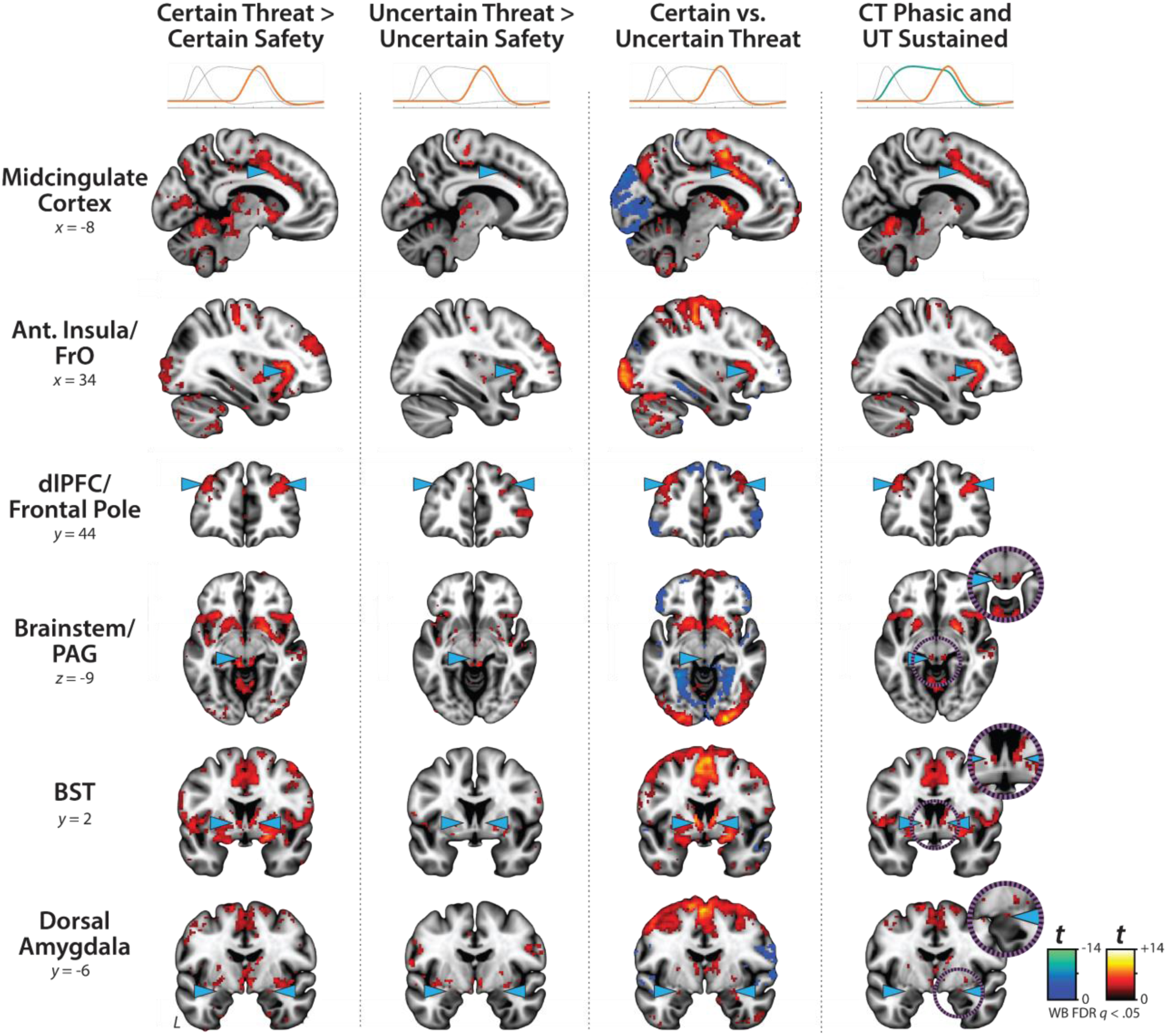
Phasic responses to certain-and-imminent threat are evident in the same regions that show sustained responses during the uncertain anticipation of threat. Regions showing significant phasic activation during the final seconds of certain-threat anticipation (*first column*) and uncertain-threat anticipation (*second column*) relative to their respective control conditions (FDR *q*<0.05, whole-brain corrected). Excepting the PAG, every key region showed significantly stronger phasic responses to certain threat (*third column*). Visual inspection suggests that the regions showing phasic responses to certain-and-imminent threat (*first column*) largely recapitulate the circuit showing sustained responses to uncertain-threat anticipation (Figure 4). Indeed, a minimum-conjunction of the two contrasts revealed voxelwise overlap in all regions (*fourth column*), suggesting that certain and uncertain threat are anatomically colocalized in a shared threat-anticipation circuit. Note: 4-mm smoothing kernel. Abbreviations—Ant., anterior; BST, bed nucleus of the stria terminalis; CT, certain-threat anticipation greater than certain-safety anticipation; dlPFC, dorsolateral prefrontal cortex; FDR, false discovery rate; FrO, frontal operculum; L, left; PAG, periaqueductal gray; t, Student’s *t*-test; UT, uncertain-threat anticipation greater than uncertain-safety anticipation; vs., versus; WB, whole-brain corrected.

While not a focus of the present report, it merits comment that exploratory analyses of the OSP Onset regressor (**Figure 2a**) revealed significant responses to both certain-*and* uncertain-threat anticipation in the right dorsal amygdala in the region of the basal and cortical nuclei, consistent with an attentional orienting or salience-related function (**Figure 6**; for detailed results see **Supplementary Tables S20-S24,** https://osf.io/e2ngf) (<u>Sokolov et al., 2002</u>; <u>Menon, 2015</u>).

**Figure 6.**
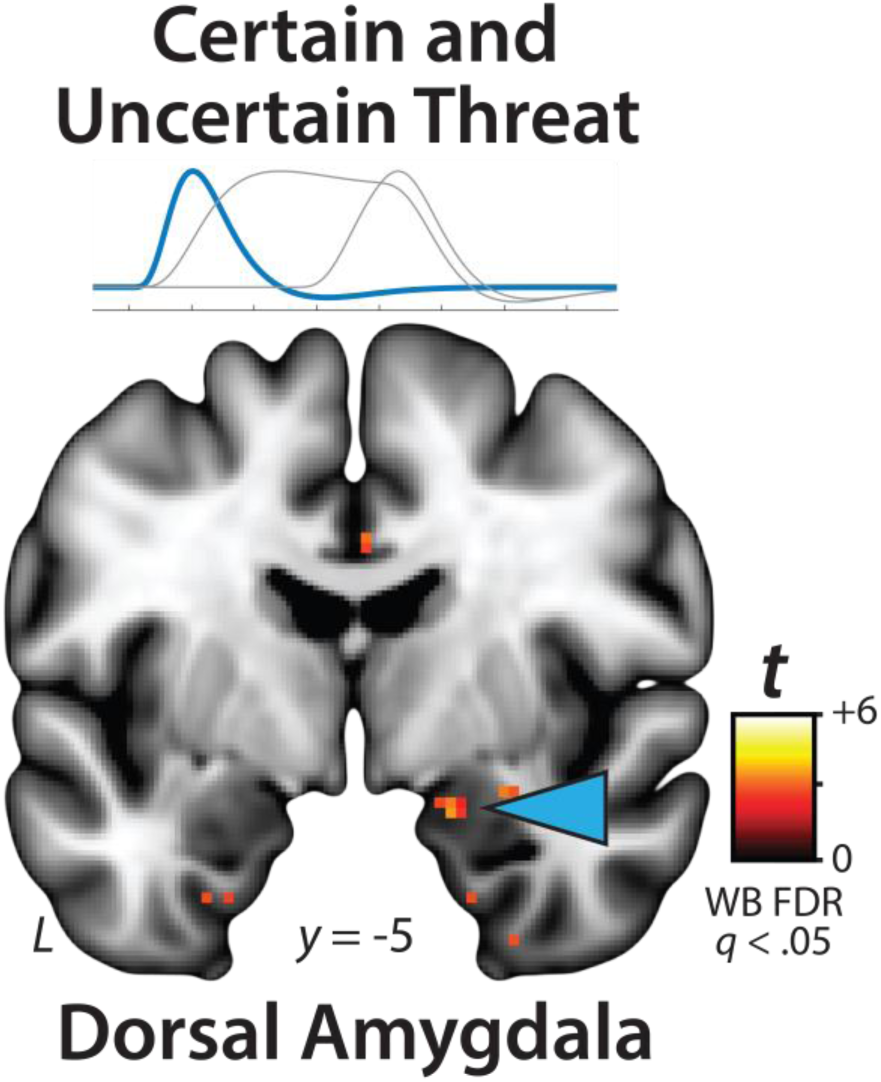
The dorsal amygdala is sensitive to the onset of the threat-anticipation epoch, independent of temporal certainty. Exploratory analyses of the OSP Onset regressor revealed significant responses to both certain- and uncertain-threat anticipation in the right dorsal amygdala in the region of the basal and cortical nuclei. Figure depicts the minimum conjunction (logical ‘AND’) of certain and uncertain threat relative to their respective control conditions (FDR *q*<0.05, whole-brain corrected). Note: 4-mm smoothing kernel. Abbreviations—FDR, False discovery rate; L, left; t, Student’s *t*-test; WB, whole-brain corrected.

### Phasic responses to acute threat reflect statistically significant surges in activation

The OSP Phasic results imply that activation significantly increased from the middle to the end of certain-threat anticipation (**Figure 5**, *first column*), and suggest that this increase is more pronounced for certain than uncertain threat (**Figure 5**, *third column*). Yet neither inference is licensed by the results, which are based on *within*-moment statistical contrasts (e.g., *OSP Phasic regressor:* certain vs. uncertain threat). The absence of *between*-moment tests reflects the fact that the partial-regression coefficients yielded by the OSP model do not allow straightforward interpretation of between-moment contrasts (**Figure 2**). To sidestep this, follow-up analyses capitalized on a second theory-driven hemodynamic model, which split the anticipation epoch into a sequence of short (6.25 s), non-overlapping rectangular functions or ‘blocks,’ each convolved with a canonical HRF (**Figure 2b**, *Convolved Blocks Model*). Although arbitrary in timing, this model yields activation estimates that are independent, inferentially intuitive, and statistically comparable across moments in time. A standard voxelwise GLM was then used to identify regions showing significant increases in activation during the late relative to the middle portion of certain-threat anticipation (FDR *q*<0.05, whole-brain corrected). Results revealed significant activation in every key region (including the BST), with the exception of the PAG (**Figure 7**, left column, and **Supplementary Tables S25-S30,** https://osf.io/e2ngf). A similar pattern was evident for the between-moments comparison of certain to uncertain threat, conceptually equivalent to testing the Threat-Certainty × Time interaction (**Figure 7**, *right column*). Taken together, these observations demonstrate that phasic responses to certain-and-imminent threat reflect statistically significant surges in activation in the seconds just before encountering threat (**Figure 7**, *left column*), and they show that this increase is significantly stronger for temporally certain encounters (**Figure 7**, *right column*).

**Figure 7.**
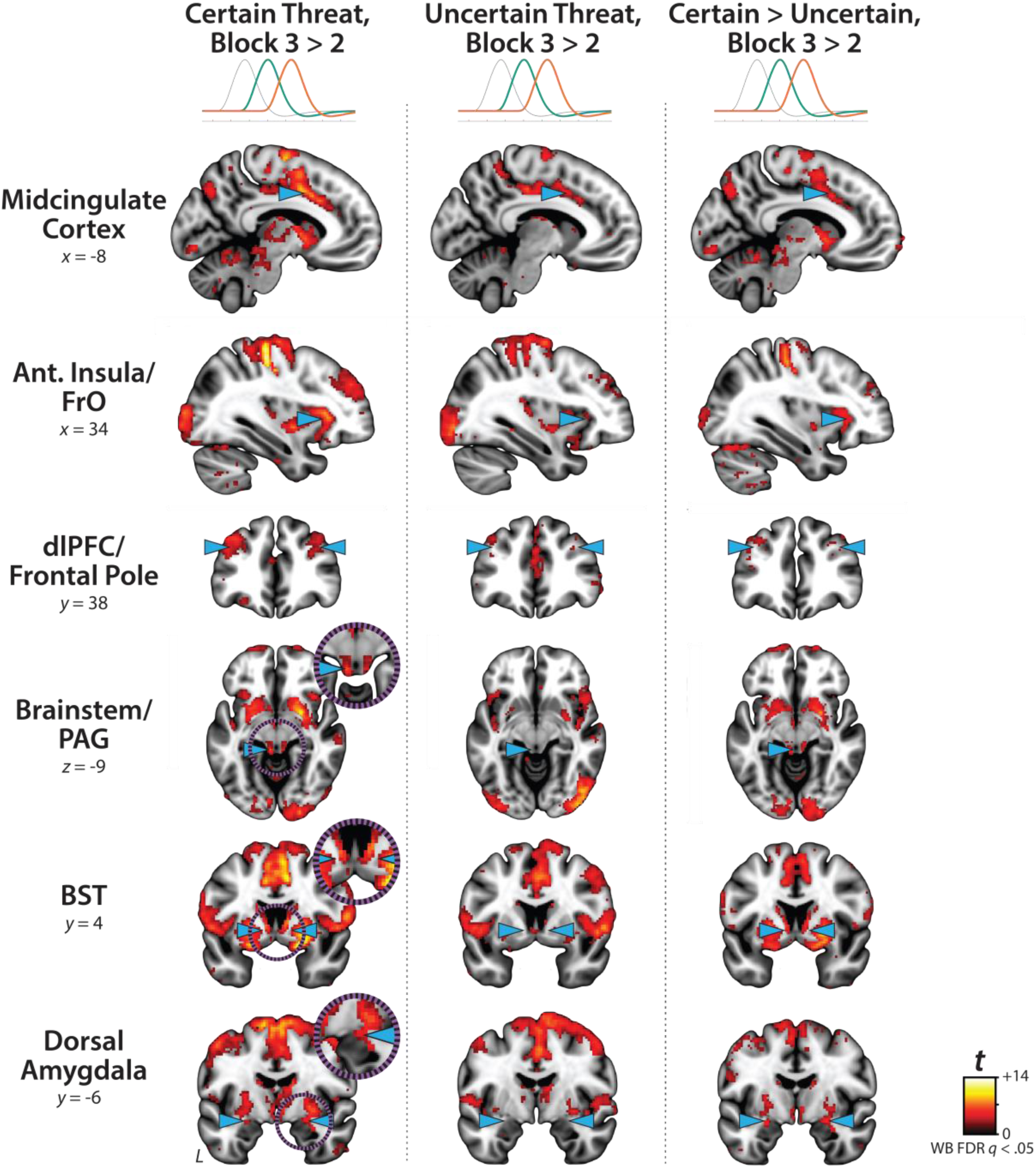
Statistically interrogating time-varying responses to certain-and-imminent threat. Regions showing significant surges in activation during the third compared to the second 6.25-s block for certain-threat anticipation (*left column*) and uncertain-threat anticipation (*middle column*; FDR *q*<0.05, whole-brain corrected). The *right column* depicts regions where activation surges are significantly stronger for certain threat. Results revealed significant surges in every key region except the PAG (*left column*), with a similar pattern evident for the between-moments comparison of certain to uncertain threat anticipation (i.e., the ‘difference of differences;’ *right column*). Note: 4-mm smoothing kernel. Abbreviations—Ant., anterior; BST, bed nucleus of the stria terminalis; dlPFC, dorsolateral prefrontal cortex; FDR, false discovery rate; FrO, frontal operculum; L, left; PAG, periaqueductal gray; t, Student’s *t*-test; WB, whole-brain corrected.

### Phasic responses to certain threat are stronger than sustained responses to uncertain threat in the BST and dorsal amygdala

The OSP results indicate that the BST, dorsal amygdala, PAG, MCC, AI/FrO, and dlPFC are recruited by anticipated threat encounters, irrespective of temporal certainty. All of these key regions show sustained levels of heightened activation when threat is uncertain and distal, and phasic surges in activation when threat is certain and imminent (**Figure 5**, *right column*). But such results do not address whether these regions differ in the strength of the two responses. Here we used activation estimates derived from the Convolved Blocks model—which enables meaningful between-moment comparisons—and a standard voxelwise GLM to compare reactivity to the second block (6.25-12.5 s) of uncertain threat, an indicator of sustained activation, to the final block of certain threat (12.5-18.75 s), an indicator of phasic surges in activation (FDR *q*<0.05, whole-brain corrected). As shown in **Figure 8**, results revealed significantly stronger activation during the final moments of certain-threat anticipation in the BST and dorsal amygdala in broad agreement with prior work (<u>Hur et al., 2020b</u>). Effects in other key regions were scattered or nil, and none of the key cortical or subcortical regions showed stronger responses to uncertain-threat anticipation (**Supplementary Tables S31-S32,** https://osf.io/e2ngf).

**Figure 8.**
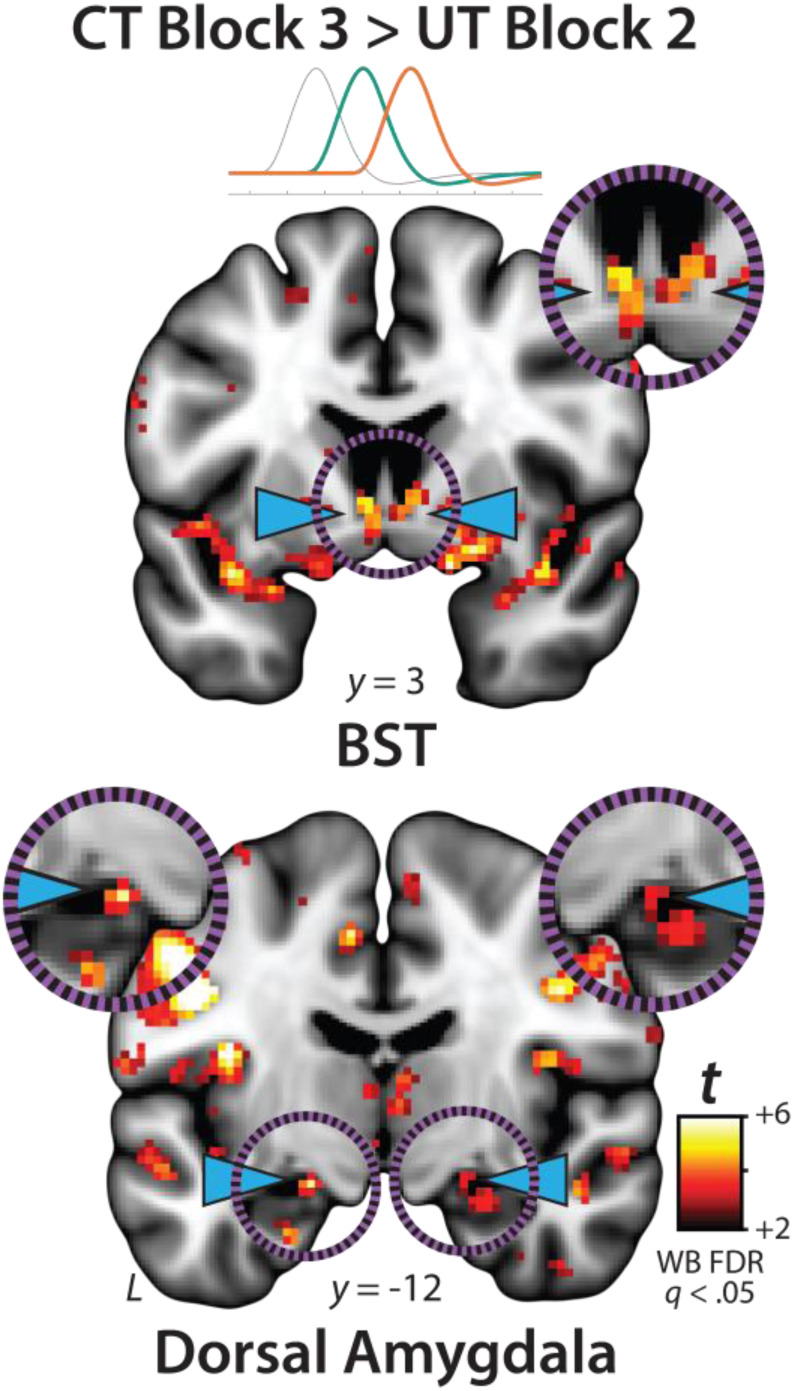
Phasic responses to certain threat are stronger than sustained responses to uncertain threat in the BST and dorsal amygdala. Regions showing significantly stronger activation during the final 6.25 s block of certain-threat anticipation compared to the second block of uncertain-threat anticipation, each relative to their respective control conditions (FDR *q*<0.05, whole-brain corrected). Note: 4-mm smoothing kernel. Abbreviations—BST, bed nucleus of the stria terminalis; FDR, false discovery rate; L, left; t, Student’s *t*-test; WB, whole-brain corrected.

### The BST and Ce show statistically indistinguishable neural dynamics

The present approach also afforded a well-powered opportunity to revisit the functional architecture of the human EAc, a macrocircuit encompassing the dorsal amygdala in the region of the central nucleus (Ce) and the neighboring bed nucleus of the stria terminalis (BST) (<u>Fox et al., 2015a</u>). There is widespread consensus that the EAc plays a critical role in assembling defensive responses to a broad spectrum of threats and contributes to the etiology of emotional illness (<u>Davis et al., 2010</u>; <u>Grupe and Nitschke, 2013</u>; <u>Tovote et al., 2015</u>; <u>Hur et al., 2019</u>; <u>Shackman and Fox, 2021</u>; <u>Moscarello and Penzo, 2022</u>; <u>Tseng et al.,</u> <u>2023</u>; <u>Fox and Shackman, 2024</u>; <u>Akiki et al., 2025</u>). Yet confusion persists about the respective contributions of its two major subdivisions (<u>Daniel-Watanabe and Fletcher, 2022</u>; <u>Shackman et al., 2024</u>).

Inspired by an earlier wave of loss-of-function studies in rats (<u>Davis, 2006</u>), it is widely believed that these regions are dissociable, with the Ce mediating phasic responses to certain-and-imminent harm and the BST mediating sustained responses to uncertain-or-remote danger (<u>Grillon, 2008</u>; <u>Öhman, 2008</u>; <u>Avery et al.,</u> <u>2016</u>; <u>LeDoux and Pine, 2016</u>; <u>Klumpers et al., 2017</u>). This hypothesized double-dissociation has even been enshrined in the National Institute of Mental Health’s (NIMH) influential Research Domain Criteria (RDoC) framework as Acute Threat (*“fear”*) and Potential Threat (*“anxiety”*) (<u>NIMH, 2011</u>, <u>2020b</u>, <u>a</u>). Yet a growing body of evidence motivates the competing hypothesis that the Ce and BST both play a role in organizing phasic and sustained responses to threat (<u>Gungor and Paré, 2016</u>; <u>Fox and Shackman, 2019</u>; <u>Hur et al.,</u> <u>2020b</u>; <u>Shackman and Fox, 2021</u>; <u>Moscarello and Penzo, 2022</u>; <u>Shackman et al., 2024</u>). Likewise, the present results demonstrate that the **(a)** dorsal amygdala (in the region of the Ce) shows sustained responses to uncertain-threat anticipation (**Figure 4**, *first column*) and **(b)** the BST shows phasic responses during the final moments of certain-threat anticipation (**Figure 7**, *left column*). Because conventional voxelwise analyses do not permit inferences about between-region differences in activation, we used *a priori* probabilistic anatomical regions of interest (ROIs) to rigorously assess these competing predictions (**Figure 9a**). This approach has the added advantage of providing statistically unbiased effect-size estimates (<u>Poldrack et al., 2017</u>), in contrast to earlier work focused on functionally defined ROIs (<u>Hur et</u> <u>al., 2020b</u>). To maximize resolution, mean activation was computed for bilateral BST and Ce ROIs using spatially unsmoothed data. Hypothesis testing focused on ROI responses to certain- and uncertain-threat anticipation, relative to their respective second-level reference conditions (e.g., uncertain threat vs. uncertain safety). To enable between-moment comparisons, activation estimates were derived using the first three blocks of the Convolved Blocks model (**Figure 2b**).

**Figure 9.**
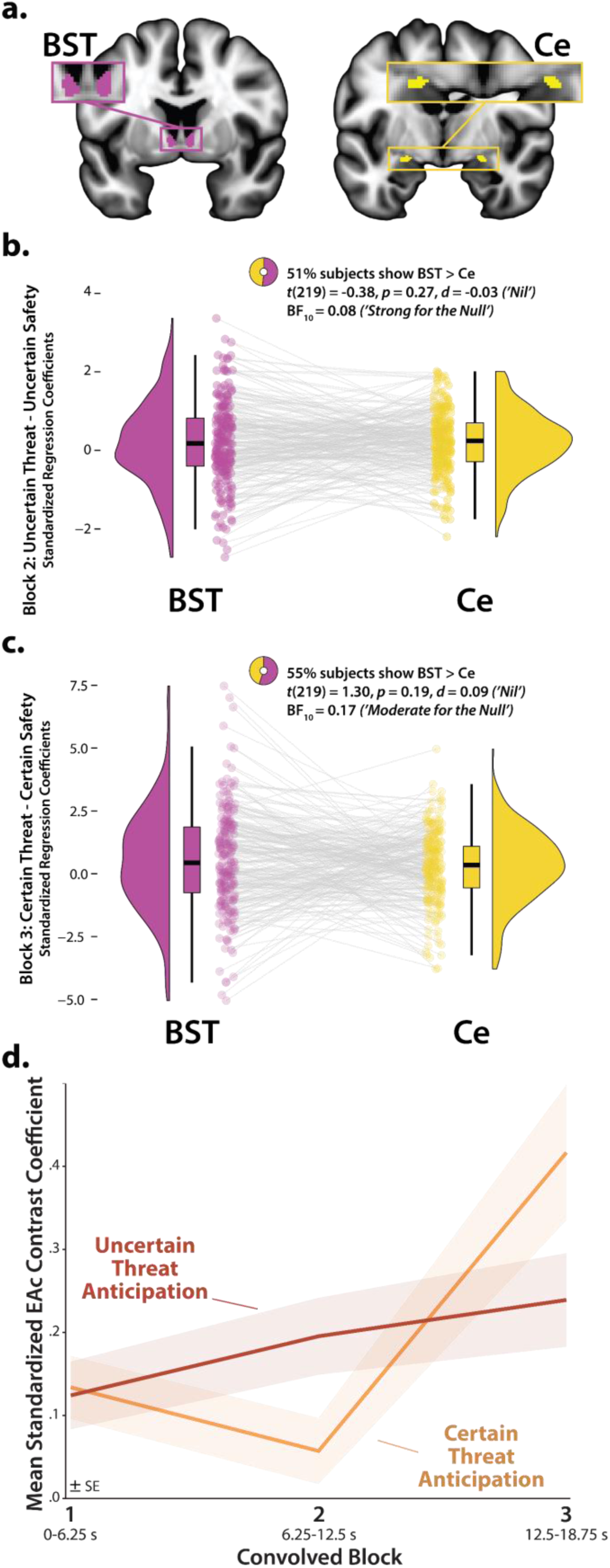
The BST and Ce show statistically indistinguishable neural dynamics. **a. *Probabilistic EAc anatomical ROIs.*** The BST (*magenta*) and Ce (*yellow*) ROIs. **b. *Uncertain-threat anticipation, second convolved block.*** The BST and Ce show negligible differences in activation during the second block (6.25-12.5 s) of uncertain-threat anticipation (a proxy for sustained activation). **c. *Certain-threat anticipation, third convolved block.*** The BST and Ce show negligible differences during the final block (12.5-18.75 s) of certain-threat anticipation (a proxy for phasic surges in activation). d. *The EAc shows context-dependent dynamics.* In aggregate, the EAc evinced a marginally significant linear increase in EAc activation during uncertain-threat anticipation (*red*; *p*=0.06) and a pronounced quadratic (‘V-shaped’) trend during certain-threat anticipation (*orange*; *p*=0.001). Colored envelopes depict the SE. Note. Raincloud plots indicate the medians (*horizontal lines*), interquartile ranges (*boxes*), and smoothed density distributions. Whiskers depict 1.5× the interquartile range. Colored dots connected by gray lines indicate mean regional activation for each participant. Note: No spatial smoothing kernel was employed for ROI analyses. Abbreviations—BF, Bayes’ factor; BST, bed nucleus of the stria terminalis; Ce, central nucleus of the amygdala; *d*, Cohen’s *dz*; EAc, central extended amygdala; SE, standard error of the mean; *t*, Student’s *t*-test.

As a precursor to hypothesis testing, we used one-sample Student’s *t*-tests to confirm that the BST and Ce are nominally engaged by anticipated threat (*p*<0.05, uncorrected). With one exception, results revealed uniformly significant activation (*t*(219)>2.08, *p*<0.04). The Ce did not show significant evidence of activation during the middle third of certain-threat anticipation (*t*(219)=-0.40, *p*=0.69). On balance, these observations indicate that both EAc subdivisions are sensitive to anticipated threat, irrespective of the temporal certainty of encounters.

Next we used a standard 2 (*Region:* BST, Ce) × 2 (*Threat Certainty:* Certain, Uncertain) × 3 (*Block:* Early, Middle, Late) repeated-measures GLM to formally test the double-dissociation hypothesis embodied in RDoC and other ‘strict-segregation’ models. None of the Regional effects were significant (*p*>0.13), including the conceptually critical Region × Threat Certainty × Block interaction (*F*(2,438)=0.72, *p*=0.46). Consistent with this, the BST and Ce showed negligible differences in activation during the second block of uncertain-threat, an indicator of sustained activation, or the final block of certain-threat, an indicator of phasic surges in activation (|*t*|(219)<1.31, *p*>0.18; **Figure 9b-c** and **Supplementary Figure S1,** https://osf.io/e2ngf). Frequentist effects were in the nil range (|*d*|=0.03-0.09).

Of course traditional null-hypothesis tests cannot addresss whether the BST and Ce show equivalent responses to certain- and uncertain-threat anticipation (<u>Wagenmakers et al., 2018</u>). Here we used Bayes Factor (*BF_10_*) to quantify the relative strength of the evidence for and against regional equivalence. The Bayesian approach provides well-established benchmarks for interpreting effect sizes and sidesteps the need to arbitrarily choose what constitutes a ‘statistically indistinguishable’ difference (<u>Wagenmakers et</u> <u>al., 2018</u>; <u>van Doorn et al., 2021</u>; <u>Bo et al., 2024</u>), unlike the equivalence tests used in prior work (<u>Hur et al.,</u> <u>^2^020^b^</u>). Bayesian results signaled moderate-to-strong evidence for the null (*BF_10_*=0.08-0.17) during the conceptually crucial second block of uncertain-threat anticipation and third block of certain-threat anticipation. Put another way, from a Bayesian perspective, the null hypothesis of equivalent regional responses is ∼6-13 times more likely than the alternative. Descriptively, participants were just as likely as not to show the RDoC-predicted regional differences; for example, 51% showed stronger BST-than-Ce activation during the second block of uncertain-threat anticipation (*H_0_*=50%).

To clarify interpretation of these results, we explored potential differences in BST and Ce activation using estimates derived from the OSP model. Because the OSP model does not permit meaningful between-moment contrasts, we separately examined sustained responses to uncertain-threat and phasic responses to certain-threat anticipation, each relative to their second-level reference conditions (e.g., certain threat vs. certain safety). Consistent with the Convolved Blocks results (**Figure 9**), neither difference was statistically significant (*t*(219)<1.84, *p*>0.06), frequentist effects were in the small-to-nil range (*d*=0.06-0.12), and Bayesian results signaled moderate-to-weak evidence for the null (*BF_10_*=0.12-0.39). From the vantage point of the OSP model, the null hypothesis of equivalent BST and Ce responses is ∼2.5-8.5 times more likely than the alternative. In sum, we uncovered no evidence for the popular double-dissociation hypothesis, despite being powered to detect small regional differences in activation (Cohen’s *d*=0.19; see the Method section for details).

### The central extended amygdala (EAc) exhibits context-dependent neural dynamics

Analyses of activation estimates derived from the convolved blocks model did, however, provide evidence that the EAc in aggregate—averaged across the BST and Ce—shows context-dependent neural dynamics, as indexed by significant Block and Threat-Certainty × Block effects (*F*(2,438)>5.18, *p*<0.01). As shown in **Figure 9d**, polynomial-trend analyses revealed a marginally significant linear increase in EAc activation during uncertain-threat anticipation (*Linear: F*(1,219)=3.58, *p*=0.06; *Quadratic: F*(1,219)=0.05, *p*=0.82). In contrast, the EAc showed a pronounced quadratic (‘V-shaped’) trend during certain-threat anticipation, manifesting as a dip in the middle third, followed by a surge of activation in the final third, when the threat encounter was most imminent (*Linear: F*(1,219)=11.30, *p*<0.001; *Quadratic: F*(1,219)=10.38, *p*=0.001).

### Brain-signature estimates of subjective distress show the same pattern of context-dependent dynamics

It is tempting to interpret our neuroimaging results in terms of conscious feelings—to infer that participants experience a sustained state of heightened anxiety when the timing of threat encounters is uncertain and a surge of fear in the seconds just before certain encounters. Yet standard fMRI analyses cannot address the momentary dynamics of threat-evoked distress, a limitation shared with other behavioral and psychophysiological measures, and with mechanistic work in animals (<u>Poldrack, 2011</u>; <u>LeDoux, 2014</u>; <u>Grogans et al., 2023</u>). Likewise, more intensive continuous or intermittent ratings have the potential to fundamentally alter momentary emotional experience (<u>Ruef and Levenson, 2007</u>; <u>Lieberman,</u> <u>2018</u>). Here we used activation estimates derived from the Convolved Blocks model and an independently trained and validated multivoxel pattern or ‘signature’ of subjective negative affect to covertly probe the momentary dynamics of threat-evoked distress for the first time (**Figure 10a**) (<u>Čeko et al., 2022</u>; <u>Peelen</u> <u>and Downing, 2023</u>). Prior work demonstrates that this signature is a sensitive indicator of distress elicited by a variety of noxious experiences—including thermal and mechanical pain, unpleasant photographs, and aversive auditory stimuli—but is unrelated to the intensity of feelings triggered by positive stimuli, showing specificity (<u>Čeko et al., 2022</u>). Conceptually similar multivoxel pattern analysis (MVPA) approaches have been successfully used in other areas of the cognitive neurosciences; for example, to unobtrusively decode the contents of working memory or the focus of selective attention without disrupting on-going performance (<u>Peelen and Downing, 2023</u>).

**Figure 10.**
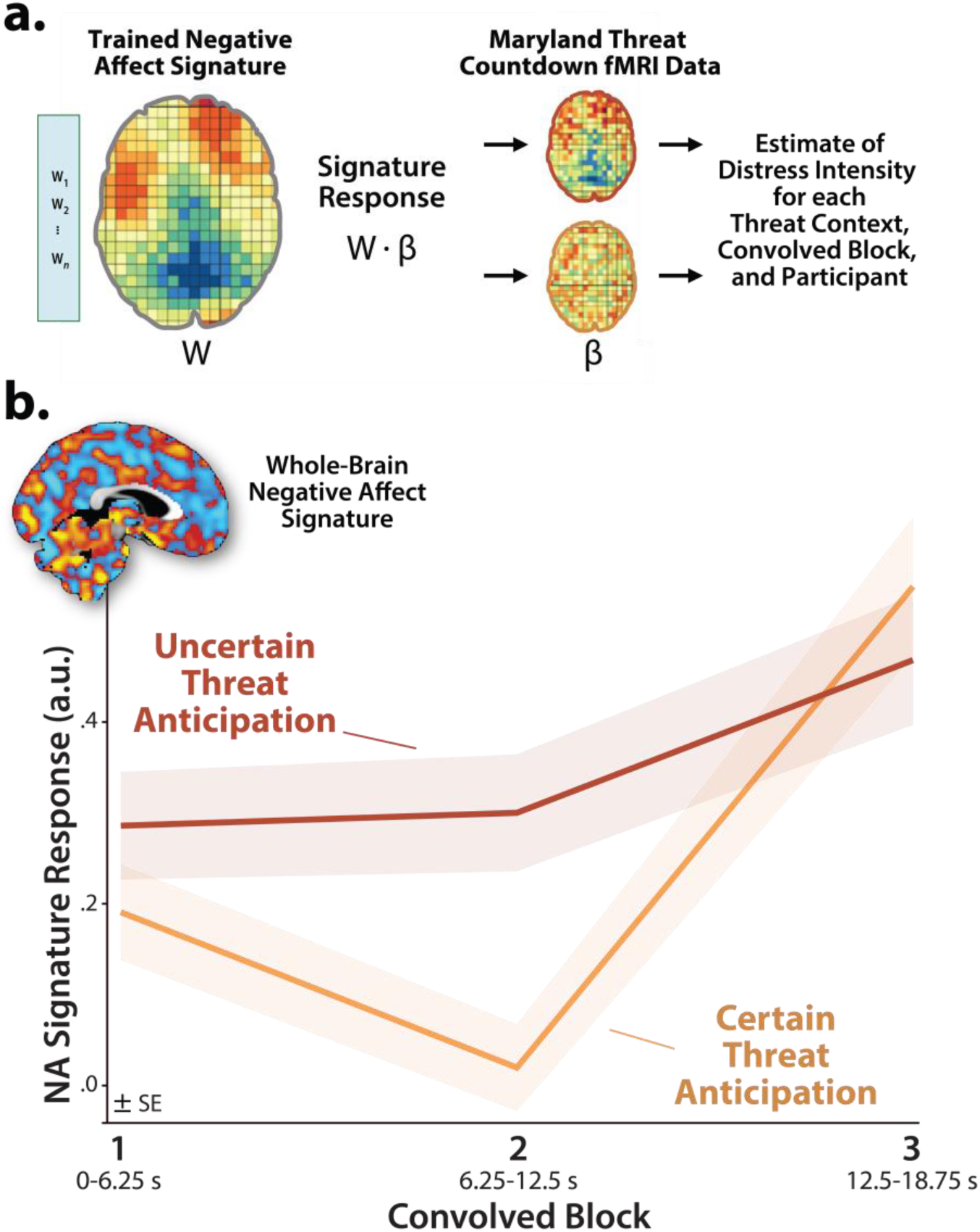
Using a multivoxel brain signature to covertly estimate momentary fluctuations in threat-elicited distress. **a. An independently trained and validated whole-brain signature of subjective negative affect was used to estimate threat-evoked distress.** Čeko, Wager, and colleagues used machine-learning to develop a whole-brain ‘signature’—a pattern of voxelwise weights (*w*)—that is predictive of negative affect intensity in unseen data across a variety of noxious stimuli (<u>Ceko et al., 2022</u>). In effect, the signature treats each voxel as a weighted source of information and the overall pattern as a collective ‘best guess.’ Computing the dot-product (⦁) between the pattern of weights (*W*) and voxelwise activation estimates (*β*) derived for the present sample using the Convolved Blocks model generates a signature response—a probabilistic estimate of distress intensity—for every combination of threat certainty, block, and participant. This made it possible to covertly estimate moment-by-moment fluctuations in threat-elicited distress and test whether distress dynamics are sensitive to the temporal certainty of threat encounters. **b. Subjective distress shows context-dependent dynamics.** The estimated intensity of distress was significantly greater, on average, when anticipating uncertain encounters with threat (*p*=0.004). Significant linear and quadratic polynomial trends were evident for both kinds of threat anticipation (*p*<0.03), but the V-shaped quadratic effect was more than an order of magnitude stronger for certain threat (*Certain: pη^2^*=0.31; *Uncertain: pη^2^*=0.02). *Inset depicts the whole-brain multivoxel signature of negative affect.* Hot and cool colors indicate positive and negative signature weights, respectively. Colored envelopes depict the SE. Note: 4-mm smoothing kernel. Abbreviations—fMRI, functional magnetic resonance imaging; NA, negative affect; SE, standard error of the mean.

As a first step, we used one-sample Student’s *t*-tests to confirm that the whole-brain signature is sensitive to threat anticipation (*p*<0.05, uncorrected). With one exception, results revealed robust signature responses, signaling more intense negative affect (*t*(219)>6.67, *p*<0.001). The signature did not show evidence of ‘sustained’ distress in the middle third of certain-threat anticipation (*t*(219)=0.62, *p*=0.54). Taken with prior work in this sample and others demonstrating that the MTC paradigm triggers robust distress and arousal (<u>Kim et al., 2023</u>; <u>Grogans et al., 2024</u>), these observations suggest that the signature is a valid index of threat-evoked anticipatory distress.

Next we used a standard 2 (*Threat Certainty:* Certain, Uncertain) × 3 (*Block:* Early, Middle, Late) GLM to estimate moment-by-moment fluctuations in probable distress across the two threat contexts. Results revealed significantly greater distress estimates, on average, when anticipating temporally uncertain threat encounters (*Threat Certainty: F*(1,219)=8.47, *p*=0.004; **Figure 10b**), consistent with prior work focused on retrospective ratings (<u>Kim et al., 2023</u>; <u>Grogans et al., 2024</u>). The Block effect and Threat Certainty × Block interaction were also significant (*F*(2,438)>19.34, *p*<0.001). Although significant linear and quadratic polynomial trends were evident for both kinds of anticipated threat (*F*(1,219)>5.00, *p*<0.03), the V-shaped (‘surge-trough-surge’) quadratic effect was more than an order of magnitude stronger when anticipating certain threat encounters (*Certain: pη^2^*=0.31, *p*=4.71 × 10^-19^; *Uncertain: pη^2^*=0.02, *p*=0.02; **Figure 10b**). In combination with the one-sample *t*-test results (see above), this suggests that temporally uncertain-threat anticipation elicits a sustained state of heightened negative affect, whereas certain threat is associated with more complex distress dynamics, with negligible distress evident in the middle period and a phasic surge when threat is most acute.

## DISCUSSION

Since the time of Freud, the distinction between fear and anxiety has been a feature of prominent models of emotion and emotional illness (<u>Freud et al., 1959</u>). Despite the enormous significance of threat-elicited emotions for public health, the neural systems underlying phasic responses to acute danger and sustained responses to uncertain harm are contentious (<u>Daniel-Watanabe and Fletcher, 2022</u>; <u>Shackman et al., 2024</u>).

Some say that *“fear”* and *“anxiety”* are phenomenologically distinct states mediated by anatomically dissociable circuits (<u>Avery et al., 2016</u>; <u>LeDoux and Pine, 2016</u>), whereas others suggest that they are more biologically alike than different (<u>Fox and Shackman, 2019</u>; <u>Hur et al., 2020b</u>). Leveraging a relatively large and racially diverse sample, translationally relevant fMRI paradigm, and theory-driven hemodynamic modeling approach, our results demonstrate that the anticipation of temporally certain and uncertain threat recruits an overlapping cortico-subcortical circuit, with co-localization evident in several previously implicated regions, including the BST, dorsal amygdala, PAG, MCC, AI/FrO, and dlPFC (**Figures 3** and **5**). This shared threat-anticipation circuit exhibits context-specific dynamics, evincing sustained levels of heightened activation when threat encounters are uncertain and distal (**Figure 4**), and phasic surges in activation when encounters are certain and imminent (**Figures 5** and **7**).

Among the regions highlighted here, the BST and Ce play a central role in prominent neurobiological models of fear and anxiety. Yet their precise contributions remain debatable (<u>Blanchard and Canteras,</u> <u>2024</u>; <u>Shackman et al., 2024</u>). Our results show that both regions exhibit activation dynamics that run counter to popular double-dissociation models, with the dorsal amygdala (in the region of the Ce) showing sustained responses to uncertain-and-distal threat and the BST showing phasic responses to acute threat (**Figures 4**-**5** and **7**-**8**). Leveraging anatomical ROIs, our results demonstrate that the BST and Ce exhibit statistically indistinguishable responses to anticipated threat—with frequentist effects in the nil range and Bayesian effects indicating moderate-to-strong evidence for the null hypothesis—reinforcing the possibility that they make broadly similar contributions to human fear and anxiety (**Figure 9**).

Our observations do not mean that the BST and Ce are functionally interchangeable. Among monkeys, BST activity is more closely related to the heritable variation (‘nature’) in trait anxiety, whereas Ce activity is more closely related to the variation in trait anxiety that is explained by differences in early-life experience (‘nurture’) (<u>Fox et al., 2015b</u>). Among humans, variation in neuroticism/negative emotionality—a key dispositional risk factor for emotional disorders—is selectively associated with heightened BST reactivity to uncertain threat (<u>Grogans et al., 2024</u>). Understanding the breadth and nature of these regional differences is a key avenue for future research.

Our voxelwise results show that the dorsal amygdala response to anticipated threat is sparse, at least when compared to popular emotional face paradigms. This was not unexpected. First, the amygdala is neither a natural kind nor a singular anatomical unit; it is a heterogeneous collection of at least 12 nuclei; and converging lines of mechanistic and imaging evidence point to the special importance of the dorsal amygdala in the region of the Ce (<u>Fox and Shackman, 2024</u>). In humans, Ce represents ∼3% of total amygdala volume (<u>Avino et al., 2018</u>). Second, the extent and location of the dorsal amygdala clusters reported here is consistent with the results of well-powered studies of certain- and uncertain-threat anticipation (<u>Hur et al., 2020b</u>; <u>Sjouwerman et al., 2020</u>; <u>Kim et al., 2023</u>; <u>Grogans et al., 2024</u>). The use of a smaller-than-typical 4-mm smoothing kernel would be expected to further reduce cluster extent. Third, confidence in our voxelwise results is reinforced by the ROI analyses, which leveraged spatially unsmoothed data to maximize resolution and inferential sharpness. In sum, our results are broadly aligned with anatomy, theory, and existing neuroimaging evidence.

Pathological fear and anxiety is largely defined, diagnosed, and treated on the basis of subjective symptoms, and for many scientists and laypeople, conscious feelings are the defining feature of these emotions (<u>Grogans et al., 2023</u>). Yet standard fMRI analyses, like animal models, do not permit strong inferences about conscious feelings. Here we used an independently trained and validated brain signature to covertly decode the momentary dynamics of threat-evoked distress for the first time. Results indicated that uncertain-threat anticipation is associated with a sustained state of elevated negative affect, whereas certain-threat anticipation elicits more complex dynamics, with a phasic surge of distress evident just before threat encounters (**Figure 10**). These observations begin to address calls for a tighter integration of subjective and neurobiological measures of human fear and anxiety (<u>Daniel-Watanabe and Fletcher,</u> <u>2022</u>; <u>Domschke, 2022</u>), and they reinforce the conclusion that human fear and anxiety, while associated with distinct functional dynamics, reflect the operation of a shared threat-anticipation circuit.

The core threat-anticipation circuit encompasses subcortical regions, such as the BST and Ce, that are critical for assembling defensive responses to anticipated threat in animals (<u>Fox and Shackman, 2019</u>; <u>Moscarello and Penzo, 2022</u>). But it also includes frontocortical regions—including the MCC, AI/FrO, and dlPFC/FP—that have received less empirical attention and are challenging or impossible to study in rodents (<u>Roberts and Mulvihill, 2024</u>). These regions have traditionally been associated with the controlled processing and regulation of emotion and cognition (<u>Shackman et al., 2011</u>; <u>Bo et al., 2024</u>) and more recently implicated in the conscious experience of emotion (<u>LeDoux, 2020</u>). Our findings extend past work focused on descriptive hemodynamic modeling approaches in smaller samples, and dovetail with meta-analytic evidence that Pavlovian fear-conditioning tasks (the prototypical experimental model of certain- and-imminent threat) and instructed threat-of-shock tasks (the prototypical experimental model of uncertain threat) recruit strongly overlapping cortico-subcortical networks in humans, including the BST (<u>Shackman and Fox, 2021</u>).

Our results provide a keyboard of regions and activation-dynamics, setting the stage for identifying the functional-neuroanatomical combinations most relevant to the development of pathological fear and anxiety and to the efficacy of established therapeutics. Consider the widely prescribed anxiolytic, diazepam. As yet, the neurodynamic mechanisms that underlie the blockade of threat-elicited distress by diazepam and other benzodiazepines remain unsettled. Does anxiolysis primarily reflect the dampening of sustained responses to uncertain threat in the Ce, as implied by recent work in mice (<u>Griessner et al., 2021</u>), or widespread changes across multiple activation metrics, as implied by our signature results?

Our findings add to a growing body of evidence that the BST and Ce both play a role in governing defensive responses to a wide variety of threats, both certain and uncertain (<u>Fox and Shackman, 2019</u>; <u>Shackman et</u> <u>al., 2024</u>). The two regions are characterized by similar patterns of anatomical connectivity, cellular composition, neurochemistry, and gene expression (<u>Fox et al., 2015a</u>). Both are poised to trigger behavioral, psychophysiological, and neuroendocrine responses to threat via dense projections to downstream effector regions (<u>Davis and Whalen, 2001</u>). Both are recruited by a broad spectrum of aversive and potentially threat-relevant stimuli (<u>Fox and Shackman, 2019</u>), and both are implicated in pathological fear and anxiety (<u>Shackman and Fox, 2021</u>). Perturbation studies in rodents demonstrate that microcircuits within and between the Ce and BST are critical for orchestrating defensive responses to both acute and uncertain threats (<u>Lange et al., 2017</u>; <u>Zelikowsky et al., 2018</u>; <u>Fox and Shackman, 2019</u>; <u>Pomrenze et al., 2019a</u>; <u>Pomrenze et al., 2019b</u>; <u>Ressler et al., 2020</u>; <u>Chen et al., 2022</u>; <u>Moscarello and Penzo,</u> <u>2022</u>; <u>Ren et al., 2022</u>; <u>Zhu et al., 2024</u>). While our understanding remains incomplete, these observations underscore the need to reformulate RDoC and other models that imply a strict segregation of certain and uncertain threat processing in the EAc.

A key challenge for the future will be to determine whether our conclusions generalize to more demographically representative samples, other types of threat (e.g., social), and other kinds of uncertainty (e.g., probability, risk, ambiguity). Although our theory-driven hemodynamic modeling approach has clear advantages over traditional ‘boxcars,’ less-constrained models promise to provide more nuanced information about threat-related neural dynamics (<u>Gonzalez-Castillo et al., 2012</u>; <u>Gonzalez-Castillo et al.,</u> <u>2015</u>). Moving forward, an enhanced emphasis on computationally tractable paradigms has the potential to address fundamental questions about the function of the regions highlighted by our results and foster a common mathematical framework (*‘lingua franca’*) for integrating research across assays, read-outs, and species (<u>Drzewiecki and Fox, 2024</u>). The Ce and BST are complex and can be subdivided into multiple subdivisions, each containing intermingled cell types with distinct, even opposing functional roles (e.g., anxiogenic vs. anxiolytic) (<u>Fox and Shackman, 2019</u>, <u>2024</u>). Animal models will be critical for generating testable hypotheses about the molecules, cell types, and microcircuits that govern activation dynamics in human-neuroimaging studies.

In conclusion, the neural circuits recruited by temporally uncertain and certain threat are not categorically different, at least when viewed through the macroscopic lens of fMRI hemodynamics. We see evidence of anatomical colocalization—*not* segregation—in the EAc and key frontocortical regions. This shared threat-anticipation circuit shows persistently elevated activation when anticipating uncertain threat encounters and acute bursts of activation in the moments before certain encounters. Subjective distress shows parallel dynamics. These observations provide a neurobiologically grounded framework for conceptualizing fear and anxiety and lay the groundwork for future prospective-longitudinal, clinical, computational, and mechanistic work.

## Supporting information

Supplementary Results

## ACKNOWLEDGEMENTS

We acknowledge assistance and critical feedback from A. Antonacci, L. Friedman, J. Furcolo, C. Grubb, J. Hassani, R. Hum, C. Kaplan, J. Kuang, M. Kuhn, C. Lejuez, D. Limon, B. Nacewicz, L. Pessoa, S. Rose, J. Swayambunathan, A. Vogel, B. Winters, J. Wedlock, two anonymous peer reviewers, members of the Affective and Translational Neuroscience Laboratory, the staff of the Maryland Neuroimaging Center, the Office of the Registrar at the University of Maryland. This work was partially supported by the California National Primate Center; National Institutes of Health (AA030042, AA031261, DA040717, MH107444, MH121409, MH121735, MH128336, MH129851, OD011107, MH131264, MH126426); National Research Foundation of Korea (2021R1F1A1063385 and 2021S1A5A2A03070229); University of California, Davis; University of Maryland; and Yonsei Signature Research Cluster Program (2021-22-0005). Authors declare no conflicts of interest.

## AUTHOR CONTRIBUTIONS

A.J.S., K.A.D., and J.F.S. designed the overall study. J.F.S. envisioned the present project. J.F.S. and A.J.S. developed and optimized the imaging paradigm. K.A.D. managed data collection and study administration. K.A.D., J.F.S., A.S.A, S.I., and R.M.T. collected data. J.F.S. and M.K. developed data processing and analytic software for imaging analyses. J.F.S., J.H., H.C.K., and R.M.T. processed imaging data. J.F.S., B.R.C., and A.J.S. analyzed imaging data. J.F.S., A.J.S., and B.R.C. developed the analytic strategy. B.R.C., P.R.D., J.F.S., A.S.F., and A.J.S. interpreted data. B.R.C., S.E.G., J.F.S., and A.J.S. wrote the paper. P.R.D., A.J.S., B.R.C., J.F.S., and Z.S.S. created figures and tables. A.J.S. funded and supervised all aspects of the study. All authors contributed to reviewing and revising the paper and approved the final version.

## RESOURCE SHARING

Raw data are available at the National Institute of Mental Health Data Archive (https://nda.nih.gov/edit_collection.html?id=2447). Neuroimaging maps are available at NeuroVault (https://neurovault.org/collections/15274). Task materials, statistical code, de-identified processed data, and supplementary neuroimaging cluster tables are available at OSF (https://osf.io/e2ngf). The negative affect brain signature is available at Github (https://github.com/canlab/Neuroimaging_Pattern_Masks/tree/master/Multivariate_signature_patterns/2021_Ceko_MPA2_multiaversive).

