## Supplementary Results for "A shared threat-anticipation circuit is dynamically engaged at different moments by certain and uncertain threat"

### Supplementary Figure and Tables

Brian R. Cornwell[1]

Paige R. Didier[2]

Shannon E. Grogans[2]

Allegra S. Anderson[5]

Samiha Islam[6,7]

Hyung Cho Kim[2,3]

Manuel Kuhn[8]

Rachael M. Tillman[9]

Juyoen Hur[10]

Zachary S. Scott[2]

Andrew S. Fox[11,12]

Kathryn A. DeYoung[2]

Jason F. Smith[2] *

Alexander J. Shackman[2,3,4] *

[1]Department of Psychological & Brain Sciences, George Washington University, Washington, DC 20006 USA. Department of [2]Psychology; [3]Neuroscience and Cognitive Science Program; and [4]Maryland Neuroimaging Center, University of Maryland, College Park, MD 20742 USA. [5]Department of Psychiatry and Human Behavior, Brown University, Providence, RI 02912 USA. [6]Department of Psychology, University of Pennsylvania, Philadelphia, PA USA. [7]Department of Child and Adolescent Psychiatry and Behavioral Sciences, Children's Hospital of Philadelphia, Philadelphia, PA 19139 USA. [8]Center for Depression, Anxiety and Stress Research, McLean Hospital, Harvard Medical School, Belmont, MA 02478 USA. [9]McGill Neuropsychology, Bethesda, MD 20814 USA. [10]Department of Psychology, Yonsei University, Seoul 03722, Republic of Korea. [11]Department of Psychology and [12]California National Primate Research Center, University of California, Davis, CA 95616 USA.

* contributed equally

**Please address manuscript correspondence to**

Alexander J. Shackman

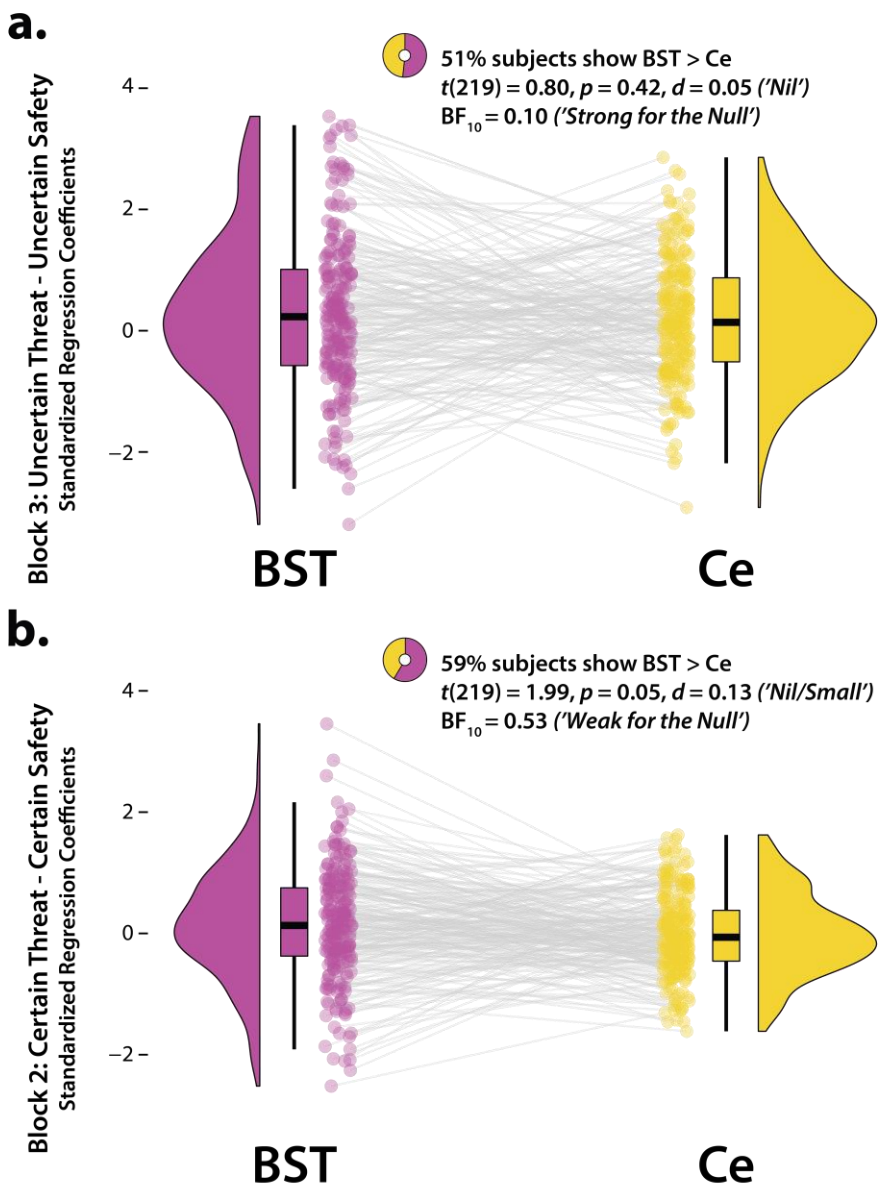


**Supplementary Figure S1. The BST and Ce show statistically similar responses to uncertain- and certain-threat anticipation.** Regional hypothesis testing focused on the conceptually crucial second block of uncertain-threat anticipation (an index of sustained responses to potential threat) and the final block of certain-threat anticipation (an index of phasic responses to acute threat). For completeness, here we describe the remaining contrasts. **a.** ***Uncertain-threat anticipation, third convolved block.*** The BST and Ce show negligible differences in activation during the third block (12.5-18.75 s) of uncertain-threat anticipation, with strong Bayesian evidence for the null hypothesis ($BF_{10}$=0.10; $H_0$ is 10 times more likely than $H_1$). **b.** ***Certain-threat anticipation, second convolved block.*** The BST exhibits greater activation than the Ce during the middle third (6.25-12.5 s) of certain-threat anticipation. While statistically significant, this regional difference runs counter to popular double-dissociation claims, which imply that the Ce is more sensitive to certain threat. On the other hand, it could be viewed as consistent with claims that the BST is more sensitive to spatially and temporally distal threat. Given this inferential ambiguity and the fact that the difference was small-to-nil in magnitude (*d*=0.13), with weak Bayesian evidence for the null ($BF_{10}$=0.53; $H_0$ is 1.9 times more likely than $H_1$), we eschew further interpretation. Raincloud plots indicate the medians (*horizontal lines*), interquartile ranges (*boxes*), and smoothed density distributions. Whiskers depict 1.5× the interquartile range. Colored dots connected by gray lines indicate mean regional for each participant. Note: No spatial smoothing kernel was employed for ROI analyses. Abbreviations—BF, Bayes' factor; BST, bed nucleus of the stria terminalis; Ce, central nucleus of the amygdala; *d*, Cohen's *d*; EAc, central extended amygdala; *t*, Student's *t*-test.

**Supplementary Table S1. [UT > US].** Descriptive statistics for clusters and local maxima from the conventional 'boxcar' analysis showing greater activation for Uncertain-Threat relative to Uncertain-Safety anticipation (FDR $q$<0.05, whole-brain corrected).

| $mm^3$ | Label | *t* | *x* | *y* | *z* |
|---|---|---|---|---|---|
| 113,264 | | | | | |
| | R Accumbens | 3.83 | 10 | 6 | -8 |
| | L Caudate | 7.17 | -16 | 4 | 22 |
| | L Bed Nucleus of Stria Terminalis | 4.79 | -6 | 2 | -4 |
| | R Caudate | 8.48 | 18 | -2 | 22 |
| | R Thalamus | 7.03 | 10 | -2 | 10 |
| | L Thalamus | 6.24 | -6 | -4 | 6 |
| | R Left Thalamus | 6.42 | 0 | -4 | 6 |
| | R Brain-Stem | 10.39 | 2 | -30 | -2 |
| | L Brain-Stem | 6.91 | -4 | -34 | -26 |
| | R Temporal Occipital Fusiform Cortex | 10.16 | 34 | -60 | -20 |
| | L Lingual Gyrus | 4.11 | -10 | -86 | -10 |
| | R Occipital Fusiform Gyrus | 4.05 | 14 | -86 | -12 |
| | R Lingual Gyrus | 3.77 | 6 | -86 | -10 |
| | R Occipital Pole | 3.66 | 10 | -92 | -12 |
| 66,328 | | | | | |
| | R Frontal Pole | 3.26 | 6 | 60 | 32 |
| | L Cingulate Gyrus, anterior division | 11.03 | -8 | 20 | 32 |
| | R Paracingulate Gyrus | 13.55 | 8 | 20 | 36 |
| | R Cingulate Gyrus, anterior division | 12.95 | 10 | 12 | 38 |
| | L Paracingulate Gyrus | 11.54 | -2 | 10 | 42 |
| | R Middle Frontal Gyrus | 3.75 | 34 | 10 | 32 |
| | L Superior Frontal Gyrus | 11.38 | -14 | 4 | 70 |
| | L Juxtapositional Lobule Cortex (formerly Supplementary Motor Cortex) | 10.42 | -4 | 4 | 46 |
| | R Juxtapositional Lobule Cortex (formerly Supplementary Motor Cortex) | 12.73 | 8 | 4 | 54 |
| | R Superior Frontal Gyrus | 11.75 | 18 | 4 | 68 |
| | R Precentral Gyrus | 11.85 | 46 | 0 | 50 |
| | L Precentral Gyrus | 7.67 | -40 | -2 | 48 |
| | R Cingulate Gyrus, posterior division | 6.35 | 8 | -20 | 42 |
| 26,672 | | | | | |
| | R Frontal Pole | 6.28 | 52 | 36 | -6 |
| | R Frontal Orbital Cortex | 15.4 | 34 | 28 | 2 |
| | R Frontal Operculum Cortex | 14.49 | 42 | 22 | 4 |
| | R Inferior Frontal Gyrus, pars opercularis | 10.21 | 54 | 12 | 2 |
| | R Precentral Gyrus | 3.32 | 60 | 12 | 24 |
| | R Putamen | 10.67 | 20 | 10 | -6 |
| | R Central Opercular Cortex | 7.49 | 46 | 2 | 10 |
| | R Pallidum | 5.72 | 14 | -2 | -6 |
| | R Amygdala | 5.59 | 30 | -8 | -14 |
| | R Amygdala | 6.36 | 26 | -10 | -12 |

| | | | | | |
|---|---|---|---|---|---|
| 22,136 | | | | | |
| | R Parietal Operculum Cortex | 6.49 | 54 | -26 | 24 |
| | R Middle Temporal Gyrus, posterior division | 10.82 | 52 | -30 | -4 |
| | R Supramarginal Gyrus, anterior division | 6.57 | 56 | -32 | 34 |
| | R Middle Temporal Gyrus, temporooccipital part | 7.99 | 56 | -40 | 4 |
| | R Supramarginal Gyrus, posterior division | 11.5 | 66 | -42 | 28 |
| | R Angular Gyrus | 6.26 | 56 | -48 | 48 |
| 20,736 | | | | | |
| | L Frontal Operculum Cortex | 14.26 | -32 | 24 | 8 |
| | L Frontal Orbital Cortex | 10.07 | -30 | 22 | -8 |
| | L Caudate | 4.17 | -16 | 18 | 4 |
| | L Putamen | 9.33 | -20 | 10 | -2 |
| | L Inferior Frontal Gyrus, pars opercularis | 9.06 | -54 | 10 | 6 |
| | L Central Opercular Cortex | 9.21 | -48 | 4 | 2 |
| | L Hippocampus | 3.39 | -30 | -18 | -12 |
| 11,008 | | | | | |
| | L Postcentral Gyrus | 4.23 | -56 | -22 | 24 |
| | L Supramarginal Gyrus, anterior division | 4.58 | -62 | -28 | 20 |
| | L Supramarginal Gyrus, posterior division | 8.96 | -54 | -42 | 32 |
| 9,696 | | | | | |
| | L Frontal Pole | 9.94 | -30 | 52 | 28 |
| | L Middle Frontal Gyrus | 6.82 | -36 | 32 | 38 |
| 7,216 | | | | | |
| | R Frontal Pole | 8.17 | 32 | 52 | 30 |
| 3,680 | | | | | |
| | R Lateral Occipital Cortex, inferior division | 3.66 | 42 | -82 | -14 |
| | R Occipital Pole | 6.62 | 24 | -96 | 14 |
| 3,608 | | | | | |
| | R Postcentral Gyrus | 2.75 | 32 | -34 | 46 |
| | R Superior Parietal Lobule | 6.73 | 26 | -44 | 66 |
| 2,496 | | | | | |
| | R Temporal Pole | 4.73 | 48 | 10 | -40 |
| 1,384 | | | | | |
| | L Temporal Pole | 4.73 | -42 | 6 | -44 |
| | L Middle Temporal Gyrus, anterior division | 3.91 | -50 | -2 | -32 |
| | L Inferior Temporal Gyrus, anterior division | 3.08 | -50 | -2 | -44 |
| 792 | | | | | |
| | R Postcentral Gyrus | 4.34 | 46 | -24 | 44 |
| 784 | | | | | |
| | R Occipital Pole | 3.62 | -18 | -100 | 8 |
| 712 | | | | | |
| | R Frontal Pole | 4.61 | 28 | 58 | -8 |

| | | | | | |
|---|---|---|---|---|---|
| 552 | | | | | |
| | L Superior Parietal Lobule | 4.46 | -22 | -50 | 64 |
| 376 | | | | | |
| | L Cingulate Gyrus, posterior division | 4.66 | -10 | -22 | 40 |
| 344 | | | | | |
| | R Hippocampus | 4.36 | 34 | -28 | -6 |
| 280 | | | | | |
| | L Pallidum | 3.86 | -16 | -6 | -2 |
| 176 | | | | | |
| | R Parahippocampal Gyrus, anterior division | 3.99 | 24 | -8 | -32 |
| 128 | | | | | |
| | L Middle Temporal Gyrus, posterior division | 3.19 | -48 | -28 | -6 |
| 120 | | | | | |
| | L Frontal Pole | 3.13 | -30 | 54 | -12 |
| 112 | | | | | |
| | R Thalamus | 4.62 | 8 | -30 | 12 |
| 104 | | | | | |
| | L Cingulate Gyrus, posterior division | 3.48 | -2 | -18 | 30 |
| 88 | | | | | |
| | L Subcallosal Cortex | 3.99 | -10 | 16 | -22 |
| 80 | | | | | |
| | L Thalamus | 4.23 | -6 | -26 | 14 |
| 72 | | | | | |
| | R Inferior Temporal Gyrus, posterior division | 3.32 | 52 | -34 | -28 |
| 56 | | | | | |
| | L Temporal Pole | 3.1 | -36 | 22 | -38 |
| 48 | | | | | |
| | L Temporal Pole | 2.87 | -32 | 16 | -44 |
| 48 | | | | | |
| | L Middle Temporal Gyrus, temporooccipital part | 2.62 | -54 | -46 | 2 |
| 40 | | | | | |
| | L Temporal Pole | 2.65 | -28 | 22 | -34 |
| 32 | | | | | |
| | L Temporal Pole | 3.64 | -32 | 12 | -34 |
| 32 | | | | | |
| | R Thalamus | 3.15 | 22 | -34 | 0 |
| 32 | | | | | |
| | Postcentral Gyrus | 2.57 | -42 | -32 | 44 |
| 32 | | | | | |
| | L Precuneus Cortex | 3.33 | -12 | -46 | 48 |
| 32 | | | | | |
| | L Precuneus Cortex | 2.55 | -10 | -54 | 60 |

| | | | | | |
|---|---|---|---|---|---|
| 32 | | | | | |
| | L Superior Parietal Lobule | 2.9 | -32 | -48 | 70 |
| 24 | | | | | |
| | L Temporal Pole | 2.41 | -20 | 2 | -40 |
| 24 | | | | | |
| | L Frontal Orbital Cortex | 2.95 | -22 | 18 | -24 |
| 24 | | | | | |
| | R Frontal Orbital Cortex | 2.58 | 12 | 18 | -22 |
| 24 | | | | | |
| | L Temporal Pole | 2.69 | -48 | 18 | -18 |
| 16 | | | | | |
| | L Temporal Pole | 2.48 | -28 | 6 | -48 |
| 16 | | | | | |
| | R Inferior Temporal Gyrus, anterior division | 2.44 | 48 | -6 | -42 |
| 16 | | | | | |
| | R Brain-Stem | 2.64 | 18 | -32 | -38 |
| 16 | | | | | |
| | R Inferior Temporal Gyrus, temporooccipital part | 2.93 | 58 | -40 | -20 |
| 16 | | | | | |
| | R Lateral Occipital Cortex, inferior division | 2.86 | 42 | -86 | -8 |
| 16 | | | | | |
| | L Middle Temporal Gyrus, posterior division | 2.67 | -58 | -28 | -6 |
| 16 | | | | | |
| | R Putamen | 3.43 | 32 | -18 | -2 |
| 16 | | | | | |
| | R Supramarginal Gyrus, anterior division | 3.36 | 52 | -22 | 32 |
| 8 | | | | | |
| | R Temporal Fusiform Cortex, anterior division | 2.51 | 26 | -4 | -42 |
| 8 | | | | | |
| | L Temporal Pole | 2.31 | -28 | 16 | -36 |
| 8 | | | | | |
| | L Inferior Temporal Gyrus, posterior division | 2.56 | -50 | -28 | -24 |
| 8 | | | | | |
| | L Inferior Temporal Gyrus, posterior division | 2.51 | -50 | -38 | -20 |
| 8 | | | | | |
| | R Cingulate Gyrus, posterior division | 2.79 | 52 | 8 | -20 |
| 8 | | | | | |
| | R Occipital Pole | 2.31 | 20 | -98 | -16 |
| 8 | | | | | |
| | L Frontal Orbital Cortex | 2.53 | -20 | 24 | -16 |

| | | | | | |
|---|---|---|---|---|---|
| 8 | | | | | |
| | L Middle Temporal Gyrus, posterior division | 2.38 | -56 | -32 | -12 |
| 8 | | | | | |
| | L Frontal Pole | 2.36 | -22 | 54 | -12 |
| 8 | | | | | |
| | R Frontal Pole | 2.58 | 36 | 54 | -8 |
| 8 | | | | | |
| | L Thalamus | 2.84 | -20 | -36 | 0 |
| 8 | | | | | |
| | L Paracingulate Gyrus | 2.49 | -10 | 54 | 12 |
| 8 | | | | | |
| | R Inferior Frontal Gyrus, pars opercularis | 2.32 | 52 | 20 | 22 |

**Supplementary Table S2. [CT > CS].** Descriptive statistics for clusters and local maxima from the conventional 'boxcar' analysis showing greater activation for Certain-Threat relative to Certain-Safety anticipation (FDR $q$<0.05, whole-brain corrected).

| mm³ | Label | *t* | *x* | *y* | *z* |
|---|---|---|---|---|---|
| 65,384 | | | | | |
| | R Brain-Stem | 5.78 | 12 | -28 | -16 |
| | L Brain-Stem | 4.15 | -4 | -40 | -22 |
| | R Lingual Gyrus | 6.87 | 6 | -72 | -10 |
| 25,304 | | | | | |
| | L Paracingulate Gyrus | 7.36 | -8 | 20 | 34 |
| | R Paracingulate Gyrus | 6.75 | 10 | 18 | 36 |
| | R Juxtapositional Lobule Cortex (formerly Supplementary Motor Cortex) | 6.99 | 8 | 6 | 64 |
| | L Superior Frontal Gyrus | 6.64 | -16 | 4 | 68 |
| | L Juxtapositional Lobule Cortex (formerly Supplementary Motor Cortex) | 6.26 | -6 | 2 | 52 |
| | R Superior Frontal Gyrus | 7.79 | 14 | 2 | 68 |
| | R Precentral Gyrus | 6.68 | 48 | 2 | 52 |
| | R Middle Frontal Gyrus | 4.52 | 38 | 0 | 62 |
| 10,168 | | | | | |
| | R Frontal Pole | 4.23 | 52 | 46 | 0 |
| | R Inferior Frontal Gyrus, pars triangularis | 4.94 | 50 | 32 | -2 |
| | R Frontal Orbital Cortex | 6.09 | 40 | 28 | 0 |
| | R Temporal Pole | 3.56 | 50 | 20 | -16 |
| | R Frontal Operculum Cortex | 8.32 | 48 | 18 | 0 |
| 10,000 | | | | | |
| | L Supramarginal Gyrus, anterior division | 2.82 | -50 | -30 | 38 |
| | L Postcentral Gyrus | 4.46 | -36 | -38 | 42 |
| | L Supramarginal Gyrus, posterior division | 6.65 | -56 | -52 | 40 |
| | L Superior Parietal Lobule | 3.52 | -32 | -54 | 44 |
| | L Angular Gyrus | 4.95 | -52 | -56 | 50 |
| | L Lateral Occipital Cortex, superior division | 3.31 | -34 | -62 | 54 |
| 9,296 | | | | | |
| | L Frontal Pole | 6.84 | -40 | 54 | 14 |
| | L Middle Frontal Gyrus | 5.39 | -34 | 30 | 38 |
| 9,192 | | | | | |
| | R Postcentral Gyrus | 4.25 | 40 | -28 | 42 |
| | R Supramarginal Gyrus, anterior division | 3.46 | 50 | -28 | 42 |
| | R Supramarginal Gyrus, posterior division | 6.52 | 64 | -40 | 26 |
| | R Superior Parietal Lobule | 3.71 | 36 | -44 | 58 |
| | R Angular Gyrus | 5.64 | 52 | -46 | 34 |
| 7,016 | | | | | |
| | L Frontal Orbital Cortex | 5.51 | -34 | 28 | 0 |
| | L Insular Cortex | 4.91 | -32 | 26 | 4 |

| | | | | | |
|---|---|---|---|---|---|
| | L Inferior Frontal Gyrus, pars opercularis | 5.49 | -54 | 18 | 0 |
| | L Frontal Operculum Cortex | 8.84 | -40 | 14 | 2 |
| | L Precentral Gyrus | 3.05 | -56 | 8 | 22 |
| 6,936 | | | | | |
| | R Frontal Pole | 6.91 | 30 | 50 | 28 |
| | R Middle Frontal Gyrus | 3.27 | 36 | 34 | 36 |
| 5,232 | | | | | |
| | R Accumbens | 4.2 | 10 | 12 | -4 |
| | R Putamen | 6.58 | 22 | 10 | -4 |
| | R Insular Cortex | 3.1 | 38 | 10 | -6 |
| | L Bed Nucleus of Stria Terminalis | 4.4 | -6 | 4 | 0 |
| | L Caudate | 3.12 | -8 | 2 | 6 |
| | L Thalamus | 4.48 | -2 | -2 | 0 |
| | R Left Thalamus | 6.03 | 0 | -4 | 6 |
| | R Thalamus | 6.7 | 2 | -18 | 0 |
| | L Brain-Stem | 5.16 | -2 | -30 | -4 |
| 3,608 | | | | | |
| | R Middle Temporal Gyrus, posterior division | 4.19 | 60 | -24 | -6 |
| | R Superior Temporal Gyrus, posterior division | 5.95 | 48 | -26 | -4 |
| | R Middle Temporal Gyrus, temporooccipital part | 4.84 | 52 | -42 | 4 |
| | R Supramarginal Gyrus, posterior division | 4.19 | 48 | -42 | 12 |
| 2,904 | | | | | |
| | R Brain-Stem | 6.14 | 6 | -46 | -58 |
| | L Brain-Stem | 4.33 | -6 | -48 | -58 |
| 1,984 | | | | | |
| | L Caudate | 3.62 | -18 | 20 | -2 |
| | L Putamen | 6 | -22 | 16 | -4 |
| 1,744 | | | | | |
| | R Lateral Occipital Cortex, inferior division | 3.83 | 30 | -86 | 0 |
| | R Occipital Pole | 5.12 | 22 | -102 | 12 |
| 1,048 | | | | | |
| | R Caudate | 6.9 | 18 | 8 | 20 |
| 848 | | | | | |
| | L Precentral Gyrus | 4.58 | -48 | 4 | 42 |
| 640 | | | | | |
| | L Frontal Pole | 4 | -32 | 48 | -16 |
| 632 | | | | | |
| | R Hippocampus | 4.45 | 24 | -32 | -4 |
| | R Thalamus | 5.17 | 12 | -34 | 8 |
| 576 | | | | | |
| | L Caudate | 5.65 | -16 | 2 | 22 |
| 368 | | | | | |

| | | | | | |
|---|---|---|---|---|---|
| | L Middle Frontal Gyrus | 3.24 | -26 | -4 | 50 |
| | L Precentral Gyrus | 3.93 | -28 | -8 | 48 |
| 360 | | | | | |
| | R Amygdala | 3.79 | 30 | -4 | -20 |
| | R Amygdala | 4.93 | 30 | -8 | -16 |
| | R Amygdala | 4.79 | 28 | -12 | -14 |
| 240 | | | | | |
| | R Lateral Occipital Cortex, inferior division | 3.65 | 48 | -72 | -12 |
| 200 | | | | | |
| | L Thalamus | 4.43 | -20 | -36 | 2 |
| 192 | | | | | |
| | L Amygdala | 4.43 | -24 | -12 | -10 |
| | L Amygdala | 3.53 | -24 | -12 | -14 |
| | L Hippocampus | 3.08 | -30 | -16 | -14 |
| 192 | | | | | |
| | L Caudate | 4.69 | -18 | 16 | 12 |
| 168 | | | | | |
| | L Brain-Stem | 4.24 | -14 | -26 | -14 |
| 160 | | | | | |
| | L Thalamus | 4.33 | -12 | -10 | 6 |
| 152 | | | | | |
| | L Precuneus Cortex | 2.71 | -10 | -72 | 44 |
| 136 | | | | | |
| | R Cingulate Gyrus, anterior division | 3.53 | 4 | 16 | 26 |
| | L Cingulate Gyrus, anterior division | 3.54 | -2 | 14 | 26 |
| 136 | | | | | |
| | R Precuneus Cortex | 3.44 | 12 | -68 | 48 |
| | R Lateral Occipital Cortex, superior division | 3.24 | 14 | -70 | 52 |
| 128 | | | | | |
| | L Frontal Pole | 3.61 | -52 | 44 | 0 |
| 104 | | | | | |
| | L Brain-Stem | 3.57 | -6 | -38 | -30 |
| 96 | | | | | |
| | L Inferior Temporal Gyrus, posterior division | 4.13 | -54 | -36 | -18 |
| 80 | | | | | |
| | R Putamen | 3.38 | 28 | -2 | 12 |
| 72 | | | | | |
| | R Frontal Pole | 3.18 | 38 | 62 | -4 |
| 72 | | | | | |
| | L Caudate | 3.43 | -12 | 12 | 2 |
| 72 | | | | | |
| | R Caudate | 3.04 | 10 | 2 | 12 |
| | R Thalamus | 2.59 | 10 | -2 | 10 |

| | | | | | |
|---|---|---|---|---|---|
| 72 | | | | | |
| | R Precuneus Cortex | 2.82 | 16 | -70 | 38 |
| 64 | | | | | |
| | L Pallidum | 3.5 | -14 | -2 | -4 |
| 64 | | | | | |
| | R Precentral Gyrus | 3.1 | 38 | -22 | 62 |
| 56 | | | | | |
| | L Temporal Fusiform Cortex, anterior division | 4.28 | -32 | 0 | -50 |
| 56 | | | | | |
| | R Frontal Pole | 3.33 | 26 | 60 | -12 |
| 56 | | | | | |
| | L Caudate | 2.66 | -18 | -14 | 24 |
| 56 | | | | | |
| | R Superior Parietal Lobule | 3.05 | 26 | -48 | 62 |
| 48 | | | | | |
| | L Brain-Stem | 2.61 | -12 | -16 | -22 |
| 48 | | | | | |
| | R Hippocampus/Amygdala | 2.76 | 16 | -12 | -16 |
| 48 | | | | | |
| | R Hippocampus | 3.85 | 36 | -32 | -6 |
| 48 | | | | | |
| | L Lateral Occipital Cortex, inferior division | 3.62 | -42 | -80 | -4 |
| 48 | | | | | |
| | R Inferior Frontal Gyrus, pars opercularis | 3.06 | 56 | 20 | 14 |
| 48 | | | | | |
| | L Cingulate Gyrus, anterior division | 3.22 | -4 | 22 | 20 |
| 40 | | | | | |
| | L Brain-Stem | 4.11 | -10 | -34 | -14 |
| 40 | | | | | |
| | R Frontal Pole | 3.46 | 32 | 60 | -6 |
| 40 | | | | | |
| | L Frontal Pole | 3.19 | -32 | 64 | 6 |
| 40 | | | | | |
| | R Thalamus | 3.37 | 12 | -10 | 8 |
| 40 | | | | | |
| | R Frontal Pole | 2.78 | 8 | 62 | 26 |
| 40 | | | | | |
| | R Middle Frontal Gyrus | 2.89 | 40 | 18 | 34 |
| 32 | | | | | |
| | R Inferior Temporal Gyrus, temporooccipital part | 2.88 | 58 | -58 | -18 |
| 32 | | | | | |
| | L Thalamus | 2.94 | -4 | -28 | 6 |
| 32 | | | | | |

| | | | | | |
|---|---|---|---|---|---|
| | L Precentral Gyrus | 2.88 | -48 | 2 | 20 |
| 32 | | | | | |
| | L Supramarginal Gyrus, anterior division | 2.96 | -54 | -34 | 36 |
| 32 | | | | | |
| | L Superior Frontal Gyrus | 2.9 | -10 | 32 | 52 |
| 32 | | | | | |
| | R Frontal Pole | 2.87 | 10 | 40 | 56 |
| 32 | | | | | |
| | L Precentral Gyrus | 2.65 | -32 | -12 | 64 |
| 24 | | | | | |
| | L Subcallosal Cortex | 2.8 | -8 | 28 | -24 |
| 24 | | | | | |
| | R Inferior Temporal Gyrus, temporooccipital part | 2.87 | 56 | -40 | -18 |
| 24 | | | | | |
| | R Frontal Pole | 2.9 | 36 | 52 | -14 |
| 24 | | | | | |
| | L Frontal Pole | 2.97 | -50 | 46 | -12 |
| 24 | | | | | |
| | R Lateral Occipital Cortex, inferior division | 2.89 | 38 | -86 | -10 |
| 24 | | | | | |
| | R Hippocampus | 2.73 | 34 | -26 | -8 |
| 24 | | | | | |
| | L Middle Temporal Gyrus, posterior division | 2.57 | -70 | -28 | -4 |
| 24 | | | | | |
| | L Occipital Pole | 2.71 | -28 | -92 | 0 |
| | L Lateral Occipital Cortex, inferior division | 2.61 | -30 | -90 | -2 |
| 24 | | | | | |
| | R Inferior Frontal Gyrus, pars triangularis | 2.7 | 52 | 30 | 6 |
| 24 | | | | | |
| | L Cingulate Gyrus, posterior division | 2.74 | -2 | -18 | 32 |
| 24 | | | | | |
| | R Superior Parietal Lobule | 2.62 | 30 | -48 | 46 |
| 24 | | | | | |
| | L Lateral Occipital Cortex, superior division | 2.75 | -26 | -64 | 62 |
| 16 | | | | | |
| | R Temporal Pole | 2.85 | 34 | 20 | -46 |
| 16 | | | | | |
| | R Temporal Pole | 2.99 | 28 | 22 | -40 |
| 16 | | | | | |
| | R Temporal Pole | 2.73 | 22 | 16 | -38 |
| 16 | | | | | |
| | L Temporal Pole | 2.61 | -50 | 12 | -36 |

| | | | | | |
|---|---|---|---|---|---|
| 16 | | | | | |
| | R Temporal Pole | 3.05 | 30 | 24 | -34 |
| 16 | | | | | |
| | L Temporal Pole | 3.26 | -26 | 18 | -34 |
| 16 | | | | | |
| | R Brain-Stem | 2.89 | 0 | -38 | -30 |
| 16 | | | | | |
| | R Brain-Stem | 2.6 | 8 | -38 | -28 |
| 16 | | | | | |
| | L Middle Temporal Gyrus, anterior division | 2.98 | -54 | -8 | -26 |
| 16 | | | | | |
| | R Frontal Pole | 2.82 | 8 | 46 | -28 |
| 16 | | | | | |
| | R Frontal Orbital Cortex | 2.99 | 38 | 20 | -16 |
| 16 | | | | | |
| | L Occipital Fusiform Gyrus | 2.77 | -14 | -86 | -12 |
| 16 | | | | | |
| | L Middle Temporal Gyrus, posterior division | 3.15 | -62 | -22 | -12 |
| 16 | | | | | |
| | L Putamen | 2.73 | -32 | -14 | -10 |
| 16 | | | | | |
| | L Middle Temporal Gyrus, posterior division | 2.88 | -54 | -32 | -6 |
| 16 | | | | | |
| | L Pallidum | 2.85 | -20 | -10 | -4 |
| 16 | | | | | |
| | R Frontal Pole | 2.76 | 26 | 52 | -2 |
| 16 | | | | | |
| | L Frontal Pole | 2.65 | -34 | 60 | 2 |
| 16 | | | | | |
| | R Supramarginal Gyrus, posterior division | 2.86 | 58 | -42 | 14 |
| 16 | | | | | |
| | L Inferior Frontal Gyrus, pars triangularis | 3.18 | -50 | 24 | 14 |
| 16 | | | | | |
| | R Precentral Gyrus | 3.57 | 54 | 4 | 20 |
| 16 | | | | | |
| | R Lateral Occipital Cortex, superior division | 2.62 | 26 | -78 | 30 |
| 16 | | | | | |
| | L Lateral Occipital Cortex, superior division | 2.77 | -26 | -60 | 46 |
| 16 | | | | | |
| | R Superior Parietal Lobule | 2.75 | 30 | -56 | 50 |

| | | | | | |
|---|---|---|---|---|---|
| 16 | | | | | |
| | L Superior Frontal Gyrus | 2.66 | -24 | -8 | 60 |
| 16 | | | | | |
| | L Lateral Occipital Cortex, superior division | 3.02 | -24 | -64 | 68 |
| 16 | | | | | |
| | L Precentral Gyrus | 2.59 | -34 | -6 | 66 |
| 8 | | | | | |
| | R Brain-Stem | 2.57 | 0 | -32 | -60 |
| 8 | | | | | |
| | L Brain-Stem | 2.6 | -2 | -36 | -56 |
| 8 | | | | | |
| | R Brain-Stem | 2.58 | 4 | -38 | -40 |
| 8 | | | | | |
| | L Temporal Pole | 2.52 | -38 | 22 | -40 |
| 8 | | | | | |
| | R Temporal Fusiform Cortex, posterior division | 2.59 | 36 | -18 | -32 |
| 8 | | | | | |
| | L Inferior Temporal Gyrus, posterior division | 2.63 | -62 | -24 | -30 |
| 8 | | | | | |
| | L Brain-Stem | 2.75 | -6 | -20 | -24 |
| 8 | | | | | |
| | L Parahippocampal Gyrus, anterior division | 2.57 | -12 | -4 | -24 |
| 8 | | | | | |
| | L Frontal Orbital Cortex | 2.95 | -32 | 24 | -24 |
| 8 | | | | | |
| | R Temporal Occipital Fusiform Cortex | 2.55 | 48 | -54 | -22 |
| 8 | | | | | |
| | L Temporal Pole | 2.64 | -50 | 16 | -18 |
| 8 | | | | | |
| | R Inferior Temporal Gyrus, temporooccipital part | 2.6 | 62 | -46 | -16 |
| 8 | | | | | |
| | L Middle Temporal Gyrus, posterior division | 2.6 | -56 | -26 | -16 |
| 8 | | | | | |
| | L Insular Cortex | 2.89 | -28 | 14 | -12 |
| 8 | | | | | |
| | L Middle Temporal Gyrus, posterior division | 2.85 | -70 | -26 | -10 |
| 8 | | | | | |
| | L Lingual Gyrus | 2.58 | -12 | -84 | -8 |

| | | | | | |
|---|---|---|---|---|---|
| 8 | | | | | |
| | R Frontal Pole | 2.59 | 28 | 58 | -8 |
| 8 | | | | | |
| | R Thalamus | 2.55 | 6 | -10 | 0 |
| 8 | | | | | |
| | L Caudate | 2.54 | -8 | 10 | 6 |
| 8 | | | | | |
| | L Thalamus | 2.84 | -8 | -34 | 8 |
| 8 | | | | | |
| | L Inferior Frontal Gyrus, pars triangularis | 2.6 | -54 | 22 | 8 |
| 8 | | | | | |
| | L Occipital Pole | 2.7 | -20 | -102 | 10 |
| 8 | | | | | |
| | L Occipital Pole | 2.53 | -26 | -98 | 12 |
| 8 | | | | | |
| | R Central Opercular Cortex | 2.63 | 46 | 2 | 12 |
| 8 | | | | | |
| | L Caudate | 2.82 | -8 | 4 | 12 |
| 8 | | | | | |
| | R Supramarginal Gyrus, posterior division | 2.6 | 66 | -38 | 14 |
| 8 | | | | | |
| | R Frontal Pole | 2.51 | 46 | 46 | 16 |
| 8 | | | | | |
| | R Caudate | 2.72 | 18 | -18 | 22 |
| 8 | | | | | |
| | R Cingulate Gyrus, anterior division | 2.65 | 6 | 20 | 22 |
| 8 | | | | | |
| | R Frontal Pole | 3.18 | 44 | 36 | 24 |
| 8 | | | | | |
| | R Lateral Occipital Cortex, superior division | 2.78 | 28 | -74 | 26 |
| 8 | | | | | |
| | R Cingulate Gyrus, posterior division | 2.76 | 6 | -24 | 28 |
| 8 | | | | | |
| | L Angular Gyrus | 2.76 | -58 | -60 | 30 |
| 8 | | | | | |
| | R Lateral Occipital Cortex, superior division | 2.88 | 26 | -72 | 32 |
| 8 | | | | | |
| | R Superior Frontal Gyrus | 2.54 | 6 | 56 | 32 |
| 8 | | | | | |
| | R Superior Frontal Gyrus | 2.72 | 8 | 48 | 34 |
| 8 | | | | | |
| | R Precuneus Cortex | 2.9 | 8 | -42 | 46 |
| 8 | | | | | |

| | | | | | |
|---|---|---|---|---|---|
| | L Middle Frontal Gyrus | 2.53 | -40 | 0 | 52 |
| 8 | | | | | |
| | L Precentral Gyrus | 2.67 | -40 | -4 | 56 |
| 8 | | | | | |
| | L Superior Frontal Gyrus | 2.61 | -20 | -8 | 60 |
| 8 | | | | | |
| | R Superior Parietal Lobule | 2.52 | 14 | -52 | 76 |

**Supplementary Table S3. [UT > US] ∩ [CT > CS].** Descriptive statistics for clusters and local maxima from the conventional 'boxcar' analysis showing greater activation for Uncertain-Threat relative to Uncertain-Safety anticipation AND greater activation for Certain-Threat relative to Certain-Safety anticipation (FDR $q<0.05$, whole-brain corrected).

| ***mm³*** | **Label** | ***t*** | ***x*** | ***y*** | ***z*** |
|---|---|---|---|---|---|
| 47,064 | | | | | |
| | L Brain-Stem | 3.80 | -4 | -40 | -22 |
| | R Brain-Stem | 2.89 | 6 | -42 | -26 |
| 24,856 | | | | | |
| | L Paracingulate Gyrus | 7.37 | -8 | 20 | 34 |
| | R Paracingulate Gyrus | 6.76 | 10 | 18 | 36 |
| | R Juxtapositional Lobule Cortex (formerly Supplementary Motor Cortex) | 6.99 | 8 | 6 | 64 |
| | L Superior Frontal Gyrus | 6.64 | -16 | 4 | 68 |
| | L Juxtapositional Lobule Cortex (formerly Supplementary Motor Cortex) | 6.27 | -6 | 2 | 52 |
| | R Superior Frontal Gyrus | 7.79 | 14 | 2 | 68 |
| | R Precentral Gyrus | 6.68 | 48 | 2 | 52 |
| | R Middle Frontal Gyrus | 4.52 | 38 | 0 | 62 |
| 9,712 | | | | | |
| | R Frontal Pole | 3.82 | 54 | 40 | -2 |
| | R Inferior Frontal Gyrus, pars triangularis | 4.94 | 50 | 32 | -2 |
| | R Frontal Orbital Cortex | 6.09 | 40 | 28 | 0 |
| | R Frontal Operculum Cortex | 8.33 | 48 | 18 | 0 |
| 7,320 | | | | | |
| | L Frontal Pole | 6.78 | -32 | 54 | 24 |
| | L Middle Frontal Gyrus | 5.39 | -34 | 30 | 38 |
| 6,984 | | | | | |
| | L Frontal Orbital Cortex | 5.52 | -34 | 28 | 0 |
| | L Insular Cortex | 4.91 | -32 | 26 | 4 |
| | L Inferior Frontal Gyrus, pars opercularis | 5.49 | -54 | 18 | 0 |
| | L Frontal Operculum Cortex | 8.85 | -40 | 14 | 2 |
| | L Precentral Gyrus | 3.06 | -56 | 8 | 22 |
| 5,696 | | | | | |
| | R Frontal Pole | 6.91 | 30 | 50 | 28 |
| | R Middle Frontal Gyrus | 3.28 | 36 | 34 | 36 |
| 5,264 | | | | | |
| | R Supramarginal Gyrus, posterior division | 6.53 | 64 | -40 | 26 |
| | R Angular Gyrus | 5.64 | 52 | -46 | 34 |
| 5,152 | | | | | |
| | L Supramarginal Gyrus, posterior division | 6.66 | -56 | -52 | 40 |
| 3,528 | | | | | |
| | R Middle Temporal Gyrus, posterior division | 4.20 | 60 | -24 | -6 |

| | | | | | |
|---|---|---|---|---|---|
| | R Superior Temporal Gyrus, posterior division | 5.96 | 48 | -26 | -4 |
| | R Middle Temporal Gyrus, temporooccipital part | 4.73 | 52 | -42 | 4 |
| | R Supramarginal Gyrus, posterior division | 4.19 | 48 | -42 | 12 |
| 2,648 | | | | | |
| | R Brain-Stem | 5.78 | 8 | -34 | -54 |
| | L Brain-Stem | 4.33 | -6 | -48 | -58 |
| 2,360 | | | | | |
| | R Putamen | 6.59 | 22 | 10 | -4 |
| 1,584 | | | | | |
| | L Putamen | 5.55 | -20 | 14 | -4 |
| 1,424 | | | | | |
| | R Occipital Pole | 4.58 | 20 | -102 | 12 |
| 920 | | | | | |
| | R Caudate | 6.50 | 18 | 8 | 20 |
| 648 | | | | | |
| | L Thalamus | 3.63 | -2 | -18 | -2 |
| | R Thalamus | 4.65 | 2 | -20 | 0 |
| | L Brain-Stem | 5.16 | -2 | -30 | -4 |
| 552 | | | | | |
| | L Thalamus | 3.59 | -2 | -4 | -2 |
| | R Thalamus | 6.04 | 0 | -4 | 6 |
| 536 | | | | | |
| | R Brain-Stem | 5.52 | 12 | -28 | -14 |
| | L Periaqueductal gray | 3.26 | -2 | -36 | -14 |
| | R Periaqueductal gray | 2.85 | 0 | -36 | -10 |
| 504 | | | | | |
| | R Postcentral Gyrus | 3.64 | 46 | -28 | 50 |
| 472 | | | | | |
| | L Precentral Gyrus | 4.12 | -48 | 2 | 40 |
| 448 | | | | | |
| | L Caudate | 5.65 | -16 | 2 | 22 |
| 328 | | | | | |
| | L Middle Frontal Gyrus | 3.25 | -26 | -4 | 50 |
| | L Precentral Gyrus | 3.94 | -28 | -8 | 48 |
| 160 | | | | | |
| | R Superior Parietal Lobule | 3.33 | 34 | -42 | 62 |
| 152 | | | | | |
| | L Thalamus | 4.17 | -10 | -10 | 6 |
| 144 | | | | | |
| | R Amygdala | 3.61 | 30 | -6 | -16 |
| | R Amygdala | 4.09 | 30 | -10 | -14 |
| 136 | | | | | |
| | L Brain-Stem | 4.24 | -14 | -26 | -14 |

| | | | | | |
|---|---|---|---|---|---|
| 104 | | | | | |
| | L Caudate | 3.13 | -8 | 2 | 6 |
| | L Bed Nucleus of the Stria Terminalis | 3.62 | -6 | 2 | 0 |
| 104 | | | | | |
| | R Cingulate Gyrus, anterior division | 3.53 | 4 | 16 | 26 |
| | L Cingulate Gyrus, anterior division | 3.54 | -2 | 14 | 26 |
| 96 | | | | | |
| | L Amygdala | 3.34 | -24 | -12 | -10 |
| | L Amygdala | 2.92 | -26 | -14 | -14 |
| | L Hippocampus | 2.92 | -26 | -14 | -14 |
| 80 | | | | | |
| | R Putamen | 3.39 | 28 | -2 | 12 |
| 72 | | | | | |
| | R Caudate | 3.04 | 10 | 2 | 12 |
| | R Thalamus | 2.59 | 10 | -2 | 10 |
| 72 | | | | | |
| | L Precentral Gyrus | 3.33 | -48 | 0 | 54 |
| 56 | | | | | |
| | L Caudate | 2.67 | -18 | -14 | 24 |
| 56 | | | | | |
| | R Superior Parietal Lobule | 3.05 | 26 | -48 | 62 |
| 48 | | | | | |
| | L Brain-Stem | 3.57 | -6 | -38 | -30 |
| 48 | | | | | |
| | L Pallidum | 3.05 | -14 | -2 | -4 |
| 48 | | | | | |
| | R Thalamus | 3.10 | 2 | -14 | 14 |
| 48 | | | | | |
| | R Inferior Frontal Gyrus, pars opercularis | 3.06 | 56 | 20 | 14 |
| 48 | | | | | |
| | R Precentral Gyrus | 3.11 | 38 | -22 | 62 |
| 40 | | | | | |
| | L Brain-Stem | 2.69 | -8 | -18 | -20 |
| 40 | | | | | |
| | R Hippocampus | 3.78 | 36 | -34 | -6 |
| 40 | | | | | |
| | R Thalamus | 3.37 | 12 | -10 | 8 |
| 40 | | | | | |
| | L Caudate | 3.57 | -16 | 14 | 14 |
| 32 | | | | | |
| | R Accumbens | 3.30 | 8 | 6 | -8 |
| 32 | | | | | |
| | R Thalamus | 3.16 | 22 | -34 | 0 |
| 24 | | | | | |
| | R Hippocampus | 2.73 | 34 | -26 | -8 |
| 24 | | | | | |

| | | | | | |
|---|---|---|---|---|---|
| | R Frontal Pole | 2.80 | 32 | 58 | -6 |
| 24 | | | | | |
| | L Caudate | 2.74 | -18 | 18 | 6 |
| 24 | | | | | |
| | R Thalamus | 3.08 | 6 | -30 | 10 |
| 24 | | | | | |
| | R Middle Frontal Gyrus | 2.73 | 38 | 16 | 34 |
| 24 | | | | | |
| | R Superior Frontal Gyrus | 2.80 | 2 | 30 | 46 |
| 24 | | | | | |
| | L Angular Gyrus | 2.80 | -42 | -54 | 48 |
| 16 | | | | | |
| | R Brain-Stem | 2.60 | 8 | -38 | -28 |
| 16 | | | | | |
| | R Frontal Orbital Cortex | 2.99 | 38 | 20 | -16 |
| 16 | | | | | |
| | L Brain-Stem | 2.61 | -10 | -32 | -12 |
| 16 | | | | | |
| | L Frontal Pole | 2.39 | -32 | 52 | -14 |
| 16 | | | | | |
| | L Pallidum | 2.86 | -20 | -10 | -4 |
| 16 | | | | | |
| | R Thalamus | 2.64 | 14 | -34 | 8 |
| 16 | | | | | |
| | R Supramarginal Gyrus, posterior division | 2.87 | 58 | -42 | 14 |
| 16 | | | | | |
| | L Inferior Frontal Gyrus, pars triangularis | 2.92 | -50 | 22 | 14 |
| 16 | | | | | |
| | L Cingulate Gyrus, anterior division | 3.05 | -2 | 22 | 20 |
| 16 | | | | | |
| | L Cingulate Gyrus, posterior division | 2.69 | -4 | -20 | 30 |
| 16 | | | | | |
| | L Postcentral Gyrus | 2.39 | -42 | -30 | 46 |
| 16 | | | | | |
| | L Precentral Gyrus | 2.63 | -30 | -12 | 64 |
| 16 | | | | | |
| | L Precentral Gyrus | 2.60 | -34 | -6 | 66 |
| 8 | | | | | |
| | R Brain-Stem | 2.57 | 0 | -44 | -60 |
| 8 | | | | | |
| | R Brain-Stem | 2.58 | 0 | -32 | -60 |
| 8 | | | | | |
| | L Brain-Stem | 2.60 | -2 | -36 | -56 |
| 8 | | | | | |
| | R Brain-Stem | 2.58 | 4 | -38 | -40 |
| 8 | | | | | |

| | | | | | |
|---|---|---|---|---|---|
| | R Frontal Pole | 2.46 | 26 | 60 | -12 |
| 8 | | | | | |
| | R Lateral Occipital Cortex, inferior division | 2.47 | 38 | -88 | -10 |
| 8 | | | | | |
| | R Lateral Occipital Cortex, inferior division | 2.56 | 42 | -78 | -10 |
| 8 | | | | | |
| | L Lingual Gyrus | 2.58 | -12 | -84 | -8 |
| 8 | | | | | |
| | R Insular Cortex | 2.54 | 36 | 14 | -8 |
| 8 | | | | | |
| | R Frontal Pole | 2.60 | 28 | 58 | -8 |
| 8 | | | | | |
| | R Insular Cortex | 2.54 | 38 | 16 | -6 |
| 8 | | | | | |
| | L Thalamus | 2.84 | -20 | -36 | 0 |
| 8 | | | | | |
| | R Thalamus | 2.39 | 6 | -10 | 0 |
| 8 | | | | | |
| | L Caudate | 2.67 | -8 | 6 | 0 |
| 8 | | | | | |
| | L Caudate | 2.53 | -8 | 10 | 6 |
| 8 | | | | | |
| | L Inferior Frontal Gyrus, pars triangularis | 2.60 | -54 | 22 | 8 |
| 8 | | | | | |
| | L Occipital Pole | 2.58 | -20 | -102 | 10 |
| 8 | | | | | |
| | R Central Opercular Cortex | 2.64 | 46 | 2 | 12 |
| 8 | | | | | |
| | R Supramarginal Gyrus, posterior division | 2.60 | 66 | -38 | 14 |
| 8 | | | | | |
| | R Cingulate Gyrus, anterior division | 2.66 | 6 | 20 | 22 |
| 8 | | | | | |
| | R Frontal Pole | 2.48 | 6 | 60 | 28 |
| 8 | | | | | |
| | L Postcentral Gyrus | 2.33 | -42 | -34 | 44 |
| 8 | | | | | |
| | L Angular Gyrus | 2.65 | -42 | -56 | 52 |
| 8 | | | | | |
| | L Middle Frontal Gyrus | 2.54 | -40 | 0 | 52 |
| 8 | | | | | |
| | L Precentral Gyrus | 2.68 | -40 | -4 | 56 |

**Supplementary Table S4. [US > UT].** Descriptive statistics for clusters and local maxima from the conventional 'boxcar' analysis showing greater activation for Uncertain-Safety relative to Uncertain-Threat anticipation (FDR $q$<0.05, whole-brain corrected).

| **mm³** | **Label** | ***t*** | ***x*** | ***y*** | ***z*** |
|---|---|---|---|---|---|
| 378,128 | | | | | |
| | L Temporal Pole | 5.92 | -46 | 12 | -14 |
| | R Temporal Pole | 5.59 | 48 | 12 | -12 |
| | R Superior Temporal Gyrus, anterior division | 7.1 | 62 | 2 | -2 |
| | R Amygdala | 7.91 | 22 | 0 | -16 |
| | L Amygdala | 8.5 | -22 | 0 | -18 |
| | R Temporal Fusiform Cortex, anterior division | 5.29 | 30 | 0 | -38 |
| | R Amygdala | 6.12 | 28 | -2 | -12 |
| | L Temporal Fusiform Cortex, anterior division | 4.3 | -30 | -2 | -36 |
| | R Middle Temporal Gyrus, anterior division | 8.09 | 60 | -2 | -18 |
| | L Middle Temporal Gyrus, anterior division | 7.22 | -62 | -4 | -16 |
| | L Superior Temporal Gyrus, anterior division | 5.99 | -60 | -4 | -4 |
| | R Planum Polare | 9.36 | 52 | -4 | -2 |
| | R Amygdala | 8.13 | 16 | -6 | -16 |
| | L Amygdala | 7.13 | -16 | -8 | -16 |
| | L Insular Cortex | 15.08 | -40 | -8 | 6 |
| | L Planum Polare | 9.43 | -52 | -8 | 2 |
| | R Insular Cortex | 15.55 | 38 | -8 | 12 |
| | R Planum Temporale | 12.17 | 64 | -8 | 4 |
| | L Precentral Gyrus | 15.16 | -46 | -12 | 32 |
| | L Putamen | 6.25 | -30 | -12 | -6 |
| | R Heschls Gyrus (includes H1 and H2) | 11.06 | 54 | -12 | 6 |
| | R Hippocampus | 10.69 | 18 | -12 | -22 |
| | R Putamen | 4.54 | 28 | -12 | -6 |
| | L Amygdala | 5.86 | -20 | -14 | -14 |
| | L Central Opercular Cortex | 17.56 | -38 | -14 | 16 |
| | L Juxtapositional Lobule Cortex (formerly Supplementary Motor Cortex) | 8.1 | -8 | -14 | 48 |
| | L Middle Temporal Gyrus, posterior division | 4.48 | -62 | -14 | -22 |
| | R Middle Temporal Gyrus, posterior division | 5.04 | 64 | -16 | -22 |
| | L Planum Temporale | 12.09 | -62 | -20 | 10 |
| | L Parahippocampal Gyrus, anterior division | 3.97 | -32 | -20 | -28 |
| | R Temporal Fusiform Cortex, posterior division | 7.15 | 34 | -22 | -24 |

| | | | | | |
|---|---|---|---|---|---|
| | R Supramarginal Gyrus, anterior division | 4.52 | 62 | -22 | 46 |
| | L Heschls Gyrus (includes H1 and H2) | 9.02 | -46 | -24 | 12 |
| | R Precentral Gyrus | 15.08 | 4 | -26 | 62 |
| | R Inferior Temporal Gyrus, posterior division | 4.06 | 44 | -26 | -20 |
| | L Parietal Operculum Cortex | 15.22 | -34 | -28 | 18 |
| | R Thalamus | 7.84 | 16 | -28 | 2 |
| | R Superior Temporal Gyrus, posterior division | 6.09 | 66 | -28 | 8 |
| | L Thalamus | 8.9 | -14 | -30 | -4 |
| | L Supramarginal Gyrus, anterior division | 3.86 | -56 | -30 | 42 |
| | R Parahippocampal Gyrus, posterior division | 12.24 | 28 | -32 | -18 |
| | R Postcentral Gyrus | 17.12 | 2 | -34 | 62 |
| | L Postcentral Gyrus | 17.03 | -2 | -36 | 62 |
| | L Temporal Fusiform Cortex, posterior division | 11.21 | -28 | -38 | -16 |
| | L Hippocampus | 12.72 | -10 | -42 | 4 |
| | L Inferior Temporal Gyrus, posterior division | 3.56 | -64 | -42 | -20 |
| | R Cingulate Gyrus, posterior division | 13.82 | 10 | -42 | 4 |
| | R Middle Temporal Gyrus, temporooccipital part | 5.89 | 64 | -42 | -12 |
| | L Inferior Temporal Gyrus, temporooccipital part | 5.93 | -46 | -50 | -18 |
| | L Middle Temporal Gyrus, temporooccipital part | 6.57 | -58 | -52 | -10 |
| | L Superior Parietal Lobule | 4.66 | -34 | -52 | 56 |
| | L Cingulate Gyrus, posterior division | 13.01 | -6 | -54 | 24 |
| | L Precuneus Cortex | 14.64 | -2 | -56 | 14 |
| | R Superior Parietal Lobule | 4.72 | 28 | -56 | 62 |
| | R Temporal Occipital Fusiform Cortex | 3.3 | 38 | -56 | -16 |
| | L Lingual Gyrus | 13.56 | -12 | -58 | 4 |
| | L Temporal Occipital Fusiform Cortex | 9.46 | -28 | -58 | -8 |
| | L Angular Gyrus | 4.56 | -42 | -58 | 26 |
| | R Angular Gyrus | 7.24 | 48 | -58 | 22 |
| | R Inferior Temporal Gyrus, temporooccipital part | 5.05 | 60 | -60 | -12 |
| | R Occipital Fusiform Gyrus | 7.61 | 26 | -66 | -10 |
| | R Precuneus Cortex | 19.91 | 2 | -68 | 22 |
| | R Lateral Occipital Cortex, superior division | 12.64 | 52 | -68 | 34 |
| | R Intracalcarine Cortex | 20.12 | 12 | -70 | 12 |
| | R Lateral Occipital Cortex, inferior division | 6.75 | 54 | -72 | 4 |
| | L Occipital Fusiform Gyrus | 5.68 | -26 | -74 | -10 |
| | R Lingual Gyrus | 18.65 | 0 | -74 | 8 |

| | | | | | |
|---|---|---|---|---|---|
| | L Lateral Occipital Cortex, superior division | 10.88 | -46 | -76 | 30 |
| | L Lateral Occipital Cortex, inferior division | 7.64 | -50 | -78 | 4 |
| | L Intracalcarine Cortex | 21 | -14 | -80 | 6 |
| | R Occipital Pole | 7.27 | 8 | -90 | 32 |
| | L Occipital Pole | 8.91 | -2 | -96 | 18 |
| 33,528 | | | | | |
| | L Frontal Pole | 7.6 | -6 | 72 | 4 |
| | R Frontal Pole | 12.12 | 2 | 64 | -4 |
| | L Frontal Medial Cortex | 7.7 | -4 | 52 | -6 |
| | L Cingulate Gyrus, anterior division | 3.65 | -6 | 44 | 4 |
| | R Frontal Medial Cortex | 8.1 | 6 | 40 | -16 |
| | R Paracingulate Gyrus | 6.68 | 6 | 34 | -10 |
| | R Cingulate Gyrus, anterior division | 6.28 | 4 | 34 | 8 |
| | L Paracingulate Gyrus | 5.52 | -6 | 32 | -14 |
| | R Subcallosal Cortex | 7.06 | 0 | 30 | -24 |
| | L Subcallosal Cortex | 4.91 | -6 | 24 | -16 |
| 31,912 | | | | | |
| | L Frontal Orbital Cortex | 9.76 | -34 | 36 | -12 |
| | L Superior Frontal Gyrus | 9.68 | -24 | 24 | 50 |
| | L Middle Frontal Gyrus | 8.8 | -26 | 16 | 50 |
| | L Inferior Frontal Gyrus, pars triangularis | 8.18 | -44 | 30 | 20 |
| | L Frontal Pole | 7.36 | -46 | 36 | 14 |
| 11,928 | | | | | |
| | R Frontal Pole | 7.27 | 52 | 40 | 12 |
| | R Inferior Frontal Gyrus, pars triangularis | 6.42 | 46 | 34 | 16 |
| | R Superior Frontal Gyrus | 12.09 | 26 | 30 | 52 |
| | R Middle Frontal Gyrus | 9.86 | 28 | 20 | 50 |
| 3,896 | | | | | |
| | R Frontal Pole | 10.18 | 36 | 38 | -14 |
| | R Frontal Orbital Cortex | 6.98 | 24 | 32 | -14 |
| 2,864 | | | | | |
| | R Caudate | 5.7 | 14 | 24 | -2 |
| | L Caudate | 5.58 | -12 | 22 | 2 |
| | R Accumbens | 6.62 | 10 | 20 | -4 |
| | R Subcallosal Cortex | 6.31 | 2 | 10 | -4 |
| 1,688 | | | | | |
| | R Brain-Stem | 3.81 | 2 | -18 | -44 |
| | L Brain-Stem | 4.38 | -2 | -26 | -44 |
| 616 | | | | | |
| | R Brain-Stem | 4.47 | 10 | -22 | -42 |
| 616 | | | | | |
| | L Cingulate Gyrus, anterior division | 5.59 | -4 | 0 | 28 |
| | R Cingulate Gyrus, anterior division | 6.05 | 4 | -4 | 30 |
| 600 | | | | | |
| | R Occipital Pole | 3.75 | 18 | -100 | -8 |

| | | | | | |
|---|---|---|---|---|---|
| 312 | | | | | |
| | L Occipital Pole | 2.84 | -24 | -100 | -6 |
| 112 | | | | | |
| | L Temporal Pole | 3.52 | -20 | 12 | -36 |
| 104 | | | | | |
| | R Frontal Pole | 2.86 | 38 | 52 | -4 |
| 96 | | | | | |
| | L Paracingulate Gyrus | 2.72 | -10 | 44 | 18 |
| 72 | | | | | |
| | L Frontal Pole | 3.06 | -10 | 44 | -28 |
| 72 | | | | | |
| | L Frontal Pole | 3.12 | -14 | 52 | -22 |
| 64 | | | | | |
| | R Frontal Pole | 3.88 | 46 | 50 | -16 |
| 64 | | | | | |
| | R Postcentral Gyrus | 3.13 | 38 | -30 | 62 |
| 56 | | | | | |
| | L Superior Parietal Lobule | 2.3 | -40 | -46 | 48 |
| 40 | | | | | |
| | L Temporal Pole | 2.56 | -28 | 4 | -40 |
| 40 | | | | | |
| | R Inferior Temporal Gyrus, posterior division | 3.11 | 52 | -32 | -22 |
| 32 | | | | | |
| | R Temporal Pole | 2.77 | 24 | 10 | -36 |
| 32 | | | | | |
| | L Insular Cortex | 3.13 | -32 | 6 | 8 |
| 24 | | | | | |
| | R Frontal Pole | 2.48 | 28 | 62 | 2 |
| 24 | | | | | |
| | L Superior Parietal Lobule | 2.15 | -36 | -44 | 42 |
| 16 | | | | | |
| | L Temporal Pole | 2.41 | -34 | 6 | -36 |
| 16 | | | | | |
| | R Frontal Pole | 2.29 | 24 | 56 | 10 |
| 16 | | | | | |
| | R Precentral Gyrus | 2.23 | 60 | 10 | 34 |
| 16 | | | | | |
| | R Cingulate Gyrus, anterior division | 2.35 | 8 | -8 | 40 |
| 8 | | | | | |
| | R Parahippocampal Gyrus, anterior division | 2.19 | 32 | -16 | -32 |
| 8 | | | | | |
| | L Inferior Temporal Gyrus, posterior division | 2.12 | -54 | -38 | -28 |
| 8 | | | | | |

| | | | | | |
|---|---|---|---|---|---|
| | L Frontal Orbital Cortex | 2.35 | -18 | 28 | -22 |
| 8 | | | | | |
| | R Frontal Pole | 2.61 | 18 | 54 | -22 |
| 8 | | | | | |
| | L Frontal Medial Cortex | 2.13 | -8 | 46 | -20 |
| 8 | | | | | |
| | L Frontal Pole | 2.43 | -20 | 44 | -16 |
| 8 | | | | | |
| | R Frontal Pole | 2.15 | 28 | 62 | -16 |
| 8 | | | | | |
| | R Frontal Pole | 2.2 | 46 | 52 | -6 |
| 8 | | | | | |
| | L Brain-Stem | 2.19 | -6 | -36 | -4 |
| 8 | | | | | |
| | L Frontal Pole | 2.27 | -38 | 44 | -2 |
| 8 | | | | | |
| | L Caudate | 2.38 | -6 | 16 | 2 |
| 8 | | | | | |
| | L Putamen | 2.27 | -28 | -12 | 12 |
| 8 | | | | | |
| | R Cingulate Gyrus, anterior division | 2.16 | 4 | -12 | 34 |
| 8 | | | | | |
| | R Supramarginal Gyrus, posterior division | 2.12 | 44 | -38 | 38 |
| 8 | | | | | |
| | L Cingulate Gyrus, posterior division | 2.54 | -10 | -32 | 38 |
| 8 | | | | | |
| | R Middle Frontal Gyrus | 2.13 | 40 | 18 | 38 |
| 8 | | | | | |
| | R Superior Parietal Lobule | 2.14 | 28 | -56 | 44 |
| 8 | | | | | |
| | R Superior Frontal Gyrus | 2.31 | 28 | 2 | 58 |

**Supplementary Table S5. [CS > CT].** Descriptive statistics for clusters and local maxima from the conventional 'boxcar' analysis showing greater activation for Certain-Safety relative to Certain-Threat anticipation (FDR $q$<0.05, whole-brain corrected).

| ***mm³*** | **Label** | ***t*** | ***x*** | ***y*** | ***z*** |
|---|---|---|---|---|---|
| 244,512 | | | | | |
| | L Frontal Orbital Cortex | 5.13 | -28 | 12 | -24 |
| | L Temporal Pole | 4.82 | -32 | 10 | -26 |
| | L Amygdala | 3.89 | -24 | 0 | -14 |
| | L Superior Temporal Gyrus, anterior division | 4.94 | -60 | 0 | -10 |
| | L Amygdala | 3.87 | -26 | -2 | -12 |
| | L Pallidum | 2.89 | -20 | -4 | 0 |
| | L Middle Temporal Gyrus, anterior division | 6.23 | -60 | -6 | -12 |
| | R Temporal Fusiform Cortex, anterior division | 2.91 | 38 | -8 | -38 |
| | L Planum Polare | 6.93 | -48 | -10 | 0 |
| | L Juxtapositional Lobule Cortex (formerly Supplementary Motor Cortex) | 5.85 | -10 | -10 | 42 |
| | R Cingulate Gyrus, anterior division | 5.15 | 10 | -10 | 42 |
| | R Juxtapositional Lobule Cortex (formerly Supplementary Motor Cortex) | 5.08 | 0 | -10 | 48 |
| | L Left Putamen | 5.91 | -32 | -12 | 2 |
| | L Postcentral Gyrus | 8.18 | -54 | -16 | 50 |
| | L Central Opercular Cortex | 6.6 | -52 | -16 | 10 |
| | L Parahippocampal Gyrus, anterior division | 4.98 | -20 | -16 | -24 |
| | L Middle Temporal Gyrus, posterior division | 4.19 | -52 | -16 | -10 |
| | R Inferior Temporal Gyrus, posterior division | 3.44 | 44 | -16 | -32 |
| | L Planum Temporale | 7.29 | -62 | -18 | 8 |
| | R Parahippocampal Gyrus, anterior division | 6.58 | 20 | -20 | -22 |
| | L Heschls Gyrus (includes H1 and H2) | 8.09 | -50 | -22 | 8 |
| | L Hippocampus | 4.73 | -24 | -24 | -10 |
| | L Precentral Gyrus | 9.2 | -26 | -26 | 66 |
| | R Precentral Gyrus | 8.59 | 6 | -26 | 56 |
| | R Hippocampus | 7.93 | 24 | -26 | -8 |
| | R Temporal Fusiform Cortex, posterior division | 3.61 | 38 | -26 | -30 |
| | L Insular Cortex | 8.85 | -32 | -28 | 16 |
| | L Parietal Operculum Cortex | 3.76 | -42 | -30 | 18 |
| | R Thalamus | 5.84 | 18 | -30 | -4 |
| | L Thalamus | 4.76 | -12 | -34 | 0 |

| | | | | | |
|---|---|---|---|---|---|
| | R Parahippocampal Gyrus, posterior division | 10.16 | 22 | -34 | -18 |
| | L Parahippocampal Gyrus, posterior division | 9.42 | -30 | -40 | -12 |
| | L Temporal Fusiform Cortex, posterior division | 3.22 | -34 | -42 | -20 |
| | R Postcentral Gyrus | 7.07 | 10 | -42 | 74 |
| | L Cingulate Gyrus, posterior division | 9.65 | -10 | -44 | 2 |
| | L Superior Parietal Lobule | 4.9 | -22 | -44 | 66 |
| | R Lingual Gyrus | 12 | 14 | -44 | -2 |
| | L Lingual Gyrus | 12.19 | -20 | -46 | -10 |
| | R Cingulate Gyrus, posterior division | 9.68 | 10 | -46 | 4 |
| | R Angular Gyrus | 5.03 | 48 | -54 | 20 |
| | R Superior Parietal Lobule | 3.47 | 14 | -54 | 68 |
| | L Angular Gyrus | 3.39 | -60 | -58 | 14 |
| | R Temporal Occipital Fusiform Cortex | 5.37 | 32 | -58 | -14 |
| | R Middle Temporal Gyrus, temporooccipital part | 2.75 | 52 | -58 | 10 |
| | L Precuneus Cortex | 9.3 | -8 | -60 | 14 |
| | L Inferior Temporal Gyrus, temporooccipital part | 3.77 | -54 | -60 | -12 |
| | L Temporal Occipital Fusiform Cortex | 3.61 | -38 | -60 | -18 |
| | L Middle Temporal Gyrus, temporooccipital part | 3.52 | -50 | -60 | 0 |
| | R Precuneus Cortex | 15.18 | 0 | -68 | 18 |
| | R Occipital Fusiform Gyrus | 9.69 | 28 | -68 | -6 |
| | L Occipital Fusiform Gyrus | 8.43 | -28 | -74 | -8 |
| | R Intracalcarine Cortex | 20.88 | 12 | -74 | 12 |
| | R Lateral Occipital Cortex, inferior division | 6.65 | 42 | -74 | 10 |
| | L Intracalcarine Cortex | 21.08 | -10 | -76 | 10 |
| | L Lateral Occipital Cortex, inferior division | 4.86 | -52 | -76 | 4 |
| | R Lateral Occipital Cortex, superior division | 7.02 | 50 | -76 | 16 |
| | L Lateral Occipital Cortex, superior division | 6.96 | -46 | -80 | 18 |
| | R Cuneal Cortex | 8.42 | 0 | -86 | 34 |
| | L Occipital Pole | 10.01 | -2 | -96 | 10 |
| | R Occipital Pole | 8.8 | 8 | -96 | 18 |
| 56,792 | | | | | |
| | R Frontal Pole | 7.85 | 0 | 64 | -10 |
| | L Frontal Pole | 6.29 | -6 | 58 | -12 |
| | R Frontal Medial Cortex | 7.04 | 0 | 50 | -12 |
| | L Paracingulate Gyrus | 5.03 | -6 | 46 | -6 |
| | R Paracingulate Gyrus | 6.58 | 0 | 42 | -10 |
| | L Cingulate Gyrus, anterior division | 4.99 | -2 | 40 | -2 |

| | | | | | |
|---|---|---|---|---|---|
| | L Frontal Medial Cortex | 6.29 | -8 | 36 | -12 |
| | R Cingulate Gyrus, anterior division | 4.97 | 2 | 34 | 14 |
| | R Frontal Orbital Cortex | 4.54 | 22 | 28 | -20 |
| | R Right Accumbens | 5.15 | 8 | 18 | -6 |
| | R Temporal Pole | 4.59 | 36 | 18 | -28 |
| | L Subcallosal Cortex | 5.65 | -2 | 16 | -6 |
| | R Subcallosal Cortex | 5.13 | 4 | 14 | -10 |
| | R Superior Temporal Gyrus, anterior division | 4.59 | 60 | -2 | -12 |
| | R Middle Temporal Gyrus, anterior division | 4.48 | 56 | -2 | -20 |
| | R Middle Temporal Gyrus, posterior division | 4.24 | 54 | -10 | -16 |
| | R Pallidum | 3.68 | 24 | -10 | 6 |
| | R Postcentral Gyrus | 6.62 | 50 | -12 | 50 |
| | R Central Opercular Cortex | 7.15 | 38 | -14 | 18 |
| | R Precentral Gyrus | 6.94 | 36 | -14 | 44 |
| | R Putamen | 4.35 | 30 | -14 | 2 |
| | R Planum Polare | 6.5 | 40 | -20 | 0 |
| | R Insular Cortex | 9.1 | 36 | -22 | 14 |
| | R Heschls Gyrus (includes H1 and H2) | 6.98 | 38 | -22 | 6 |
| | R Planum Temporale | 6.2 | 62 | -22 | 12 |
| | R Superior Temporal Gyrus, posterior division | 3.63 | 70 | -24 | 4 |
| 6,584 | | | | | |
| | R Frontal Pole | 2.79 | 20 | 44 | 46 |
| | R Superior Frontal Gyrus | 6.03 | 20 | 30 | 42 |
| | R Middle Frontal Gyrus | 7.68 | 26 | 16 | 46 |
| 6,400 | | | | | |
| | L Middle Frontal Gyrus | 4.59 | -28 | 26 | 46 |
| | L Superior Frontal Gyrus | 6.29 | -22 | 24 | 40 |
| 1,272 | | | | | |
| | L Inferior Frontal Gyrus, pars triangularis | 4.19 | -48 | 30 | 16 |
| | L Middle Frontal Gyrus | 3.62 | -46 | 22 | 26 |
| | L Inferior Frontal Gyrus, pars opercularis | 3.16 | -42 | 14 | 26 |
| 976 | | | | | |
| | R Frontal Pole | 3.01 | 52 | 36 | 12 |
| | R Inferior Frontal Gyrus, pars triangularis | 2.98 | 46 | 32 | 14 |
| | R Middle Frontal Gyrus | 3.52 | 52 | 28 | 28 |
| 912 | | | | | |
| | L Frontal Orbital Cortex | 5.05 | -32 | 34 | -12 |
| 704 | | | | | |
| | L Brain-Stem | 3.78 | -14 | -26 | -34 |
| 448 | | | | | |
| | R Frontal Pole | 4.52 | 36 | 34 | -16 |

| | | | | | |
|---|---|---|---|---|---|
| | R Frontal Orbital Cortex | 3.17 | 40 | 28 | -14 |
| 320 | | | | | |
| | R Amygdala | 3.87 | 22 | 2 | -16 |
| | R Amygdala | 2.72 | 18 | 0 | -24 |
| | R Amygdala | 4.85 | 26 | 0 | -12 |
| 248 | | | | | |
| | L Middle Frontal Gyrus | 3.49 | -54 | 16 | 34 |
| 232 | | | | | |
| | R Temporal Pole | 3.27 | 30 | 6 | -38 |
| | R Parahippocampal Gyrus, anterior division | 3.35 | 26 | 2 | -34 |
| 168 | | | | | |
| | L Angular Gyrus | 2.81 | -56 | -52 | 16 |
| 160 | | | | | |
| | R Temporal Fusiform Cortex, anterior division | 3.35 | 26 | 0 | -44 |
| 144 | | | | | |
| | L Frontal Pole | 2.77 | -12 | 46 | 44 |
| 120 | | | | | |
| | R Thalamus | 3.55 | 6 | -14 | 12 |
| 104 | | | | | |
| | R Brain-Stem | 4.12 | 10 | -14 | -34 |
| 104 | | | | | |
| | R Lateral Occipital Cortex, superior division | 3.18 | 34 | -72 | 50 |
| 96 | | | | | |
| | R Brain-Stem | 3.68 | 0 | -18 | -30 |
| | L Brain-Stem | 2.83 | -2 | -20 | -32 |
| 88 | | | | | |
| | L Left Thalamus | 3.17 | -18 | -24 | 10 |
| 88 | | | | | |
| | R Insular Cortex | 2.56 | 34 | 8 | 8 |
| 80 | | | | | |
| | R Inferior Temporal Gyrus, anterior division | 3.16 | 42 | 2 | -42 |
| 64 | | | | | |
| | L Temporal Fusiform Cortex, anterior division | 2.93 | -34 | -6 | -46 |
| 64 | | | | | |
| | R Temporal Fusiform Cortex, anterior division | 3.33 | 34 | -2 | -42 |
| 64 | | | | | |
| | R Inferior Temporal Gyrus, posterior division | 2.96 | 52 | -18 | -34 |
| 64 | | | | | |

| | | | | | |
|---|---|---|---|---|---|
| | R Frontal Pole | 3.83 | 12 | 36 | -20 |
| 64 | | | | | |
| | R Middle Temporal Gyrus, posterior division | 2.93 | 70 | -24 | -14 |
| 64 | | | | | |
| | R Thalamus | 3.1 | 14 | -24 | 18 |
| 56 | | | | | |
| | R Brain-Stem | 3.35 | 8 | -44 | -36 |
| 56 | | | | | |
| | R Frontal Orbital Cortex | 2.95 | 14 | 30 | -18 |
| | R Subcallosal Cortex | 2.81 | 10 | 28 | -20 |
| 56 | | | | | |
| | R Middle Temporal Gyrus, temporooccipital part | 2.76 | 54 | -56 | 2 |
| 56 | | | | | |
| | R Thalamus | 2.84 | 20 | -26 | 4 |
| 56 | | | | | |
| | R Cingulate Gyrus, anterior division | 3.84 | 4 | 10 | 36 |
| 56 | | | | | |
| | R Cingulate Gyrus, posterior division | 2.41 | 14 | -32 | 38 |
| 48 | | | | | |
| | R Frontal Pole | 2.87 | 14 | 56 | -20 |
| 48 | | | | | |
| | L Left Thalamus | 2.86 | -14 | -24 | 18 |
| 48 | | | | | |
| | R Middle Frontal Gyrus | 2.82 | 42 | 18 | 52 |
| 40 | | | | | |
| | R Frontal Pole | 2.98 | 26 | 66 | 0 |
| 40 | | | | | |
| | L Superior Temporal Gyrus, posterior division | 2.81 | -56 | -36 | 4 |
| 40 | | | | | |
| | L Superior Parietal Lobule | 2.71 | -22 | -56 | 60 |
| 32 | | | | | |
| | L Brain-Stem | 2.91 | -4 | -42 | -36 |
| 32 | | | | | |
| | R Frontal Pole | 2.74 | 14 | 36 | -26 |
| 32 | | | | | |
| | L Frontal Pole | 2.52 | -12 | 36 | -26 |
| 32 | | | | | |
| | R Amygdala | 3.31 | 18 | -6 | -16 |
| 32 | | | | | |
| | L Temporal Pole | 3.04 | -48 | 10 | -10 |
| 32 | | | | | |
| | L Frontal Pole | 2.51 | -18 | 70 | -8 |

| | | | | | |
|---|---|---|---|---|---|
| 32 | | | | | |
| | L Frontal Pole | 3.53 | -14 | 62 | 16 |
| 24 | | | | | |
| | R Brain-Stem | 2.53 | 4 | -30 | -46 |
| 24 | | | | | |
| | R Inferior Temporal Gyrus, posterior division | 2.42 | 60 | -16 | -38 |
| 24 | | | | | |
| | L Temporal Fusiform Cortex, anterior division | 2.6 | -36 | -10 | -38 |
| 24 | | | | | |
| | L Temporal Pole | 2.78 | -34 | 10 | -36 |
| 24 | | | | | |
| | L Inferior Temporal Gyrus, posterior division | 2.51 | -58 | -40 | -28 |
| 24 | | | | | |
| | R Temporal Pole | 3.1 | 44 | 20 | -26 |
| 24 | | | | | |
| | L Frontal Orbital Cortex | 2.84 | -22 | 16 | -20 |
| 24 | | | | | |
| | L Frontal Pole | 2.57 | -14 | 50 | -20 |
| 24 | | | | | |
| | L Inferior Temporal Gyrus, temporooccipital part | 2.59 | -58 | -48 | -16 |
| 24 | | | | | |
| | L Frontal Orbital Cortex | 2.41 | -12 | 14 | -18 |
| 24 | | | | | |
| | R Planum Polare | 2.85 | 44 | -6 | -12 |
| 24 | | | | | |
| | R Lateral Occipital Cortex, inferior division | 2.55 | 52 | -76 | -6 |
| 24 | | | | | |
| | L Frontal Pole | 2.78 | -6 | 50 | 48 |
| 16 | | | | | |
| | R Temporal Fusiform Cortex, anterior division | 2.73 | 30 | -6 | -46 |
| 16 | | | | | |
| | L Temporal Fusiform Cortex, anterior division | 2.37 | -26 | -8 | -42 |
| 16 | | | | | |
| | L Temporal Pole | 2.44 | -26 | 2 | -42 |
| 16 | | | | | |
| | R Brain-Stem | 2.94 | 10 | -24 | -40 |
| 16 | | | | | |
| | L Brain-Stem | 2.86 | -2 | -24 | -38 |

| | | | | | |
|---|---|---|---|---|---|
| 16 | | | | | |
| | R Temporal Pole | 2.52 | 24 | 12 | -36 |
| 16 | | | | | |
| | L Inferior Temporal Gyrus, posterior division | 2.82 | -50 | -20 | -30 |
| 16 | | | | | |
| | R Temporal Pole | 2.69 | 52 | 20 | -26 |
| 16 | | | | | |
| | L Brain-Stem | 2.54 | -14 | -32 | -26 |
| 16 | | | | | |
| | R Brain-Stem | 2.79 | 10 | -28 | -24 |
| 16 | | | | | |
| | R Frontal Pole | 2.39 | 14 | 48 | -26 |
| 16 | | | | | |
| | R Frontal Pole | 2.44 | 14 | 42 | -24 |
| 16 | | | | | |
| | L Lateral Occipital Cortex, inferior division | 2.59 | -38 | -84 | -20 |
| 16 | | | | | |
| | L Frontal Orbital Cortex | 2.95 | -42 | 22 | -16 |
| 16 | | | | | |
| | R Frontal Pole | 2.49 | 24 | 54 | -16 |
| 16 | | | | | |
| | L Inferior Temporal Gyrus, temporooccipital part | 2.56 | -52 | -50 | -12 |
| 16 | | | | | |
| | L Insular Cortex | 2.55 | -38 | -4 | -12 |
| 16 | | | | | |
| | R Frontal Pole | 2.72 | 40 | 44 | -12 |
| 16 | | | | | |
| | R Pallidum | 2.64 | 24 | -4 | -6 |
| 16 | | | | | |
| | R Middle Temporal Gyrus, temporooccipital part | 2.62 | 66 | -54 | -2 |
| 16 | | | | | |
| | L Caudate | 2.74 | -14 | 24 | -2 |
| 16 | | | | | |
| | R Pallidum | 2.79 | 20 | -6 | 4 |
| 16 | | | | | |
| | R Thalamus | 2.58 | 16 | -22 | 12 |
| 16 | | | | | |
| | R Caudate | 2.45 | 14 | 18 | 14 |
| 16 | | | | | |
| | R Cingulate Gyrus, anterior division | 2.41 | 4 | -8 | 30 |
| 16 | | | | | |
| | R Cingulate Gyrus, anterior division | 2.79 | 0 | -2 | 30 |

| | | | | | |
|---|---|---|---|---|---|
| 16 | | | | | |
| | L Frontal Pole | 3.03 | -14 | 52 | 30 |
| 16 | | | | | |
| | R Superior Frontal Gyrus | 2.53 | 0 | 50 | 36 |
| 16 | | | | | |
| | R Middle Frontal Gyrus | 2.71 | 42 | 18 | 40 |
| 8 | | | | | |
| | L Temporal Pole | 2.28 | -34 | 8 | -48 |
| 8 | | | | | |
| | R Inferior Temporal Gyrus, posterior division | 2.41 | 44 | -14 | -40 |
| 8 | | | | | |
| | L Parahippocampal Gyrus, anterior division | 2.71 | -22 | -2 | -38 |
| 8 | | | | | |
| | R Inferior Temporal Gyrus, anterior division | 2.46 | 38 | 2 | -38 |
| 8 | | | | | |
| | L Temporal Pole | 2.42 | -32 | 12 | -38 |
| 8 | | | | | |
| | R Brain-Stem | 2.44 | 12 | -24 | -36 |
| 8 | | | | | |
| | L Parahippocampal Gyrus, anterior division | 2.42 | -30 | -16 | -34 |
| 8 | | | | | |
| | R Brain-Stem | 2.55 | 12 | -30 | -32 |
| 8 | | | | | |
| | L Brain-Stem | 2.29 | -2 | -26 | -28 |
| 8 | | | | | |
| | L Temporal Fusiform Cortex, posterior division | 2.5 | -40 | -22 | -28 |
| 8 | | | | | |
| | L Temporal Fusiform Cortex, posterior division | 2.75 | -42 | -40 | -24 |
| 8 | | | | | |
| | L Temporal Fusiform Cortex, posterior division | 2.38 | -46 | -40 | -24 |
| 8 | | | | | |
| | R Inferior Temporal Gyrus, posterior division | 2.38 | 60 | -30 | -24 |
| 8 | | | | | |
| | L Frontal Orbital Cortex | 2.3 | -28 | 26 | -22 |
| 8 | | | | | |
| | R Temporal Fusiform Cortex, posterior division | 2.51 | 42 | -24 | -20 |
| 8 | | | | | |

| | | | | | |
|---|---|---|---|---|---|
| | L Frontal Orbital Cortex | 2.43 | -24 | 24 | -20 |
| 8 | | | | | |
| | R Frontal Pole | 2.63 | 16 | 40 | -20 |
| 8 | | | | | |
| | L Inferior Temporal Gyrus, temporooccipital part | 2.39 | -48 | -46 | -18 |
| 8 | | | | | |
| | R Temporal Occipital Fusiform Cortex | 2.68 | 42 | -44 | -16 |
| 8 | | | | | |
| | R Inferior Temporal Gyrus, temporooccipital part | 2.28 | 46 | -42 | -16 |
| 8 | | | | | |
| | L Amygdala | 2.37 | -14 | -8 | -16 |
| 8 | | | | | |
| | L Frontal Orbital Cortex | 2.6 | -14 | 20 | -16 |
| 8 | | | | | |
| | L Brain-Stem | 2.29 | -8 | -34 | -8 |
| 8 | | | | | |
| | L Inferior Temporal Gyrus, temporooccipital part | 2.34 | -42 | -58 | -6 |
| 8 | | | | | |
| | L Superior Temporal Gyrus, posterior division | 2.42 | -60 | -22 | -2 |
| 8 | | | | | |
| | R Middle Temporal Gyrus, posterior division | 2.37 | 70 | -38 | 0 |
| 8 | | | | | |
| | R Paracingulate Gyrus | 2.3 | 12 | 52 | 2 |
| 8 | | | | | |
| | R Thalamus | 2.39 | 8 | -14 | 4 |
| 8 | | | | | |
| | R Thalamus | 2.36 | 6 | -20 | 8 |
| 8 | | | | | |
| | L Caudate | 2.53 | -12 | 20 | 8 |
| 8 | | | | | |
| | R Middle Temporal Gyrus, temporooccipital part | 2.34 | 68 | -46 | 10 |
| 8 | | | | | |
| | L Cingulate Gyrus, anterior division | 2.93 | -4 | 26 | 16 |
| 8 | | | | | |
| | R Angular Gyrus | 2.28 | 58 | -48 | 18 |
| 8 | | | | | |
| | L Thalamus | 2.3 | -6 | -14 | 18 |
| 8 | | | | | |
| | L Cingulate Gyrus, anterior division | 2.66 | -4 | 0 | 28 |

| | | | | | |
|---|---|---|---|---|---|
| 8 | | | | | |
| | R Cingulate Gyrus, anterior division | 2.56 | 2 | 2 | 28 |
| 8 | | | | | |
| | L Supramarginal Gyrus, anterior division | 2.27 | -60 | -28 | 32 |
| 8 | | | | | |
| | R Cingulate Gyrus, anterior division | 2.4 | 4 | -14 | 34 |
| 8 | | | | | |
| | L Lateral Occipital Cortex, superior division | 2.33 | -34 | -68 | 38 |
| 8 | | | | | |
| | R Supramarginal Gyrus, anterior division | 2.33 | 58 | -24 | 40 |
| 8 | | | | | |
| | L Superior Frontal Gyrus | 2.51 | -2 | 42 | 42 |
| 8 | | | | | |
| | L Superior Frontal Gyrus | 2.45 | -2 | 48 | 46 |
| 8 | | | | | |
| | R Frontal Pole | 2.38 | 12 | 50 | 48 |
| 8 | | | | | |
| | L Superior Frontal Gyrus | 2.46 | -6 | 38 | 56 |

**Supplementary Table S6. [$UT_{sustained}$ > $US_{sustained}$].** Descriptive statistics for clusters and local maxima showing greater activation for Uncertain-Threat relative to Uncertain-Safety anticipation for the OSP Sustained regressor (FDR *q*<0.05, whole-brain corrected).

| ***mm³*** | **Label** | ***t*** | ***x*** | ***y*** | ***z*** |
|---|---|---|---|---|---|
| 125,152 | | | | | |
| | R Frontal Operculum Cortex | 13.66 | 38 | 22 | 8 |
| | R Frontal Orbital Cortex | 9.86 | 34 | 20 | -10 |
| | R Central Opercular Cortex | 11.39 | 42 | 10 | 4 |
| | R Inferior Frontal Gyrus, pars opercularis | 9.27 | 54 | 10 | 4 |
| | R Precentral Gyrus | 6.78 | 64 | 8 | 10 |
| | R Putamen | 9.99 | 24 | 6 | -8 |
| | L Caudate | 7.01 | -16 | 4 | 22 |
| | R Pallidum | 5.59 | 14 | 0 | -4 |
| | R Caudate | 8.51 | 18 | -2 | 22 |
| | R Thalamus | 7.42 | 10 | -2 | 10 |
| | R Thalamus | 5.13 | 0 | -4 | 6 |
| | L Thalamus | 6.51 | -10 | -10 | 16 |
| | R Amygdala | 5.84 | 26 | -12 | -12 |
| | R Brain-Stem | 7.68 | 2 | -30 | -2 |
| | R Periaqueductal gray | 4.29 | 2 | -34 | -14 |
| | L Brain-Stem | 6.38 | -10 | -38 | -48 |
| | R Temporal Occipital Fusiform Cortex | 9.94 | 34 | -60 | -20 |
| | R Lingual Gyrus | 2.96 | 6 | -86 | -10 |
| 60,160 | | | | | |
| | R Paracingulate Gyrus | 12.26 | 10 | 20 | 34 |
| | L Paracingulate Gyrus | 10.42 | -8 | 12 | 38 |
| | R Cingulate Gyrus, anterior division | 12.74 | 10 | 12 | 38 |
| | L Cingulate Gyrus, anterior division | 10.45 | -6 | 10 | 40 |
| | R Juxtapositional Lobule Cortex (formerly Supplementary Motor Cortex) | 11.99 | 6 | 6 | 54 |
| | R Superior Frontal Gyrus | 10.76 | 16 | 6 | 68 |
| | L Superior Frontal Gyrus | 10.47 | -14 | 4 | 70 |
| | L Juxtapositional Lobule Cortex (formerly Supplementary Motor Cortex) | 10.02 | -4 | 4 | 46 |
| | L Middle Frontal Gyrus | 5.62 | -42 | 0 | 56 |
| | R Precentral Gyrus | 11.4 | 46 | 0 | 50 |
| | L Precentral Gyrus | 8.03 | -40 | -4 | 48 |
| | R Cingulate Gyrus, posterior division | 5.95 | 8 | -20 | 42 |
| 19,400 | | | | | |
| | L Frontal Orbital Cortex | 8.31 | -34 | 24 | -8 |
| | L Caudate | 3.79 | -16 | 18 | 4 |
| | L Frontal Operculum Cortex | 12.89 | -40 | 10 | 4 |

| | | | | | |
|---|---|---|---|---|---|
| | L Inferior Frontal Gyrus, pars opercularis | 5.48 | -56 | 10 | 14 |
| | L Putamen | 8.25 | -22 | 6 | -4 |
| | L Central Opercular Cortex | 8.1 | -52 | 4 | 4 |
| 18,064 | | | | | |
| | R Parietal Operculum Cortex | 5.87 | 46 | -22 | 18 |
| | R Middle Temporal Gyrus, posterior division | 8.33 | 52 | -30 | -4 |
| | R Middle Temporal Gyrus, temporooccipital part | 5.75 | 48 | -40 | 4 |
| | R Supramarginal Gyrus, posterior division | 10.15 | 66 | -42 | 28 |
| | R Angular Gyrus | 8.98 | 52 | -46 | 34 |
| 8,912 | | | | | |
| | L Frontal Pole | 9.63 | -28 | 48 | 26 |
| | L Middle Frontal Gyrus | 6.29 | -36 | 34 | 40 |
| 8,688 | | | | | |
| | L Postcentral Gyrus | 4.34 | -56 | -22 | 24 |
| | L Parietal Operculum Cortex | 5.45 | -56 | -32 | 26 |
| | L Supramarginal Gyrus, anterior division | 5.61 | -66 | -38 | 26 |
| | L Supramarginal Gyrus, posterior division | 7.83 | -60 | -48 | 34 |
| 6,440 | | | | | |
| | R Frontal Pole | 7.43 | 32 | 44 | 28 |
| | R Middle Frontal Gyrus | 4.74 | 36 | 34 | 38 |
| 4,112 | | | | | |
| | R Postcentral Gyrus | 2.49 | 36 | -36 | 52 |
| | R Superior Parietal Lobule | 6.92 | 28 | -44 | 70 |
| 2,552 | | | | | |
| | R Occipital Pole | 5.73 | 24 | -96 | 14 |
| 2,088 | | | | | |
| | R Temporal Pole | 4.61 | 44 | 6 | -36 |
| 1,000 | | | | | |
| | L Temporal Pole | 4.48 | -44 | 6 | -44 |
| | L Inferior Temporal Gyrus, anterior division | 3.66 | -44 | 0 | -36 |
| | L Middle Temporal Gyrus, anterior division | 3.52 | -48 | -2 | -32 |
| 944 | | | | | |
| | L Superior Parietal Lobule | 5.06 | -22 | -50 | 64 |
| 760 | | | | | |
| | R Postcentral Gyrus | 4.01 | 48 | -26 | 50 |
| 448 | | | | | |
| | L Occipital Pole | 3.63 | -20 | -100 | 8 |
| 328 | | | | | |
| | R Frontal Pole | 3.67 | 28 | 58 | -8 |
| 288 | | | | | |

| | | | | | |
|---|---|---|---|---|---|
| | L Pallidum | 4.05 | -18 | -8 | -2 |
| 240 | | | | | |
| | L Hippocampus | 3.61 | -34 | -28 | -8 |
| 232 | | | | | |
| | R Occipital Fusiform Gyrus | 3.31 | 14 | -86 | -12 |
| 224 | | | | | |
| | R Hippocampus | 3.6 | 36 | -34 | -6 |
| 208 | | | | | |
| | L Cingulate Gyrus, posterior division | 3.66 | -10 | -22 | 40 |
| 184 | | | | | |
| | R Inferior Frontal Gyrus, pars opercularis | 2.75 | 38 | 14 | 26 |
| 128 | | | | | |
| | R Occipital Pole | 3.17 | 8 | -100 | -6 |
| 112 | | | | | |
| | L Lingual Gyrus | 3.05 | -10 | -86 | -10 |
| 72 | | | | | |
| | L Frontal Pole | 3.43 | -32 | 58 | -12 |
| 72 | | | | | |
| | L Caudate | 2.84 | -18 | -16 | 24 |
| 64 | | | | | |
| | R Inferior Temporal Gyrus, posterior division | 2.9 | 50 | -30 | -28 |
| 64 | | | | | |
| | L Amygdala | 2.8 | -24 | -10 | -12 |
| 56 | | | | | |
| | L Temporal Pole | 2.94 | -30 | 22 | -34 |
| 56 | | | | | |
| | L Subcallosal Cortex | 3.88 | -10 | 16 | -22 |
| 48 | | | | | |
| | L Frontal Orbital Cortex | 3.34 | -22 | 18 | -24 |
| 48 | | | | | |
| | R Accumbens | 3.55 | 10 | 6 | -8 |
| 40 | | | | | |
| | R Inferior Temporal Gyrus, posterior division | 2.96 | 56 | -10 | -40 |
| 40 | | | | | |
| | L Hippocampus | 3.43 | -28 | -18 | -12 |
| 40 | | | | | |
| | L Thalamus | 4.05 | -6 | -26 | 14 |
| 40 | | | | | |
| | L Precuneus Cortex | 2.71 | -12 | -46 | 50 |
| 32 | | | | | |
| | L Parahippocampal Gyrus, anterior division | 2.96 | -20 | 0 | -38 |
| 32 | | | | | |

| | | | | | |
|---|---|---|---|---|---|
| | R Brain-Stem | 2.73 | 6 | -22 | -20 |
| 32 | | | | | |
| | L Frontal Pole | 2.68 | -30 | 52 | -14 |
| 32 | | | | | |
| | L Frontal Pole | 2.55 | -30 | 48 | 8 |
| 32 | | | | | |
| | R Thalamus | 3.66 | 8 | -30 | 12 |
| 32 | | | | | |
| | L Superior Frontal Gyrus | 2.71 | -6 | 28 | 52 |
| 32 | | | | | |
| | L Superior Parietal Lobule | 3.21 | -14 | -54 | 72 |
| 24 | | | | | |
| | L Temporal Pole | 2.54 | -36 | 22 | -38 |
| 24 | | | | | |
| | R Middle Temporal Gyrus, anterior division | 2.71 | 58 | 0 | -36 |
| 24 | | | | | |
| | L Frontal Orbital Cortex | 2.81 | -46 | 32 | -10 |
| 24 | | | | | |
| | R Hippocampus | 4.15 | 34 | -28 | -6 |
| 24 | | | | | |
| | L Frontal Pole | 2.71 | -26 | 68 | -6 |
| 24 | | | | | |
| | L Middle Temporal Gyrus, temporooccipital part | 2.73 | -50 | -50 | 4 |
| 24 | | | | | |
| | L Postcentral Gyrus | 2.6 | -44 | -28 | 40 |
| 24 | | | | | |
| | R Postcentral Gyrus | 2.72 | 32 | -34 | 46 |
| 24 | | | | | |
| | L Superior Parietal Lobule | 2.61 | -42 | -52 | 58 |
| 16 | | | | | |
| | R Inferior Temporal Gyrus, anterior division | 2.87 | 48 | -4 | -42 |
| 16 | | | | | |
| | R Temporal Fusiform Cortex, anterior division | 2.87 | 26 | -4 | -42 |
| 16 | | | | | |
| | R Parahippocampal Gyrus, anterior division | 3.06 | 24 | -10 | -32 |
| 16 | | | | | |
| | R Putamen | 3.94 | 32 | -18 | -2 |
| 16 | | | | | |
| | R Thalamus | 2.47 | 8 | -8 | -2 |
| 16 | | | | | |

| | | | | | |
|---|---|---|---|---|---|
| | R Putamen | 2.57 | 30 | -18 | 8 |
| 16 | | | | | |
| | R Thalamus | 2.89 | 12 | -34 | 8 |
| 16 | | | | | |
| | L Precentral Gyrus | 3 | -50 | 2 | 28 |
| 8 | | | | | |
| | R Parahippocampal Gyrus, anterior division | 2.69 | 22 | -8 | -36 |
| 8 | | | | | |
| | R Inferior Temporal Gyrus, anterior division | 2.4 | 54 | -6 | -34 |
| 8 | | | | | |
| | L Temporal Pole | 3.32 | -32 | 12 | -34 |
| 8 | | | | | |
| | R Inferior Temporal Gyrus, temporooccipital part | 2.62 | 58 | -42 | -20 |
| 8 | | | | | |
| | R Lateral Occipital Cortex, inferior division | 2.4 | 38 | -88 | -12 |
| 8 | | | | | |
| | R Thalamus | 2.36 | 22 | -34 | 0 |
| 8 | | | | | |
| | L Pallidum | 2.46 | -12 | 2 | 0 |
| 8 | | | | | |
| | L Paracingulate Gyrus | 2.38 | -10 | 54 | 12 |
| 8 | | | | | |
| | R Supramarginal Gyrus, anterior division | 2.58 | 52 | -22 | 32 |
| 8 | | | | | |
| | R Superior Frontal Gyrus | 2.36 | 2 | 54 | 36 |
| 8 | | | | | |
| | L Superior Parietal Lobule | 2.36 | -30 | -46 | 50 |
| 8 | | | | | |
| | L Bed Nucleus of the Stria Terminalis | 5.68 | -6 | 2 | -4 |

**Supplementary Table S7. [$CT_{sustained}$ > $CS_{sustained}$].** Descriptive statistics for clusters and local maxima showing greater activation for Certain-Threat relative to Certain-Safety anticipation for the OSP Sustained regressor (FDR $q$<0.05, whole-brain corrected).

| *mm³* | Label | *t* | *x* | *y* | *z* |
|---|---|---|---|---|---|
| 14,880 | | | | | |
| | L Cingulate Gyrus, anterior division | 4.82 | -8 | 24 | 30 |
| | L Paracingulate Gyrus | 5.78 | -8 | 20 | 34 |
| | R Paracingulate Gyrus | 5.18 | 10 | 18 | 36 |
| | L Juxtapositional Lobule Cortex (formerly Supplementary Motor Cortex) | 5.37 | -6 | 6 | 58 |
| | R Juxtapositional Lobule Cortex (formerly Supplementary Motor Cortex) | 6.01 | 6 | 6 | 70 |
| | L Superior Frontal Gyrus | 6.07 | -16 | 4 | 68 |
| | R Superior Frontal Gyrus | 7.39 | 14 | 4 | 72 |
| | L Middle Frontal Gyrus | 3.21 | -36 | -4 | 62 |
| | L Precentral Gyrus | 3.34 | -32 | -6 | 68 |
| | R Precentral Gyrus | 4.62 | 32 | -8 | 66 |
| 5,264 | | | | | |
| | L Postcentral Gyrus | 3.43 | -40 | -32 | 48 |
| | L Superior Parietal Lobule | 3.38 | -34 | -46 | 46 |
| | L Supramarginal Gyrus, posterior division | 5.58 | -56 | -52 | 38 |
| | L Angular Gyrus | 4.27 | -44 | -54 | 56 |
| | L Lateral Occipital Cortex, superior division | 3.99 | -34 | -60 | 54 |
| 4,864 | | | | | |
| | R Frontal Pole | 4.65 | 50 | 34 | -4 |
| | R Frontal Orbital Cortex | 4.50 | 38 | 30 | -2 |
| | R Inferior Frontal Gyrus, pars triangularis | 3.73 | 54 | 24 | 6 |
| | R Frontal Operculum Cortex | 6.78 | 48 | 18 | 0 |
| | R Inferior Frontal Gyrus, pars opercularis | 3.60 | 56 | 16 | 6 |
| | R Central Opercular Cortex | 4.54 | 50 | 8 | 2 |
| 4,336 | | | | | |
| | L Frontal Pole | 6.63 | -40 | 54 | 14 |
| | L Middle Frontal Gyrus | 4.10 | -34 | 28 | 38 |
| 3,968 | | | | | |
| | R Postcentral Gyrus | 3.68 | 44 | -28 | 48 |
| | R Supramarginal Gyrus, anterior division | 4.10 | 46 | -32 | 44 |
| | R Supramarginal Gyrus, posterior division | 5.01 | 64 | -42 | 28 |
| | R Superior Parietal Lobule | 4.14 | 44 | -46 | 58 |
| | R Angular Gyrus | 3.82 | 50 | -48 | 34 |
| 3,168 | | | | | |
| | R Frontal Pole | 5.36 | 32 | 52 | 28 |
| 2,408 | | | | | |
| | L Inferior Frontal Gyrus, pars triangularis | 3.45 | -52 | 20 | -2 |
| | L Frontal Operculum Cortex | 6.77 | -38 | 14 | 6 |

| | | | | | |
|---|---|---|---|---|---|
| | L Inferior Frontal Gyrus, pars opercularis | 3.67 | -52 | 14 | 2 |
| 1,376 | | | | | |
| | R Middle Frontal Gyrus | 3.33 | 46 | 8 | 46 |
| | R Precentral Gyrus | 4.95 | 50 | 2 | 52 |
| 1,288 | | | | | |
| | R Superior Temporal Gyrus, posterior division | 4.86 | 48 | -26 | -4 |
| | R Middle Temporal Gyrus, posterior division | 4.15 | 50 | -28 | -8 |
| | R Supramarginal Gyrus, posterior division | 3.40 | 46 | -38 | 8 |
| | R Middle Temporal Gyrus, temporooccipital part | 4.59 | 52 | -42 | 4 |
| 968 | | | | | |
| | R Brain-Stem | 5.75 | 6 | -46 | -60 |
| | L Brain-Stem | 3.54 | -6 | -48 | -58 |
| 544 | | | | | |
| | L Precentral Gyrus | 3.98 | -54 | 0 | 42 |
| 432 | | | | | |
| | R Caudate | 6.01 | 18 | -6 | 24 |
| 248 | | | | | |
| | R Putamen | 3.61 | 18 | 8 | -8 |
| 240 | | | | | |
| | R Superior Frontal Gyrus | 3.86 | 12 | 28 | 58 |
| 216 | | | | | |
| | L Frontal Pole | 3.74 | -44 | 56 | -8 |
| 192 | | | | | |
| | L Brain-Stem | 3.87 | -4 | -30 | -54 |
| 192 | | | | | |
| | L Caudate | 4.86 | -16 | 2 | 22 |
| 176 | | | | | |
| | R Frontal Pole | 4.01 | 32 | 60 | -6 |
| 176 | | | | | |
| | R Thalamus | 4.40 | 2 | -18 | 0 |
| 160 | | | | | |
| | R Brain-Stem | 4.73 | 10 | -32 | -16 |
| 160 | | | | | |
| | L Putamen | 3.84 | -22 | 14 | -4 |
| 160 | | | | | |
| | R Occipital Pole | 4.23 | 22 | -102 | 12 |
| 152 | | | | | |
| | R Amygdala | 3.53 | 30 | -4 | -20 |
| | R Amygdala | 4.26 | 30 | -8 | -16 |
| 144 | | | | | |
| | L Frontal Orbital Cortex | 4.13 | -34 | 30 | -2 |
| 144 | | | | | |
| | R Frontal Pole | 3.82 | 52 | 46 | 0 |

| | | | | | |
|---|---|---|---|---|---|
| 136 | | | | | |
| | L Frontal Pole | 4.10 | -50 | 42 | -2 |
| 104 | | | | | |
| | L Periaqueductal Grey | 4.38 | -2 | -36 | -16 |
| 104 | | | | | |
| | R Frontal Pole | 3.83 | 18 | 54 | 30 |
| 88 | | | | | |
| | L Brain-Stem | 4.18 | -6 | -44 | -64 |
| 88 | | | | | |
| | R Brain-Stem | 3.77 | 0 | -28 | -4 |
| 88 | | | | | |
| | L Precentral Gyrus | 3.48 | -28 | -8 | 48 |
| 80 | | | | | |
| | L Brain-Stem | 3.36 | -6 | -38 | -62 |
| 80 | | | | | |
| | R Thalamus | 3.30 | 22 | -34 | -2 |
| | R Hippocampus | 4.46 | 20 | -36 | 6 |
| 72 | | | | | |
| | R Superior Temporal Gyrus, posterior division | 3.66 | 56 | -20 | -4 |
| 64 | | | | | |
| | L Frontal Pole | 3.72 | -32 | 48 | -16 |
| 64 | | | | | |
| | R Accumbens | 3.05 | 6 | 6 | -6 |
| 64 | | | | | |
| | R Putamen | 3.20 | 24 | 16 | -6 |
| 64 | | | | | |
| | R Putamen | 3.81 | 24 | 14 | 4 |
| 64 | | | | | |
| | L Caudate | 4.26 | -18 | 16 | 12 |
| 56 | | | | | |
| | L Temporal Fusiform Cortex, anterior division | 4.38 | -32 | 0 | -50 |
| 56 | | | | | |
| | R Thalamus | 4.12 | 0 | -4 | 6 |
| 56 | | | | | |
| | R Thalamus | 4.14 | 6 | -30 | 8 |
| 56 | | | | | |
| | R Frontal Pole | 3.52 | 34 | 36 | 40 |
| 48 | | | | | |
| | L Amygdala | 4.05 | -24 | -12 | -10 |
| 48 | | | | | |
| | R Putamen | 3.64 | 20 | 6 | 6 |
| 48 | | | | | |
| | L Frontal Operculum Cortex | 3.68 | -36 | 26 | 8 |
| 48 | | | | | |

| | | | | | |
|---|---|---|---|---|---|
| | L Superior Frontal Gyrus | 3.31 | -2 | 18 | 62 |
| 40 | | | | | |
| | L Brain-Stem | 3.33 | -6 | -38 | -30 |
| 40 | | | | | |
| | L Putamen | 3.04 | -16 | 8 | -10 |
| 40 | | | | | |
| | R Thalamus | 4.87 | 12 | -34 | 8 |
| 40 | | | | | |
| | R Superior Parietal Lobule | 3.61 | 38 | -48 | 68 |
| 40 | | | | | |
| | L Superior Parietal Lobule | 3.20 | -34 | -46 | 68 |
| 32 | | | | | |
| | L Brain-Stem | 3.17 | -14 | -26 | -14 |
| 32 | | | | | |
| | R Temporal Pole | 3.25 | 50 | 20 | -16 |
| 32 | | | | | |
| | R Lateral Occipital Cortex, inferior division | 3.40 | 48 | -72 | -12 |
| 32 | | | | | |
| | L Frontal Pole | 3.27 | -50 | 46 | -12 |
| 32 | | | | | |
| | R Hippocampus | 3.71 | 36 | -34 | -6 |
| 32 | | | | | |
| | L Postcentral Gyrus | 2.89 | -48 | -28 | 40 |
| | L Supramarginal Gyrus, anterior division | 2.89 | -50 | -30 | 38 |
| 24 | | | | | |
| | R Frontal Pole | 3.30 | 24 | 60 | -12 |
| 24 | | | | | |
| | R Middle Temporal Gyrus, posterior division | 2.97 | 60 | -24 | -8 |
| 24 | | | | | |
| | R Thalamus | 3.95 | 0 | -20 | 10 |
| 24 | | | | | |
| | L Cingulate Gyrus, anterior division | 3.22 | -4 | 24 | 20 |
| 24 | | | | | |
| | R Angular Gyrus | 2.84 | 52 | -52 | 48 |
| 16 | | | | | |
| | L Brain-Stem | 3.08 | -2 | -32 | -60 |
| 16 | | | | | |
| | R Brain-Stem | 3.35 | 2 | -30 | -54 |
| 16 | | | | | |
| | R Temporal Pole | 3.21 | 30 | 24 | -36 |
| 16 | | | | | |
| | R Inferior Temporal Gyrus, posterior division | 2.94 | 58 | -36 | -20 |
| 16 | | | | | |

| | | | | | |
|---|---|---|---|---|---|
| | L Inferior Temporal Gyrus, posterior division | 3.59 | -54 | -36 | -18 |
| 16 | | | | | |
| | L Brain-Stem | 3.39 | -10 | -34 | -14 |
| 16 | | | | | |
| | R Frontal Orbital Cortex | 2.86 | 30 | 22 | -14 |
| 16 | | | | | |
| | L Frontal Pole | 3.23 | -42 | 54 | -14 |
| 16 | | | | | |
| | L Pallidum | 3.17 | -14 | -2 | -4 |
| 16 | | | | | |
| | L Caudate | 2.82 | -18 | 20 | -2 |
| 16 | | | | | |
| | R Caudate | 3.02 | 18 | 14 | 12 |
| 16 | | | | | |
| | R Caudate | 2.98 | 12 | 0 | 20 |
| 16 | | | | | |
| | R Supramarginal Gyrus, posterior division | 2.89 | 40 | -40 | 38 |
| 8 | | | | | |
| | L Temporal Pole | 2.91 | -38 | 22 | -40 |
| 8 | | | | | |
| | L Temporal Pole | 2.79 | -34 | 14 | -34 |
| 8 | | | | | |
| | R Brain-Stem | 2.78 | 6 | -42 | -26 |
| 8 | | | | | |
| | L Brain-Stem | 2.83 | -10 | -38 | -26 |
| 8 | | | | | |
| | L Subcallosal Cortex | 2.88 | -8 | 28 | -24 |
| 8 | | | | | |
| | L Brain-Stem | 2.85 | -12 | -26 | -20 |
| 8 | | | | | |
| | R Inferior Temporal Gyrus, temporooccipital part | 2.88 | 62 | -46 | -16 |
| 8 | | | | | |
| | L Frontal Orbital Cortex | 2.91 | -36 | 22 | -16 |
| 8 | | | | | |
| | L Hippocampus | 2.86 | -28 | -14 | -14 |
| 8 | | | | | |
| | R Frontal Pole | 2.81 | 42 | 56 | -10 |
| 8 | | | | | |
| | R Hippocampus | 2.86 | 34 | -26 | -8 |
| 8 | | | | | |
| | L Middle Temporal Gyrus, posterior division | 3.03 | -54 | -32 | -6 |
| 8 | | | | | |
| | R Insular Cortex | 2.80 | 38 | 10 | -6 |

| | | | | | |
|---|---|---|---|---|---|
| 8 | | | | | |
| | L Frontal Orbital Cortex | 2.89 | -32 | 26 | -6 |
| 8 | | | | | |
| | L Lateral Occipital Cortex, inferior division | 3.09 | -42 | -80 | -4 |
| 8 | | | | | |
| | R Hippocampus | 3.23 | 24 | -32 | -4 |
| 8 | | | | | |
| | L Pallidum | 3.14 | -20 | -10 | -4 |
| 8 | | | | | |
| | L Middle Temporal Gyrus, posterior division | 2.97 | -52 | -42 | -2 |
| 8 | | | | | |
| | L Pallidum | 2.92 | -16 | -6 | -2 |
| 8 | | | | | |
| | L Thalamus | 2.85 | -2 | -2 | 0 |
| 8 | | | | | |
| | L Thalamus | 3.12 | -20 | -36 | 2 |
| 8 | | | | | |
| | R Occipital Pole | 2.84 | 32 | -96 | 6 |
| 8 | | | | | |
| | L Thalamus | 2.82 | -12 | -10 | 6 |
| 8 | | | | | |
| | L Frontal Pole | 2.78 | -32 | 64 | 6 |
| 8 | | | | | |
| | L Supramarginal Gyrus, posterior division | 3.33 | -54 | -44 | 20 |
| 8 | | | | | |
| | R Precentral Gyrus | 3.36 | 54 | 4 | 20 |
| 8 | | | | | |
| | R Caudate | 2.89 | 18 | -20 | 22 |
| 8 | | | | | |
| | L Middle Frontal Gyrus | 3.29 | -40 | 30 | 26 |
| 8 | | | | | |
| | L Paracingulate Gyrus | 3.01 | -6 | 20 | 46 |
| 8 | | | | | |
| | L Superior Frontal Gyrus | 2.99 | -14 | 30 | 52 |
| 8 | | | | | |
| | R Frontal Pole | 2.80 | 10 | 40 | 56 |
| 8 | | | | | |
| | R Precentral Gyrus | 2.79 | 28 | -12 | 60 |
| 8 | | | | | |
| | R Superior Parietal Lobule | 3.06 | 28 | -48 | 62 |
| 8 | | | | | |
| | R Precentral Gyrus | 2.87 | 40 | -20 | 62 |
| 8 | | | | | |
| | R Superior Frontal Gyrus | 3.01 | 24 | 8 | 64 |
| 8 | | | | | |

| | | | | | |
|---|---|---|---|---|---|
| | R Lateral Occipital Cortex, superior division | 2.83 | 8 | -66 | 66 |
| 8 | | | | | |
| | R Superior Parietal Lobule | 2.83 | 14 | -52 | 76 |

**Supplementary Table S8. [$UT_{sustained}$ > $US_{sustained}$] ∩ [$CT_{sustained}$ > $CS_{sustained}$].** Descriptive statistics for clusters and local maxima greater activation for Uncertain-Threat relative to Uncertain-Safety anticipation for the OSP Sustained regressor AND greater sustained activation for Certain-Threat relative to Certain-Safety anticipation for the OSP Sustained regressor (FDR $q$<0.05, whole-brain corrected).

| mm³ | Label | *t* | *x* | *y* | *z* |
|---|---|---|---|---|---|
| 14,616 | | | | | |
| | L Cingulate Gyrus, anterior division | 4.82 | -8 | 24 | 30 |
| | L Paracingulate Gyrus | 5.78 | -8 | 20 | 34 |
| | R Paracingulate Gyrus | 5.18 | 10 | 18 | 36 |
| | L Juxtapositional Lobule Cortex (formerly Supplementary Motor Cortex) | 5.28 | -4 | 6 | 58 |
| | R Juxtapositional Lobule Cortex (formerly Supplementary Motor Cortex) | 6.01 | 6 | 6 | 70 |
| | L Superior Frontal Gyrus | 6.07 | -16 | 4 | 68 |
| | R Superior Frontal Gyrus | 7.39 | 14 | 4 | 72 |
| | L Middle Frontal Gyrus | 3.21 | -36 | -4 | 62 |
| | L Precentral Gyrus | 3.34 | -32 | -6 | 68 |
| | R Precentral Gyrus | 4.62 | 32 | -8 | 66 |
| 4,856 | | | | | |
| | R Frontal Pole | 4.63 | 50 | 34 | -4 |
| | R Frontal Orbital Cortex | 4.50 | 38 | 30 | -2 |
| | R Inferior Frontal Gyrus, pars triangularis | 3.73 | 54 | 24 | 6 |
| | R Frontal Operculum Cortex | 6.78 | 48 | 18 | 0 |
| | R Inferior Frontal Gyrus, pars opercularis | 3.60 | 56 | 16 | 6 |
| | R Central Opercular Cortex | 4.54 | 50 | 8 | 2 |
| 3,616 | | | | | |
| | L Frontal Pole | 5.60 | -34 | 54 | 24 |
| | L Middle Frontal Gyrus | 4.10 | -34 | 28 | 38 |
| 2,624 | | | | | |
| | R Frontal Pole | 5.36 | 32 | 52 | 28 |
| 2,384 | | | | | |
| | L Inferior Frontal Gyrus, pars triangularis | 3.45 | -52 | 20 | -2 |
| | L Frontal Operculum Cortex | 6.77 | -38 | 14 | 6 |
| | L Inferior Frontal Gyrus, pars opercularis | 3.67 | -52 | 14 | 2 |
| 2,168 | | | | | |
| | L Supramarginal Gyrus, posterior division | 5.36 | -56 | -52 | 38 |
| | L Angular Gyrus | 3.25 | -52 | -56 | 50 |
| 1,448 | | | | | |
| | R Supramarginal Gyrus, posterior division | 5.01 | 64 | -42 | 28 |
| | R Angular Gyrus | 3.82 | 50 | -48 | 34 |
| 1,376 | | | | | |
| | R Middle Frontal Gyrus | 3.33 | 46 | 8 | 46 |
| | R Precentral Gyrus | 4.95 | 50 | 2 | 52 |
| 1,248 | | | | | |

| | | | | | |
|---|---|---|---|---|---|
| | R Superior Temporal Gyrus, posterior division | 4.86 | 48 | -26 | -4 |
| | R Middle Temporal Gyrus, posterior division | 4.15 | 50 | -28 | -8 |
| | R Supramarginal Gyrus, posterior division | 3.38 | 48 | -38 | 8 |
| 816 | | | | | |
| | R Brain-Stem | 5.20 | 6 | -48 | -60 |
| 432 | | | | | |
| | R Caudate | 6.01 | 18 | -6 | 24 |
| 328 | | | | | |
| | R Postcentral Gyrus | 3.28 | 48 | -24 | 42 |
| 272 | | | | | |
| | L Precentral Gyrus | 3.72 | -48 | 2 | 40 |
| 232 | | | | | |
| | R Putamen | 3.61 | 18 | 8 | -8 |
| 216 | | | | | |
| | R Postcentral Gyrus | 3.93 | 38 | -38 | 66 |
| | R Superior Parietal Lobule | 2.52 | 38 | -44 | 62 |
| 184 | | | | | |
| | L Brain-Stem | 3.87 | -4 | -30 | -54 |
| 176 | | | | | |
| | L Caudate | 4.86 | -16 | 2 | 22 |
| 160 | | | | | |
| | R Occipital Pole | 4.03 | 22 | -102 | 12 |
| 152 | | | | | |
| | L Putamen | 3.84 | -22 | 14 | -4 |
| 144 | | | | | |
| | L Frontal Orbital Cortex | 4.13 | -34 | 30 | -2 |
| 136 | | | | | |
| | R Superior Frontal Gyrus | 3.61 | 10 | 30 | 56 |
| 104 | | | | | |
| | L Brain-Stem | 3.54 | -6 | -48 | -58 |
| 104 | | | | | |
| | R Brain-Stem | 4.58 | 12 | -28 | -14 |
| 88 | | | | | |
| | L Brain-Stem | 4.18 | -6 | -44 | -64 |
| 88 | | | | | |
| | R Brain-Stem | 3.77 | 0 | -28 | -4 |
| 80 | | | | | |
| | L Brain-Stem | 3.36 | -6 | -38 | -62 |
| 80 | | | | | |
| | L Precentral Gyrus | 3.48 | -28 | -8 | 48 |
| 64 | | | | | |
| | R Superior Temporal Gyrus, posterior division | 3.66 | 56 | -20 | -4 |
| 64 | | | | | |

| | | | | | |
|---|---|---|---|---|---|
| | R Thalamus | 4.24 | 2 | -20 | 0 |
| 64 | | | | | |
| | R Putamen | 3.81 | 24 | 14 | 4 |
| 56 | | | | | |
| | R Thalamus | 4.12 | 0 | -4 | 6 |
| 48 | | | | | |
| | R Brain-Stem | 3.24 | 0 | -36 | -14 |
| 48 | | | | | |
| | R Putamen | 2.99 | 20 | 14 | -4 |
| 48 | | | | | |
| | R Putamen | 3.64 | 20 | 6 | 6 |
| 48 | | | | | |
| | L Frontal Operculum Cortex | 3.68 | -36 | 26 | 8 |
| 40 | | | | | |
| | R Brain-Stem | 3.97 | 12 | -16 | -22 |
| 40 | | | | | |
| | R Frontal Pole | 3.52 | 34 | 36 | 40 |
| 40 | | | | | |
| | R Superior Parietal Lobule | 3.50 | 38 | -46 | 68 |
| 32 | | | | | |
| | L Brain-Stem | 3.17 | -14 | -26 | -14 |
| 32 | | | | | |
| | L Amygdala | 2.80 | -24 | -10 | -12 |
| 32 | | | | | |
| | L Caudate | 3.56 | -18 | 16 | 12 |
| 32 | | | | | |
| | L Superior Frontal Gyrus | 2.86 | -4 | 20 | 64 |
| 24 | | | | | |
| | R Amygdala | 3.21 | 28 | -12 | -14 |
| 24 | | | | | |
| | L Putamen | 2.94 | -16 | 10 | -8 |
| 24 | | | | | |
| | R Middle Temporal Gyrus, posterior division | 2.97 | 60 | -24 | -8 |
| 16 | | | | | |
| | L Brain-Stem | 3.08 | -2 | -32 | -60 |
| 16 | | | | | |
| | R Brain-Stem | 3.35 | 2 | -30 | -54 |
| 16 | | | | | |
| | L Brain-Stem | 3.09 | -4 | -42 | -26 |
| 16 | | | | | |
| | R Frontal Orbital Cortex | 2.86 | 30 | 22 | -14 |
| 16 | | | | | |
| | R Accumbens | 2.96 | 8 | 6 | -8 |
| 16 | | | | | |
| | R Hippocampus | 3.60 | 36 | -34 | -6 |

| | | | | | |
|---|---|---|---|---|---|
| 16 | | | | | |
| | R Frontal Pole | 2.84 | 32 | 58 | -6 |
| 16 | | | | | |
| | L Pallidum | 2.81 | -14 | -4 | -4 |
| 16 | | | | | |
| | R Thalamus | 2.89 | 12 | -34 | 8 |
| 16 | | | | | |
| | R Caudate | 3.02 | 18 | 14 | 12 |
| 16 | | | | | |
| | R Caudate | 2.98 | 12 | 0 | 20 |
| 16 | | | | | |
| | L Superior Parietal Lobule | 2.61 | -42 | -52 | 58 |
| 8 | | | | | |
| | L Brain-Stem | 3.33 | -6 | -38 | -30 |
| 8 | | | | | |
| | R Brain-Stem | 2.78 | 6 | -42 | -26 |
| 8 | | | | | |
| | L Brain-Stem | 2.85 | -12 | -26 | -20 |
| 8 | | | | | |
| | L Frontal Orbital Cortex | 2.89 | -32 | 26 | -6 |
| 8 | | | | | |
| | L Pallidum | 3.14 | -20 | -10 | -4 |
| 8 | | | | | |
| | L Pallidum | 2.92 | -16 | -6 | -2 |
| 8 | | | | | |
| | L Caudate | 2.68 | -18 | 18 | -2 |
| 8 | | | | | |
| | R Thalamus | 2.36 | 22 | -34 | 0 |
| 8 | | | | | |
| | L Thalamus | 2.82 | -12 | -10 | 6 |
| 8 | | | | | |
| | R Thalamus | 2.83 | 6 | -30 | 10 |
| 8 | | | | | |
| | R Caudate | 2.86 | 18 | -20 | 22 |
| 8 | | | | | |
| | R Precentral Gyrus | 2.79 | 28 | -12 | 60 |
| 8 | | | | | |
| | R Superior Parietal Lobule | 2.92 | 28 | -48 | 62 |
| 8 | | | | | |
| | R Superior Frontal Gyrus | 3.01 | 24 | 8 | 64 |
| 8 | | | | | |
| | L Superior Parietal Lobule | 2.78 | -34 | -48 | 68 |

**Supplementary Table S9. [$UT_{sustained}$ > $CT_{sustained}$].** Descriptive statistics for clusters and local maxima showing greater sustained activation for Uncertain-Threat relative to Certain-Threat anticipation for the OSP Sustained regressor (FDR *q*<0.05, whole-brain corrected).

| ***mm³*** | **Label** | ***t*** | ***x*** | ***y*** | ***z*** |
|---|---|---|---|---|---|
| 62,312 | | | | | |
| | L Paracingulate Gyrus | 7.49 | -4 | 34 | 26 |
| | R Paracingulate Gyrus | 11.43 | 2 | 18 | 38 |
| | R Cingulate Gyrus, anterior division | 11.87 | 0 | 10 | 42 |
| | R Middle Frontal Gyrus | 6.99 | 48 | 10 | 50 |
| | L Cingulate Gyrus, anterior division | 9.79 | -8 | 6 | 40 |
| | R Juxtapositional Lobule Cortex (formerly Supplementary Motor Cortex) | 12.01 | 6 | 6 | 48 |
| | R Precentral Gyrus | 11.49 | 42 | -2 | 54 |
| | L Precentral Gyrus | 5.67 | -36 | -4 | 48 |
| | L Superior Frontal Gyrus | 10.40 | -12 | -8 | 68 |
| | R Superior Frontal Gyrus | 11.88 | 18 | -10 | 70 |
| | L Cingulate Gyrus, posterior division | 5.64 | -2 | -18 | 28 |
| | R Cingulate Gyrus, posterior division | 7.64 | 8 | -22 | 46 |
| 34,376 | | | | | |
| | R Inferior Temporal Gyrus, temporooccipital part | 3.81 | 46 | -44 | -14 |
| | R Temporal Occipital Fusiform Cortex | 5.21 | 36 | -48 | -16 |
| | R Middle Temporal Gyrus, temporooccipital part | 2.66 | 54 | -52 | 4 |
| | R Lateral Occipital Cortex, inferior division | 10.89 | 42 | -82 | -14 |
| | R Occipital Fusiform Gyrus | 10.86 | 30 | -82 | -14 |
| | R Occipital Pole | 11.63 | 26 | -96 | 2 |
| 25,008 | | | | | |
| | R Frontal Pole | 4.30 | 54 | 38 | -6 |
| | R Frontal Orbital Cortex | 14.02 | 34 | 28 | 2 |
| | R Frontal Operculum Cortex | 12.74 | 42 | 24 | 4 |
| | R Inferior Frontal Gyrus, pars triangularis | 9.28 | 54 | 22 | 2 |
| | R Inferior Frontal Gyrus, pars opercularis | 8.87 | 56 | 14 | 0 |
| | R Central Opercular Cortex | 9.74 | 54 | 4 | 4 |
| | R Precentral Gyrus | 9.37 | 58 | 4 | 10 |
| | R Putamen | 10.99 | 26 | 2 | -6 |
| | R Pallidum | 3.66 | 16 | 2 | 0 |
| 22,848 | | | | | |
| | L Accumbens | 2.93 | -6 | 8 | -10 |
| | L Caudate | 5.10 | -8 | 0 | 10 |
| | R Thalamus | 6.96 | 12 | -4 | 14 |
| | L Thalamus | 5.12 | -6 | -6 | 8 |
| | R Caudate | 5.01 | 16 | -8 | 20 |
| | L Brain-Stem | 5.31 | -6 | -30 | -8 |
| | R Brain-Stem | 7.03 | 6 | -30 | -8 |

| | | | | | |
|---|---|---|---|---|---|
| | R Lingual Gyrus | 2.57 | 2 | -74 | -8 |
| 20,160 | | | | | |
| | L Frontal Orbital Cortex | 9.12 | -30 | 22 | -8 |
| | L Frontal Operculum Cortex | 13.62 | -36 | 18 | 8 |
| | L Inferior Frontal Gyrus, pars opercularis | 4.24 | -58 | 12 | 16 |
| | L Precentral Gyrus | 7.34 | -56 | 8 | 8 |
| | L Putamen | 9.11 | -24 | 6 | -6 |
| | L Insular Cortex | 7.20 | -40 | 4 | 4 |
| | L Central Opercular Cortex | 8.21 | -46 | 0 | 8 |
| 12,160 | | | | | |
| | R Parietal Operculum Cortex | 8.36 | 46 | -20 | 18 |
| | R Supramarginal Gyrus, posterior division | 8.44 | 66 | -44 | 30 |
| | R Angular Gyrus | 8.51 | 64 | -48 | 32 |
| 11,912 | | | | | |
| | L Occipital Fusiform Gyrus | 6.46 | -20 | -86 | -10 |
| | L Lateral Occipital Cortex, inferior division | 5.39 | -32 | -88 | -12 |
| | L Occipital Pole | 8.80 | -12 | -94 | -8 |
| 7,744 | | | | | |
| | L Parietal Operculum Cortex | 5.66 | -56 | -32 | 26 |
| | L Supramarginal Gyrus, anterior division | 5.29 | -58 | -36 | 30 |
| | L Supramarginal Gyrus, posterior division | 7.09 | -64 | -44 | 36 |
| | L Angular Gyrus | 2.67 | -46 | -54 | 46 |
| 5,816 | | | | | |
| | L Frontal Pole | 7.25 | -36 | 46 | 26 |
| | L Middle Frontal Gyrus | 5.20 | -40 | 36 | 36 |
| 4,200 | | | | | |
| | R Postcentral Gyrus | 6.58 | 34 | -38 | 62 |
| | R Superior Parietal Lobule | 7.70 | 24 | -44 | 64 |
| 2,736 | | | | | |
| | R Frontal Pole | 4.57 | 34 | 52 | 30 |
| | R Middle Frontal Gyrus | 4.00 | 32 | 34 | 28 |
| 1,472 | | | | | |
| | R Middle Temporal Gyrus, posterior division | 4.95 | 54 | -28 | -4 |
| 1,112 | | | | | |
| | R Temporal Pole | 4.39 | 44 | 6 | -36 |
| | R Middle Temporal Gyrus, anterior division | 3.61 | 48 | 2 | -32 |
| | R Inferior Temporal Gyrus, anterior division | 3.11 | 40 | 2 | -42 |
| 720 | | | | | |
| | L Temporal Pole | 4.53 | -44 | 6 | -42 |
| | L Inferior Temporal Gyrus, anterior division | 3.06 | -50 | -2 | -34 |
| 480 | | | | | |

| | | | | | |
|---|---|---|---|---|---|
| | L Superior Parietal Lobule | 4.68 | -20 | -50 | 66 |
| 328 | | | | | |
| | R Postcentral Gyrus | 3.88 | 46 | -26 | 50 |
| 200 | | | | | |
| | L Cingulate Gyrus, posterior division | 4.22 | -12 | -26 | 40 |
| 184 | | | | | |
| | R Frontal Pole | 4.12 | 12 | 36 | -20 |
| 136 | | | | | |
| | L Frontal Orbital Cortex | 4.05 | -22 | 18 | -26 |
| 112 | | | | | |
| | L Precuneus Cortex | 3.41 | -12 | -68 | 40 |
| 72 | | | | | |
| | L Frontal Pole | 3.35 | -30 | 58 | -12 |
| 56 | | | | | |
| | R Amygdala | 3.40 | 26 | -10 | -12 |
| 56 | | | | | |
| | L Frontal Pole | 3.04 | -22 | 68 | -2 |
| 48 | | | | | |
| | R Frontal Pole | 2.77 | 16 | 34 | -26 |
| 48 | | | | | |
| | L Pallidum | 3.93 | -12 | 2 | 0 |
| 48 | | | | | |
| | L Frontal Pole | 2.96 | -32 | 66 | -2 |
| 40 | | | | | |
| | L Frontal Orbital Cortex | 2.98 | -18 | 10 | -22 |
| 32 | | | | | |
| | L Temporal Fusiform Cortex, anterior division | 2.81 | -36 | -2 | -46 |
| 32 | | | | | |
| | R Brain-Stem | 2.79 | 12 | -14 | -28 |
| 32 | | | | | |
| | R Frontal Pole | 2.70 | 26 | 60 | -8 |
| 32 | | | | | |
| | L Precentral Gyrus | 3.06 | -40 | -12 | 54 |
| 24 | | | | | |
| | R Inferior Temporal Gyrus, posterior division | 3.24 | 56 | -10 | -40 |
| 24 | | | | | |
| | R Brain-Stem | 2.95 | 12 | -42 | -38 |
| 24 | | | | | |
| | L Parahippocampal Gyrus, anterior division | 3.09 | -20 | 0 | -38 |
| 24 | | | | | |
| | L Frontal Pole | 2.94 | -18 | 60 | -20 |
| 24 | | | | | |

| | | | | | |
|---|---|---|---|---|---|
| | L Frontal Orbital Cortex | 3.01 | -22 | 22 | -18 |
| 16 | | | | | |
| | R Inferior Temporal Gyrus, posterior division | 2.70 | 44 | -14 | -42 |
| 16 | | | | | |
| | R Subcallosal Cortex | 2.79 | 8 | 26 | -26 |
| 16 | | | | | |
| | L Brain-Stem | 2.73 | -14 | -18 | -22 |
| 16 | | | | | |
| | R Temporal Pole | 2.82 | 36 | 14 | -22 |
| 16 | | | | | |
| | R Frontal Orbital Cortex | 2.69 | 20 | 24 | -22 |
| 16 | | | | | |
| | R Subcallosal Cortex | 2.67 | 12 | 18 | -14 |
| 16 | | | | | |
| | L Pallidum | 3.23 | -20 | -12 | 0 |
| 16 | | | | | |
| | R Insular Cortex | 3.02 | 38 | -16 | 6 |
| 16 | | | | | |
| | R Thalamus | 2.94 | 4 | -20 | 14 |
| 16 | | | | | |
| | R Caudate | 2.49 | 18 | 10 | 14 |
| 16 | | | | | |
| | L Caudate | 3.14 | -16 | 2 | 24 |
| 16 | | | | | |
| | R Lateral Occipital Cortex, superior division | 2.48 | 44 | -56 | 56 |
| 8 | | | | | |
| | L Temporal Pole | 2.51 | -28 | 12 | -48 |
| 8 | | | | | |
| | R Inferior Temporal Gyrus, anterior division | 2.63 | 48 | -6 | -42 |
| 8 | | | | | |
| | R Inferior Temporal Gyrus, posterior division | 2.55 | 52 | -16 | -38 |
| 8 | | | | | |
| | L Frontal Orbital Cortex | 2.76 | -22 | 30 | -20 |
| 8 | | | | | |
| | L Occipital Fusiform Gyrus | 2.58 | -38 | -68 | -18 |
| 8 | | | | | |
| | R Frontal Pole | 2.52 | 24 | 52 | -18 |
| 8 | | | | | |
| | R Frontal Pole | 2.54 | 26 | 56 | -14 |
| 8 | | | | | |

| | | | | | |
|---|---|---|---|---|---|
| | R Middle Temporal Gyrus, posterior division | 2.44 | 68 | -24 | -10 |
| 8 | | | | | |
| | L Accumbens | 2.55 | -6 | 14 | -6 |
| 8 | | | | | |
| | L Occipital Pole | 2.53 | -20 | -104 | 12 |
| 8 | | | | | |
| | L Frontal Pole | 2.50 | -40 | 48 | 14 |
| 8 | | | | | |
| | L Caudate | 2.65 | -16 | 12 | 16 |
| 8 | | | | | |
| | L Caudate | 2.44 | -16 | 4 | 18 |
| 8 | | | | | |
| | L Caudate | 2.50 | -12 | -8 | 20 |
| 8 | | | | | |
| | R Caudate | 2.98 | 18 | -2 | 20 |
| 8 | | | | | |
| | L Frontal Pole | 2.50 | -42 | 40 | 20 |
| 8 | | | | | |
| | L Middle Frontal Gyrus | 2.43 | -36 | 6 | 46 |
| 8 | | | | | |
| | L Middle Frontal Gyrus | 2.53 | -32 | 6 | 54 |

**Supplementary Table S10. [$US_{sustained} > UT_{sustained}$].** Descriptive statistics for clusters and local maxima showing greater activation for Uncertain-Safety relative to Uncertain-Threat anticipation for the OSP Sustained regressor (FDR $q$<0.05, whole-brain corrected).

| ***mm³*** | **Label** | ***t*** | ***x*** | ***y*** | ***z*** |
|---|---|---|---|---|---|
| 382,568 | | | | | |
| | L Frontal Orbital Cortex | 2.57 | -24 | 14 | -18 |
| | L Temporal Pole | 5.83 | -46 | 12 | -14 |
| | R Temporal Pole | 5.53 | 30 | 6 | -24 |
| | L Amygdala | 7.60 | -24 | 0 | -18 |
| | R Amygdala | 7.55 | 22 | 0 | -18 |
| | L Temporal Fusiform Cortex, anterior division | 4.66 | -30 | -2 | -36 |
| | R Middle Temporal Gyrus, anterior division | 8.39 | 60 | -2 | -18 |
| | L Middle Temporal Gyrus, anterior division | 5.94 | -60 | -4 | -16 |
| | L Superior Temporal Gyrus, anterior division | 5.71 | -60 | -4 | -4 |
| | R Planum Polare | 9.04 | 52 | -4 | -2 |
| | R Temporal Fusiform Cortex, anterior division | 3.94 | 30 | -4 | -38 |
| | R Superior Temporal Gyrus, anterior division | 6.80 | 62 | -6 | -4 |
| | L Insular Cortex | 13.64 | -38 | -8 | 6 |
| | L Planum Polare | 8.77 | -52 | -8 | 2 |
| | R Insular Cortex | 14.53 | 38 | -8 | 12 |
| | R Planum Temporale | 11.25 | 64 | -8 | 4 |
| | R Putamen | 4.60 | 32 | -8 | -8 |
| | L Juxtapositional Lobule Cortex (formerly Supplementary Motor Cortex) | 7.49 | -10 | -10 | 42 |
| | L Pallidum | 4.83 | -26 | -10 | -4 |
| | R Central Opercular Cortex | 14.41 | 38 | -10 | 16 |
| | R Middle Temporal Gyrus, posterior division | 5.19 | 62 | -10 | -26 |
| | L Precentral Gyrus | 15.58 | -46 | -12 | 32 |
| | L Middle Temporal Gyrus, posterior division | 4.05 | -64 | -12 | -18 |
| | R Hippocampus | 9.61 | 18 | -12 | -22 |
| | L Central Opercular Cortex | 16.39 | -38 | -14 | 16 |
| | R Heschls Gyrus (includes H1 and H2) | 10.47 | 54 | -14 | 6 |
| | R Superior Temporal Gyrus, posterior division | 7.48 | 68 | -18 | 8 |
| | L Planum Temporale | 10.16 | -62 | -20 | 10 |
| | L Parahippocampal Gyrus, anterior division | 3.88 | -32 | -22 | -28 |
| | R Supramarginal Gyrus, anterior division | 4.82 | 62 | -22 | 46 |

| | | | | | |
|---|---|---|---|---|---|
| | L Thalamus | 8.18 | -14 | -24 | -2 |
| | L Heschls Gyrus (includes H1 and H2) | 8.43 | -48 | -26 | 8 |
| | L Superior Temporal Gyrus, posterior division | 6.29 | -66 | -26 | 8 |
| | R Precentral Gyrus | 14.47 | 4 | -26 | 62 |
| | R Inferior Temporal Gyrus, posterior division | 4.07 | 44 | -26 | -20 |
| | L Parietal Operculum Cortex | 14.99 | -34 | -28 | 18 |
| | L Supramarginal Gyrus, anterior division | 3.04 | -56 | -30 | 42 |
| | R Thalamus | 7.42 | 16 | -30 | -4 |
| | R Parahippocampal Gyrus, posterior division | 11.26 | 28 | -32 | -18 |
| | L Temporal Fusiform Cortex, posterior division | 10.45 | -30 | -34 | -18 |
| | R Postcentral Gyrus | 16.04 | 2 | -34 | 64 |
| | L Postcentral Gyrus | 17.03 | -2 | -36 | 62 |
| | R Supramarginal Gyrus, posterior division | 2.89 | 52 | -36 | 52 |
| | L Parahippocampal Gyrus, posterior division | 10.97 | -26 | -38 | -16 |
| | R Temporal Fusiform Cortex, posterior division | 10.96 | 24 | -38 | -16 |
| | L Hippocampus | 12.00 | -10 | -42 | 4 |
| | L Inferior Temporal Gyrus, posterior division | 3.92 | -64 | -42 | -20 |
| | R Cingulate Gyrus, posterior division | 13.07 | 10 | -42 | 4 |
| | R Middle Temporal Gyrus, temporooccipital part | 6.67 | 64 | -42 | -12 |
| | L Cingulate Gyrus, posterior division | 12.12 | -12 | -48 | 0 |
| | L Middle Temporal Gyrus, temporooccipital part | 6.75 | -56 | -50 | -10 |
| | R Lingual Gyrus | 13.66 | 12 | -50 | 2 |
| | L Superior Parietal Lobule | 3.73 | -34 | -52 | 56 |
| | L Inferior Temporal Gyrus, temporooccipital part | 6.33 | -46 | -54 | -20 |
| | R Temporal Occipital Fusiform Cortex | 10.23 | 30 | -54 | -6 |
| | R Superior Parietal Lobule | 4.26 | 26 | -54 | 62 |
| | L Temporal Occipital Fusiform Cortex | 8.93 | -28 | -58 | -8 |
| | L Angular Gyrus | 3.76 | -42 | -58 | 26 |
| | R Angular Gyrus | 5.80 | 54 | -58 | 16 |
| | R Inferior Temporal Gyrus, temporooccipital part | 5.22 | 60 | -58 | -12 |
| | L Lingual Gyrus | 13.60 | -12 | -60 | 4 |
| | L Precuneus Cortex | 15.60 | -6 | -68 | 20 |
| | R Precuneus Cortex | 18.89 | 2 | -68 | 22 |
| | R Supracalcarine Cortex | 18.19 | 2 | -68 | 18 |

| | | | | | |
|---|---|---|---|---|---|
| | R Lateral Occipital Cortex, superior division | 10.57 | 52 | -68 | 34 |
| | R Occipital Fusiform Gyrus | 7.45 | 26 | -68 | -10 |
| | R Intracalcarine Cortex | 20.10 | 12 | -70 | 12 |
| | L Occipital Fusiform Gyrus | 5.48 | -26 | -74 | -10 |
| | L Lateral Occipital Cortex, superior division | 10.06 | -46 | -76 | 30 |
| | L Lateral Occipital Cortex, inferior division | 8.31 | -50 | -78 | 4 |
| | R Lateral Occipital Cortex, inferior division | 7.70 | 44 | -78 | 14 |
| | L Intracalcarine Cortex | 20.46 | -14 | -80 | 6 |
| | R Cuneal Cortex | 4.27 | 16 | -80 | 30 |
| | R Occipital Pole | 5.26 | 24 | -88 | 36 |
| | L Occipital Pole | 9.29 | -2 | -96 | 16 |
| 33,072 | | | | | |
| | L Frontal Pole | 6.17 | -6 | 72 | 4 |
| | R Frontal Pole | 11.06 | 2 | 64 | -4 |
| | R Superior Frontal Gyrus | 3.65 | 6 | 54 | 20 |
| | L Frontal Medial Cortex | 6.93 | -4 | 52 | -6 |
| | L Paracingulate Gyrus | 6.15 | -8 | 48 | -4 |
| | R Frontal Medial Cortex | 8.01 | 6 | 40 | -16 |
| | L Cingulate Gyrus, anterior division | 4.86 | -2 | 34 | 0 |
| | R Paracingulate Gyrus | 6.71 | 6 | 34 | -10 |
| | R Cingulate Gyrus, anterior division | 5.52 | 4 | 32 | 8 |
| | R Subcallosal Cortex | 6.46 | 4 | 30 | -18 |
| | L Subcallosal Cortex | 5.41 | -6 | 24 | -16 |
| | R Caudate | 6.32 | 14 | 24 | -2 |
| | L Caudate | 4.78 | -12 | 22 | 2 |
| | R Accumbens | 5.58 | 10 | 20 | -4 |
| 30,608 | | | | | |
| | L Frontal Pole | 7.97 | -50 | 40 | 12 |
| | L Frontal Orbital Cortex | 8.82 | -34 | 34 | -12 |
| | L Inferior Frontal Gyrus, pars triangularis | 7.77 | -44 | 32 | 18 |
| | L Superior Frontal Gyrus | 8.29 | -24 | 24 | 50 |
| | L Middle Frontal Gyrus | 7.79 | -52 | 24 | 30 |
| 24,696 | | | | | |
| | R Frontal Pole | 10.72 | 32 | 36 | -14 |
| | R Superior Frontal Gyrus | 11.16 | 24 | 34 | 50 |
| | R Inferior Frontal Gyrus, pars triangularis | 6.93 | 46 | 34 | 16 |
| | R Middle Frontal Gyrus | 6.59 | 36 | 20 | 50 |
| 2,480 | | | | | |
| | R Brain-Stem | 4.37 | 10 | -22 | -40 |
| | L Brain-Stem | 4.44 | -2 | -26 | -44 |
| 656 | | | | | |
| | L Cingulate Gyrus, anterior division | 5.36 | -4 | 0 | 28 |
| | R Cingulate Gyrus, anterior division | 6.81 | 4 | -4 | 30 |

| | | | | | |
|---|---|---|---|---|---|
| 632 | | | | | |
| | R Occipital Pole | 4.27 | 18 | -100 | -8 |
| 288 | | | | | |
| | L Superior Parietal Lobule | 2.63 | -40 | -46 | 48 |
| 232 | | | | | |
| | L Paracingulate Gyrus | 3.45 | -10 | 44 | 18 |
| 224 | | | | | |
| | R Frontal Pole | 2.81 | 16 | 50 | 34 |
| 176 | | | | | |
| | L Occipital Pole | 3.06 | -24 | -100 | -6 |
| 128 | | | | | |
| | R Caudate | 4.47 | 12 | 18 | 12 |
| 112 | | | | | |
| | L Occipital Pole | 2.93 | -12 | -104 | -8 |
| 80 | | | | | |
| | L Temporal Pole | 2.32 | -20 | 8 | -36 |
| 80 | | | | | |
| | R Temporal Pole | 3.36 | 24 | 10 | -36 |
| 80 | | | | | |
| | R Temporal Pole | 2.82 | 46 | 10 | -24 |
| 80 | | | | | |
| | R Paracingulate Gyrus | 2.94 | 2 | 46 | 12 |
| 64 | | | | | |
| | R Frontal Pole | 3.00 | 24 | 54 | 6 |
| 56 | | | | | |
| | L Frontal Pole | 2.82 | -16 | 64 | -18 |
| 48 | | | | | |
| | R Temporal Pole | 2.80 | 30 | 14 | -44 |
| 48 | | | | | |
| | R Precentral Gyrus | 2.96 | 60 | 10 | 34 |
| 40 | | | | | |
| | L Inferior Temporal Gyrus, posterior division | 2.71 | -56 | -20 | -22 |
| 40 | | | | | |
| | R Accumbens | 2.82 | 12 | 14 | -10 |
| 40 | | | | | |
| | L Brain-Stem | 3.21 | -6 | -36 | -4 |
| 40 | | | | | |
| | L Superior Frontal Gyrus | 2.46 | -4 | 56 | 30 |
| 32 | | | | | |
| | R Inferior Temporal Gyrus, temporooccipital part | 2.66 | 60 | -54 | -22 |
| 32 | | | | | |
| | L Frontal Pole | 2.47 | -14 | 52 | -24 |
| 24 | | | | | |

| | | | | | |
|---|---|---|---|---|---|
| | R Parahippocampal Gyrus, anterior division | 2.40 | 30 | -16 | -34 |
| 24 | | | | | |
| | L Temporal Pole | 2.30 | -48 | 20 | -28 |
| 24 | | | | | |
| | L Frontal Pole | 2.82 | -10 | 44 | -28 |
| 24 | | | | | |
| | R Brain-Stem | 2.63 | 6 | -36 | -10 |
| 24 | | | | | |
| | R Cingulate Gyrus, anterior division | 3.07 | 4 | -12 | 34 |
| 24 | | | | | |
| | R Postcentral Gyrus | 2.79 | 38 | -30 | 60 |
| 16 | | | | | |
| | R Temporal Pole | 2.24 | 26 | 6 | -42 |
| 16 | | | | | |
| | R Frontal Pole | 2.40 | 6 | 60 | -26 |
| 16 | | | | | |
| | R Lateral Occipital Cortex, inferior division | 2.27 | 32 | -86 | -16 |
| 16 | | | | | |
| | L Insular Cortex | 2.58 | -32 | 6 | 8 |
| 16 | | | | | |
| | R Cingulate Gyrus, anterior division | 2.22 | 8 | 42 | 10 |
| 8 | | | | | |
| | L Brain-Stem | 2.21 | -8 | -44 | -38 |
| 8 | | | | | |
| | R Temporal Pole | 2.11 | 28 | 14 | -32 |
| 8 | | | | | |
| | R Temporal Fusiform Cortex, posterior division | 2.13 | 34 | -18 | -30 |
| 8 | | | | | |
| | L Temporal Pole | 2.49 | -38 | 8 | -28 |
| 8 | | | | | |
| | R Inferior Temporal Gyrus, posterior division | 2.28 | 56 | -24 | -26 |
| 8 | | | | | |
| | R Frontal Orbital Cortex | 2.17 | 12 | 28 | -26 |
| 8 | | | | | |
| | L Frontal Orbital Cortex | 2.24 | -18 | 28 | -22 |
| 8 | | | | | |
| | R Frontal Orbital Cortex | 2.12 | 26 | 12 | -16 |
| 8 | | | | | |
| | L Occipital Pole | 2.27 | -24 | -104 | 0 |
| 8 | | | | | |
| | R Frontal Pole | 2.26 | 40 | 62 | 0 |
| 8 | | | | | |

| | | | | | |
|---|---|---|---|---|---|
| | L Frontal Pole | 2.25 | -26 | 54 | 4 |
| 8 | | | | | |
| | L Cingulate Gyrus, anterior division | 2.69 | -4 | 30 | 12 |
| 8 | | | | | |
| | R Paracingulate Gyrus | 2.39 | 14 | 42 | 18 |
| 8 | | | | | |
| | L Cingulate Gyrus, posterior division | 2.14 | -10 | -32 | 38 |
| 8 | | | | | |
| | L Lateral Occipital Cortex, superior division | 2.38 | -14 | -66 | 56 |

**Supplementary Table S11. [$CS_{sustained}$ > $CT_{sustained}$].** Descriptive statistics for clusters and local maxima showing greater activation for Certain-Safety relative to Certain-Threat anticipation for the OSP Sustained regressor (FDR $q$<0.05, whole-brain corrected).

| $mm^3$ | Label | $t$ | $x$ | $y$ | $z$ |
|---|---|---|---|---|---|
| 273,936 | | | | | |
| | R Frontal Pole | 4.95 | 36 | 34 | -14 |
| | R Frontal Orbital Cortex | 3.79 | 22 | 28 | -20 |
| | L Frontal Orbital Cortex | 4.49 | -28 | 12 | -24 |
| | R Temporal Pole | 4.85 | 34 | 12 | -24 |
| | L Temporal Pole | 4.74 | -34 | 10 | -24 |
| | R Planum Polare | 4.06 | 50 | 4 | -8 |
| | L Amygdala | 4.32 | -24 | 0 | -14 |
| | R Superior Temporal Gyrus, anterior division | 4.80 | 58 | 0 | -12 |
| | R Middle Temporal Gyrus, anterior division | 3.68 | 58 | 0 | -26 |
| | L Amygdala | 4.84 | -26 | -2 | -12 |
| | L Amygdala | 4.84 | -26 | -2 | -12 |
| | L Pallidum | 4.34 | -20 | -4 | 0 |
| | R Temporal Fusiform Cortex, anterior division | 3.62 | 38 | -6 | -38 |
| | L Middle Temporal Gyrus, anterior division | 5.83 | -60 | -8 | -10 |
| | R Pallidum | 3.59 | 26 | -8 | 0 |
| | L Postcentral Gyrus | 6.60 | -54 | -10 | 26 |
| | L Juxtapositional Lobule Cortex (formerly Supplementary Motor Cortex) | 5.61 | -10 | -10 | 42 |
| | L Putamen | 5.14 | -28 | -10 | -4 |
| | L Cingulate Gyrus, anterior division | 5.13 | -6 | -10 | 42 |
| | R Cingulate Gyrus, anterior division | 5.49 | 10 | -10 | 42 |
| | R Middle Temporal Gyrus, posterior division | 4.72 | 54 | -10 | -16 |
| | R Postcentral Gyrus | 6.64 | 50 | -12 | 50 |
| | R Putamen | 4.09 | 30 | -12 | 4 |
| | L Central Opercular Cortex | 4.86 | -42 | -14 | 20 |
| | R Juxtapositional Lobule Cortex (formerly Supplementary Motor Cortex) | 4.30 | 8 | -14 | 48 |
| | L Thalamus | 4.90 | -4 | -16 | 12 |
| | L Middle Temporal Gyrus, posterior division | 3.96 | -68 | -16 | -16 |
| | R Central Opercular Cortex | 6.82 | 38 | -16 | 18 |
| | R Heschls Gyrus (includes H1 and H2) | 6.81 | 50 | -16 | 8 |
| | R Inferior Temporal Gyrus, posterior division | 3.22 | 46 | -18 | -32 |
| | R Parahippocampal Gyrus, anterior division | 6.61 | 20 | -20 | -22 |

| | | | | | |
|---|---|---|---|---|---|
| | L Planum Polare | 6.94 | -40 | -22 | 0 |
| | L Parahippocampal Gyrus, anterior division | 3.39 | -18 | -22 | -20 |
| | R Insular Cortex | 8.43 | 36 | -22 | 14 |
| | R Supramarginal Gyrus, anterior division | 3.42 | 64 | -22 | 20 |
| | L Insular Cortex | 7.94 | -34 | -24 | 16 |
| | L Hippocampus | 4.95 | -26 | -24 | -10 |
| | L Precentral Gyrus | 8.24 | -26 | -26 | 66 |
| | L Heschls Gyrus (includes H1 and H2) | 8.18 | -34 | -26 | 12 |
| | R Precentral Gyrus | 7.54 | 6 | -26 | 56 |
| | R Hippocampus | 6.31 | 24 | -26 | -8 |
| | R Parietal Operculum Cortex | 2.67 | 54 | -28 | 24 |
| | L Planum Temporale | 7.73 | -34 | -30 | 16 |
| | R Thalamus | 5.57 | 18 | -30 | -4 |
| | R Planum Temporale | 6.64 | 40 | -32 | 16 |
| | L Parietal Operculum Cortex | 5.00 | -44 | -34 | 22 |
| | L Brain-Stem | 3.96 | -8 | -34 | -8 |
| | L Temporal Fusiform Cortex, posterior division | 3.27 | -34 | -34 | -28 |
| | R Parahippocampal Gyrus, posterior division | 8.64 | 22 | -34 | -18 |
| | R Temporal Fusiform Cortex, posterior division | 6.67 | 30 | -34 | -18 |
| | R Brain-Stem | 3.18 | 6 | -34 | -8 |
| | L Parahippocampal Gyrus, posterior division | 8.86 | -30 | -38 | -12 |
| | R Lingual Gyrus | 10.41 | 18 | -42 | -10 |
| | R Cingulate Gyrus, posterior division | 9.33 | 10 | -42 | 4 |
| | L Lingual Gyrus | 11.68 | -12 | -52 | -2 |
| | L Cingulate Gyrus, posterior division | 8.05 | -2 | -52 | 22 |
| | L Angular Gyrus | 3.32 | -56 | -52 | 18 |
| | R Temporal Occipital Fusiform Cortex | 9.15 | 30 | -52 | -6 |
| | L Superior Parietal Lobule | 3.20 | -14 | -54 | 68 |
| | R Angular Gyrus | 4.88 | 48 | -54 | 20 |
| | L Precuneus Cortex | 8.42 | -18 | -56 | 4 |
| | R Middle Temporal Gyrus, temporooccipital part | 2.99 | 52 | -58 | 10 |
| | L Middle Temporal Gyrus, temporooccipital part | 4.05 | -50 | -60 | 0 |
| | L Inferior Temporal Gyrus, temporooccipital part | 3.75 | -54 | -60 | -14 |
| | L Temporal Occipital Fusiform Cortex | 5.36 | -34 | -62 | -14 |
| | R Occipital Fusiform Gyrus | 9.20 | 28 | -66 | -12 |
| | R Cuneal Cortex | 14.49 | 2 | -70 | 20 |
| | L Occipital Fusiform Gyrus | 7.22 | -28 | -74 | -8 |

| | | | | | |
|---|---|---|---|---|---|
| | L Lateral Occipital Cortex, inferior division | 4.43 | -52 | -74 | 8 |
| | R Intracalcarine Cortex | 20.22 | 12 | -74 | 12 |
| | L Intracalcarine Cortex | 20.48 | -10 | -76 | 10 |
| | R Lateral Occipital Cortex, superior division | 6.39 | 46 | -76 | 16 |
| | R Lateral Occipital Cortex, inferior division | 6.28 | 42 | -76 | 10 |
| | L Lateral Occipital Cortex, superior division | 6.51 | -42 | -78 | 20 |
| | R Precuneus Cortex | 9.62 | 2 | -78 | 38 |
| | L Cuneal Cortex | 7.80 | -2 | -86 | 34 |
| | L Occipital Pole | 10.28 | -2 | -96 | 10 |
| | R Occipital Pole | 8.67 | 8 | -96 | 18 |
| 23,496 | | | | | |
| | L Frontal Pole | 5.71 | -2 | 66 | 0 |
| | R Frontal Pole | 7.66 | 0 | 64 | -10 |
| | R Frontal Medial Cortex | 6.78 | 2 | 50 | -12 |
| | L Paracingulate Gyrus | 4.68 | -6 | 46 | -6 |
| | R Paracingulate Gyrus | 5.95 | 2 | 40 | -10 |
| | L Frontal Medial Cortex | 5.47 | -8 | 36 | -12 |
| | R Cingulate Gyrus, anterior division | 5.33 | 0 | 36 | 4 |
| | L Cingulate Gyrus, anterior division | 4.64 | -2 | 32 | -6 |
| | R Subcallosal Cortex | 4.86 | 2 | 30 | -14 |
| | L Caudate | 4.18 | -14 | 24 | -2 |
| | L Subcallosal Cortex | 4.65 | -4 | 20 | -10 |
| | R Accumbens | 3.77 | 8 | 18 | -6 |
| | L Accumbens | 4.43 | -6 | 16 | -4 |
| 4,872 | | | | | |
| | R Superior Frontal Gyrus | 5.39 | 22 | 28 | 42 |
| | R Middle Frontal Gyrus | 6.87 | 26 | 18 | 44 |
| 4,808 | | | | | |
| | L Frontal Pole | 2.56 | -20 | 46 | 44 |
| | L Superior Frontal Gyrus | 6.24 | -22 | 24 | 40 |
| | L Middle Frontal Gyrus | 3.36 | -32 | 24 | 50 |
| 1,080 | | | | | |
| | L Frontal Orbital Cortex | 5.18 | -24 | 30 | -20 |
| 928 | | | | | |
| | L Frontal Pole | 2.55 | -48 | 38 | 18 |
| | L Inferior Frontal Gyrus, pars triangularis | 3.98 | -48 | 30 | 16 |
| 712 | | | | | |
| | R Frontal Pole | 2.80 | 52 | 36 | 12 |
| | R Inferior Frontal Gyrus, pars triangularis | 2.73 | 48 | 34 | 14 |
| | R Middle Frontal Gyrus | 3.74 | 46 | 30 | 20 |
| 632 | | | | | |
| | L Brain-Stem | 3.71 | -12 | -28 | -38 |
| 560 | | | | | |

| | | | | | |
|---|---|---|---|---|---|
| | R Amygdala | 4.40 | 22 | 2 | -16 |
| | R Amygdala | 5.40 | 28 | 0 | -12 |
| | R Amygdala | 3.20 | 18 | 0 | -24 |
| 248 | | | | | |
| | R Subcallosal Cortex | 3.66 | 12 | 18 | -14 |
| | R Frontal Orbital Cortex | 3.71 | 16 | 12 | -18 |
| 240 | | | | | |
| | R Superior Parietal Lobule | 3.13 | 12 | -54 | 66 |
| 224 | | | | | |
| | L Supramarginal Gyrus, anterior division | 3.11 | -62 | -28 | 32 |
| 208 | | | | | |
| | R Brain-Stem | 4.40 | 0 | -18 | -30 |
| | L Brain-Stem | 3.25 | -2 | -24 | -40 |
| 168 | | | | | |
| | R Middle Frontal Gyrus | 3.35 | 38 | 10 | 56 |
| 160 | | | | | |
| | R Frontal Pole | 3.13 | 14 | 72 | 14 |
| 144 | | | | | |
| | R Cingulate Gyrus, anterior division | 4.11 | 4 | 10 | 36 |
| 136 | | | | | |
| | R Brain-Stem | 3.14 | 4 | -30 | -40 |
| 128 | | | | | |
| | L Cingulate Gyrus, anterior division | 3.26 | -4 | 0 | 28 |
| | R Cingulate Gyrus, anterior division | 3.71 | 0 | -2 | 30 |
| 120 | | | | | |
| | L Brain-Stem | 3.34 | -8 | -28 | -18 |
| 120 | | | | | |
| | L Middle Frontal Gyrus | 2.96 | -52 | 16 | 34 |
| 112 | | | | | |
| | R Temporal Pole | 2.79 | 44 | 4 | -44 |
| | R Inferior Temporal Gyrus, anterior division | 3.09 | 42 | 2 | -42 |
| 104 | | | | | |
| | R Temporal Fusiform Cortex, anterior division | 2.79 | 30 | -6 | -46 |
| 104 | | | | | |
| | R Frontal Orbital Cortex | 3.59 | 14 | 30 | -18 |
| | R Subcallosal Cortex | 3.24 | 10 | 28 | -20 |
| 96 | | | | | |
| | L Temporal Pole | 4.02 | -28 | 12 | -48 |
| 96 | | | | | |
| | L Temporal Fusiform Cortex, posterior division | 2.56 | -46 | -40 | -24 |
| | L Inferior Temporal Gyrus, posterior division | 2.56 | -48 | -42 | -22 |

| | | | | | |
|---|---|---|---|---|---|
| | L Inferior Temporal Gyrus, temporooccipital part | 3.01 | -48 | -46 | -18 |
| 96 | | | | | |
| | L Inferior Temporal Gyrus, temporooccipital part | 3.02 | -58 | -46 | -18 |
| | L Middle Temporal Gyrus, temporooccipital part | 2.52 | -62 | -46 | -12 |
| 96 | | | | | |
| | R Middle Temporal Gyrus, temporooccipital part | 3.19 | 66 | -54 | -2 |
| 96 | | | | | |
| | R Thalamus | 3.39 | 10 | -20 | 18 |
| 88 | | | | | |
| | R Temporal Occipital Fusiform Cortex | 2.91 | 42 | -40 | -26 |
| 88 | | | | | |
| | R Brain-Stem | 3.84 | 10 | -28 | -26 |
| 80 | | | | | |
| | L Thalamus | 3.52 | -10 | -22 | 18 |
| 72 | | | | | |
| | R Brain-Stem | 3.64 | 8 | -44 | -36 |
| 72 | | | | | |
| | L Inferior Temporal Gyrus, posterior division | 3.39 | -64 | -38 | -24 |
| 72 | | | | | |
| | R Middle Temporal Gyrus, temporooccipital part | 3.00 | 54 | -52 | 2 |
| 64 | | | | | |
| | R Inferior Temporal Gyrus, posterior division | 3.09 | 52 | -16 | -38 |
| 64 | | | | | |
| | R Brain-Stem | 2.95 | 14 | -24 | -36 |
| 64 | | | | | |
| | R Brain-Stem | 2.76 | 2 | -16 | -22 |
| 64 | | | | | |
| | R Amygdala | 3.67 | 18 | -6 | -16 |
| 64 | | | | | |
| | L Middle Frontal Gyrus | 3.03 | -52 | 16 | 42 |
| 56 | | | | | |
| | L Parahippocampal Gyrus, anterior division | 3.27 | -22 | -2 | -38 |
| 56 | | | | | |
| | R Brain-Stem | 3.57 | 10 | -14 | -34 |
| 56 | | | | | |
| | L Temporal Pole | 3.22 | -34 | 8 | -36 |
| 56 | | | | | |

| | | | | | |
|---|---|---|---|---|---|
| | R Middle Temporal Gyrus, anterior division | 2.52 | 54 | -2 | -28 |
| 56 | | | | | |
| | R Frontal Pole | 3.54 | 12 | 36 | -20 |
| 56 | | | | | |
| | R Middle Frontal Gyrus | 3.07 | 52 | 24 | 36 |
| 48 | | | | | |
| | R Temporal Fusiform Cortex, anterior division | 2.85 | 26 | 0 | -44 |
| 48 | | | | | |
| | L Temporal Pole | 2.71 | -30 | 14 | -38 |
| 48 | | | | | |
| | L Brain-Stem | 3.35 | -4 | -42 | -36 |
| 48 | | | | | |
| | R Temporal Pole | 3.17 | 46 | 14 | -26 |
| 48 | | | | | |
| | R Middle Frontal Gyrus | 2.70 | 42 | 24 | 36 |
| 32 | | | | | |
| | R Brain-Stem | 3.14 | 10 | -24 | -40 |
| 32 | | | | | |
| | L Inferior Temporal Gyrus, posterior division | 2.51 | -52 | -18 | -36 |
| 32 | | | | | |
| | R Temporal Pole | 3.21 | 52 | 20 | -26 |
| 32 | | | | | |
| | R Pallidum | 2.98 | 16 | 2 | 0 |
| 32 | | | | | |
| | R Lateral Occipital Cortex, superior division | 2.44 | 34 | -72 | 50 |
| 24 | | | | | |
| | R Inferior Temporal Gyrus, posterior division | 2.79 | 60 | -14 | -38 |
| 24 | | | | | |
| | R Inferior Temporal Gyrus, posterior division | 2.85 | 58 | -30 | -24 |
| 24 | | | | | |
| | L Amygdala | 2.85 | -14 | -6 | -16 |
| 24 | | | | | |
| | L Middle Temporal Gyrus, posterior division | 2.63 | -62 | -30 | -12 |
| 24 | | | | | |
| | L Temporal Pole | 2.68 | -48 | 10 | -10 |
| 24 | | | | | |
| | L Hippocampus | 2.49 | -28 | -36 | -6 |
| 24 | | | | | |

| | | | | | |
|---|---|---|---|---|---|
| | R Pallidum | 2.69 | 22 | -4 | -6 |
| 24 | | | | | |
| | R Caudate | 2.65 | 14 | 16 | 14 |
| 24 | | | | | |
| | R Middle Frontal Gyrus | 2.69 | 42 | 18 | 40 |
| 24 | | | | | |
| | R Superior Frontal Gyrus | 3.09 | 22 | -2 | 54 |
| 16 | | | | | |
| | L Brain-Stem | 3.08 | -10 | -38 | -40 |
| 16 | | | | | |
| | L Temporal Fusiform Cortex, anterior division | 2.39 | -36 | -10 | -36 |
| 16 | | | | | |
| | R Brain-Stem | 2.54 | 12 | -30 | -32 |
| 16 | | | | | |
| | L Inferior Temporal Gyrus, posterior division | 2.54 | -50 | -20 | -30 |
| 16 | | | | | |
| | R Brain-Stem | 2.55 | 12 | -38 | -30 |
| 16 | | | | | |
| | R Brain-Stem | 2.96 | 14 | -30 | -28 |
| 16 | | | | | |
| | R Brain-Stem | 2.61 | 12 | -16 | -28 |
| 16 | | | | | |
| | L Inferior Temporal Gyrus, posterior division | 2.61 | -56 | -44 | -24 |
| 16 | | | | | |
| | L Brain-Stem | 2.67 | -14 | -32 | -26 |
| 16 | | | | | |
| | R Inferior Temporal Gyrus, temporooccipital part | 2.39 | 64 | -50 | -18 |
| 16 | | | | | |
| | L Planum Polare | 2.87 | -40 | -4 | -18 |
| 16 | | | | | |
| | L Frontal Orbital Cortex | 2.70 | -12 | 14 | -18 |
| 16 | | | | | |
| | L Frontal Orbital Cortex | 2.61 | -44 | 32 | -18 |
| 16 | | | | | |
| | L Inferior Temporal Gyrus, temporooccipital part | 2.75 | -52 | -50 | -12 |
| 16 | | | | | |
| | L Inferior Temporal Gyrus, temporooccipital part | 2.50 | -42 | -58 | -6 |
| 16 | | | | | |
| | R Frontal Pole | 2.73 | 24 | 54 | 8 |

| | | | | | |
|---|---|---|---|---|---|
| 16 | | | | | |
| | L Frontal Pole | 3.65 | -14 | 62 | 16 |
| 16 | | | | | |
| | L Thalamus | 3.09 | -6 | -14 | 18 |
| 16 | | | | | |
| | L Cingulate Gyrus, anterior division | 2.78 | -2 | 8 | 26 |
| 16 | | | | | |
| | L Frontal Pole | 3.29 | -14 | 52 | 30 |
| 16 | | | | | |
| | R Postcentral Gyrus | 2.37 | 46 | -18 | 64 |
| 8 | | | | | |
| | L Brain-Stem | 2.30 | -4 | -40 | -48 |
| 8 | | | | | |
| | R Inferior Temporal Gyrus, anterior division | 2.31 | 48 | 0 | -46 |
| 8 | | | | | |
| | L Temporal Fusiform Cortex, anterior division | 2.30 | -30 | -6 | -44 |
| 8 | | | | | |
| | L Temporal Pole | 2.38 | -38 | 14 | -40 |
| 8 | | | | | |
| | R Middle Temporal Gyrus, anterior division | 2.62 | 54 | 2 | -36 |
| 8 | | | | | |
| | R Brain-Stem | 2.36 | 8 | -26 | -34 |
| 8 | | | | | |
| | L Brain-Stem | 2.35 | -2 | -26 | -28 |
| 8 | | | | | |
| | R Middle Temporal Gyrus, posterior division | 2.34 | 58 | -12 | -28 |
| 8 | | | | | |
| | L Inferior Temporal Gyrus, posterior division | 2.48 | -56 | -38 | -24 |
| 8 | | | | | |
| | R Parahippocampal Gyrus, anterior division | 2.28 | 16 | -8 | -24 |
| 8 | | | | | |
| | L Brain-Stem | 2.36 | -2 | -22 | -22 |
| 8 | | | | | |
| | R Hippocampus | 2.40 | 12 | -10 | -22 |
| 8 | | | | | |
| | R Temporal Fusiform Cortex, posterior division | 2.29 | 42 | -24 | -20 |
| 8 | | | | | |
| | L Frontal Orbital Cortex | 2.35 | -22 | 16 | -20 |

| | | | | | |
|---|---|---|---|---|---|
| 8 | | | | | |
| | R Frontal Pole | 2.39 | 16 | 40 | -20 |
| 8 | | | | | |
| | R Frontal Pole | 2.32 | 48 | 36 | -18 |
| 8 | | | | | |
| | R Inferior Temporal Gyrus, temporooccipital part | 2.48 | 46 | -42 | -16 |
| 8 | | | | | |
| | L Frontal Orbital Cortex | 2.52 | -14 | 20 | -16 |
| 8 | | | | | |
| | R Hippocampus | 2.43 | 34 | -24 | -14 |
| 8 | | | | | |
| | L Amygdala | 2.32 | -18 | -12 | -14 |
| 8 | | | | | |
| | R Amygdala | 2.33 | 22 | -14 | -12 |
| 8 | | | | | |
| | R Middle Temporal Gyrus, temporooccipital part | 2.37 | 58 | -44 | -8 |
| 8 | | | | | |
| | L Middle Temporal Gyrus, posterior division | 2.45 | -54 | -26 | -8 |
| 8 | | | | | |
| | L Frontal Pole | 2.32 | -12 | 64 | -6 |
| 8 | | | | | |
| | R Putamen | 2.30 | 32 | -2 | -4 |
| 8 | | | | | |
| | L Superior Temporal Gyrus, posterior division | 2.40 | -60 | -22 | -2 |
| 8 | | | | | |
| | R Insular Cortex | 2.39 | 34 | 20 | 0 |
| 8 | | | | | |
| | R Caudate | 2.31 | 10 | 20 | 6 |
| 8 | | | | | |
| | L Caudate | 2.85 | -12 | 20 | 8 |
| 8 | | | | | |
| | R Middle Temporal Gyrus, temporooccipital part | 2.29 | 68 | -46 | 10 |
| 8 | | | | | |
| | L Caudate | 2.83 | -8 | 12 | 12 |
| 8 | | | | | |
| | L Cingulate Gyrus, anterior division | 2.59 | -4 | 26 | 16 |
| 8 | | | | | |
| | R Frontal Pole | 2.30 | 16 | 62 | 16 |
| 8 | | | | | |
| | R Precentral Gyrus | 2.31 | 60 | 8 | 22 |

| | | | | | |
|---|---|---|---|---|---|
| 8 | | | | | |
| | L Cingulate Gyrus, posterior division | 2.86 | -4 | -24 | 26 |
| 8 | | | | | |
| | L Inferior Frontal Gyrus, pars opercularis | 2.37 | -42 | 14 | 26 |
| 8 | | | | | |
| | L Precentral Gyrus | 2.59 | -40 | -2 | 32 |
| 8 | | | | | |
| | L Cingulate Gyrus, anterior division | 2.81 | -8 | 12 | 32 |
| 8 | | | | | |
| | R Superior Frontal Gyrus | 2.45 | 0 | 48 | 36 |
| 8 | | | | | |
| | R Precuneus Cortex | 2.84 | 12 | -64 | 38 |
| 8 | | | | | |
| | R Frontal Pole | 2.48 | 20 | 36 | 38 |
| 8 | | | | | |
| | R Cingulate Gyrus, anterior division | 2.33 | 4 | 2 | 40 |
| 8 | | | | | |
| | L Frontal Pole | 2.40 | -14 | 50 | 40 |
| 8 | | | | | |
| | R Precuneus Cortex | 2.44 | 14 | -42 | 42 |
| 8 | | | | | |
| | L Frontal Pole | 2.28 | -12 | 46 | 42 |
| 8 | | | | | |
| | L Postcentral Gyrus | 2.56 | -42 | -28 | 52 |
| 8 | | | | | |
| | L Lateral Occipital Cortex, superior division | 2.42 | -16 | -76 | 54 |

**Supplementary Table S12. [$CT_{sustained}$ > $UT_{sustained}$].** Descriptive statistics for clusters and local maxima showing greater activation for Certain-Threat relative to Uncertain-Threat anticipation for the OSP Sustained regressor (FDR $q$<0.05, whole-brain corrected).

| ***mm³*** | **Label** | ***t*** | ***x*** | ***y*** | ***z*** |
|---|---|---|---|---|---|
| 384,256 | | | | | |
| | L Frontal Pole | 3.75 | -50 | 40 | 10 |
| | L Inferior Frontal Gyrus, pars triangularis | 4.77 | -44 | 32 | 18 |
| | L Middle Frontal Gyrus | 5.83 | -40 | 24 | 24 |
| | L Temporal Pole | 4.36 | -58 | 8 | -22 |
| | R Temporal Pole | 4.53 | 52 | 8 | -10 |
| | R Amygdala | 3.94 | 28 | 4 | -26 |
| | L Amygdala | 4.85 | -32 | 2 | -18 |
| | R Superior Temporal Gyrus, anterior division | 6.25 | 60 | 2 | -10 |
| | L Amygdala | 4.48 | -24 | 0 | -20 |
| | L Temporal Fusiform Cortex, anterior division | 3.86 | -30 | 0 | -36 |
| | R Middle Temporal Gyrus, anterior division | 8.28 | 58 | 0 | -20 |
| | R Amygdala | 5.10 | 26 | -2 | -18 |
| | L Amygdala | 4.22 | -22 | -2 | -18 |
| | R Planum Polare | 4.76 | 50 | -2 | -6 |
| | R Amygdala | 4.65 | 18 | -4 | -18 |
| | R Amygdala | 4.62 | 14 | -4 | -16 |
| | L Middle Temporal Gyrus, anterior division | 5.86 | -58 | -4 | -14 |
| | L Superior Temporal Gyrus, anterior division | 3.75 | -52 | -4 | -10 |
| | R Precentral Gyrus | 12.47 | 52 | -6 | 28 |
| | L Amygdala | 6.50 | -16 | -8 | -20 |
| | L Amygdala | 4.18 | -22 | -8 | -22 |
| | R Postcentral Gyrus | 13.05 | 66 | -8 | 32 |
| | R Insular Cortex | 11.89 | 38 | -8 | 12 |
| | L Insular Cortex | 13.21 | -36 | -10 | 14 |
| | R Central Opercular Cortex | 7.75 | 44 | -10 | 16 |
| | R Planum Temporale | 6.46 | 66 | -10 | 6 |
| | L Central Opercular Cortex | 13.50 | -38 | -12 | 16 |
| | L Precentral Gyrus | 12.10 | -48 | -12 | 36 |
| | L Parahippocampal Gyrus, anterior division | 5.92 | -18 | -12 | -26 |
| | L Planum Polare | 4.41 | -48 | -12 | -2 |
| | R Middle Temporal Gyrus, posterior division | 5.93 | 60 | -12 | -8 |
| | L Putamen | 4.82 | -32 | -14 | -8 |
| | L Hippocampus | 8.19 | -22 | -20 | -18 |

| | | | | | |
|---|---|---|---|---|---|
| | L Middle Temporal Gyrus, posterior division | 4.51 | -62 | -20 | -12 |
| | R Hippocampus | 9.41 | 22 | -20 | -18 |
| | R Superior Temporal Gyrus, posterior division | 6.71 | 66 | -20 | 4 |
| | R Parahippocampal Gyrus, anterior division | 4.28 | 28 | -20 | -28 |
| | R Parietal Operculum Cortex | 6.59 | 36 | -22 | 18 |
| | L Thalamus | 7.30 | -16 | -24 | -2 |
| | R Supramarginal Gyrus, anterior division | 6.81 | 60 | -24 | 44 |
| | R Temporal Fusiform Cortex, posterior division | 6.62 | 34 | -24 | -22 |
| | R Heschls Gyrus (includes H1 and H2) | 4.40 | 48 | -24 | 8 |
| | L Heschls Gyrus (includes H1 and H2) | 5.86 | -48 | -26 | 6 |
| | R Inferior Temporal Gyrus, posterior division | 3.28 | 60 | -26 | -22 |
| | R Parahippocampal Gyrus, posterior division | 7.90 | 28 | -28 | -22 |
| | L Parietal Operculum Cortex | 9.43 | -36 | -30 | 18 |
| | L Superior Temporal Gyrus, posterior division | 6.67 | -68 | -30 | 6 |
| | L Inferior Temporal Gyrus, posterior division | 5.43 | -46 | -30 | -28 |
| | L Supramarginal Gyrus, anterior division | 4.53 | -56 | -30 | 42 |
| | L Temporal Fusiform Cortex, posterior division | 9.15 | -30 | -34 | -18 |
| | L Brain-Stem | 2.56 | -20 | -34 | -32 |
| | R Thalamus | 6.75 | 14 | -34 | 4 |
| | L Postcentral Gyrus | 13.03 | -2 | -36 | 62 |
| | L Planum Temporale | 8.69 | -62 | -36 | 16 |
| | L Parahippocampal Gyrus, posterior division | 10.23 | -24 | -38 | -16 |
| | R Lingual Gyrus | 12.21 | 32 | -40 | -10 |
| | R Supramarginal Gyrus, posterior division | 5.20 | 38 | -40 | 44 |
| | L Supramarginal Gyrus, posterior division | 4.48 | -44 | -42 | 46 |
| | L Cingulate Gyrus, posterior division | 8.70 | -8 | -44 | 4 |
| | R Superior Parietal Lobule | 4.42 | 38 | -44 | 54 |
| | L Lingual Gyrus | 11.44 | -28 | -46 | -8 |
| | R Cingulate Gyrus, posterior division | 9.97 | 2 | -48 | 18 |
| | L Superior Parietal Lobule | 8.23 | -34 | -52 | 58 |
| | L Inferior Temporal Gyrus, temporooccipital part | 4.48 | -48 | -54 | -20 |
| | L Middle Temporal Gyrus, temporooccipital part | 7.28 | -60 | -56 | -10 |
| | L Temporal Occipital Fusiform Cortex | 3.43 | -42 | -56 | -14 |
| | L Angular Gyrus | 5.23 | -44 | -58 | 22 |

| | | | | | |
|---|---|---|---|---|---|
| | R Angular Gyrus | 2.55 | 46 | -58 | 44 |
| | R Intracalcarine Cortex | 10.36 | 24 | -60 | 6 |
| | L Intracalcarine Cortex | 10.72 | -14 | -64 | 10 |
| | R Precuneus Cortex | 15.56 | 10 | -64 | 18 |
| | L Lateral Occipital Cortex, inferior division | 8.51 | -56 | -66 | -8 |
| | L Precuneus Cortex | 12.26 | -6 | -68 | 20 |
| | R Cuneal Cortex | 13.46 | 18 | -68 | 22 |
| | R Supracalcarine Cortex | 8.90 | 0 | -78 | 14 |
| | L Lateral Occipital Cortex, superior division | 11.21 | -24 | -82 | 32 |
| | L Cuneal Cortex | 8.78 | -4 | -82 | 30 |
| | R Lateral Occipital Cortex, superior division | 12.64 | 38 | -82 | 32 |
| | L Supracalcarine Cortex | 7.68 | -2 | -84 | 10 |
| | R Occipital Pole | 8.24 | 14 | -88 | 34 |
| | L Occipital Pole | 7.18 | -2 | -90 | 24 |
| 66,152 | | | | | |
| | R Frontal Pole | 8.59 | 2 | 58 | -8 |
| | R Paracingulate Gyrus | 4.67 | 6 | 54 | 18 |
| | R Frontal Medial Cortex | 8.08 | 6 | 50 | -10 |
| | L Frontal Medial Cortex | 5.84 | -2 | 48 | -16 |
| | L Paracingulate Gyrus | 5.69 | -8 | 46 | -8 |
| | L Frontal Pole | 5.91 | -18 | 40 | 50 |
| | L Cingulate Gyrus, anterior division | 3.69 | -2 | 36 | -6 |
| | R Superior Frontal Gyrus | 10.97 | 26 | 32 | 50 |
| | R Cingulate Gyrus, anterior division | 3.47 | 6 | 32 | 10 |
| | L Superior Frontal Gyrus | 8.16 | -24 | 24 | 46 |
| | L Subcallosal Cortex | 5.83 | -4 | 24 | -18 |
| | R Caudate | 3.88 | 14 | 24 | -2 |
| | R Middle Frontal Gyrus | 8.25 | 28 | 20 | 54 |
| | R Accumbens | 2.57 | 12 | 16 | -6 |
| | R Subcallosal Cortex | 4.85 | 2 | 14 | -6 |
| | L Middle Frontal Gyrus | 2.44 | -32 | 8 | 64 |
| 10,376 | | | | | |
| | L Lingual Gyrus | 2.75 | -8 | -86 | -16 |
| 2,912 | | | | | |
| | R Inferior Temporal Gyrus, posterior division | 3.09 | 58 | -34 | -16 |
| | R Middle Temporal Gyrus, temporooccipital part | 4.67 | 58 | -50 | -10 |
| | R Inferior Temporal Gyrus, temporooccipital part | 5.04 | 60 | -58 | -12 |
| | R Lateral Occipital Cortex, inferior division | 3.45 | 58 | -60 | -4 |
| 2,896 | | | | | |
| | L Frontal Pole | 4.05 | -48 | 46 | -12 |

| | | | | | |
|---|---|---|---|---|---|
| | L Frontal Orbital Cortex | 6.69 | -34 | 36 | -12 |
| 456 | | | | | |
| | L Juxtapositional Lobule Cortex (formerly Supplementary Motor Cortex) | 6.39 | -4 | 0 | 64 |
| 376 | | | | | |
| | R Temporal Fusiform Cortex, anterior division | 3.19 | 32 | -4 | -40 |
| | R Parahippocampal Gyrus, anterior division | 2.88 | 26 | -6 | -36 |
| 360 | | | | | |
| | R Temporal Pole | 3.67 | 24 | 16 | -44 |
| 296 | | | | | |
| | L Brain-Stem | 2.47 | -2 | -36 | -6 |
| | R Brain-Stem | 3.91 | 6 | -38 | -12 |
| 208 | | | | | |
| | L Cingulate Gyrus, anterior division | 3.62 | -4 | -2 | 30 |
| | R Cingulate Gyrus, anterior division | 4.24 | 4 | -4 | 32 |
| 184 | | | | | |
| | R Temporal Fusiform Cortex, posterior division | 3.72 | 40 | -22 | -32 |
| 168 | | | | | |
| | L Brain-Stem | 3.21 | -6 | -26 | -44 |
| 160 | | | | | |
| | L Caudate | 3.24 | -18 | 18 | 12 |
| 96 | | | | | |
| | L Temporal Pole | 2.35 | -32 | 6 | -28 |
| 96 | | | | | |
| | L Middle Frontal Gyrus | 2.93 | -40 | 14 | 46 |
| 80 | | | | | |
| | L Middle Temporal Gyrus, temporooccipital part | 2.70 | -48 | -52 | 6 |
| 72 | | | | | |
| | R Brain-Stem | 3.06 | 0 | -18 | -40 |
| 64 | | | | | |
| | L Temporal Pole | 2.72 | -44 | 22 | -32 |
| 64 | | | | | |
| | L Inferior Temporal Gyrus, posterior division | 2.48 | -60 | -24 | -26 |
| 56 | | | | | |
| | L Brain-Stem | 3.20 | -12 | -24 | -32 |
| 48 | | | | | |
| | R Inferior Temporal Gyrus, temporooccipital part | 2.38 | 58 | -56 | -22 |
| 48 | | | | | |
| | L Frontal Orbital Cortex | 2.56 | -34 | 30 | -22 |

| | | | | | |
|---|---|---|---|---|---|
| 48 | | | | | |
| | L Frontal Pole | 2.44 | -10 | 54 | -24 |
| 40 | | | | | |
| | L Temporal Pole | 2.85 | -28 | 18 | -40 |
| 40 | | | | | |
| | R Brain-Stem | 2.82 | 0 | -20 | -24 |
| 40 | | | | | |
| | L Frontal Pole | 2.97 | -12 | 64 | -18 |
| 32 | | | | | |
| | R Temporal Pole | 2.45 | 44 | 14 | -20 |
| 32 | | | | | |
| | R Paracingulate Gyrus | 2.81 | 12 | 44 | 18 |
| 32 | | | | | |
| | R Caudate | 2.76 | 16 | 2 | 22 |
| 24 | | | | | |
| | R Temporal Fusiform Cortex, posterior division | 2.24 | 36 | -16 | -28 |
| 24 | | | | | |
| | L Temporal Pole | 2.51 | -48 | 12 | -28 |
| 24 | | | | | |
| | L Caudate | 2.27 | -12 | 20 | -4 |
| 24 | | | | | |
| | L Frontal Pole | 2.59 | -36 | 48 | 4 |
| 24 | | | | | |
| | R Angular Gyrus | 2.32 | 42 | -52 | 48 |
| 16 | | | | | |
| | L Temporal Fusiform Cortex, posterior division | 2.21 | -36 | -14 | -38 |
| 16 | | | | | |
| | R Temporal Pole | 2.20 | 24 | 4 | -40 |
| 16 | | | | | |
| | L Brain-Stem | 2.84 | -6 | -14 | -38 |
| 16 | | | | | |
| | L Brain-Stem | 2.45 | -10 | -32 | -36 |
| 16 | | | | | |
| | R Brain-Stem | 2.63 | 4 | -18 | -32 |
| 16 | | | | | |
| | R Temporal Pole | 2.26 | 24 | 10 | -32 |
| 16 | | | | | |
| | R Brain-Stem | 3.11 | 6 | -42 | -28 |
| 16 | | | | | |
| | L Brain-Stem | 2.20 | -8 | -28 | -28 |
| 16 | | | | | |
| | R Brain-Stem | 2.44 | 2 | -24 | -26 |
| 16 | | | | | |

| | | | | | |
|---|---|---|---|---|---|
| | R Parahippocampal Gyrus, anterior division | 2.34 | 20 | 2 | -28 |
| 16 | | | | | |
| | L Temporal Pole | 2.72 | -38 | 10 | -28 |
| 16 | | | | | |
| | R Frontal Orbital Cortex | 2.95 | 10 | 18 | -26 |
| 16 | | | | | |
| | L Temporal Occipital Fusiform Cortex | 2.37 | -42 | -48 | -24 |
| 16 | | | | | |
| | L Frontal Medial Cortex | 2.48 | -6 | 48 | -24 |
| 16 | | | | | |
| | R Frontal Orbital Cortex | 2.39 | 26 | 12 | -18 |
| 16 | | | | | |
| | R Frontal Pole | 2.79 | 38 | 60 | -10 |
| 16 | | | | | |
| | L Insular Cortex | 3.01 | -32 | 6 | 8 |
| 16 | | | | | |
| | L Frontal Pole | 3.17 | -18 | 48 | 30 |
| 8 | | | | | |
| | R Brain-Stem | 2.23 | 10 | -20 | -44 |
| 8 | | | | | |
| | R Brain-Stem | 2.31 | 6 | -44 | -42 |
| 8 | | | | | |
| | R Brain-Stem | 2.23 | 8 | -26 | -42 |
| 8 | | | | | |
| | L Temporal Fusiform Cortex, posterior division | 2.37 | -28 | -12 | -42 |
| 8 | | | | | |
| | L Brain-Stem | 2.49 | -10 | -18 | -40 |
| 8 | | | | | |
| | R Brain-Stem | 2.16 | 12 | -26 | -38 |
| 8 | | | | | |
| | L Temporal Pole | 2.24 | -20 | 8 | -36 |
| 8 | | | | | |
| | R Temporal Pole | 2.24 | 24 | 20 | -36 |
| 8 | | | | | |
| | L Temporal Pole | 2.31 | -42 | 14 | -34 |
| 8 | | | | | |
| | L Temporal Fusiform Cortex, posterior division | 2.35 | -40 | -30 | -30 |
| 8 | | | | | |
| | R Inferior Temporal Gyrus, posterior division | 2.16 | 54 | -24 | -26 |
| 8 | | | | | |

| | | | | | |
|---|---|---|---|---|---|
| | R Inferior Temporal Gyrus, posterior division | 2.18 | 52 | -38 | -18 |
| 8 | | | | | |
| | L Middle Temporal Gyrus, posterior division | 2.21 | -64 | -30 | -18 |
| 8 | | | | | |
| | L Middle Temporal Gyrus, posterior division | 2.13 | -66 | -28 | -16 |
| 8 | | | | | |
| | R Frontal Orbital Cortex | 2.28 | 22 | 14 | -16 |
| 8 | | | | | |
| | L Frontal Pole | 2.68 | -36 | 58 | -16 |
| 8 | | | | | |
| | R Frontal Pole | 2.20 | 26 | 48 | -12 |
| 8 | | | | | |
| | L Frontal Pole | 2.13 | -36 | 54 | -10 |
| 8 | | | | | |
| | R Frontal Pole | 2.13 | 14 | 62 | -10 |
| 8 | | | | | |
| | L Middle Temporal Gyrus, posterior division | 2.18 | -68 | -24 | -8 |
| 8 | | | | | |
| | R Putamen | 2.21 | 32 | -14 | -8 |
| 8 | | | | | |
| | L Frontal Pole | 2.68 | -18 | 60 | -8 |
| 8 | | | | | |
| | R Cingulate Gyrus, anterior division | 2.15 | 2 | 32 | -6 |
| 8 | | | | | |
| | L Brain-Stem | 2.20 | -6 | -36 | -4 |
| 8 | | | | | |
| | L Cingulate Gyrus, anterior division | 2.21 | -8 | 36 | -4 |
| 8 | | | | | |
| | L Caudate | 2.57 | -6 | 16 | 2 |
| 8 | | | | | |
| | R Heschls Gyrus (includes H1 and H2) | 2.30 | 44 | -20 | 8 |
| 8 | | | | | |
| | L Thalamus | 2.20 | -4 | -28 | 10 |
| 8 | | | | | |
| | R Middle Frontal Gyrus | 2.14 | 42 | 28 | 36 |
| 8 | | | | | |
| | L Frontal Pole | 2.39 | -4 | 44 | 52 |

**Supplementary Table S13. [CT$_{phasic}$ > CS$_{phasic}$].** Descriptive statistics for clusters and local maxima showing greater activation for Certain-Threat relative to Certain-Safety anticipation for the OSP Phasic regressor (FDR *q*<0.05, whole-brain corrected).

| *mm³* | Label | *t* | *x* | *y* | *z* |
|---|---|---|---|---|---|
| 80,016 | | | | | |
| | R Insular Cortex | 7.02 | 34 | 24 | 2 |
| | L Caudate | 6.60 | -14 | 22 | -2 |
| | L Frontal Orbital Cortex | 6.44 | -30 | 22 | -8 |
| | L Inferior Frontal Gyrus, pars triangularis | 3.92 | -50 | 22 | 12 |
| | R Caudate | 7.34 | 14 | 20 | -2 |
| | R Frontal Operculum Cortex | 8.62 | 36 | 18 | 8 |
| | R Frontal Orbital Cortex | 4.93 | 44 | 18 | -8 |
| | L Frontal Operculum Cortex | 8.12 | -42 | 16 | -2 |
| | L Insular Cortex | 7.89 | -36 | 14 | 4 |
| | R Inferior Frontal Gyrus, pars opercularis | 5.09 | 60 | 14 | 8 |
| | R Accumbens | 4.38 | 6 | 14 | -2 |
| | L Inferior Frontal Gyrus, pars opercularis | 6.44 | -58 | 12 | 4 |
| | L Accumbens | 6.19 | -6 | 10 | -4 |
| | L Putamen | 6.01 | -20 | 8 | 0 |
| | R Central Opercular Cortex | 7.32 | 48 | 8 | -2 |
| | R Temporal Pole | 3.37 | 38 | 8 | -22 |
| | L Temporal Pole | 3.88 | -30 | 6 | -20 |
| | R Putamen | 8.94 | 22 | 4 | -10 |
| | R Precentral Gyrus | 4.96 | 60 | 2 | 12 |
| | L Central Opercular Cortex | 5.87 | -50 | 0 | 2 |
| | L Amygdala | 3.83 | -32 | -2 | -18 |
| | L Pallidum | 5.58 | -20 | -2 | 0 |
| | R Thalamus | 4.32 | 0 | -8 | 2 |
| | L Planum Polare | 4.80 | -42 | -10 | -8 |
| | L Postcentral Gyrus | 4.73 | -46 | -10 | 30 |
| | L Precentral Gyrus | 5.94 | -36 | -16 | 50 |
| | R Planum Polare | 2.95 | 40 | -16 | -4 |
| | L Thalamus | 7.43 | -4 | -26 | 0 |
| | R Thalamus | 6.28 | 8 | -26 | 4 |
| | L Brain-Stem | 7.20 | -6 | -34 | -8 |
| | L Parahippocampal Gyrus, posterior division | 2.97 | -14 | -34 | -10 |
| | R Brain-Stem | 5.44 | 8 | -34 | -8 |
| | L Temporal Occipital Fusiform Cortex | 3.75 | -34 | -54 | -22 |
| | R Temporal Occipital Fusiform Cortex | 3.48 | 36 | -60 | -20 |
| | L Occipital Fusiform Gyrus | 3.95 | -24 | -68 | -16 |
| | R Occipital Fusiform Gyrus | 3.33 | 26 | -70 | -16 |
| | L Lingual Gyrus | 3.62 | -10 | -76 | -14 |
| 21,328 | | | | | |

| | | | | | |
|---|---|---|---|---|---|
| | L Paracingulate Gyrus | 4.46 | -6 | 36 | 22 |
| | L Cingulate Gyrus, anterior division | 5.79 | -8 | 14 | 36 |
| | R Superior Frontal Gyrus | 3.60 | 4 | 14 | 60 |
| | R Paracingulate Gyrus | 6.22 | 0 | 12 | 46 |
| | R Cingulate Gyrus, anterior division | 7.24 | 10 | 8 | 40 |
| | L Juxtapositional Lobule Cortex (formerly Supplementary Motor Cortex) | 6.32 | -10 | 2 | 42 |
| | R Juxtapositional Lobule Cortex (formerly Supplementary Motor Cortex) | 6.89 | 4 | -2 | 50 |
| | L Superior Frontal Gyrus | 5.59 | -12 | -8 | 66 |
| | L Precentral Gyrus | 3.86 | -14 | -20 | 72 |
| | R Precentral Gyrus | 4.22 | 8 | -28 | 50 |
| | R Cingulate Gyrus, posterior division | 4.64 | 10 | -32 | 46 |
| 10,120 | | | | | |
| | R Supramarginal Gyrus, anterior division | 4.92 | 62 | -28 | 26 |
| | R Parietal Operculum Cortex | 5.58 | 52 | -32 | 28 |
| | R Angular Gyrus | 5.48 | 56 | -46 | 32 |
| | R Supramarginal Gyrus, posterior division | 5.39 | 58 | -46 | 38 |
| 7,712 | | | | | |
| | L Postcentral Gyrus | 3.80 | -60 | -20 | 20 |
| | L Planum Temporale | 3.52 | -60 | -30 | 10 |
| | L Parietal Operculum Cortex | 7.12 | -44 | -34 | 22 |
| | L Supramarginal Gyrus, anterior division | 5.69 | -64 | -36 | 26 |
| | L Supramarginal Gyrus, posterior division | 4.89 | -64 | -48 | 34 |
| | L Angular Gyrus | 3.35 | -58 | -54 | 34 |
| 7,120 | | | | | |
| | R Middle Frontal Gyrus | 6.09 | 46 | 2 | 56 |
| | R Precentral Gyrus | 6.24 | 36 | -18 | 46 |
| | R Postcentral Gyrus | 5.19 | 30 | -28 | 60 |
| 4,624 | | | | | |
| | R Frontal Pole | 5.94 | 26 | 46 | 26 |
| 4,320 | | | | | |
| | R Lateral Occipital Cortex, inferior division | 4.62 | 38 | -78 | -12 |
| | R Occipital Fusiform Gyrus | 3.35 | 32 | -80 | -10 |
| | R Lateral Occipital Cortex, superior division | 3.37 | 32 | -88 | 16 |
| | R Occipital Pole | 4.45 | 36 | -90 | 8 |
| 4,280 | | | | | |
| | L Frontal Pole | 6.22 | -36 | 48 | 30 |
| | L Middle Frontal Gyrus | 4.18 | -32 | 30 | 42 |
| 3,128 | | | | | |
| | R Lingual Gyrus | 3.70 | 26 | -58 | 4 |
| | R Intracalcarine Cortex | 5.98 | 12 | -72 | 14 |
| | R Cuneal Cortex | 3.84 | 4 | -80 | 20 |
| 2,272 | | | | | |

| | | | | | |
|---|---|---|---|---|---|
| | L Lingual Gyrus | 3.40 | -10 | -62 | 2 |
| | L Intracalcarine Cortex | 6.45 | -14 | -72 | 8 |
| 1,768 | | | | | |
| | R Superior Temporal Gyrus, posterior division | 5.01 | 62 | -22 | -2 |
| | R Middle Temporal Gyrus, posterior division | 4.72 | 56 | -28 | -4 |
| 1,528 | | | | | |
| | R Superior Frontal Gyrus | 4.66 | 14 | -10 | 68 |
| | R Precentral Gyrus | 3.30 | 14 | -16 | 72 |
| 1,264 | | | | | |
| | L Precuneus Cortex | 5.77 | -6 | -72 | 46 |
| | L Cuneal Cortex | 3.23 | -18 | -72 | 26 |
| 1,032 | | | | | |
| | R Precuneus Cortex | 4.78 | 18 | -60 | 28 |
| | R Lateral Occipital Cortex, superior division | 3.36 | 20 | -70 | 48 |
| 920 | | | | | |
| | L Precentral Gyrus | 4.94 | -14 | -30 | 62 |
| 816 | | | | | |
| | L Occipital Fusiform Gyrus | 3.73 | -20 | -86 | -8 |
| | L Lateral Occipital Cortex, inferior division | 2.96 | -36 | -86 | -14 |
| | L Occipital Pole | 3.99 | -32 | -92 | -14 |
| 728 | | | | | |
| | L Precentral Gyrus | 3.69 | -10 | -22 | 46 |
| | L Cingulate Gyrus, posterior division | 4.55 | -10 | -26 | 40 |
| 504 | | | | | |
| | L Middle Temporal Gyrus, temporooccipital part | 4.56 | -58 | -60 | 10 |
| 440 | | | | | |
| | R Lateral Occipital Cortex, superior division | 4.42 | 22 | -86 | 30 |
| | R Occipital Pole | 3.49 | 12 | -90 | 26 |
| 360 | | | | | |
| | L Precuneus Cortex | 3.65 | -6 | -50 | 48 |
| 344 | | | | | |
| | L Thalamus | 4.08 | -16 | -26 | 8 |
| 328 | | | | | |
| | L Supramarginal Gyrus, posterior division | 3.47 | -52 | -50 | 14 |
| | L Angular Gyrus | 3.69 | -56 | -52 | 18 |
| 320 | | | | | |
| | R Parahippocampal Gyrus, anterior division | 4.16 | 24 | -8 | -32 |
| | R Temporal Fusiform Cortex, posterior division | 4.20 | 34 | -14 | -32 |

| | | | | | |
|---|---|---|---|---|---|
| 304 | | | | | |
| | R Temporal Pole | 3.95 | 46 | 8 | -40 |
| 280 | | | | | |
| | L Lateral Occipital Cortex, superior division | 2.93 | -26 | -88 | 18 |
| | L Occipital Pole | 3.26 | -30 | -92 | 18 |
| 232 | | | | | |
| | R Insular Cortex | 4.18 | 36 | -20 | 2 |
| 216 | | | | | |
| | L Insular Cortex | 3.84 | -40 | -18 | -2 |
| | L Putamen | 3.32 | -30 | -18 | -2 |
| 208 | | | | | |
| | R Hippocampus | 4.10 | 26 | -40 | 2 |
| 208 | | | | | |
| | L Precentral Gyrus | 3.53 | -22 | -14 | 58 |
| 200 | | | | | |
| | L Thalamus | 5.19 | -22 | -36 | 0 |
| 200 | | | | | |
| | L Frontal Pole | 3.99 | -34 | 42 | 18 |
| 192 | | | | | |
| | R Cingulate Gyrus, anterior division | 3.74 | 0 | 34 | 4 |
| | L Subcallosal Cortex | 4.13 | -2 | 28 | -2 |
| 176 | | | | | |
| | R Precentral Gyrus | 4.40 | 32 | -14 | 64 |
| 168 | | | | | |
| | R Frontal Medial Cortex | 3.48 | 2 | 36 | -28 |
| 168 | | | | | |
| | L Lateral Occipital Cortex, inferior division | 3.32 | -30 | -90 | -4 |
| | L Occipital Pole | 3.75 | -34 | -92 | -6 |
| 168 | | | | | |
| | L Caudate | 3.41 | -12 | 4 | 12 |
| 160 | | | | | |
| | L Middle Temporal Gyrus, posterior division | 3.18 | -60 | -22 | -14 |
| 160 | | | | | |
| | L Supramarginal Gyrus, posterior division | 3.34 | -56 | -48 | 50 |
| 144 | | | | | |
| | L Frontal Pole | 4.15 | -26 | 64 | -12 |
| 144 | | | | | |
| | R Middle Temporal Gyrus, temporooccipital part | 3.63 | 68 | -40 | 4 |
| 144 | | | | | |
| | R Superior Parietal Lobule | 3.46 | 24 | -42 | 60 |
| 136 | | | | | |
| | L Hippocampus | 3.76 | -26 | -34 | -8 |

| | | | | | |
|---|---|---|---|---|---|
| 136 | | | | | |
| | L Occipital Pole | 2.78 | -32 | -96 | 2 |
| 136 | | | | | |
| | L Frontal Pole | 3.38 | -26 | 50 | 16 |
| 136 | | | | | |
| | R Cingulate Gyrus, posterior division | 3.53 | 4 | -20 | 30 |
| 136 | | | | | |
| | L Postcentral Gyrus | 3.10 | -36 | -28 | 50 |
| 128 | | | | | |
| | R Temporal Occipital Fusiform Cortex | 3.32 | 28 | -56 | -18 |
| 128 | | | | | |
| | R Frontal Pole | 3.38 | 26 | 68 | -4 |
| 120 | | | | | |
| | R Middle Temporal Gyrus, anterior division | 3.11 | 54 | 2 | -36 |
| | R Inferior Temporal Gyrus, anterior division | 3.54 | 48 | 0 | -34 |
| 120 | | | | | |
| | L Brain-Stem | 4.11 | -12 | -30 | -30 |
| 120 | | | | | |
| | L Lateral Occipital Cortex, superior division | 3.29 | -32 | -60 | 40 |
| 104 | | | | | |
| | R Brain-Stem | 3.15 | 14 | -20 | -28 |
| 104 | | | | | |
| | R Occipital Fusiform Gyrus | 3.12 | 28 | -72 | -8 |
| 104 | | | | | |
| | L Insular Cortex | 3.66 | -36 | -20 | 12 |
| 96 | | | | | |
| | R Superior Temporal Gyrus, anterior division | 3.06 | 56 | 0 | -14 |
| 96 | | | | | |
| | R Putamen | 3.14 | 28 | -16 | 8 |
| 96 | | | | | |
| | R Inferior Frontal Gyrus, pars opercularis | 3.54 | 48 | 18 | 22 |
| 96 | | | | | |
| | L Precentral Gyrus | 3.51 | -6 | -26 | 50 |
| 88 | | | | | |
| | L Occipital Pole | 3.31 | -12 | -98 | 16 |
| 88 | | | | | |
| | R Precentral Gyrus | 3.05 | 62 | 4 | 32 |
| 80 | | | | | |
| | R Frontal Orbital Cortex | 3.58 | 36 | 26 | -16 |
| 80 | | | | | |
| | R Amygdala | 3.70 | 22 | -14 | -12 |

| | | | | | |
|---|---|---|---|---|---|
| 80 | | | | | |
| | R Frontal Pole | 3.32 | 28 | 62 | -10 |
| 80 | | | | | |
| | R Lateral Occipital Cortex, superior division | 2.77 | 42 | -76 | 20 |
| | R Lateral Occipital Cortex, inferior division | 2.77 | 42 | -78 | 14 |
| 72 | | | | | |
| | L Occipital Fusiform Gyrus | 3.10 | -36 | -78 | -14 |
| 72 | | | | | |
| | R Precuneus Cortex | 3.47 | 2 | -44 | 48 |
| 64 | | | | | |
| | R Brain-Stem | 3.24 | 16 | -32 | -42 |
| 64 | | | | | |
| | R Inferior Temporal Gyrus, posterior division | 3.59 | 58 | -20 | -28 |
| 64 | | | | | |
| | L Amygdala | 4.88 | -18 | -12 | -14 |
| 64 | | | | | |
| | R Precentral Gyrus | 2.56 | 40 | 2 | 32 |
| 56 | | | | | |
| | L Middle Temporal Gyrus, posterior division | 2.81 | -62 | -30 | -12 |
| 56 | | | | | |
| | R Paracingulate Gyrus | 3.27 | 2 | 46 | -2 |
| 56 | | | | | |
| | L Thalamus | 3.40 | -10 | -10 | 6 |
| 56 | | | | | |
| | R Frontal Pole | 3.53 | 26 | 50 | 42 |
| 48 | | | | | |
| | R Brain-Stem | 3.47 | 6 | -28 | -42 |
| 48 | | | | | |
| | R Middle Temporal Gyrus, anterior division | 3.03 | 58 | -4 | -34 |
| 48 | | | | | |
| | L Hippocampus | 3.36 | -30 | -10 | -26 |
| 48 | | | | | |
| | L Lateral Occipital Cortex, inferior division | 2.94 | -48 | -78 | -16 |
| 48 | | | | | |
| | L Frontal Pole | 2.99 | -24 | 38 | -16 |
| 48 | | | | | |
| | L Frontal Pole | 3.42 | -22 | 66 | -2 |
| 48 | | | | | |
| | R Superior Temporal Gyrus, posterior division | 3.36 | 70 | -20 | 6 |
| 48 | | | | | |

| | | | | | |
|---|---|---|---|---|---|
| | R Angular Gyrus | 2.99 | 60 | -56 | 18 |
| 48 | | | | | |
| | L Lateral Occipital Cortex, superior division | 2.85 | -40 | -66 | 20 |
| 48 | | | | | |
| | R Caudate | 3.39 | 18 | -12 | 22 |
| 48 | | | | | |
| | R Cuneal Cortex | 3.39 | 10 | -78 | 30 |
| 48 | | | | | |
| | R Lateral Occipital Cortex, superior division | 3.58 | 26 | -78 | 32 |
| 48 | | | | | |
| | R Precentral Gyrus | 2.82 | 50 | -8 | 32 |
| 48 | | | | | |
| | L Precentral Gyrus | 3.08 | -8 | -22 | 58 |
| 48 | | | | | |
| | R Superior Frontal Gyrus | 2.79 | 26 | 2 | 62 |
| 48 | | | | | |
| | L Superior Frontal Gyrus | 3.14 | -14 | 2 | 68 |
| 40 | | | | | |
| | L Temporal Pole | 2.95 | -52 | 4 | -32 |
| 40 | | | | | |
| | L Caudate | 3.16 | -14 | -4 | 20 |
| 40 | | | | | |
| | R Precentral Gyrus | 3.09 | 56 | 6 | 28 |
| 40 | | | | | |
| | R Lateral Occipital Cortex, superior division | 2.80 | 16 | -84 | 30 |
| | R Occipital Pole | 2.94 | 14 | -88 | 32 |
| 40 | | | | | |
| | L Middle Frontal Gyrus | 3.09 | -42 | 20 | 34 |
| 40 | | | | | |
| | R Superior Parietal Lobule | 3.17 | 16 | -48 | 64 |
| 32 | | | | | |
| | L Inferior Temporal Gyrus, anterior division | 3.25 | -50 | -2 | -34 |
| 32 | | | | | |
| | R Temporal Pole | 2.94 | 54 | 10 | -34 |
| 32 | | | | | |
| | L Inferior Temporal Gyrus, posterior division | 3.25 | -60 | -30 | -20 |
| 32 | | | | | |
| | R Hippocampus | 2.66 | 36 | -30 | -12 |
| 32 | | | | | |
| | L Lateral Occipital Cortex, inferior division | 3.14 | -40 | -70 | -10 |

| | | | | | |
|---|---|---|---|---|---|
| 32 | | | | | |
| | R Middle Temporal Gyrus, temporooccipital part | 2.76 | 66 | -46 | 2 |
| 32 | | | | | |
| | L Caudate | 3.18 | -8 | 12 | 6 |
| 32 | | | | | |
| | R Supramarginal Gyrus, posterior division | 3.04 | 66 | -42 | 10 |
| 32 | | | | | |
| | L Lateral Occipital Cortex, superior division | 2.83 | -36 | -84 | 14 |
| 32 | | | | | |
| | L Lateral Occipital Cortex, superior division | 2.72 | -56 | -66 | 22 |
| 32 | | | | | |
| | L Caudate | 3.36 | -14 | -10 | 20 |
| 32 | | | | | |
| | L Frontal Pole | 3.56 | -46 | 46 | 20 |
| 32 | | | | | |
| | L Cingulate Gyrus, posterior division | 3.59 | -6 | -38 | 26 |
| 32 | | | | | |
| | R Cingulate Gyrus, anterior division | 3.86 | 4 | 6 | 28 |
| 32 | | | | | |
| | R Cingulate Gyrus, posterior division | 3.13 | 2 | -42 | 40 |
| 32 | | | | | |
| | R Postcentral Gyrus | 2.94 | 20 | -36 | 62 |
| 32 | | | | | |
| | L Precentral Gyrus | 2.76 | -28 | -14 | 66 |
| 24 | | | | | |
| | L Brain-Stem | 3.11 | -10 | -30 | -40 |
| 24 | | | | | |
| | L Parahippocampal Gyrus, anterior division | 3.32 | -22 | -6 | -36 |
| 24 | | | | | |
| | R Brain-Stem | 3.31 | 8 | -24 | -32 |
| 24 | | | | | |
| | R Inferior Temporal Gyrus, posterior division | 2.84 | 62 | -32 | -26 |
| 24 | | | | | |
| | R Frontal Orbital Cortex | 2.84 | 16 | 24 | -26 |
| 24 | | | | | |
| | L Hippocampus | 2.99 | -26 | -18 | -20 |
| 24 | | | | | |
| | R Middle Temporal Gyrus, posterior division | 3.47 | 48 | -14 | -16 |
| 24 | | | | | |

| | | | | | |
|---|---|---|---|---|---|
| | R Frontal Pole | 2.83 | 34 | 34 | -14 |
| 24 | | | | | |
| | L Occipital Pole | 3.00 | -16 | -92 | -10 |
| 24 | | | | | |
| | L Lateral Occipital Cortex, inferior division | 2.94 | -50 | -80 | -10 |
| 24 | | | | | |
| | L Planum Polare | 3.24 | -46 | -10 | -2 |
| 24 | | | | | |
| | R Cingulate Gyrus, anterior division | 2.76 | 4 | 34 | 10 |
| 24 | | | | | |
| | L Frontal Pole | 2.82 | -42 | 50 | 14 |
| 24 | | | | | |
| | L Supramarginal Gyrus, posterior division | 2.88 | -64 | -50 | 18 |
| 24 | | | | | |
| | L Occipital Pole | 3.26 | -14 | -92 | 22 |
| 24 | | | | | |
| | L Lateral Occipital Cortex, superior division | 2.97 | -34 | -78 | 28 |
| 24 | | | | | |
| | R Precentral Gyrus | 2.64 | 44 | -14 | 38 |
| 24 | | | | | |
| | R Postcentral Gyrus | 3.79 | 46 | -28 | 54 |
| 16 | | | | | |
| | R Temporal Fusiform Cortex, anterior division | 2.59 | 30 | -6 | -44 |
| 16 | | | | | |
| | R Middle Temporal Gyrus, posterior division | 2.91 | 60 | -10 | -30 |
| 16 | | | | | |
| | L Temporal Pole | 3.40 | -34 | 10 | -30 |
| 16 | | | | | |
| | R Temporal Fusiform Cortex, posterior division | 2.61 | 42 | -32 | -26 |
| 16 | | | | | |
| | R Hippocampus | 2.81 | 32 | -8 | -26 |
| 16 | | | | | |
| | L Middle Temporal Gyrus, anterior division | 2.90 | -56 | -8 | -26 |
| 16 | | | | | |
| | L Inferior Temporal Gyrus, posterior division | 2.66 | -50 | -42 | -24 |
| 16 | | | | | |
| | L Temporal Pole | 3.21 | -42 | 14 | -24 |
| 16 | | | | | |

| | | | | | |
|---|---|---|---|---|---|
| | L Middle Temporal Gyrus, posterior division | 2.71 | -64 | -16 | -22 |
| 16 | | | | | |
| | R Hippocampus | 2.68 | 26 | -20 | -14 |
| 16 | | | | | |
| | R Hippocampus | 2.64 | 28 | -28 | -12 |
| 16 | | | | | |
| | L Frontal Pole | 2.65 | -16 | 62 | -12 |
| 16 | | | | | |
| | L Middle Temporal Gyrus, posterior division | 2.78 | -66 | -24 | -10 |
| 16 | | | | | |
| | L Frontal Pole | 2.96 | -14 | 68 | -6 |
| 16 | | | | | |
| | R Lateral Occipital Cortex, inferior division | 3.13 | 36 | -80 | 2 |
| 16 | | | | | |
| | L Lingual Gyrus | 2.87 | -20 | -60 | 2 |
| 16 | | | | | |
| | R Insular Cortex | 2.63 | 42 | -14 | 2 |
| 16 | | | | | |
| | R Paracingulate Gyrus | 3.14 | 8 | 46 | 4 |
| 16 | | | | | |
| | L Occipital Pole | 2.68 | -34 | -94 | 6 |
| 16 | | | | | |
| | R Middle Temporal Gyrus, temporooccipital part | 2.76 | 62 | -52 | 6 |
| 16 | | | | | |
| | L Central Opercular Cortex | 3.18 | -46 | -10 | 6 |
| 16 | | | | | |
| | L Insular Cortex | 2.72 | -32 | -26 | 8 |
| 16 | | | | | |
| | R Cingulate Gyrus, anterior division | 3.07 | 12 | 40 | 8 |
| 16 | | | | | |
| | L Middle Temporal Gyrus, temporooccipital part | 2.64 | -46 | -54 | 10 |
| 16 | | | | | |
| | L Thalamus | 2.87 | -8 | -2 | 10 |
| 16 | | | | | |
| | R Occipital Pole | 2.89 | 10 | -90 | 14 |
| 16 | | | | | |
| | R Frontal Pole | 2.65 | 38 | 42 | 18 |
| 16 | | | | | |
| | R Parietal Operculum Cortex | 2.92 | 44 | -20 | 18 |
| 16 | | | | | |
| | R Parietal Operculum Cortex | 2.67 | 38 | -30 | 20 |

| | | | | | |
|---|---|---|---|---|---|
| 16 | | | | | |
| | R Precentral Gyrus | 2.82 | 62 | 2 | 22 |
| 16 | | | | | |
| | R Lateral Occipital Cortex, superior division | 3.22 | 40 | -74 | 24 |
| 16 | | | | | |
| | R Cingulate Gyrus, posterior division | 2.64 | 6 | -44 | 34 |
| 16 | | | | | |
| | L Cingulate Gyrus, posterior division | 2.59 | -12 | -46 | 34 |
| 16 | | | | | |
| | L Middle Frontal Gyrus | 2.74 | -38 | 20 | 40 |
| 16 | | | | | |
| | L Precentral Gyrus | 3.15 | -8 | -34 | 48 |
| 16 | | | | | |
| | R Superior Frontal Gyrus | 3.37 | 6 | 36 | 48 |
| 16 | | | | | |
| | R Postcentral Gyrus | 2.81 | 54 | -14 | 50 |
| 16 | | | | | |
| | R Superior Frontal Gyrus | 2.63 | 22 | 0 | 56 |
| 16 | | | | | |
| | R Precuneus Cortex | 2.53 | 12 | -56 | 56 |
| 8 | | | | | |
| | L Brain-Stem | 2.67 | -4 | -44 | -60 |
| 8 | | | | | |
| | L Temporal Pole | 2.77 | -36 | 14 | -44 |
| 8 | | | | | |
| | L Brain-Stem | 2.69 | -12 | -34 | -42 |
| 8 | | | | | |
| | L Brain-Stem | 2.87 | -2 | -26 | -42 |
| 8 | | | | | |
| | L Brain-Stem | 2.54 | -6 | -22 | -40 |
| 8 | | | | | |
| | R Temporal Fusiform Cortex, posterior division | 2.68 | 42 | -16 | -38 |
| 8 | | | | | |
| | L Brain-Stem | 2.65 | -20 | -30 | -34 |
| 8 | | | | | |
| | L Brain-Stem | 2.59 | -20 | -26 | -34 |
| 8 | | | | | |
| | R Inferior Temporal Gyrus, posterior division | 2.56 | 46 | -20 | -34 |
| 8 | | | | | |
| | R Inferior Temporal Gyrus, posterior division | 2.67 | 46 | -22 | -28 |
| 8 | | | | | |

| | | | | | |
|---|---|---|---|---|---|
| | L Brain-Stem | 2.57 | -6 | -20 | -28 |
| 8 | | | | | |
| | L Inferior Temporal Gyrus, posterior division | 2.58 | -56 | -44 | -24 |
| 8 | | | | | |
| | R Amygdala | 2.97 | 30 | 4 | -22 |
| | R Amygdala | 2.97 | 30 | 4 | -22 |
| 8 | | | | | |
| | R Inferior Temporal Gyrus, temporooccipital part | 2.55 | 60 | -50 | -20 |
| 8 | | | | | |
| | R Inferior Temporal Gyrus, posterior division | 2.60 | 58 | -28 | -20 |
| 8 | | | | | |
| | L Middle Temporal Gyrus, posterior division | 2.54 | -66 | -24 | -18 |
| 8 | | | | | |
| | L Hippocampus | 2.81 | -34 | -16 | -18 |
| 8 | | | | | |
| | R Middle Temporal Gyrus, posterior division | 2.54 | 52 | -12 | -18 |
| 8 | | | | | |
| | R Hippocampus | 2.62 | 24 | -12 | -18 |
| 8 | | | | | |
| | L Inferior Temporal Gyrus, temporooccipital part | 2.81 | -52 | -60 | -16 |
| 8 | | | | | |
| | R Lateral Occipital Cortex, inferior division | 2.65 | 50 | -76 | -14 |
| 8 | | | | | |
| | R Occipital Fusiform Gyrus | 2.71 | 32 | -72 | -14 |
| 8 | | | | | |
| | R Parahippocampal Gyrus, posterior division | 2.71 | 18 | -30 | -14 |
| 8 | | | | | |
| | L Frontal Pole | 2.56 | -28 | 50 | -14 |
| 8 | | | | | |
| | L Frontal Pole | 3.37 | -18 | 54 | -14 |
| 8 | | | | | |
| | L Frontal Orbital Cortex | 2.82 | -26 | 36 | -12 |
| 8 | | | | | |
| | R Middle Temporal Gyrus, posterior division | 2.62 | 60 | -12 | -10 |
| 8 | | | | | |
| | L Superior Temporal Gyrus, anterior division | 2.54 | -56 | -4 | -10 |
| 8 | | | | | |

| | | | | | |
|---|---|---|---|---|---|
| | R Planum Polare | 2.57 | 50 | 4 | -8 |
| 8 | | | | | |
| | L Subcallosal Cortex | 2.54 | -6 | 26 | -8 |
| 8 | | | | | |
| | L Planum Polare | 2.61 | -48 | -8 | -6 |
| 8 | | | | | |
| | L Middle Temporal Gyrus, temporooccipital part | 2.73 | -52 | -60 | -4 |
| 8 | | | | | |
| | R Middle Temporal Gyrus, temporooccipital part | 2.54 | 60 | -54 | -2 |
| 8 | | | | | |
| | L Lingual Gyrus | 2.73 | -18 | -50 | -2 |
| 8 | | | | | |
| | L Frontal Pole | 2.61 | -14 | 70 | -2 |
| 8 | | | | | |
| | L Pallidum | 2.74 | -24 | -10 | 0 |
| 8 | | | | | |
| | L Lateral Occipital Cortex, inferior division | 2.52 | -40 | -74 | 4 |
| 8 | | | | | |
| | L Lingual Gyrus | 2.72 | -4 | -70 | 4 |
| 8 | | | | | |
| | R Inferior Frontal Gyrus, pars triangularis | 2.56 | 50 | 32 | 4 |
| 8 | | | | | |
| | R Superior Temporal Gyrus, posterior division | 2.52 | 70 | -30 | 8 |
| 8 | | | | | |
| | L Heschls Gyrus (includes H1 and H2) | 2.51 | -42 | -18 | 10 |
| 8 | | | | | |
| | L Putamen | 2.57 | -24 | 4 | 12 |
| 8 | | | | | |
| | L Occipital Pole | 2.76 | -20 | -92 | 14 |
| 8 | | | | | |
| | L Lateral Occipital Cortex, superior division | 2.53 | -44 | -70 | 16 |
| 8 | | | | | |
| | R Cingulate Gyrus, anterior division | 2.56 | 4 | 24 | 16 |
| 8 | | | | | |
| | L Frontal Pole | 2.54 | -26 | 56 | 16 |
| 8 | | | | | |
| | L Thalamus | 2.56 | -8 | -14 | 18 |
| 8 | | | | | |
| | L Central Opercular Cortex | 2.82 | -40 | -8 | 18 |
| 8 | | | | | |
| | L Caudate | 2.59 | -18 | 6 | 18 |

| | | | | | |
|---|---|---|---|---|---|
| 8 | | | | | |
| | R Precentral Gyrus | 2.65 | 58 | 4 | 20 |
| 8 | | | | | |
| | R Frontal Pole | 2.62 | 42 | 42 | 20 |
| 8 | | | | | |
| | L Occipital Pole | 2.62 | -20 | -96 | 22 |
| 8 | | | | | |
| | R Cingulate Gyrus, posterior division | 2.60 | 4 | -40 | 24 |
| 8 | | | | | |
| | L Postcentral Gyrus | 3.02 | -56 | -14 | 26 |
| 8 | | | | | |
| | L Frontal Pole | 2.53 | -26 | 44 | 26 |
| 8 | | | | | |
| | R Frontal Pole | 3.12 | 10 | 58 | 26 |
| 8 | | | | | |
| | R Frontal Pole | 2.56 | 46 | 44 | 28 |
| 8 | | | | | |
| | L Frontal Pole | 2.64 | -24 | 46 | 28 |
| 8 | | | | | |
| | R Frontal Pole | 2.65 | 24 | 38 | 32 |
| 8 | | | | | |
| | R Middle Frontal Gyrus | 2.72 | 38 | 26 | 34 |
| 8 | | | | | |
| | L Cingulate Gyrus, posterior division | 2.51 | -10 | -38 | 36 |
| 8 | | | | | |
| | R Precentral Gyrus | 2.60 | 56 | 8 | 36 |
| 8 | | | | | |
| | R Superior Frontal Gyrus | 2.62 | 8 | 40 | 38 |
| 8 | | | | | |
| | L Precuneus Cortex | 2.65 | -10 | -48 | 42 |
| 8 | | | | | |
| | L Cingulate Gyrus, posterior division | 2.69 | -4 | -18 | 42 |
| 8 | | | | | |
| | L Frontal Pole | 2.54 | -26 | 44 | 44 |
| 8 | | | | | |
| | L Lateral Occipital Cortex, superior division | 2.53 | -26 | -60 | 46 |
| 8 | | | | | |
| | L Postcentral Gyrus | 2.63 | -38 | -24 | 46 |
| 8 | | | | | |
| | R Middle Frontal Gyrus | 2.59 | 34 | 28 | 46 |
| 8 | | | | | |
| | L Precentral Gyrus | 2.53 | -32 | -10 | 58 |
| 8 | | | | | |

| | | | | | |
|---|---|---|---|---|---|
| | R Superior Frontal Gyrus | 2.62 | 20 | 0 | 60 |
| 8 | | | | | |
| | L Precentral Gyrus | 2.60 | -24 | -20 | 68 |
| 8 | | | | | |
| | L Precentral Gyrus | 2.64 | -18 | -24 | 70 |

**Supplementary Table S14. [$UT_{phasic}$ > $US_{phasic}$].** Descriptive statistics for clusters and local maxima showing greater activation for Uncertain-Threat relative to Uncertain-Safety anticipation for the OSP Phasic regressor (FDR $q<0.05$, whole-brain corrected).

| ***mm³*** | **Label** | ***t*** | ***x*** | ***y*** | ***z*** |
|---|---|---|---|---|---|
| 4,088 | | | | | |
| | R Inferior Frontal Gyrus, pars triangularis | 3.47 | 50 | 24 | 8 |
| | R Insular Cortex | 6.17 | 38 | 20 | -4 |
| | R Temporal Pole | 4.40 | 50 | 20 | -14 |
| | R Frontal Orbital Cortex | 3.82 | 38 | 20 | -16 |
| | R Frontal Operculum Cortex | 5.87 | 46 | 14 | -2 |
| | R Inferior Frontal Gyrus, pars opercularis | 3.43 | 56 | 12 | 4 |
| | R Central Opercular Cortex | 4.52 | 38 | 10 | 8 |
| | R Precentral Gyrus | 3.56 | 60 | 6 | 6 |
| 3,944 | | | | | |
| | R Postcentral Gyrus | 3.82 | 64 | -18 | 22 |
| | R Supramarginal Gyrus, anterior division | 4.77 | 64 | -26 | 26 |
| | R Parietal Operculum Cortex | 4.87 | 56 | -28 | 24 |
| | R Supramarginal Gyrus, posterior division | 4.01 | 56 | -40 | 20 |
| | R Middle Temporal Gyrus, temporooccipital part | 5.93 | 64 | -44 | 8 |
| | R Angular Gyrus | 4.40 | 54 | -48 | 16 |
| 3,776 | | | | | |
| | R Thalamus | 6.64 | 6 | -22 | 6 |
| | L Thalamus | 5.01 | -6 | -24 | 10 |
| | L Brain-Stem | 6.22 | -4 | -34 | -6 |
| | R Brain-Stem | 5.78 | 4 | -36 | -8 |
| 3,008 | | | | | |
| | L Postcentral Gyrus | 3.70 | -62 | -18 | 20 |
| | L Supramarginal Gyrus, anterior division | 5.60 | -64 | -28 | 20 |
| | L Supramarginal Gyrus, posterior division | 5.64 | -60 | -44 | 18 |
| | L Angular Gyrus | 4.17 | -54 | -54 | 44 |
| 2,744 | | | | | |
| | R Frontal Pole | 5.60 | 32 | 50 | 28 |
| | R Middle Frontal Gyrus | 3.68 | 42 | 34 | 36 |
| 2,640 | | | | | |
| | L Frontal Orbital Cortex | 4.80 | -32 | 24 | -6 |
| | L Frontal Operculum Cortex | 5.14 | -40 | 14 | 0 |
| | L Insular Cortex | 4.40 | -44 | 10 | -6 |
| | L Central Opercular Cortex | 4.59 | -38 | 8 | 6 |
| | L Putamen | 5.17 | -30 | 6 | 0 |
| | L Pallidum | 3.80 | -16 | 4 | -2 |

| | | | | | |
|---|---|---|---|---|---|
| | L Planum Polare | 3.55 | -54 | -2 | -2 |
| 2,176 | | | | | |
| | R Postcentral Gyrus | 4.44 | 60 | -8 | 30 |
| | R Precentral Gyrus | 5.65 | 40 | -12 | 48 |
| 2,144 | | | | | |
| | L Precentral Gyrus | 6.15 | -46 | -12 | 32 |
| | L Postcentral Gyrus | 4.39 | -40 | -18 | 42 |
| 1,824 | | | | | |
| | L Precentral Gyrus | 6.53 | -20 | -30 | 58 |
| | L Postcentral Gyrus | 4.27 | -20 | -34 | 68 |
| 1,624 | | | | | |
| | R Precentral Gyrus | 7.06 | 16 | -26 | 70 |
| | R Postcentral Gyrus | 4.42 | 24 | -28 | 58 |
| 1,360 | | | | | |
| | R Intracalcarine Cortex | 4.97 | 10 | -76 | 14 |
| 1,184 | | | | | |
| | R Frontal Pole | 4.69 | 42 | 44 | 4 |
| 1,136 | | | | | |
| | L Intracalcarine Cortex | 5.15 | -8 | -70 | 8 |
| 1,104 | | | | | |
| | L Insular Cortex | 4.82 | -42 | -10 | -4 |
| | L Putamen | 4.53 | -32 | -10 | -4 |
| | L Planum Polare | 4.93 | -44 | -14 | -2 |
| 1,000 | | | | | |
| | R Putamen | 4.20 | 22 | 10 | 0 |
| | R Pallidum | 4.33 | 20 | 2 | -2 |
| 992 | | | | | |
| | R Supramarginal Gyrus, anterior division | 3.79 | 58 | -34 | 48 |
| | R Supramarginal Gyrus, posterior division | 5.04 | 60 | -46 | 42 |
| | R Angular Gyrus | 3.54 | 50 | -48 | 50 |
| 464 | | | | | |
| | L Central Opercular Cortex | 3.62 | -38 | -20 | 20 |
| | L Insular Cortex | 4.88 | -32 | -28 | 16 |
| | L Parietal Operculum Cortex | 4.21 | -36 | -30 | 20 |
| 408 | | | | | |
| | L Cingulate Gyrus, posterior division | 4.67 | -6 | -20 | 46 |
| | L Precentral Gyrus | 4.23 | -12 | -24 | 44 |
| 400 | | | | | |
| | R Insular Cortex | 3.43 | 42 | -10 | 4 |
| | R Planum Polare | 4.14 | 42 | -14 | -4 |
| 352 | | | | | |
| | R Paracingulate Gyrus | 4.58 | 8 | 18 | 36 |
| 344 | | | | | |
| | L Precentral Gyrus | 5.35 | -8 | -26 | 58 |

| | | | | | |
|---|---|---|---|---|---|
| 312 | | | | | |
| | R Precentral Gyrus | 4.99 | 6 | -30 | 60 |
| 272 | | | | | |
| | R Cingulate Gyrus, anterior division | 5.21 | 0 | 32 | 0 |
| | R Subcallosal Cortex | 3.90 | 0 | 26 | -4 |
| 264 | | | | | |
| | L Middle Temporal Gyrus, temporooccipital part | 4.28 | -54 | -54 | 6 |
| 264 | | | | | |
| | L Paracingulate Gyrus | 4.79 | -4 | 30 | 32 |
| 248 | | | | | |
| | R Lateral Occipital Cortex, superior division | 3.79 | 50 | -58 | 46 |
| | R Angular Gyrus | 3.19 | 52 | -58 | 32 |
| 224 | | | | | |
| | L Postcentral Gyrus | 4.93 | -60 | -6 | 16 |
| 200 | | | | | |
| | L Inferior Temporal Gyrus, posterior division | 3.88 | -52 | -42 | -24 |
| 200 | | | | | |
| | R Thalamus | 5.73 | 6 | -4 | -2 |
| 160 | | | | | |
| | L Frontal Pole | 4.76 | -26 | 52 | 10 |
| 152 | | | | | |
| | R Middle Temporal Gyrus, posterior division | 3.88 | 64 | -30 | 0 |
| 152 | | | | | |
| | R Caudate | 5.06 | 14 | 24 | -2 |
| 144 | | | | | |
| | L Cingulate Gyrus, anterior division | 4.22 | -8 | 26 | 26 |
| 136 | | | | | |
| | R Frontal Pole | 4.12 | 26 | 34 | -10 |
| 112 | | | | | |
| | L Insular Cortex | 4.33 | -40 | 10 | -12 |
| 112 | | | | | |
| | L Parietal Operculum Cortex | 4.20 | -48 | -38 | 22 |
| 104 | | | | | |
| | R Frontal Pole | 3.61 | 26 | 64 | -2 |
| 104 | | | | | |
| | R Angular Gyrus | 3.86 | 60 | -52 | 24 |
| 96 | | | | | |
| | R Brain-Stem | 3.64 | 4 | -20 | -24 |
| 96 | | | | | |
| | L Inferior Temporal Gyrus, temporooccipital part | 3.44 | -60 | -48 | -14 |

| | | | | | |
|---|---|---|---|---|---|
| 96 | | | | | |
| | L Angular Gyrus | 3.76 | -48 | -54 | 28 |
| | L Lateral Occipital Cortex, superior division | 3.61 | -46 | -62 | 26 |
| 96 | | | | | |
| | L Lateral Occipital Cortex, superior division | 3.96 | -30 | -80 | 32 |
| 96 | | | | | |
| | R Cingulate Gyrus, posterior division | 3.53 | 6 | -36 | 34 |
| 88 | | | | | |
| | R Cingulate Gyrus, posterior division | 3.94 | 0 | -22 | 28 |
| 88 | | | | | |
| | R Paracingulate Gyrus | 3.28 | 2 | 16 | 46 |
| | L Paracingulate Gyrus | 3.54 | -2 | 12 | 46 |
| 80 | | | | | |
| | R Frontal Pole | 3.79 | 22 | 44 | -14 |
| 80 | | | | | |
| | L Frontal Pole | 3.51 | -28 | 52 | 30 |
| 72 | | | | | |
| | L Brain-Stem | 3.42 | -8 | -38 | -26 |
| 72 | | | | | |
| | R Putamen | 4.47 | 24 | -4 | 6 |
| 64 | | | | | |
| | R Insular Cortex | 3.85 | 38 | -18 | 2 |
| 64 | | | | | |
| | L Frontal Pole | 3.16 | -50 | 38 | 10 |
| | L Inferior Frontal Gyrus, pars triangularis | 3.34 | -50 | 34 | 10 |
| 56 | | | | | |
| | L Brain-Stem | 3.83 | -8 | -28 | -12 |
| 56 | | | | | |
| | R Middle Temporal Gyrus, posterior division | 3.67 | 64 | -22 | -6 |
| 56 | | | | | |
| | R Frontal Pole | 3.32 | 48 | 48 | -4 |
| 56 | | | | | |
| | L Frontal Pole | 4.18 | -26 | 52 | 4 |
| 56 | | | | | |
| | R Middle Frontal Gyrus | 3.65 | 52 | 28 | 26 |
| 56 | | | | | |
| | R Cingulate Gyrus, anterior division | 3.34 | 0 | 16 | 32 |
| 56 | | | | | |
| | R Superior Frontal Gyrus | 4.21 | 4 | 34 | 58 |
| 48 | | | | | |

| | | | | | |
|---|---|---|---|---|---|
| | L Brain-Stem | 3.45 | -6 | -40 | -52 |
| | R Brain-Stem | 3.54 | 0 | -40 | -52 |
| 48 | | | | | |
| | R Middle Temporal Gyrus, posterior division | 3.59 | 58 | -28 | -2 |
| 48 | | | | | |
| | L Insular Cortex | 4.22 | -34 | -24 | 6 |
| 48 | | | | | |
| | L Precentral Gyrus | 3.53 | -58 | 4 | 10 |
| 48 | | | | | |
| | R Insular Cortex | 3.39 | 36 | -12 | 14 |
| 48 | | | | | |
| | L Postcentral Gyrus | 3.86 | -60 | -20 | 34 |
| 48 | | | | | |
| | L Precuneus Cortex | 3.18 | -8 | -68 | 34 |
| 48 | | | | | |
| | R Frontal Pole | 4.19 | 24 | 44 | 46 |
| 40 | | | | | |
| | R Frontal Orbital Cortex | 3.92 | 28 | 26 | -16 |
| 40 | | | | | |
| | L Frontal Orbital Cortex | 3.29 | -40 | 20 | -14 |
| 40 | | | | | |
| | L Parahippocampal Gyrus, posterior division | 3.72 | -20 | -38 | -12 |
| 40 | | | | | |
| | L Putamen | 3.67 | -20 | 2 | -8 |
| 40 | | | | | |
| | R Frontal Pole | 4.30 | 26 | 52 | -2 |
| 40 | | | | | |
| | L Frontal Pole | 3.10 | -48 | 46 | -2 |
| 40 | | | | | |
| | R Frontal Pole | 3.56 | 12 | 58 | 26 |
| 40 | | | | | |
| | L Middle Frontal Gyrus | 3.55 | -46 | 12 | 36 |
| 40 | | | | | |
| | L Lateral Occipital Cortex, superior division | 3.17 | -42 | -70 | 42 |
| 40 | | | | | |
| | R Precentral Gyrus | 3.82 | 24 | -14 | 64 |
| 40 | | | | | |
| | L Precentral Gyrus | 3.19 | -16 | -16 | 66 |
| 32 | | | | | |
| | R Temporal Pole | 3.60 | 34 | 24 | -42 |
| 32 | | | | | |
| | L Brain-Stem | 3.33 | -4 | -16 | -22 |
| 32 | | | | | |

| | | | | | |
|---|---|---|---|---|---|
| | R Lateral Occipital Cortex, inferior division | 3.40 | 58 | -62 | -4 |
| 32 | | | | | |
| | L Insular Cortex | 3.46 | -42 | -4 | 2 |
| 32 | | | | | |
| | R Occipital Pole | 3.60 | 0 | -98 | 12 |
| 32 | | | | | |
| | L Lateral Occipital Cortex, superior division | 3.36 | -34 | -84 | 8 |
| 32 | | | | | |
| | L Central Opercular Cortex | 3.86 | -44 | -14 | 10 |
| 32 | | | | | |
| | R Cingulate Gyrus, anterior division | 3.68 | 8 | 34 | 20 |
| 32 | | | | | |
| | L Inferior Frontal Gyrus, pars opercularis | 3.23 | -58 | 14 | 22 |
| 32 | | | | | |
| | R Paracingulate Gyrus | 3.60 | 0 | 38 | 22 |
| 32 | | | | | |
| | R Middle Frontal Gyrus | 3.59 | 50 | 34 | 28 |
| 32 | | | | | |
| | L Paracingulate Gyrus | 3.36 | -4 | 42 | 28 |
| 32 | | | | | |
| | R Lateral Occipital Cortex, superior division | 3.97 | 28 | -76 | 32 |
| 32 | | | | | |
| | R Frontal Pole | 3.27 | 4 | 58 | 34 |
| 32 | | | | | |
| | L Frontal Pole | 3.60 | -36 | 40 | 38 |
| 32 | | | | | |
| | L Cingulate Gyrus, anterior division | 3.12 | -6 | -8 | 44 |
| 32 | | | | | |
| | R Precentral Gyrus | 3.60 | 6 | -20 | 48 |
| 32 | | | | | |
| | R Supramarginal Gyrus, posterior division | 3.02 | 52 | -38 | 50 |
| 32 | | | | | |
| | R Precentral Gyrus | 3.71 | 44 | -8 | 56 |
| 32 | | | | | |
| | L Precentral Gyrus | 4.18 | -4 | -36 | 58 |
| 32 | | | | | |
| | R Precentral Gyrus | 3.45 | 6 | -22 | 66 |
| 32 | | | | | |
| | R Postcentral Gyrus | 3.49 | 24 | -32 | 70 |
| 24 | | | | | |

| | | | | | |
|---|---|---|---|---|---|
| | L Brain-Stem | 3.56 | -8 | -44 | -38 |
| 24 | | | | | |
| | L Lateral Occipital Cortex, inferior division | 3.45 | -36 | -82 | -18 |
| 24 | | | | | |
| | R Superior Temporal Gyrus, anterior division | 3.84 | 60 | 4 | -14 |
| 24 | | | | | |
| | L Frontal Orbital Cortex | 3.74 | -26 | 34 | -14 |
| 24 | | | | | |
| | L Superior Temporal Gyrus, anterior division | 3.26 | -50 | -4 | -12 |
| 24 | | | | | |
| | R Parahippocampal Gyrus, posterior division | 3.59 | 18 | -34 | -10 |
| 24 | | | | | |
| | R Lateral Occipital Cortex, inferior division | 3.83 | 48 | -78 | -8 |
| 24 | | | | | |
| | L Insular Cortex | 3.30 | -36 | 0 | -6 |
| 24 | | | | | |
| | L Temporal Pole | 3.41 | -56 | 4 | -6 |
| 24 | | | | | |
| | R Lateral Occipital Cortex, inferior division | 3.27 | 38 | -90 | -4 |
| 24 | | | | | |
| | R Frontal Pole | 3.43 | 38 | 56 | -2 |
| 24 | | | | | |
| | L Lateral Occipital Cortex, inferior division | 3.21 | -46 | -86 | -2 |
| 24 | | | | | |
| | R Hippocampus | 3.89 | 26 | -36 | 0 |
| 24 | | | | | |
| | L Lateral Occipital Cortex, inferior division | 3.48 | -56 | -70 | 6 |
| 24 | | | | | |
| | L Thalamus | 3.26 | -14 | -22 | 8 |
| 24 | | | | | |
| | R Frontal Pole | 3.20 | 24 | 60 | 6 |
| 24 | | | | | |
| | R Frontal Pole | 3.42 | 34 | 64 | 10 |
| 24 | | | | | |
| | L Thalamus | 3.08 | -18 | -26 | 12 |
| 24 | | | | | |
| | R Putamen | 3.58 | 26 | 8 | 10 |
| 24 | | | | | |
| | R Planum Temporale | 3.41 | 46 | -28 | 14 |

| | | | | | |
|---|---|---|---|---|---|
| 24 | | | | | |
| | R Cingulate Gyrus, anterior division | 3.43 | 0 | 14 | 24 |
| 24 | | | | | |
| | R Inferior Frontal Gyrus, pars opercularis | 3.55 | 52 | 20 | 24 |
| 24 | | | | | |
| | L Lateral Occipital Cortex, superior division | 3.16 | -44 | -60 | 36 |
| 24 | | | | | |
| | R Cingulate Gyrus, posterior division | 3.08 | 0 | -28 | 34 |
| | L Cingulate Gyrus, posterior division | 3.16 | -2 | -30 | 36 |
| 24 | | | | | |
| | L Superior Frontal Gyrus | 3.44 | -6 | 36 | 42 |
| 24 | | | | | |
| | R Superior Frontal Gyrus | 3.24 | 4 | 48 | 42 |
| 24 | | | | | |
| | L Postcentral Gyrus | 3.38 | -54 | -20 | 50 |
| 24 | | | | | |
| | L Juxtapositional Lobule Cortex (formerly Supplementary Motor Cortex) | 3.62 | -6 | -14 | 52 |
| 16 | | | | | |
| | R Brain-Stem | 3.23 | 4 | -44 | -56 |
| 16 | | | | | |
| | L Brain-Stem | 3.69 | -4 | -38 | -46 |
| 16 | | | | | |
| | R Brain-Stem | 3.00 | 6 | -14 | -38 |
| 16 | | | | | |
| | R Brain-Stem | 3.04 | 6 | -14 | -28 |
| 16 | | | | | |
| | L Brain-Stem | 3.23 | -4 | -36 | -16 |
| 16 | | | | | |
| | L Lateral Occipital Cortex, inferior division | 3.22 | -42 | -78 | -14 |
| 16 | | | | | |
| | R Frontal Orbital Cortex | 3.20 | 38 | 26 | -14 |
| 16 | | | | | |
| | R Lateral Occipital Cortex, inferior division | 3.38 | 46 | -76 | -12 |
| 16 | | | | | |
| | L Hippocampus | 4.08 | -28 | -34 | -8 |
| 16 | | | | | |
| | L Frontal Orbital Cortex | 3.33 | -42 | 32 | -8 |
| 16 | | | | | |

| | | | | | |
|---|---|---|---|---|---|
| | R Middle Temporal Gyrus, posterior division | 3.34 | 54 | -24 | -6 |
| 16 | | | | | |
| | L Hippocampus | 3.23 | -24 | -40 | 0 |
| 16 | | | | | |
| | R Hippocampus | 3.42 | 18 | -38 | 2 |
| 16 | | | | | |
| | R Putamen | 3.18 | 30 | -14 | 0 |
| 16 | | | | | |
| | L Central Opercular Cortex | 3.39 | -54 | -4 | 8 |
| 16 | | | | | |
| | L Planum Temporale | 3.42 | -64 | -28 | 10 |
| 16 | | | | | |
| | R Planum Temporale | 3.02 | 56 | -28 | 12 |
| 16 | | | | | |
| | R Precentral Gyrus | 3.26 | 64 | -2 | 12 |
| 16 | | | | | |
| | L Frontal Pole | 3.04 | -44 | 52 | 14 |
| 16 | | | | | |
| | R Frontal Pole | 3.15 | 40 | 60 | 12 |
| 16 | | | | | |
| | R Postcentral Gyrus | 3.53 | 66 | -8 | 18 |
| 16 | | | | | |
| | L Lateral Occipital Cortex, superior division | 3.10 | -40 | -82 | 18 |
| 16 | | | | | |
| | L Angular Gyrus | 3.24 | -40 | -54 | 20 |
| 16 | | | | | |
| | R Inferior Frontal Gyrus, pars opercularis | 3.27 | 58 | 12 | 20 |
| 16 | | | | | |
| | L Middle Frontal Gyrus | 3.44 | -42 | 22 | 24 |
| 16 | | | | | |
| | R Cingulate Gyrus, anterior division | 3.70 | 4 | 24 | 26 |
| 16 | | | | | |
| | R Frontal Pole | 3.02 | 46 | 42 | 28 |
| 16 | | | | | |
| | R Paracingulate Gyrus | 3.28 | 6 | 32 | 30 |
| 16 | | | | | |
| | R Middle Frontal Gyrus | 3.09 | 46 | 22 | 32 |
| 16 | | | | | |
| | R Paracingulate Gyrus | 3.33 | 8 | 42 | 34 |
| 16 | | | | | |
| | L Paracingulate Gyrus | 3.15 | -2 | 22 | 36 |
| 16 | | | | | |
| | R Paracingulate Gyrus | 3.25 | 2 | 28 | 42 |

| | | | | | |
|---|---|---|---|---|---|
| 16 | | | | | |
| | L Middle Frontal Gyrus | 3.13 | -40 | 22 | 42 |
| 16 | | | | | |
| | L Precuneus Cortex | 3.09 | -12 | -42 | 44 |
| 8 | | | | | |
| | R Brain-Stem | 3.21 | 2 | -28 | -46 |
| 8 | | | | | |
| | R Brain-Stem | 3.23 | 6 | -42 | -44 |
| 8 | | | | | |
| | R Brain-Stem | 3.21 | 12 | -38 | -40 |
| 8 | | | | | |
| | R Brain-Stem | 3.09 | 0 | -20 | -38 |
| 8 | | | | | |
| | L Temporal Pole | 3.10 | -22 | 16 | -38 |
| 8 | | | | | |
| | R Temporal Pole | 3.12 | 48 | 6 | -34 |
| 8 | | | | | |
| | R Brain-Stem | 3.15 | 8 | -34 | -24 |
| 8 | | | | | |
| | L Temporal Pole | 3.08 | -52 | 8 | -22 |
| 8 | | | | | |
| | R Amygdala | 2.96 | 26 | -4 | -20 |
| | R Amygdala | 2.96 | 26 | -4 | -20 |
| 8 | | | | | |
| | R Middle Temporal Gyrus, anterior division | 3.07 | 58 | 2 | -20 |
| 8 | | | | | |
| | R Parahippocampal Gyrus, posterior division | 2.98 | 22 | -36 | -18 |
| 8 | | | | | |
| | R Planum Polare | 3.11 | 42 | 4 | -18 |
| 8 | | | | | |
| | R Inferior Temporal Gyrus, temporooccipital part | 3.25 | 64 | -40 | -16 |
| 8 | | | | | |
| | L Temporal Pole | 3.03 | -50 | 16 | -16 |
| 8 | | | | | |
| | R Frontal Orbital Cortex | 3.07 | 42 | 20 | -16 |
| 8 | | | | | |
| | L Frontal Pole | 2.99 | -24 | 42 | -14 |
| 8 | | | | | |
| | L Amygdala | 3.10 | -18 | -6 | -12 |
| | L Amygdala | 3.10 | -18 | -6 | -12 |
| 8 | | | | | |
| | R Temporal Pole | 3.03 | 56 | 10 | -10 |
| 8 | | | | | |
| | R Hippocampus | 3.02 | 26 | -32 | -6 |

| | | | | | |
|---|---|---|---|---|---|
| 8 | | | | | |
| | R Middle Temporal Gyrus, posterior division | 3.08 | 50 | -26 | -6 |
| 8 | | | | | |
| | L Superior Temporal Gyrus, anterior division | 3.09 | -56 | -6 | -6 |
| 8 | | | | | |
| | R Insular Cortex | 3.09 | 42 | 6 | -6 |
| 8 | | | | | |
| | R Frontal Orbital Cortex | 2.98 | 48 | 28 | -6 |
| 8 | | | | | |
| | L Paracingulate Gyrus | 3.38 | -6 | 46 | -4 |
| 8 | | | | | |
| | R Frontal Pole | 2.96 | 26 | 60 | -4 |
| 8 | | | | | |
| | L Superior Temporal Gyrus, posterior division | 3.39 | -62 | -22 | -2 |
| 8 | | | | | |
| | L Precentral Gyrus | 3.47 | -56 | 6 | 2 |
| 8 | | | | | |
| | L Thalamus | 2.97 | -20 | -26 | 6 |
| 8 | | | | | |
| | R Insular Cortex | 2.96 | 34 | -24 | 8 |
| 8 | | | | | |
| | R Central Opercular Cortex | 3.46 | 56 | -2 | 8 |
| 8 | | | | | |
| | R Frontal Pole | 3.37 | 10 | 64 | 8 |
| 8 | | | | | |
| | L Planum Temporale | 3.42 | -50 | -30 | 10 |
| 8 | | | | | |
| | R Precentral Gyrus | 3.16 | 60 | 2 | 10 |
| 8 | | | | | |
| | L Precentral Gyrus | 3.00 | -60 | 10 | 10 |
| 8 | | | | | |
| | L Central Opercular Cortex | 2.98 | -46 | -18 | 12 |
| 8 | | | | | |
| | R Central Opercular Cortex | 2.99 | 50 | -14 | 12 |
| 8 | | | | | |
| | R Caudate | 3.10 | 16 | 10 | 12 |
| 8 | | | | | |
| | L Inferior Frontal Gyrus, pars opercularis | 3.16 | -54 | 10 | 12 |
| 8 | | | | | |
| | R Frontal Pole | 3.02 | 6 | 66 | 12 |
| 8 | | | | | |
| | L Planum Temporale | 3.33 | -36 | -34 | 14 |
| 8 | | | | | |

| | | | | | |
|---|---|---|---|---|---|
| | L Inferior Frontal Gyrus, pars triangularis | 3.04 | -48 | 34 | 14 |
| 8 | | | | | |
| | L Parietal Operculum Cortex | 2.97 | -44 | -26 | 16 |
| 8 | | | | | |
| | L Frontal Pole | 3.20 | -48 | 42 | 16 |
| 8 | | | | | |
| | R Caudate | 3.26 | 16 | -4 | 18 |
| 8 | | | | | |
| | R Parietal Operculum Cortex | 2.97 | 40 | -30 | 20 |
| 8 | | | | | |
| | L Precentral Gyrus | 3.43 | -58 | 0 | 24 |
| 8 | | | | | |
| | L Cingulate Gyrus, anterior division | 3.13 | -2 | 10 | 26 |
| 8 | | | | | |
| | R Postcentral Gyrus | 3.00 | 64 | -18 | 30 |
| 8 | | | | | |
| | R Precentral Gyrus | 3.03 | 62 | 6 | 30 |
| 8 | | | | | |
| | R Superior Frontal Gyrus | 3.03 | 6 | 50 | 30 |
| 8 | | | | | |
| | L Frontal Pole | 2.97 | -14 | 52 | 30 |
| 8 | | | | | |
| | R Lateral Occipital Cortex, superior division | 3.14 | 24 | -82 | 34 |
| 8 | | | | | |
| | L Precuneus Cortex | 2.96 | -14 | -76 | 34 |
| 8 | | | | | |
| | L Cingulate Gyrus, anterior division | 2.96 | -8 | 12 | 34 |
| 8 | | | | | |
| | L Supramarginal Gyrus, anterior division | 3.04 | -60 | -28 | 36 |
| 8 | | | | | |
| | R Paracingulate Gyrus | 3.15 | 0 | 38 | 36 |
| 8 | | | | | |
| | L Middle Frontal Gyrus | 2.97 | -30 | 26 | 38 |
| 8 | | | | | |
| | L Supramarginal Gyrus, posterior division | 3.02 | -60 | -42 | 40 |
| 8 | | | | | |
| | L Supramarginal Gyrus, anterior division | 2.97 | -62 | -38 | 40 |
| 8 | | | | | |
| | L Precuneus Cortex | 2.99 | -6 | -50 | 42 |
| 8 | | | | | |

| | | | | | |
|---|---|---|---|---|---|
| | L Supramarginal Gyrus, posterior division | 3.40 | -50 | -48 | 42 |
| 8 | | | | | |
| | L Middle Frontal Gyrus | 3.22 | -40 | 20 | 50 |
| 8 | | | | | |
| | R Frontal Pole | 3.13 | 12 | 44 | 50 |
| 8 | | | | | |
| | R Frontal Pole | 2.98 | 12 | 40 | 52 |
| 8 | | | | | |
| | L Precuneus Cortex | 3.18 | -10 | -50 | 54 |
| 8 | | | | | |
| | R Postcentral Gyrus | 3.09 | 22 | -38 | 58 |
| 8 | | | | | |
| | L Precentral Gyrus | 3.00 | -36 | -12 | 60 |
| 8 | | | | | |
| | L Postcentral Gyrus | 3.19 | -20 | -46 | 66 |
| 8 | | | | | |
| | R Superior Parietal Lobule | 3.27 | 22 | -42 | 66 |
| 8 | | | | | |
| | L Precentral Gyrus | 3.37 | -6 | -16 | 72 |

**Supplementary Table S15. [$CT_{phasic}$ > $UT_{phasic}$].** Descriptive statistics for clusters and local maxima showing greater activation for Certain-Threat relative to Uncertain-Threat anticipation for the OSP Phasic regressor (FDR *q*<0.05, whole-brain corrected).

| *mm³* | Label | *t* | *x* | *y* | *z* |
|---|---|---|---|---|---|
| 73,416 | | | | | |
| | R Frontal Pole | 4.02 | 2 | 58 | -2 |
| | L Paracingulate Gyrus | 7.53 | -8 | 22 | 32 |
| | L Insular Cortex | 6.46 | -32 | 20 | 8 |
| | R Paracingulate Gyrus | 5.26 | 10 | 20 | 34 |
| | L Frontal Operculum Cortex | 6.72 | -34 | 16 | 8 |
| | L Cingulate Gyrus, anterior division | 7.97 | -8 | 14 | 36 |
| | L Inferior Frontal Gyrus, pars opercularis | 4.60 | -58 | 10 | 4 |
| | R Cingulate Gyrus, anterior division | 9.81 | 10 | 10 | 40 |
| | L Central Opercular Cortex | 3.07 | -52 | 2 | 4 |
| | R Juxtapositional Lobule Cortex (formerly Supplementary Motor Cortex) | 9.66 | 0 | 0 | |
| | L Middle Frontal Gyrus | 4.86 | -38 | -2 | 56 |
| | L Precentral Gyrus | 7.27 | -48 | -4 | 54 |
| | L Juxtapositional Lobule Cortex (formerly Supplementary Motor Cortex) | 9.77 | -2 | -6 | 64 |
| | L Superior Frontal Gyrus | 8.30 | -14 | -6 | 68 |
| | R Superior Frontal Gyrus | 7.18 | 16 | -10 | 72 |
| | R Precentral Gyrus | 8.80 | 32 | -26 | 52 |
| | R Postcentral Gyrus | 7.75 | 36 | -26 | 62 |
| | R Cingulate Gyrus, posterior division | 5.26 | 4 | -32 | 26 |
| | L Cingulate Gyrus, posterior division | 5.75 | -4 | -36 | 26 |
| | R Superior Parietal Lobule | 5.29 | 26 | -42 | 62 |
| | R Lateral Occipital Cortex, superior division | 4.34 | 32 | -62 | 64 |
| 27,752 | | | | | |
| | R Frontal Pole | 4.05 | 12 | 34 | -20 |
| | R Frontal Orbital Cortex | 5.09 | 22 | 28 | -20 |
| | L Accumbens | 5.22 | -8 | 18 | -2 |
| | R Subcallosal Cortex | 3.82 | 12 | 18 | -16 |
| | L Subcallosal Cortex | 3.63 | -6 | 16 | -14 |
| | L Putamen | 7.48 | -16 | 12 | -12 |
| | R Putamen | 8.32 | 20 | 12 | 0 |
| | L Frontal Orbital Cortex | 7.17 | -18 | 10 | -16 |
| | R Accumbens | 7.92 | 6 | 10 | -4 |
| | R Caudate | 10.69 | 8 | 6 | 2 |
| | L Caudate | 9.46 | -8 | 4 | 4 |
| | R Amygdala | 4.47 | 16 | 0 | -20 |
| | R Thalamus | 6.30 | 8 | -2 | 6 |
| | R Amygdala | 4.17 | 12 | -4 | -18 |
| | R Amygdala | 2.81 | 18 | -4 | -24 |
| | R Amygdala | 4.50 | 28 | -8 | -12 |
| | R Amygdala | 4.20 | 30 | -8 | -18 |

| | | | | | |
|---|---|---|---|---|---|
| | L Amygdala | 5.54 | -28 | -10 | -16 |
| | L Hippocampus | 5.54 | -22 | -14 | -18 |
| | L Amygdala | 5.46 | -20 | -16 | -14 |
| | L Thalamus | 5.44 | -2 | -16 | 12 |
| | R Hippocampus | 4.58 | 22 | -20 | -16 |
| 26,104 | | | | | |
| | R Middle Temporal Gyrus, temporooccipital part | 3.95 | 54 | -52 | -6 |
| | R Temporal Occipital Fusiform Cortex | 2.91 | 40 | -52 | -16 |
| | R Lingual Gyrus | 2.75 | 12 | -70 | -12 |
| | R Occipital Fusiform Gyrus | 3.19 | 20 | -84 | -6 |
| | R Lateral Occipital Cortex, inferior division | 9.84 | 32 | -88 | -2 |
| | R Occipital Pole | 13.54 | 24 | -98 | 0 |
| 7,992 | | | | | |
| | L Postcentral Gyrus | 3.68 | -48 | -30 | 46 |
| | L Supramarginal Gyrus, anterior division | 5.73 | -42 | -38 | 40 |
| | L Supramarginal Gyrus, posterior division | 4.64 | -58 | -46 | 50 |
| | L Superior Parietal Lobule | 5.44 | -34 | -54 | 50 |
| | L Angular Gyrus | 3.09 | -42 | -58 | 46 |
| | L Lateral Occipital Cortex, superior division | 4.00 | -30 | -62 | 40 |
| 6,144 | | | | | |
| | L Frontal Pole | 6.33 | -36 | 48 | 30 |
| | L Middle Frontal Gyrus | 5.58 | -38 | 36 | 40 |
| 5,992 | | | | | |
| | L Lateral Occipital Cortex, inferior division | 4.64 | -40 | -72 | 2 |
| | L Occipital Pole | 7.77 | -26 | -96 | -6 |
| 3,856 | | | | | |
| | L Frontal Pole | 4.61 | -8 | 68 | -16 |
| | R Frontal Pole | 4.83 | 2 | 66 | -10 |
| 3,448 | | | | | |
| | R Precuneus Cortex | 3.90 | 4 | -60 | 44 |
| | L Precuneus Cortex | 6.75 | -10 | -70 | 40 |
| 2,664 | | | | | |
| | R Precuneus Cortex | 6.46 | 14 | -64 | 34 |
| | R Lateral Occipital Cortex, superior division | 3.55 | 12 | -74 | 52 |
| 2,248 | | | | | |
| | R Frontal Orbital Cortex | 4.93 | 34 | 28 | 2 |
| | R Frontal Operculum Cortex | 6.10 | 36 | 20 | 8 |
| | R Central Opercular Cortex | 3.11 | 48 | 4 | 2 |
| 1,880 | | | | | |
| | R Frontal Pole | 5.08 | 30 | 38 | 36 |
| 1,384 | | | | | |
| | R Temporal Pole | 4.27 | 42 | 6 | -40 |
| | R Inferior Temporal Gyrus, anterior division | 4.14 | 52 | -4 | -34 |

| | | | | | |
|---|---|---|---|---|---|
| 1,136 | | | | | |
| | L Frontal Pole | 4.30 | -26 | 40 | -18 |
| | L Frontal Orbital Cortex | 6.91 | -24 | 28 | -22 |
| 888 | | | | | |
| | L Temporal Pole | 4.01 | -46 | 6 | -40 |
| | L Inferior Temporal Gyrus, anterior division | 5.33 | -46 | 0 | -38 |
| | L Temporal Fusiform Cortex, anterior division | 3.29 | -38 | -2 | -40 |
| 824 | | | | | |
| | R Angular Gyrus | 4.13 | 56 | -46 | 26 |
| 592 | | | | | |
| | R Precentral Gyrus | 4.24 | 60 | 4 | 14 |
| 576 | | | | | |
| | R Brain-Stem | 3.27 | 8 | -48 | -54 |
| 392 | | | | | |
| | L Brain-Stem | 4.11 | -8 | -46 | -52 |
| 344 | | | | | |
| | R Precuneus Cortex | 4.60 | 2 | -44 | 42 |
| | R Cingulate Gyrus, posterior division | 3.13 | 4 | -44 | 36 |
| 304 | | | | | |
| | R Temporal Pole | 4.80 | 38 | 18 | -32 |
| 296 | | | | | |
| | L Brain-Stem | 4.45 | -10 | -24 | -42 |
| 288 | | | | | |
| | L Cingulate Gyrus, posterior division | 4.57 | -12 | -28 | 40 |
| 280 | | | | | |
| | R Central Opercular Cortex | 3.49 | 50 | -18 | 18 |
| | R Parietal Operculum Cortex | 4.15 | 46 | -22 | 18 |
| 272 | | | | | |
| | L Hippocampus | 4.27 | -22 | -38 | 0 |
| 240 | | | | | |
| | L Lateral Occipital Cortex, superior division | 3.80 | -28 | -80 | 50 |
| 216 | | | | | |
| | R Insular Cortex | 4.14 | 36 | 16 | -10 |
| 192 | | | | | |
| | R Lateral Occipital Cortex, superior division | 3.37 | 26 | -64 | 42 |
| 192 | | | | | |
| | L Precentral Gyrus | 3.44 | -16 | -30 | 68 |
| 168 | | | | | |
| | L Postcentral Gyrus | 3.77 | -62 | -22 | 28 |
| 152 | | | | | |
| | R Brain-Stem | 3.26 | 0 | -36 | -40 |
| 144 | | | | | |

| | | | | | |
|---|---|---|---|---|---|
| | R Temporal Fusiform Cortex, anterior division | 3.28 | 32 | -8 | -42 |
| | R Temporal Fusiform Cortex, posterior division | 3.04 | 32 | -10 | -38 |
| 144 | | | | | |
| | R Brain-Stem | 3.03 | 14 | -20 | -38 |
| 144 | | | | | |
| | R Brain-Stem | 3.74 | 8 | -24 | -22 |
| 136 | | | | | |
| | L Hippocampus | 3.81 | -32 | -18 | -14 |
| 128 | | | | | |
| | R Temporal Pole | 3.03 | 26 | 6 | -34 |
| | R Parahippocampal Gyrus, anterior division | 3.37 | 26 | 2 | -38 |
| 128 | | | | | |
| | R Lateral Occipital Cortex, superior division | 3.20 | 30 | -70 | 26 |
| 120 | | | | | |
| | L Temporal Pole | 3.45 | -32 | 4 | -22 |
| | L Amygdala | 4.22 | -32 | -2 | -20 |
| 120 | | | | | |
| | R Supramarginal Gyrus, posterior division | 2.77 | 66 | -40 | 34 |
| 112 | | | | | |
| | L Frontal Pole | 3.37 | -16 | 50 | -20 |
| 104 | | | | | |
| | R Hippocampus | 4.24 | 26 | -36 | 4 |
| 96 | | | | | |
| | L Bed Nucleus of the Stria Terminalis | 9.84 | -6 | 4 | 0 |
| | R Brain-Stem | 3.18 | 4 | -32 | -48 |
| 96 | | | | | |
| | L Brain-Stem | 2.87 | -2 | -32 | -28 |
| | R Brain-Stem | 3.29 | 0 | -38 | -28 |
| 96 | | | | | |
| | L Lateral Occipital Cortex, superior division | 3.00 | -26 | -70 | 60 |
| 88 | | | | | |
| | R Temporal Fusiform Cortex, posterior division | 3.48 | 34 | -14 | -34 |
| 88 | | | | | |
| | L Brain-Stem | 3.51 | -4 | -38 | -22 |
| 88 | | | | | |
| | R Middle Temporal Gyrus, posterior division | 3.77 | 64 | -20 | -14 |
| 88 | | | | | |
| | L Insular Cortex | 3.21 | -32 | 18 | -8 |
| 88 | | | | | |
| | L Frontal Pole | 2.80 | -10 | 58 | 4 |

| | | | | | |
|---|---|---|---|---|---|
| | L Paracingulate Gyrus | 3.30 | -8 | 54 | 2 |
| 88 | | | | | |
| | L Supramarginal Gyrus, posterior division | 3.79 | -50 | -50 | 38 |
| | L Angular Gyrus | 2.69 | -48 | -52 | 34 |
| 80 | | | | | |
| | R Brain-Stem | 3.85 | 8 | -24 | -34 |
| 80 | | | | | |
| | L Frontal Pole | 3.38 | -26 | 64 | -12 |
| 80 | | | | | |
| | R Paracingulate Gyrus | 3.14 | 2 | 36 | -10 |
| | R Frontal Medial Cortex | 3.08 | 8 | 36 | -12 |
| 72 | | | | | |
| | L Inferior Temporal Gyrus, anterior division | 2.97 | -52 | -8 | -34 |
| 72 | | | | | |
| | L Middle Temporal Gyrus, posterior division | 2.69 | -66 | -22 | -20 |
| 72 | | | | | |
| | L Paracingulate Gyrus | 3.25 | -6 | 46 | 6 |
| 72 | | | | | |
| | L Frontal Pole | 3.33 | -32 | 62 | 16 |
| 72 | | | | | |
| | L Lateral Occipital Cortex, superior division | 3.18 | -36 | -66 | 56 |
| 64 | | | | | |
| | R Temporal Pole | 2.88 | 36 | 14 | -22 |
| 64 | | | | | |
| | L Middle Temporal Gyrus, posterior division | 3.36 | -66 | -32 | -20 |
| 64 | | | | | |
| | L Hippocampus | 3.31 | -24 | -26 | -12 |
| 64 | | | | | |
| | R Lateral Occipital Cortex, superior division | 3.51 | 46 | -74 | 42 |
| 56 | | | | | |
| | R Parahippocampal Gyrus, anterior division | 3.46 | 22 | -10 | -32 |
| 56 | | | | | |
| | L Cingulate Gyrus, anterior division | 3.58 | -4 | 40 | -4 |
| 56 | | | | | |
| | L Pallidum | 3.28 | -20 | -10 | 0 |
| 56 | | | | | |
| | R Angular Gyrus | 2.95 | 62 | -50 | 42 |
| 56 | | | | | |
| | L Precentral Gyrus | 3.51 | -10 | -18 | 42 |
| 48 | | | | | |
| | L Brain-Stem | 3.01 | -6 | -20 | -28 |

| | | | | | |
|---|---|---|---|---|---|
| 48 | | | | | |
| | L Frontal Orbital Cortex | 3.00 | -22 | 18 | -26 |
| 48 | | | | | |
| | R Brain-Stem | 3.16 | 2 | -18 | -20 |
| | L Brain-Stem | 2.84 | -4 | -22 | -18 |
| 48 | | | | | |
| | L Middle Temporal Gyrus, posterior division | 3.43 | -70 | -20 | -14 |
| 48 | | | | | |
| | L Lateral Occipital Cortex, inferior division | 3.15 | -46 | -70 | -8 |
| 48 | | | | | |
| | R Middle Temporal Gyrus, posterior division | 3.32 | 62 | -32 | -6 |
| 48 | | | | | |
| | L Lateral Occipital Cortex, superior division | 3.54 | -36 | -82 | 44 |
| 40 | | | | | |
| | L Parahippocampal Gyrus, anterior division | 3.05 | -26 | 0 | -32 |
| 40 | | | | | |
| | R Hippocampus | 3.50 | 32 | -8 | -26 |
| 40 | | | | | |
| | L Inferior Temporal Gyrus, temporooccipital part | 3.01 | -46 | -58 | -6 |
| 40 | | | | | |
| | R Precuneus Cortex | 2.73 | 6 | -56 | 24 |
| 40 | | | | | |
| | R Frontal Pole | 3.37 | 38 | 52 | 26 |
| 40 | | | | | |
| | R Lateral Occipital Cortex, superior division | 2.89 | 52 | -68 | 40 |
| 40 | | | | | |
| | R Angular Gyrus | 2.84 | 50 | -52 | 40 |
| 40 | | | | | |
| | L Precuneus Cortex | 3.09 | -8 | -54 | 58 |
| 32 | | | | | |
| | R Brain-Stem | 3.17 | 8 | -40 | -52 |
| 32 | | | | | |
| | L Temporal Pole | 3.42 | -38 | 12 | -36 |
| 32 | | | | | |
| | L Brain-Stem | 2.56 | -18 | -24 | -34 |
| 32 | | | | | |
| | L Frontal Orbital Cortex | 3.30 | -14 | 32 | -22 |
| 32 | | | | | |
| | L Frontal Pole | 3.00 | -14 | 38 | -24 |
| 32 | | | | | |

| | | | | | |
|---|---|---|---|---|---|
| | L Middle Temporal Gyrus, posterior division | 3.19 | -60 | -26 | -12 |
| 32 | | | | | |
| | R Middle Temporal Gyrus, posterior division | 2.97 | 66 | -16 | -8 |
| 32 | | | | | |
| | L Thalamus | 2.77 | -12 | -32 | 8 |
| 32 | | | | | |
| | R Paracingulate Gyrus | 2.71 | 14 | 48 | 8 |
| 32 | | | | | |
| | R Caudate | 3.18 | 16 | -14 | 22 |
| 32 | | | | | |
| | R Frontal Pole | 2.67 | 28 | 42 | 22 |
| 32 | | | | | |
| | L Middle Frontal Gyrus | 2.69 | -44 | 20 | 36 |
| 32 | | | | | |
| | R Precuneus Cortex | 2.65 | 8 | -56 | 52 |
| 32 | | | | | |
| | R Middle Frontal Gyrus | 2.69 | 32 | 2 | 54 |
| 32 | | | | | |
| | L Lateral Occipital Cortex, superior division | 2.84 | -18 | -68 | 66 |
| 24 | | | | | |
| | L Temporal Pole | 2.64 | -40 | 8 | -44 |
| 24 | | | | | |
| | L Temporal Pole | 2.59 | -34 | 10 | -30 |
| 24 | | | | | |
| | R Inferior Temporal Gyrus, posterior division | 2.97 | 56 | -18 | -26 |
| 24 | | | | | |
| | R Temporal Fusiform Cortex, posterior division | 2.76 | 38 | -18 | -26 |
| 24 | | | | | |
| | R Subcallosal Cortex | 3.23 | 2 | 20 | -18 |
| 24 | | | | | |
| | R Middle Temporal Gyrus, posterior division | 2.74 | 62 | -34 | -14 |
| 24 | | | | | |
| | R Temporal Pole | 2.79 | 52 | 10 | -14 |
| 24 | | | | | |
| | L Frontal Pole | 2.79 | -22 | 52 | -14 |
| 24 | | | | | |
| | R Hippocampus | 3.22 | 36 | -30 | -12 |
| 24 | | | | | |
| | R Thalamus | 2.97 | 14 | -36 | 4 |
| 24 | | | | | |
| | R Thalamus | 2.58 | 6 | -16 | 6 |

| | | | | | |
|---|---|---|---|---|---|
| 24 | | | | | |
| | R Paracingulate Gyrus | 2.72 | 4 | 50 | 12 |
| 24 | | | | | |
| | L Parietal Operculum Cortex | 3.00 | -42 | -30 | 22 |
| 24 | | | | | |
| | L Parietal Operculum Cortex | 2.61 | -44 | -34 | 24 |
| 24 | | | | | |
| | R Precentral Gyrus | 2.77 | 46 | 6 | 30 |
| 24 | | | | | |
| | R Superior Frontal Gyrus | 2.98 | 6 | 42 | 38 |
| 24 | | | | | |
| | R Cingulate Gyrus, posterior division | 2.82 | 10 | -38 | 42 |
| 24 | | | | | |
| | L Lateral Occipital Cortex, superior division | 2.63 | -12 | -76 | 58 |
| 24 | | | | | |
| | R Lateral Occipital Cortex, superior division | 2.71 | 16 | -66 | 64 |
| 24 | | | | | |
| | L Lateral Occipital Cortex, superior division | 2.75 | -14 | -64 | 64 |
| 16 | | | | | |
| | R Brain-Stem | 2.66 | 12 | -24 | -42 |
| 16 | | | | | |
| | R Inferior Temporal Gyrus, posterior division | 2.68 | 60 | -16 | -38 |
| 16 | | | | | |
| | L Inferior Temporal Gyrus, posterior division | 2.61 | -48 | -16 | -38 |
| 16 | | | | | |
| | L Temporal Fusiform Cortex, posterior division | 2.81 | -34 | -18 | -36 |
| 16 | | | | | |
| | L Temporal Pole | 2.88 | -36 | 6 | -36 |
| 16 | | | | | |
| | L Temporal Pole | 2.72 | -52 | 4 | -34 |
| 16 | | | | | |
| | L Temporal Pole | 2.70 | -50 | 14 | -34 |
| 16 | | | | | |
| | L Temporal Fusiform Cortex, posterior division | 2.66 | -36 | -34 | -28 |
| 16 | | | | | |
| | L Inferior Temporal Gyrus, posterior division | 2.79 | -50 | -34 | -28 |
| 16 | | | | | |
| | L Amygdala | 3.25 | -14 | -2 | -20 |
| 16 | | | | | |

| | | | | | |
|---|---|---|---|---|---|
| | R Frontal Pole | 2.76 | 0 | 54 | -20 |
| 16 | | | | | |
| | L Frontal Pole | 2.77 | -32 | 58 | -14 |
| 16 | | | | | |
| | R Frontal Medial Cortex | 2.87 | 2 | 50 | -12 |
| 16 | | | | | |
| | L Parahippocampal Gyrus, posterior division | 3.33 | -14 | -32 | -10 |
| 16 | | | | | |
| | L Hippocampus | 3.25 | -28 | -38 | -8 |
| 16 | | | | | |
| | R Insular Cortex | 2.87 | 36 | 4 | -8 |
| 16 | | | | | |
| | L Frontal Orbital Cortex | 3.25 | -28 | 28 | -4 |
| 16 | | | | | |
| | R Hippocampus | 3.19 | 26 | -38 | -2 |
| 16 | | | | | |
| | R Thalamus | 2.77 | 4 | -22 | -2 |
| 16 | | | | | |
| | R Thalamus | 2.60 | 6 | -30 | 4 |
| 16 | | | | | |
| | L Lateral Occipital Cortex, inferior division | 2.77 | -42 | -66 | 6 |
| 16 | | | | | |
| | L Insular Cortex | 2.68 | -36 | -16 | 10 |
| 16 | | | | | |
| | R Thalamus | 3.17 | 20 | -20 | 14 |
| 16 | | | | | |
| | R Cingulate Gyrus, anterior division | 2.75 | 10 | 38 | 16 |
| 16 | | | | | |
| | R Angular Gyrus | 2.97 | 44 | -48 | 20 |
| 16 | | | | | |
| | L Frontal Pole | 2.66 | -20 | 56 | 18 |
| 16 | | | | | |
| | L Caudate | 3.38 | -14 | -10 | 20 |
| 16 | | | | | |
| | L Caudate | 2.71 | -14 | -6 | 20 |
| 16 | | | | | |
| | L Frontal Pole | 2.86 | -26 | 46 | 20 |
| 16 | | | | | |
| | R Precentral Gyrus | 2.93 | 52 | 4 | 22 |
| 16 | | | | | |
| | R Precentral Gyrus | 3.17 | 58 | 6 | 26 |
| 16 | | | | | |
| | L Supramarginal Gyrus, posterior division | 2.87 | -54 | -44 | 28 |
| 16 | | | | | |
| | R Precentral Gyrus | 2.70 | 42 | 2 | 30 |
| 16 | | | | | |

| | | | | | |
|---|---|---|---|---|---|
| | R Supramarginal Gyrus, posterior division | 2.73 | 58 | -40 | 38 |
| 16 | | | | | |
| | R Angular Gyrus | 2.56 | 44 | -48 | 42 |
| 16 | | | | | |
| | R Supramarginal Gyrus, anterior division | 3.10 | 54 | -28 | 40 |
| 16 | | | | | |
| | R Postcentral Gyrus | 2.87 | 54 | -10 | 50 |
| 16 | | | | | |
| | L Superior Parietal Lobule | 2.87 | -36 | -42 | 60 |
| 16 | | | | | |
| | R Lateral Occipital Cortex, superior division | 2.66 | 16 | -74 | 60 |
| 16 | | | | | |
| | L Lateral Occipital Cortex, superior division | 2.83 | -10 | -66 | 64 |
| 8 | | | | | |
| | L Brain-Stem | 2.50 | -2 | -36 | -48 |
| 8 | | | | | |
| | R Inferior Temporal Gyrus, anterior division | 2.48 | 50 | -2 | -46 |
| 8 | | | | | |
| | R Brain-Stem | 2.71 | 14 | -26 | -40 |
| 8 | | | | | |
| | L Temporal Pole | 2.64 | -46 | 12 | -40 |
| 8 | | | | | |
| | R Temporal Pole | 2.58 | 32 | 4 | -38 |
| 8 | | | | | |
| | L Brain-Stem | 2.55 | -12 | -30 | -30 |
| 8 | | | | | |
| | R Inferior Temporal Gyrus, posterior division | 2.58 | 54 | -12 | -30 |
| 8 | | | | | |
| | R Temporal Pole | 2.80 | 28 | 10 | -30 |
| 8 | | | | | |
| | L Brain-Stem | 2.88 | -16 | -22 | -28 |
| 8 | | | | | |
| | L Inferior Temporal Gyrus, posterior division | 2.59 | -52 | -36 | -26 |
| 8 | | | | | |
| | R Temporal Pole | 2.76 | 32 | 8 | -26 |
| 8 | | | | | |
| | R Brain-Stem | 2.61 | 12 | -30 | -22 |
| 8 | | | | | |
| | L Temporal Fusiform Cortex, posterior division | 3.24 | -38 | -24 | -20 |
| 8 | | | | | |
| | L Frontal Orbital Cortex | 2.58 | -14 | 20 | -20 |

| | | | | | |
|---|---|---|---|---|---|
| 8 | | | | | |
| | R Middle Temporal Gyrus, posterior division | 2.49 | 64 | -8 | -18 |
| 8 | | | | | |
| | R Frontal Orbital Cortex | 2.56 | 18 | 22 | -18 |
| 8 | | | | | |
| | R Occipital Fusiform Gyrus | 2.49 | 30 | -80 | -16 |
| 8 | | | | | |
| | R Temporal Occipital Fusiform Cortex | 2.49 | 26 | -58 | -16 |
| 8 | | | | | |
| | R Frontal Medial Cortex | 2.85 | 4 | 34 | -16 |
| 8 | | | | | |
| | L Frontal Pole | 2.57 | -30 | 50 | -14 |
| 8 | | | | | |
| | L Hippocampus | 2.61 | -34 | -32 | -12 |
| 8 | | | | | |
| | L Middle Temporal Gyrus, posterior division | 2.51 | -56 | -30 | -12 |
| 8 | | | | | |
| | L Subcallosal Cortex | 2.50 | -4 | 24 | -12 |
| 8 | | | | | |
| | R Subcallosal Cortex | 2.77 | 4 | 28 | -12 |
| 8 | | | | | |
| | R Frontal Pole | 2.60 | 24 | 66 | -12 |
| 8 | | | | | |
| | L Hippocampus | 2.48 | -32 | -28 | -10 |
| 8 | | | | | |
| | R Middle Temporal Gyrus, posterior division | 2.58 | 70 | -22 | -10 |
| 8 | | | | | |
| | L Subcallosal Cortex | 2.89 | -6 | 28 | -6 |
| 8 | | | | | |
| | R Middle Temporal Gyrus, posterior division | 2.55 | 56 | -36 | -4 |
| 8 | | | | | |
| | R Inferior Frontal Gyrus, pars triangularis | 2.69 | 50 | 20 | -4 |
| 8 | | | | | |
| | R Middle Temporal Gyrus, temporooccipital part | 2.55 | 60 | -46 | -2 |
| 8 | | | | | |
| | R Middle Temporal Gyrus, temporooccipital part | 2.53 | 54 | -48 | 4 |
| 8 | | | | | |
| | R Frontal Pole | 2.58 | 0 | 64 | 4 |
| 8 | | | | | |
| | L Frontal Pole | 2.56 | -28 | 66 | 6 |
| 8 | | | | | |

| | | | | | |
|---|---|---|---|---|---|
| | L Cingulate Gyrus, posterior division | 2.54 | -8 | -44 | 8 |
| 8 | | | | | |
| | L Central Opercular Cortex | 2.83 | -52 | 0 | 8 |
| 8 | | | | | |
| | R Cingulate Gyrus, posterior division | 2.56 | 8 | -44 | 10 |
| 8 | | | | | |
| | R Middle Temporal Gyrus, temporooccipital part | 2.87 | 48 | -48 | 14 |
| 8 | | | | | |
| | R Central Opercular Cortex | 2.50 | 46 | -4 | 14 |
| 8 | | | | | |
| | L Central Opercular Cortex | 2.67 | -42 | -4 | 14 |
| 8 | | | | | |
| | R Paracingulate Gyrus | 2.56 | 6 | 48 | 14 |
| 8 | | | | | |
| | R Lateral Occipital Cortex, superior division | 2.67 | 44 | -60 | 20 |
| 8 | | | | | |
| | R Paracingulate Gyrus | 2.62 | 10 | 40 | 20 |
| 8 | | | | | |
| | L Precuneus Cortex | 2.67 | -8 | -54 | 22 |
| 8 | | | | | |
| | R Inferior Frontal Gyrus, pars opercularis | 2.52 | 38 | 14 | 24 |
| 8 | | | | | |
| | L Supramarginal Gyrus, anterior division | 2.56 | -62 | -34 | 30 |
| 8 | | | | | |
| | R Precuneus Cortex | 2.67 | 12 | -54 | 36 |
| 8 | | | | | |
| | L Precuneus Cortex | 2.51 | -8 | -56 | 38 |
| 8 | | | | | |
| | R Angular Gyrus | 2.64 | 58 | -48 | 38 |
| 8 | | | | | |
| | L Precuneus Cortex | 3.37 | -10 | -60 | 40 |
| 8 | | | | | |
| | R Precuneus Cortex | 2.58 | 4 | -52 | 44 |
| 8 | | | | | |
| | R Precuneus Cortex | 2.49 | 8 | -40 | 46 |
| 8 | | | | | |
| | R Superior Frontal Gyrus | 2.72 | 6 | 36 | 50 |
| 8 | | | | | |
| | L Middle Frontal Gyrus | 2.69 | -30 | 6 | 56 |
| 8 | | | | | |
| | R Superior Frontal Gyrus | 2.68 | 22 | 10 | 58 |
| 8 | | | | | |
| | L Precentral Gyrus | 2.58 | -20 | -26 | 74 |

**Supplementary Table S16. [$UT_{sustained}$ > $US_{sustained}$] ∩ [$CT_{phasic}$ > $CS_{phasic}$].** Descriptive statistics for clusters and local maxima showing greater activation for Uncertain-Threat relative to Uncertain-Safety anticipation for the OSP Sustained regressor AND greater activation for Certain-Threat relative to Certain-Safety anticipation for the OSP Phasic regressor (FDR $q$<0.05, whole-brain corrected).

| *mm³* | Label | *t* | *x* | *y* | *z* |
|---|---|---|---|---|---|
| 18,304 | | | | | |
| | L Left Caudate | 4.00 | -8 | 4 | 4 |
| | R Right Thalamus | 4.22 | 8 | -2 | 6 |
| | R Right Caudate | 5.32 | 12 | -4 | 16 |
| | R Left Thalamus | 4.05 | 0 | -6 | 8 |
| | L Left Thalamus | 3.79 | -2 | -14 | -2 |
| | R Brain-Stem | 4.05 | 10 | -28 | -26 |
| | L Brain-Stem | 6.24 | -2 | -30 | -2 |
| | L Temporal Occipital Fusiform Cortex | 3.75 | -34 | -54 | -22 |
| | L Occipital Fusiform Gyrus | 3.95 | -24 | -68 | -16 |
| | L Lingual Gyrus | 3.62 | -10 | -76 | -14 |
| 15,736 | | | | | |
| | L Paracingulate Gyrus | 3.99 | -8 | 30 | 26 |
| | L Cingulate Gyrus, anterior division | 5.79 | -8 | 14 | 36 |
| | R Superior Frontal Gyrus | 3.60 | 4 | 14 | 60 |
| | R Paracingulate Gyrus | 6.22 | 0 | 12 | 46 |
| | R Cingulate Gyrus, anterior division | 7.24 | 10 | 8 | 40 |
| | L Juxtapositional Lobule Cortex (formerly Supplementary Motor Cortex) | 5.74 | -6 | 4 | 46 |
| | L Bed Nucleus of the Stria Terminalis | 4.24 | -6 | 4 | -2 |
| | R Juxtapositional Lobule Cortex (formerly Supplementary Motor Cortex) | 6.89 | 4 | -2 | 50 |
| | L Superior Frontal Gyrus | 5.59 | -12 | -8 | 66 |
| 13,920 | | | | | |
| | R Insular Cortex | 6.93 | 34 | 24 | 2 |
| | R Frontal Orbital Cortex | 5.41 | 36 | 20 | -10 |
| | R Frontal Operculum Cortex | 8.51 | 36 | 18 | 8 |
| | R Inferior Frontal Gyrus, pars opercularis | 4.95 | 60 | 14 | 8 |
| | R Right Putamen | 8.65 | 22 | 10 | -4 |
| | R Central Opercular Cortex | 7.32 | 48 | 8 | -2 |
| | R Precentral Gyrus | 4.58 | 60 | 4 | 14 |
| | R Right Pallidum | 3.40 | 16 | 2 | -4 |
| 9,016 | | | | | |
| | L Frontal Orbital Cortex | 6.44 | -30 | 22 | -8 |
| | L Inferior Frontal Gyrus, pars triangularis | 3.92 | -50 | 22 | 12 |
| | L Frontal Operculum Cortex | 8.12 | -42 | 16 | -2 |
| | L Insular Cortex | 7.89 | -36 | 14 | 4 |
| | L Inferior Frontal Gyrus, pars opercularis | 6.44 | -58 | 12 | 4 |
| | L Precentral Gyrus | 4.64 | -56 | 6 | 2 |
| | L Central Opercular Cortex | 5.40 | -50 | 2 | 2 |

| | | | | | |
|---|---|---|---|---|---|
| 7,440 | | | | | |
| | R Supramarginal Gyrus, anterior division | 3.55 | 64 | -28 | 30 |
| | R Parietal Operculum Cortex | 5.12 | 54 | -30 | 26 |
| | R Angular Gyrus | 5.48 | 56 | -46 | 32 |
| | R Supramarginal Gyrus, posterior division | 5.39 | 58 | -46 | 38 |
| 4,712 | | | | | |
| | L Postcentral Gyrus | 2.94 | -58 | -22 | 22 |
| | L Parietal Operculum Cortex | 4.14 | -54 | -34 | 24 |
| | L Supramarginal Gyrus, anterior division | 5.09 | -66 | -38 | 26 |
| | L Supramarginal Gyrus, posterior division | 4.89 | -64 | -48 | 34 |
| | L Angular Gyrus | 3.35 | -58 | -54 | 34 |
| 4,088 | | | | | |
| | R Frontal Pole | 5.87 | 26 | 46 | 26 |
| 3,608 | | | | | |
| | L Frontal Pole | 6.22 | -36 | 48 | 30 |
| | L Middle Frontal Gyrus | 4.12 | -32 | 30 | 40 |
| 3,040 | | | | | |
| | L Putamen | 6.01 | -20 | 8 | 0 |
| 2,664 | | | | | |
| | R Middle Frontal Gyrus | 6.09 | 46 | 2 | 56 |
| | R Precentral Gyrus | 4.46 | 40 | -14 | 54 |
| 2,640 | | | | | |
| | R Temporal Occipital Fusiform Cortex | 3.48 | 36 | -60 | -20 |
| 1,448 | | | | | |
| | R Superior Frontal Gyrus | 4.66 | 14 | -10 | 68 |
| 1,416 | | | | | |
| | R Middle Temporal Gyrus, posterior division | 4.72 | 56 | -28 | -4 |
| | R Middle Temporal Gyrus, temporooccipital part | 2.70 | 64 | -38 | 0 |
| 1,040 | | | | | |
| | R Brain-Stem | 4.69 | 2 | -34 | -50 |
| | L Brain-Stem | 3.78 | -8 | -38 | -48 |
| 840 | | | | | |
| | L Precentral Gyrus | 4.26 | -40 | -2 | 52 |
| 640 | | | | | |
| | R Occipital Pole | 3.89 | 24 | -94 | 16 |
| 408 | | | | | |
| | R Cingulate Gyrus, posterior division | 4.18 | 10 | -24 | 42 |
| 296 | | | | | |
| | R Temporal Pole | 3.63 | 48 | 10 | -40 |
| 280 | | | | | |
| | R Precentral Gyrus | 3.38 | 34 | -22 | 52 |
| 176 | | | | | |

| | | | | | |
|---|---|---|---|---|---|
| | R Precentral Gyrus | 4.40 | 32 | -14 | 64 |
| 136 | | | | | |
| | L Cingulate Gyrus, posterior division | 3.58 | -10 | -24 | 40 |
| 136 | | | | | |
| | L Supramarginal Gyrus, posterior division | 3.34 | -56 | -48 | 50 |
| 128 | | | | | |
| | R Temporal Occipital Fusiform Cortex | 3.32 | 28 | -56 | -18 |
| 128 | | | | | |
| | R Superior Parietal Lobule | 3.46 | 24 | -42 | 60 |
| 112 | | | | | |
| | L Precentral Gyrus | 3.34 | -26 | -10 | 58 |
| 104 | | | | | |
| | L Frontal Pole | 3.26 | -26 | 48 | 16 |
| 72 | | | | | |
| | L Caudate | 3.79 | -16 | 18 | 4 |
| 64 | | | | | |
| | R Inferior Temporal Gyrus, anterior division | 3.54 | 48 | 0 | -34 |
| 56 | | | | | |
| | L Thalamus | 3.40 | -10 | -10 | 6 |
| 56 | | | | | |
| | R Thalamus | 3.85 | 4 | -14 | 16 |
| 48 | | | | | |
| | R Caudate | 3.02 | 16 | -14 | 24 |
| 48 | | | | | |
| | L Superior Frontal Gyrus | 3.14 | -14 | 2 | 68 |
| 40 | | | | | |
| | L Caudate | 3.16 | -14 | -4 | 20 |
| 40 | | | | | |
| | L Precentral Gyrus | 2.82 | -50 | -2 | 42 |
| 32 | | | | | |
| | R Frontal Pole | 2.97 | 28 | 62 | -12 |
| 32 | | | | | |
| | R Supramarginal Gyrus, posterior division | 2.87 | 66 | -42 | 10 |
| 32 | | | | | |
| | L Caudate | 2.81 | -14 | 8 | 18 |
| 32 | | | | | |
| | R Thalamus | 3.19 | 14 | -20 | 16 |
| 32 | | | | | |
| | L Thalamus | 3.26 | -14 | -18 | 18 |
| 24 | | | | | |
| | L Inferior Temporal Gyrus, anterior division | 3.25 | -50 | -2 | -34 |
| 24 | | | | | |
| | R Amygdala | 3.07 | 24 | -14 | -12 |
| 24 | | | | | |

| | | | | | |
|---|---|---|---|---|---|
| | R Amygdala | 3.29 | 22 | -6 | -12 |
| 24 | | | | | |
| | R Middle Temporal Gyrus, temporooccipital part | 2.84 | 64 | -38 | 4 |
| 24 | | | | | |
| | L Frontal Pole | 2.70 | -42 | 52 | 16 |
| 24 | | | | | |
| | L Caudate | 2.91 | -14 | -10 | 20 |
| 24 | | | | | |
| | L Precuneus Cortex | 2.59 | -10 | -46 | 50 |
| 16 | | | | | |
| | R Parahippocampal Gyrus, anterior division | 3.06 | 24 | -10 | -32 |
| 16 | | | | | |
| | R Brain-Stem | 2.75 | 4 | -34 | -22 |
| 16 | | | | | |
| | R Brain-Stem | 2.89 | 8 | -12 | -24 |
| 16 | | | | | |
| | R Brain-Stem | 2.73 | 6 | -22 | -20 |
| 16 | | | | | |
| | L Thalamus | 2.87 | -8 | -2 | 10 |
| 16 | | | | | |
| | L Thalamus | 3.48 | -6 | -26 | 14 |
| 16 | | | | | |
| | R Precentral Gyrus | 2.43 | 60 | 10 | 24 |
| 16 | | | | | |
| | R Precentral Gyrus | 2.56 | 40 | 2 | 32 |
| 16 | | | | | |
| | R Cingulate Gyrus, anterior division | 2.83 | 2 | -10 | 42 |
| 16 | | | | | |
| | R Postcentral Gyrus | 2.70 | 46 | -28 | 52 |
| 16 | | | | | |
| | R Superior Parietal Lobule | 2.69 | 16 | -50 | 64 |
| 8 | | | | | |
| | L Brain-Stem | 2.67 | -4 | -44 | -60 |
| 8 | | | | | |
| | R Brain-Stem | 2.60 | 4 | -46 | -58 |
| 8 | | | | | |
| | L Brain-Stem | 2.54 | -2 | -40 | -56 |
| 8 | | | | | |
| | R Brain-Stem | 2.76 | 8 | -44 | -48 |
| 8 | | | | | |
| | R Parahippocampal Gyrus, anterior division | 2.69 | 22 | -8 | -36 |
| 8 | | | | | |
| | R Temporal Pole | 2.62 | 54 | 10 | -36 |

| | | | | | |
|---|---|---|---|---|---|
| 8 | | | | | |
| | R Brain-Stem | 2.58 | 12 | -16 | -28 |
| 8 | | | | | |
| | L Brain-Stem | 2.43 | -4 | -16 | -20 |
| 8 | | | | | |
| | R Frontal Orbital Cortex | 2.46 | 34 | 26 | -14 |
| 8 | | | | | |
| | R Amygdala | 2.72 | 28 | -8 | -12 |
| 8 | | | | | |
| | R Thalamus | 2.44 | 8 | -6 | -2 |
| 8 | | | | | |
| | L Caudate | 2.38 | -16 | 18 | -2 |
| 8 | | | | | |
| | R Thalamus | 2.82 | 0 | -8 | 0 |
| 8 | | | | | |
| | L Pallidum | 2.46 | -12 | 2 | 0 |
| 8 | | | | | |
| | R Occipital Pole | 2.52 | 32 | -90 | 4 |
| 8 | | | | | |
| | R Thalamus | 2.72 | 14 | -12 | 6 |
| 8 | | | | | |
| | L Caudate | 2.73 | -8 | 10 | 6 |
| 8 | | | | | |
| | R Putamen | 2.62 | 28 | -12 | 8 |
| 8 | | | | | |
| | R Thalamus | 2.65 | 12 | -10 | 8 |
| 8 | | | | | |
| | R Thalamus | 2.97 | 4 | -18 | 14 |
| 8 | | | | | |
| | R Parietal Operculum Cortex | 2.92 | 44 | -20 | 18 |
| 8 | | | | | |
| | L Frontal Pole | 2.50 | -34 | 44 | 20 |
| 8 | | | | | |
| | R Precentral Gyrus | 2.97 | 56 | 6 | 26 |
| 8 | | | | | |
| | L Frontal Pole | 2.53 | -26 | 44 | 26 |
| 8 | | | | | |
| | R Cingulate Gyrus, anterior division | 2.61 | 4 | 8 | 28 |
| 8 | | | | | |
| | L Frontal Pole | 2.64 | -24 | 46 | 28 |
| 8 | | | | | |
| | L Precentral Gyrus | 2.53 | -32 | -10 | 58 |
| 8 | | | | | |
| | R Superior Frontal Gyrus | 2.54 | 26 | 2 | 64 |
| 8 | | | | | |
| | R Precentral Gyrus | 2.88 | 28 | -20 | 68 |

**Supplementary Table S17. [CSphasic > CTphasic].** Descriptive statistics for clusters and local maxima showing greater activation for Certain-Safety relative to Certain-Threat anticipation for the OSP Phasic regressor (FDR $q$<0.05, whole-brain corrected).

| $mm^3$ | Label | *t* | *x* | *y* | *z* |
|---|---|---|---|---|---|
| 5,896 | | | | | |
| | L Precentral Gyrus | 6.35 | -8 | -20 | 80 |
| | L Superior Parietal Lobule | 4.92 | -30 | -42 | 68 |
| | L Postcentral Gyrus | 6.59 | -10 | -44 | 72 |
| 2,456 | | | | | |
| | R Precentral Gyrus | 6.66 | 12 | -24 | 80 |
| | R Postcentral Gyrus | 5.13 | 10 | -42 | 72 |
| | R Precuneus Cortex | 4.96 | 6 | -44 | 68 |
| 632 | | | | | |
| | R Frontal Pole | 4.18 | 26 | 38 | 50 |
| | R Superior Frontal Gyrus | 5.19 | 24 | 30 | 48 |
| 432 | | | | | |
| | R Cingulate Gyrus, posterior division | 4.69 | 0 | -46 | 14 |
| 312 | | | | | |
| | R Postcentral Gyrus | 4.78 | 66 | -12 | 32 |
| 288 | | | | | |
| | R Superior Parietal Lobule | 5.58 | 30 | -56 | 60 |
| 280 | | | | | |
| | R Superior Parietal Lobule | 4.79 | 44 | -44 | 62 |
| 272 | | | | | |
| | L Postcentral Gyrus | 4.96 | -58 | -12 | 46 |
| 152 | | | | | |
| | L Lateral Occipital Cortex, superior division | 4.21 | -32 | -64 | 58 |
| 120 | | | | | |
| | L Temporal Pole | 4.58 | -36 | 22 | -34 |
| 96 | | | | | |
| | R Precuneus Cortex | 4.76 | 2 | -60 | 16 |
| 96 | | | | | |
| | R Lateral Occipital Cortex, superior division | 3.94 | 30 | -70 | 60 |
| 88 | | | | | |
| | R Middle Frontal Gyrus | 4.13 | 30 | 18 | 60 |
| 72 | | | | | |
| | R Lateral Occipital Cortex, superior division | 4.23 | 18 | -82 | 48 |
| 72 | | | | | |
| | R Lateral Occipital Cortex, superior division | 4.04 | 26 | -78 | 50 |
| 64 | | | | | |

| | | | | | |
|---|---|---|---|---|---|
| | R Lateral Occipital Cortex, superior division | 3.67 | 50 | -72 | 34 |
| 64 | | | | | |
| | L Lateral Occipital Cortex, superior division | 4.01 | -26 | -82 | 44 |
| 56 | | | | | |
| | L Frontal Pole | 4.11 | -48 | 44 | -12 |
| 56 | | | | | |
| | R Superior Frontal Gyrus | 4.35 | 12 | 32 | 58 |
| 48 | | | | | |
| | R Temporal Pole | 3.84 | 40 | 24 | -26 |
| 48 | | | | | |
| | L Superior Frontal Gyrus | 3.51 | -24 | 16 | 52 |
| 40 | | | | | |
| | L Subcallosal Cortex | 3.67 | -2 | 14 | -10 |
| 40 | | | | | |
| | R Frontal Pole | 4.40 | 46 | 54 | -8 |
| 40 | | | | | |
| | R Postcentral Gyrus | 3.97 | 58 | -18 | 36 |
| 40 | | | | | |
| | L Superior Frontal Gyrus | 3.89 | -4 | 16 | 66 |
| 32 | | | | | |
| | R Frontal Pole | 3.96 | 12 | 42 | 48 |
| 32 | | | | | |
| | L Postcentral Gyrus | 3.89 | -54 | -28 | 52 |
| 24 | | | | | |
| | R Temporal Pole | 3.95 | 36 | 22 | -36 |
| 24 | | | | | |
| | L Temporal Pole | 3.98 | -30 | 12 | -34 |
| 24 | | | | | |
| | R Frontal Pole | 3.86 | 14 | 58 | -22 |
| 24 | | | | | |
| | R Frontal Medial Cortex | 4.51 | 4 | 34 | -20 |
| 24 | | | | | |
| | R Parahippocampal Gyrus, posterior division | 3.84 | 28 | -34 | -14 |
| 24 | | | | | |
| | R Lingual Gyrus | 3.78 | 4 | -86 | -4 |
| 24 | | | | | |
| | L Frontal Pole | 3.80 | -44 | 48 | -6 |
| 24 | | | | | |
| | L Frontal Pole | 3.79 | -22 | 62 | -6 |
| 24 | | | | | |
| | R Lingual Gyrus | 3.66 | 12 | -44 | -4 |
| 24 | | | | | |

| | | | | | |
|---|---|---|---|---|---|
| | R Postcentral Gyrus | 3.67 | 58 | -14 | 30 |
| 24 | | | | | |
| | R Cuneal Cortex | 3.56 | 4 | -82 | 42 |
| 24 | | | | | |
| | R Lateral Occipital Cortex, superior division | 3.46 | 32 | -76 | 42 |
| 24 | | | | | |
| | R Lateral Occipital Cortex, superior division | 3.93 | 36 | -70 | 52 |
| 24 | | | | | |
| | L Superior Parietal Lobule | 3.75 | -34 | -54 | 58 |
| 24 | | | | | |
| | L Lateral Occipital Cortex, superior division | 3.49 | -22 | -64 | 62 |
| 16 | | | | | |
| | L Frontal Orbital Cortex | 3.78 | -36 | 20 | -24 |
| 16 | | | | | |
| | R Parahippocampal Gyrus, posterior division | 3.66 | 22 | -30 | -16 |
| 16 | | | | | |
| | L Frontal Pole | 3.69 | -42 | 58 | -8 |
| 16 | | | | | |
| | L Paracingulate Gyrus | 3.45 | -10 | 46 | -6 |
| 16 | | | | | |
| | R Precentral Gyrus | 3.49 | 60 | -6 | 44 |
| 16 | | | | | |
| | R Lateral Occipital Cortex, superior division | 3.59 | 40 | -62 | 54 |
| 16 | | | | | |
| | L Superior Frontal Gyrus | 3.92 | -6 | 38 | 56 |
| 16 | | | | | |
| | L Lateral Occipital Cortex, superior division | 3.55 | -24 | -66 | 58 |
| 16 | | | | | |
| | R Lateral Occipital Cortex, superior division | 3.53 | 8 | -66 | 66 |
| 16 | | | | | |
| | L Superior Frontal Gyrus | 3.59 | -6 | 20 | 68 |
| 8 | | | | | |
| | R Parahippocampal Gyrus, posterior division | 3.56 | 26 | -28 | -22 |
| 8 | | | | | |
| | L Frontal Pole | 3.50 | -16 | 38 | -22 |
| 8 | | | | | |
| | R Subcallosal Cortex | 3.44 | 4 | 26 | -20 |

| | | | | | |
|---|---|---|---|---|---|
| 8 | | | | | |
| | R Temporal Fusiform Cortex, posterior division | 3.72 | 40 | -26 | -18 |
| 8 | | | | | |
| | L Lingual Gyrus | 3.41 | -4 | -86 | -14 |
| 8 | | | | | |
| | R Parahippocampal Gyrus, posterior division | 3.62 | 30 | -36 | -12 |
| 8 | | | | | |
| | R Frontal Pole | 4.00 | 46 | 48 | -12 |
| 8 | | | | | |
| | L Lingual Gyrus | 3.53 | -2 | -78 | -10 |
| 8 | | | | | |
| | R Occipital Pole | 3.54 | 10 | -96 | -4 |
| 8 | | | | | |
| | R Frontal Pole | 3.40 | 40 | 60 | 2 |
| 8 | | | | | |
| | L Heschls Gyrus (includes H1 and H2) | 3.43 | -50 | -16 | 10 |
| 8 | | | | | |
| | L Frontal Pole | 3.49 | -10 | 70 | 10 |
| 8 | | | | | |
| | R Lateral Occipital Cortex, superior division | 3.43 | 50 | -78 | 26 |
| 8 | | | | | |
| | L Lateral Occipital Cortex, superior division | 3.52 | -42 | -80 | 38 |
| 8 | | | | | |
| | R Lateral Occipital Cortex, superior division | 3.38 | 34 | -80 | 42 |
| 8 | | | | | |
| | R Lateral Occipital Cortex, superior division | 3.57 | 40 | -76 | 44 |
| 8 | | | | | |
| | R Frontal Pole | 3.37 | 14 | 50 | 44 |
| 8 | | | | | |
| | L Superior Frontal Gyrus | 3.40 | -24 | 18 | 46 |
| 8 | | | | | |
| | R Frontal Pole | 3.61 | 18 | 46 | 46 |
| 8 | | | | | |
| | L Frontal Pole | 3.39 | -18 | 42 | 50 |
| 8 | | | | | |
| | L Superior Frontal Gyrus | 3.41 | -16 | 18 | 58 |
| 8 | | | | | |
| | R Lateral Occipital Cortex, superior division | 3.36 | 22 | -64 | 64 |

| | | | | | |
|---|---|---|---|---|---|
| 8 | | | | | |
| | R Lateral Occipital Cortex, superior division | 3.42 | 26 | -62 | 66 |
| 8 | | | | | |
| | L Superior Parietal Lobule | 3.36 | -28 | -56 | 66 |

**Supplementary Table S18. [$US_{phasic}$ > $UT_{phasic}$].** Descriptive statistics for clusters and local maxima showing greater activation for Uncertain-Safety relative to Uncertain-Threat anticipation for the OSP Phasic regressor (FDR $q$<0.05, whole-brain corrected).

| ***mm³*** | **Label** | ***t*** | ***x*** | ***y*** | ***z*** |
|---|---|---|---|---|---|
| 2,776 | | | | | |
| | L Postcentral Gyrus | 6.74 | -46 | -16 | 62 |
| | L Precentral Gyrus | 5.91 | -42 | -18 | 66 |
| | L Superior Parietal Lobule | 4.94 | -38 | -42 | 62 |
| 1,432 | | | | | |
| | R Occipital Pole | 6.22 | 14 | -96 | -2 |
| 1,304 | | | | | |
| | L Frontal Pole | 4.98 | -12 | 68 | -16 |
| | R Frontal Pole | 5.39 | 2 | 68 | -14 |
| 888 | | | | | |
| | L Lateral Occipital Cortex, superior division | 5.69 | -18 | -68 | 64 |
| 856 | | | | | |
| | R Postcentral Gyrus | 5.85 | 46 | -20 | 64 |
| 784 | | | | | |
| | L Occipital Pole | 5.78 | -20 | -98 | -10 |
| 760 | | | | | |
| | R Frontal Orbital Cortex | 5.56 | 14 | 16 | -16 |
| 528 | | | | | |
| | L Middle Frontal Gyrus | 4.46 | -32 | 0 | 66 |
| | L Precentral Gyrus | 4.45 | -36 | -6 | 66 |
| 432 | | | | | |
| | R Postcentral Gyrus | 4.39 | 42 | -36 | 60 |
| 344 | | | | | |
| | R Lingual Gyrus | 4.68 | 10 | -70 | -10 |
| 344 | | | | | |
| | L Occipital Pole | 5.25 | -20 | -102 | 6 |
| 280 | | | | | |
| | L Frontal Orbital Cortex | 4.99 | -26 | 30 | -20 |
| 224 | | | | | |
| | L Frontal Orbital Cortex | 4.15 | -18 | 10 | -22 |
| 200 | | | | | |
| | R Subcallosal Cortex | 5.15 | 2 | 18 | -6 |
| | L Subcallosal Cortex | 5.28 | -2 | 16 | -6 |
| 192 | | | | | |
| | R Frontal Pole | 4.76 | 6 | 74 | 2 |
| 192 | | | | | |
| | R Precentral Gyrus | 4.39 | 38 | -6 | 66 |
| 152 | | | | | |
| | L Brain-Stem | 4.27 | -2 | -34 | -58 |
| | R Brain-Stem | 3.91 | 8 | -36 | -60 |
| 144 | | | | | |

| | | | | | |
|---|---|---|---|---|---|
| | L Temporal Pole | 4.57 | -46 | 2 | -38 |
| 128 | | | | | |
| | L Hippocampus | 5.13 | -20 | -22 | -16 |
| 128 | | | | | |
| | R Occipital Pole | 3.85 | 28 | -98 | 6 |
| 120 | | | | | |
| | L Frontal Pole | 4.80 | -12 | 40 | -24 |
| 120 | | | | | |
| | R Hippocampus | 4.66 | 20 | -20 | -16 |
| 120 | | | | | |
| | L Lateral Occipital Cortex, superior division | 3.68 | -32 | -88 | 36 |
| 120 | | | | | |
| | R Precentral Gyrus | 3.47 | 4 | -16 | 76 |
| 104 | | | | | |
| | L Frontal Pole | 4.17 | -14 | 72 | 0 |
| 96 | | | | | |
| | L Occipital Fusiform Gyrus | 3.93 | -20 | -82 | -14 |
| 88 | | | | | |
| | R Temporal Pole | 4.67 | 40 | 20 | -30 |
| 88 | | | | | |
| | L Frontal Pole | 3.57 | -10 | 48 | -22 |
| 88 | | | | | |
| | R Lateral Occipital Cortex, superior division | 4.19 | 46 | -74 | 42 |
| 88 | | | | | |
| | R Postcentral Gyrus | 4.16 | 56 | -22 | 54 |
| 72 | | | | | |
| | L Frontal Pole | 4.48 | -10 | 52 | -28 |
| 72 | | | | | |
| | R Lateral Occipital Cortex, superior division | 4.15 | 14 | -78 | 56 |
| 64 | | | | | |
| | L Lingual Gyrus | 4.43 | -2 | -74 | -8 |
| 64 | | | | | |
| | L Precentral Gyrus | 4.15 | -32 | -10 | 50 |
| 64 | | | | | |
| | L Superior Parietal Lobule | 4.40 | -30 | -48 | 48 |
| 56 | | | | | |
| | L Lateral Occipital Cortex, superior division | 3.80 | -22 | -86 | 44 |
| 56 | | | | | |
| | L Superior Parietal Lobule | 3.57 | -22 | -50 | 72 |
| 56 | | | | | |
| | L Superior Parietal Lobule | 3.74 | -10 | -54 | 72 |
| 56 | | | | | |

| | | | | | |
|---|---|---|---|---|---|
| | R Superior Frontal Gyrus | 3.50 | 22 | -8 | 72 |
| 56 | | | | | |
| | L Juxtapositional Lobule Cortex (formerly Supplementary Motor Cortex) | 3.89 | -4 | 2 | 72 |
| 56 | | | | | |
| | R Superior Parietal Lobule | 3.92 | 22 | -50 | 74 |
| 48 | | | | | |
| | R Temporal Pole | 3.74 | 26 | 6 | -34 |
| 48 | | | | | |
| | L Lingual Gyrus | 3.67 | -12 | -78 | -12 |
| | L Occipital Fusiform Gyrus | 3.67 | -16 | -78 | -16 |
| 48 | | | | | |
| | L Cingulate Gyrus, posterior division | 4.70 | -6 | -44 | 4 |
| 48 | | | | | |
| | L Caudate | 4.33 | -8 | 0 | 8 |
| 48 | | | | | |
| | L Postcentral Gyrus | 3.83 | -58 | -10 | 46 |
| 40 | | | | | |
| | L Frontal Orbital Cortex | 3.78 | -48 | 28 | -12 |
| 32 | | | | | |
| | R Inferior Temporal Gyrus, anterior division | 3.68 | 42 | -2 | -40 |
| 32 | | | | | |
| | L Subcallosal Cortex | 3.98 | -4 | 14 | -12 |
| 32 | | | | | |
| | R Lingual Gyrus | 4.57 | 12 | -58 | -4 |
| 32 | | | | | |
| | R Paracingulate Gyrus | 4.14 | 14 | 48 | 2 |
| 32 | | | | | |
| | L Frontal Pole | 3.99 | -10 | 74 | 6 |
| 32 | | | | | |
| | L Postcentral Gyrus | 3.66 | -56 | -26 | 52 |
| 32 | | | | | |
| | L Postcentral Gyrus | 3.75 | -48 | -36 | 58 |
| 32 | | | | | |
| | R Lateral Occipital Cortex, superior division | 3.70 | 14 | -64 | 66 |
| 32 | | | | | |
| | R Superior Parietal Lobule | 3.99 | 38 | -44 | 68 |
| 24 | | | | | |
| | L Temporal Pole | 3.56 | -34 | 16 | -30 |
| 24 | | | | | |
| | R Frontal Pole | 3.66 | 28 | 34 | -20 |
| 24 | | | | | |
| | R Lingual Gyrus | 3.62 | 22 | -58 | -10 |
| 24 | | | | | |

| | | | | | |
|---|---|---|---|---|---|
| | R Lingual Gyrus | 3.78 | 2 | -72 | -6 |
| 24 | | | | | |
| | R Lingual Gyrus | 3.65 | 2 | -64 | 2 |
| 24 | | | | | |
| | R Occipital Pole | 3.45 | 24 | -96 | 20 |
| 24 | | | | | |
| | L Superior Frontal Gyrus | 3.69 | -24 | -4 | 56 |
| 16 | | | | | |
| | L Brain-Stem | 3.66 | -8 | -48 | -52 |
| 16 | | | | | |
| | R Brain-Stem | 4.17 | 8 | -30 | -36 |
| 16 | | | | | |
| | R Temporal Pole | 3.73 | 36 | 14 | -32 |
| 16 | | | | | |
| | R Frontal Pole | 3.52 | 18 | 36 | -20 |
| 16 | | | | | |
| | L Frontal Orbital Cortex | 3.61 | -14 | 20 | -16 |
| 16 | | | | | |
| | R Occipital Fusiform Gyrus | 3.45 | 16 | -88 | -12 |
| 16 | | | | | |
| | R Hippocampus | 4.21 | 28 | -18 | -12 |
| 16 | | | | | |
| | R Lingual Gyrus | 4.12 | 30 | -46 | -4 |
| 16 | | | | | |
| | R Hippocampus | 3.93 | 32 | -34 | -2 |
| 16 | | | | | |
| | Bed Nucleus of the Stria Terminalis | 3.37 | 8 | 4 | 2 |
| | R Occipital Pole | 3.43 | 38 | -94 | 8 |
| 16 | | | | | |
| | R Lateral Occipital Cortex, inferior division | 3.50 | 56 | -72 | 8 |
| 16 | | | | | |
| | L Precentral Gyrus | 3.57 | -52 | 0 | 24 |
| 16 | | | | | |
| | R Lateral Occipital Cortex, superior division | 3.40 | 22 | -62 | 64 |
| 16 | | | | | |
| | R Postcentral Gyrus | 3.54 | 12 | -42 | 74 |
| 16 | | | | | |
| | R Postcentral Gyrus | 3.43 | 10 | -38 | 80 |
| 16 | | | | | |
| | R Precentral Gyrus | 3.71 | 10 | -26 | 80 |
| 8 | | | | | |
| | L Brain-Stem | 3.29 | -8 | -40 | -60 |
| 8 | | | | | |
| | R Temporal Fusiform Cortex, anterior division | 3.42 | 24 | -6 | -42 |

| | | | | | |
|---|---|---|---|---|---|
| 8 | | | | | |
| | R Inferior Temporal Gyrus, anterior division | 3.38 | 42 | 2 | -40 |
| 8 | | | | | |
| | R Temporal Pole | 3.35 | 30 | 10 | -34 |
| 8 | | | | | |
| | L Temporal Pole | 3.56 | -50 | 14 | -32 |
| 8 | | | | | |
| | L Temporal Pole | 3.47 | -24 | 4 | -26 |
| 8 | | | | | |
| | R Frontal Pole | 3.39 | 14 | 34 | -26 |
| 8 | | | | | |
| | R Parahippocampal Gyrus, anterior division | 3.79 | 16 | -6 | -24 |
| 8 | | | | | |
| | L Frontal Pole | 3.47 | -4 | 62 | -24 |
| 8 | | | | | |
| | R Amygdala | 3.61 | 12 | -4 | -18 |
| 8 | | | | | |
| | L Hippocampus | 3.81 | -36 | -26 | -12 |
| 8 | | | | | |
| | L Lingual Gyrus | 3.75 | -12 | -56 | -6 |
| 8 | | | | | |
| | L Frontal Medial Cortex | 3.31 | -8 | 54 | -6 |
| 8 | | | | | |
| | Bed Nucleus of the Stria Terminalis | 3.46 | -6 | 2 | -4 |
| 8 | | | | | |
| | L Frontal Pole | 3.59 | -2 | 66 | 0 |
| 8 | | | | | |
| | R Lateral Occipital Cortex, inferior division | 3.29 | 44 | -64 | 8 |
| 8 | | | | | |
| | L Caudate | 3.66 | -18 | 14 | 14 |
| 8 | | | | | |
| | R Postcentral Gyrus | 3.35 | 64 | -12 | 40 |
| 8 | | | | | |
| | L Postcentral Gyrus | 3.77 | -40 | -34 | 56 |
| 8 | | | | | |
| | R Lateral Occipital Cortex, superior division | 3.45 | 26 | -58 | 60 |

**Supplementary Table S19. [UT$_{phasic}$ > CT$_{phasic}$].** Descriptive statistics for clusters and local maxima showing greater activation for Uncertain-Threat relative to Certain-Threat anticipation for the OSP Phasic regressor (FDR $q$<0.05, whole-brain corrected).

| ***mm³*** | **Label** | ***t*** | ***x*** | ***y*** | ***z*** |
|---|---|---|---|---|---|
| 43,448 | | | | | |
| | R Thalamus | 4.15 | 14 | -24 | -4 |
| | R Temporal Fusiform Cortex, posterior division | 3.42 | 34 | -30 | -18 |
| | R Brain-Stem | 3.28 | 8 | -30 | -14 |
| | L Temporal Fusiform Cortex, posterior division | 2.85 | -30 | -34 | -20 |
| | R Parahippocampal Gyrus, posterior division | 3.1 | 20 | -36 | -18 |
| | L Parahippocampal Gyrus, posterior division | 5.41 | -20 | -40 | -12 |
| | R Lingual Gyrus | 6.72 | 32 | -40 | -10 |
| | L Lingual Gyrus | 6.96 | -24 | -46 | -6 |
| | L Temporal Occipital Fusiform Cortex | 8.24 | -30 | -48 | -8 |
| | R Temporal Occipital Fusiform Cortex | 6.63 | 28 | -52 | -8 |
| | L Precuneus Cortex | 4.36 | -18 | -58 | 8 |
| | R Precuneus Cortex | 3.52 | 16 | -60 | 12 |
| | L Supracalcarine Cortex | 4.52 | -18 | -64 | 14 |
| | R Occipital Fusiform Gyrus | 4.93 | 24 | -66 | -10 |
| | L Occipital Fusiform Gyrus | 5.84 | -22 | -68 | -10 |
| | R Supracalcarine Cortex | 4.73 | 2 | -76 | 14 |
| | R Intracalcarine Cortex | 5.52 | 16 | -80 | 8 |
| | R Lateral Occipital Cortex, superior division | 5.09 | 18 | -80 | 40 |
| | L Lateral Occipital Cortex, superior division | 5.63 | -42 | -82 | 18 |
| | L Cuneal Cortex | 6.42 | -2 | -86 | 34 |
| | R Cuneal Cortex | 6.14 | 6 | -86 | 36 |
| | L Intracalcarine Cortex | 6.32 | -8 | -88 | 6 |
| | L Occipital Pole | 6.51 | -8 | -90 | 36 |
| | R Occipital Pole | 7.96 | 2 | -96 | -2 |
| 4,432 | | | | | |
| | R Frontal Pole | 5.4 | 48 | 48 | -4 |
| | R Frontal Orbital Cortex | 5.21 | 44 | 30 | -12 |
| | R Temporal Pole | 4.19 | 56 | 16 | -12 |
| 3,888 | | | | | |
| | R Superior Temporal Gyrus, anterior division | 3.25 | 62 | 0 | -12 |
| | R Planum Polare | 5.21 | 42 | -12 | -6 |
| | R Superior Temporal Gyrus, posterior division | 3.88 | 68 | -14 | 2 |
| | R Central Opercular Cortex | 4.84 | 58 | -16 | 10 |

| | | | | | |
|---|---|---|---|---|---|
| | R Planum Temporale | 4.66 | 54 | -22 | 8 |
| | R Heschls Gyrus (includes H1 and H2) | 5.78 | 44 | -26 | 10 |
| 2,528 | | | | | |
| | L Superior Temporal Gyrus, anterior division | 3.13 | -62 | -8 | -2 |
| | L Central Opercular Cortex | 4.68 | -40 | -18 | 20 |
| | L Heschls Gyrus (includes H1 and H2) | 4.61 | -48 | -26 | 6 |
| | L Insular Cortex | 4.5 | -32 | -28 | 16 |
| | L Parietal Operculum Cortex | 4.92 | -34 | -30 | 18 |
| | L Planum Temporale | 4.78 | -38 | -34 | 12 |
| 2,224 | | | | | |
| | L Frontal Pole | 5.29 | -46 | 46 | -12 |
| | L Inferior Frontal Gyrus, pars triangularis | 4.46 | -48 | 32 | -4 |
| | L Frontal Orbital Cortex | 4.23 | -46 | 28 | -6 |
| 1,816 | | | | | |
| | R Frontal Pole | 4.97 | 10 | 50 | 46 |
| 1,368 | | | | | |
| | R Lateral Occipital Cortex, inferior division | 5.06 | 48 | -74 | 12 |
| | R Lateral Occipital Cortex, superior division | 4.5 | 50 | -78 | 22 |
| 1,176 | | | | | |
| | R Precentral Gyrus | 4.07 | 52 | -6 | 28 |
| | R Postcentral Gyrus | 5.05 | 64 | -8 | 32 |
| 1,168 | | | | | |
| | L Inferior Frontal Gyrus, pars triangularis | 3.99 | -56 | 28 | 16 |
| | L Inferior Frontal Gyrus, pars opercularis | 3.68 | -58 | 20 | 16 |
| 1,144 | | | | | |
| | L Frontal Pole | 5.66 | -14 | 58 | 28 |
| | L Superior Frontal Gyrus | 5.71 | -4 | 56 | 30 |
| 936 | | | | | |
| | L Temporal Pole | 3.65 | -58 | 4 | -4 |
| | L Planum Polare | 4.99 | -48 | -12 | -2 |
| | L Insular Cortex | 4.19 | -40 | -14 | -6 |
| 856 | | | | | |
| | L Superior Temporal Gyrus, posterior division | 3.62 | -56 | -28 | -2 |
| | L Middle Temporal Gyrus, posterior division | 4.71 | -50 | -38 | 0 |
| 680 | | | | | |
| | L Superior Frontal Gyrus | 4.56 | -2 | 48 | 46 |
| | L Frontal Pole | 4.16 | -12 | 44 | 48 |
| 448 | | | | | |
| | R Frontal Pole | 3.35 | 50 | 36 | 22 |
| | R Middle Frontal Gyrus | 4.15 | 52 | 28 | 26 |
| 376 | | | | | |
| | R Superior Frontal Gyrus | 3.93 | 22 | 30 | 48 |

| | | | | | |
|---|---|---|---|---|---|
| | R Middle Frontal Gyrus | 3.22 | 28 | 20 | 54 |
| 312 | | | | | |
| | L Superior Parietal Lobule | 5.83 | -26 | -56 | 64 |
| 256 | | | | | |
| | L Frontal Orbital Cortex | 4.67 | -40 | 22 | -14 |
| 256 | | | | | |
| | L Postcentral Gyrus | 3.89 | -54 | -18 | 50 |
| 200 | | | | | |
| | L Temporal Pole | 4.38 | -52 | 8 | -20 |
| 192 | | | | | |
| | L Precentral Gyrus | 4.38 | -52 | -8 | 32 |
| 192 | | | | | |
| | R Supramarginal Gyrus, anterior division | 3.98 | 62 | -24 | 46 |
| 192 | | | | | |
| | L Middle Frontal Gyrus | 4.06 | -38 | 16 | 48 |
| 176 | | | | | |
| | R Frontal Pole | 4.55 | 10 | 68 | 20 |
| 176 | | | | | |
| | R Precentral Gyrus | 3.81 | 38 | -12 | 44 |
| 168 | | | | | |
| | L Precentral Gyrus | 4.8 | -34 | -22 | 44 |
| | L Postcentral Gyrus | 3.31 | -42 | -26 | 48 |
| 152 | | | | | |
| | L Superior Temporal Gyrus, posterior division | 3.38 | -68 | -32 | 12 |
| | L Planum Temporale | 3.64 | -62 | -36 | 14 |
| 144 | | | | | |
| | R Temporal Pole | 3.72 | 40 | 26 | -34 |
| 144 | | | | | |
| | L Postcentral Gyrus | 4.36 | -20 | -32 | 72 |
| 136 | | | | | |
| | L Frontal Pole | 3.86 | -24 | 60 | -8 |
| 128 | | | | | |
| | L Temporal Pole | 4.03 | -38 | 24 | -36 |
| 128 | | | | | |
| | R Inferior Frontal Gyrus, pars opercularis | 4.09 | 60 | 20 | 14 |
| 120 | | | | | |
| | L Angular Gyrus | 3.45 | -46 | -54 | 26 |
| 120 | | | | | |
| | L Precentral Gyrus | 3.36 | -26 | -26 | 52 |
| 120 | | | | | |
| | L Superior Frontal Gyrus | 4.25 | -4 | 16 | 62 |
| 112 | | | | | |
| | L Supramarginal Gyrus, posterior division | 3.99 | -60 | -46 | 12 |
| 104 | | | | | |
| | L Cingulate Gyrus, posterior division | 3.8 | -2 | -46 | 12 |

| | | | | | |
|---|---|---|---|---|---|
| 104 | | | | | |
| | L Postcentral Gyrus | 3.71 | -42 | -26 | 58 |
| 104 | | | | | |
| | R Precentral Gyrus | 3.91 | 12 | -28 | 70 |
| 96 | | | | | |
| | R Temporal Pole | 3.65 | 52 | 8 | -20 |
| | R Superior Temporal Gyrus, anterior division | 4.13 | 50 | 2 | -20 |
| 96 | | | | | |
| | R Frontal Pole | 3.41 | 38 | 62 | -10 |
| 96 | | | | | |
| | R Supracalcarine Cortex | 3.78 | 22 | -64 | 16 |
| 96 | | | | | |
| | L Postcentral Gyrus | 3.85 | -6 | -42 | 64 |
| 88 | | | | | |
| | R Cingulate Gyrus, posterior division | 3.77 | 18 | -46 | 2 |
| | R Lingual Gyrus | 3.25 | 16 | -50 | 0 |
| 88 | | | | | |
| | L Postcentral Gyrus | 3.65 | -66 | -8 | 26 |
| 80 | | | | | |
| | L Superior Temporal Gyrus, anterior division | 3.85 | -52 | 0 | -16 |
| 80 | | | | | |
| | R Superior Temporal Gyrus, posterior division | 3.11 | 68 | -32 | 4 |
| 80 | | | | | |
| | R Precuneus Cortex | 4.51 | 22 | -54 | 8 |
| 80 | | | | | |
| | R Middle Temporal Gyrus, temporooccipital part | 3.33 | 68 | -40 | 6 |
| 80 | | | | | |
| | R Inferior Frontal Gyrus, pars opercularis | 3.36 | 52 | 16 | 28 |
| 72 | | | | | |
| | R Periaqueductal gray | 3.15 | 0 | -32 | -10 |
| | R Brain-Stem | 3.24 | 4 | -36 | -8 |
| 72 | | | | | |
| | L Putamen | 4.8 | -32 | -12 | -6 |
| 64 | | | | | |
| | R Inferior Temporal Gyrus, temporooccipital part | 4.31 | 56 | -54 | -24 |
| 64 | | | | | |
| | L Temporal Fusiform Cortex, posterior division | 3.61 | -34 | -26 | -22 |
| 64 | | | | | |
| | L Superior Frontal Gyrus | 3.5 | -4 | 28 | 62 |
| 64 | | | | | |

| | | | | | |
|---|---|---|---|---|---|
| | L Postcentral Gyrus | 3.19 | -4 | -40 | 72 |
| 56 | | | | | |
| | L Temporal Pole | 3.07 | -36 | 20 | -26 |
| 56 | | | | | |
| | R Middle Temporal Gyrus, anterior division | 3.37 | 58 | 4 | -22 |
| 56 | | | | | |
| | L Frontal Pole | 3.28 | -40 | 56 | -4 |
| 56 | | | | | |
| | R Lingual Gyrus | 3.92 | 12 | -56 | 2 |
| 56 | | | | | |
| | R Central Opercular Cortex | 3.63 | 42 | -8 | 12 |
| 56 | | | | | |
| | L Lateral Occipital Cortex, superior division | 3.31 | -50 | -70 | 26 |
| 56 | | | | | |
| | R Superior Frontal Gyrus | 3.77 | 4 | 36 | 58 |
| 56 | | | | | |
| | L Postcentral Gyrus | 3.3 | -26 | -40 | 68 |
| 48 | | | | | |
| | L Brain-Stem | 3.05 | -6 | -36 | -60 |
| 48 | | | | | |
| | L Temporal Pole | 3.16 | -48 | 12 | -12 |
| 48 | | | | | |
| | L Precentral Gyrus | 3.56 | -14 | -24 | 72 |
| 40 | | | | | |
| | L Temporal Pole | 3.45 | -50 | 20 | -18 |
| 40 | | | | | |
| | R Supramarginal Gyrus, posterior division | 3.7 | 56 | -38 | 14 |
| 40 | | | | | |
| | R Cingulate Gyrus, posterior division | 3.16 | 0 | -44 | 20 |
| 40 | | | | | |
| | L Precentral Gyrus | 3.86 | -36 | -14 | 46 |
| 40 | | | | | |
| | R Superior Frontal Gyrus | 2.98 | 24 | 26 | 58 |
| 32 | | | | | |
| | R Temporal Pole | 4.19 | 48 | 24 | -26 |
| 32 | | | | | |
| | L Middle Temporal Gyrus, posterior division | 3.1 | -64 | -36 | -4 |
| 32 | | | | | |
| | L Central Opercular Cortex | 3.54 | -48 | -12 | 16 |
| 32 | | | | | |
| | L Angular Gyrus | 3.22 | -60 | -56 | 24 |
| 32 | | | | | |
| | L Superior Frontal Gyrus | 3.22 | -26 | 22 | 50 |

| | | | | | |
|---|---|---|---|---|---|
| 32 | | | | | |
| | R Middle Frontal Gyrus | 3.26 | 38 | 24 | 52 |
| 32 | | | | | |
| | L Middle Frontal Gyrus | 3.16 | -38 | 18 | 56 |
| 32 | | | | | |
| | L Superior Parietal Lobule | 3.27 | -32 | -50 | 60 |
| 24 | | | | | |
| | R Brain-Stem | 2.88 | 14 | -22 | -24 |
| 24 | | | | | |
| | R Parahippocampal Gyrus, posterior division | 3.44 | 20 | -28 | -18 |
| 24 | | | | | |
| | L Temporal Occipital Fusiform Cortex | 3.24 | -34 | -58 | -16 |
| 24 | | | | | |
| | R Temporal Pole | 3.28 | 60 | 6 | -16 |
| 24 | | | | | |
| | R Frontal Pole | 3.16 | 20 | 44 | -16 |
| 24 | | | | | |
| | L Thalamus | 4.28 | -16 | -32 | -4 |
| 24 | | | | | |
| | R Temporal Pole | 3.7 | 60 | 12 | -6 |
| 24 | | | | | |
| | L Cingulate Gyrus, posterior division | 3.03 | -14 | -48 | 2 |
| 24 | | | | | |
| | R Frontal Pole | 2.91 | 36 | 54 | 4 |
| 24 | | | | | |
| | L Frontal Pole | 3.64 | -2 | 64 | 12 |
| 24 | | | | | |
| | R Inferior Frontal Gyrus, pars opercularis | 3.02 | 48 | 14 | 30 |
| 24 | | | | | |
| | L Superior Frontal Gyrus | 2.98 | -22 | 20 | 44 |
| 24 | | | | | |
| | L Superior Frontal Gyrus | 3.7 | -4 | 36 | 42 |
| 24 | | | | | |
| | L Superior Frontal Gyrus | 3.06 | -2 | 38 | 48 |
| 24 | | | | | |
| | L Superior Parietal Lobule | 3.23 | -18 | -48 | 64 |
| 24 | | | | | |
| | L Postcentral Gyrus | 3.46 | -32 | -38 | 64 |
| 16 | | | | | |
| | R Brain-Stem | 3.29 | 10 | -38 | -58 |
| 16 | | | | | |
| | L Frontal Medial Cortex | 2.94 | -6 | 48 | -26 |
| 16 | | | | | |
| | R Inferior Temporal Gyrus, temporooccipital part | 3.01 | 48 | -42 | -22 |

| | | | | | |
|---|---|---|---|---|---|
| 16 | | | | | |
| | R Temporal Pole | 3.3 | 48 | 20 | -22 |
| 16 | | | | | |
| | R Occipital Fusiform Gyrus | 2.88 | 20 | -86 | -18 |
| 16 | | | | | |
| | R Frontal Orbital Cortex | 3.35 | 38 | 22 | -16 |
| 16 | | | | | |
| | L Planum Polare | 3.04 | -44 | -6 | -14 |
| 16 | | | | | |
| | R Frontal Pole | 2.91 | 30 | 64 | -8 |
| 16 | | | | | |
| | L Middle Temporal Gyrus, temporooccipital part | 2.9 | -58 | -54 | -4 |
| 16 | | | | | |
| | R Thalamus | 2.95 | 16 | -20 | 0 |
| 16 | | | | | |
| | R Cingulate Gyrus, anterior division | 3.43 | 0 | 32 | 0 |
| 16 | | | | | |
| | L Lateral Occipital Cortex, inferior division | 2.83 | -46 | -84 | 10 |
| 16 | | | | | |
| | L Frontal Pole | 3.46 | -14 | 68 | 12 |
| 16 | | | | | |
| | R Central Opercular Cortex | 3.1 | 38 | -14 | 22 |
| 16 | | | | | |
| | L Caudate | 3.21 | -16 | -12 | 24 |
| 16 | | | | | |
| | L Middle Frontal Gyrus | 2.93 | -42 | 22 | 24 |
| 16 | | | | | |
| | L Frontal Pole | 2.95 | -20 | 54 | 32 |
| 16 | | | | | |
| | L Postcentral Gyrus | 3.4 | -56 | -14 | 36 |
| 16 | | | | | |
| | R Postcentral Gyrus | 2.97 | 60 | -12 | 38 |
| 16 | | | | | |
| | R Frontal Pole | 3.04 | 22 | 44 | 46 |
| 16 | | | | | |
| | R Superior Frontal Gyrus | 2.99 | 14 | 28 | 56 |
| 16 | | | | | |
| | R Lateral Occipital Cortex, superior division | 3.07 | 18 | -58 | 60 |
| 16 | | | | | |
| | R Superior Parietal Lobule | 3.03 | 26 | -54 | 62 |
| 16 | | | | | |
| | L Postcentral Gyrus | 3.17 | -12 | -40 | 70 |
| 16 | | | | | |
| | R Postcentral Gyrus | 3.15 | 6 | -32 | 80 |

| | | | | | |
|---|---|---|---|---|---|
| 8 | | | | | |
| | R Brain-Stem | 2.9 | 8 | -32 | -56 |
| 8 | | | | | |
| | R Brain-Stem | 2.98 | 4 | -30 | -54 |
| 8 | | | | | |
| | R Temporal Pole | 2.92 | 34 | 24 | -42 |
| 8 | | | | | |
| | R Inferior Temporal Gyrus, temporooccipital part | 2.84 | 54 | -46 | -24 |
| 8 | | | | | |
| | R Inferior Temporal Gyrus, temporooccipital part | 2.81 | 52 | -40 | -22 |
| 8 | | | | | |
| | R Parahippocampal Gyrus, posterior division | 2.91 | 26 | -28 | -22 |
| 8 | | | | | |
| | R Frontal Orbital Cortex | 3 | 12 | 16 | -22 |
| 8 | | | | | |
| | L Lateral Occipital Cortex, inferior division | 2.94 | -36 | -82 | -18 |
| 8 | | | | | |
| | L Parahippocampal Gyrus, posterior division | 2.8 | -16 | -30 | -18 |
| 8 | | | | | |
| | R Planum Polare | 2.86 | 42 | -2 | -16 |
| 8 | | | | | |
| | L Frontal Pole | 3.16 | -4 | 60 | -14 |
| 8 | | | | | |
| | L Frontal Pole | 3.17 | -12 | 60 | -14 |
| 8 | | | | | |
| | L Occipital Fusiform Gyrus | 3.39 | -32 | -68 | -12 |
| 8 | | | | | |
| | L Frontal Pole | 2.82 | -38 | 52 | -12 |
| 8 | | | | | |
| | L Paracingulate Gyrus | 2.88 | -12 | 44 | -4 |
| 8 | | | | | |
| | R Insular Cortex | 3.38 | 44 | 2 | -2 |
| 8 | | | | | |
| | L Superior Temporal Gyrus, anterior division | 2.92 | -62 | -2 | 2 |
| 8 | | | | | |
| | R Insular Cortex | 2.8 | 40 | -10 | 4 |
| 8 | | | | | |
| | L Insular Cortex | 2.81 | -38 | -8 | 4 |
| 8 | | | | | |
| | L Intracalcarine Cortex | 2.97 | -16 | -76 | 10 |
| 8 | | | | | |

| | | | | | |
|---|---|---|---|---|---|
| | R Central Opercular Cortex | 2.82 | 50 | -12 | 10 |
| 8 | | | | | |
| | L Intracalcarine Cortex | 3.08 | -2 | -66 | 12 |
| 8 | | | | | |
| | L Precuneus Cortex | 2.8 | -2 | -62 | 14 |
| 8 | | | | | |
| | L Central Opercular Cortex | 2.94 | -38 | -10 | 14 |
| 8 | | | | | |
| | L Postcentral Gyrus | 3.05 | -62 | -8 | 14 |
| 8 | | | | | |
| | R Central Opercular Cortex | 2.84 | 50 | -8 | 16 |
| 8 | | | | | |
| | R Lateral Occipital Cortex, superior division | 3.08 | 24 | -86 | 20 |
| 8 | | | | | |
| | L Angular Gyrus | 2.81 | -54 | -60 | 22 |
| 8 | | | | | |
| | L Lateral Occipital Cortex, superior division | 2.85 | -50 | -62 | 24 |
| 8 | | | | | |
| | L Inferior Frontal Gyrus, pars opercularis | 2.84 | -54 | 16 | 24 |
| 8 | | | | | |
| | L Inferior Frontal Gyrus, pars triangularis | 2.81 | -52 | 24 | 24 |
| 8 | | | | | |
| | L Cuneal Cortex | 2.93 | -10 | -80 | 26 |
| 8 | | | | | |
| | R Superior Frontal Gyrus | 2.84 | 6 | 56 | 26 |
| 8 | | | | | |
| | R Cuneal Cortex | 2.95 | 14 | -76 | 30 |
| 8 | | | | | |
| | L Frontal Pole | 3.18 | -20 | 54 | 38 |
| 8 | | | | | |
| | L Superior Frontal Gyrus | 2.83 | -18 | 36 | 44 |
| 8 | | | | | |
| | L Middle Frontal Gyrus | 2.83 | -28 | 28 | 48 |
| 8 | | | | | |
| | R Precuneus Cortex | 2.83 | 2 | -76 | 50 |
| 8 | | | | | |
| | L Postcentral Gyrus | 3.08 | -44 | -20 | 50 |
| 8 | | | | | |
| | L Precentral Gyrus | 2.88 | -36 | -20 | 52 |
| 8 | | | | | |
| | L Postcentral Gyrus | 2.81 | -28 | -36 | 54 |
| 8 | | | | | |
| | L Postcentral Gyrus | 2.98 | -42 | -24 | 54 |
| 8 | | | | | |

| | | | | | |
|---|---|---|---|---|---|
| | R Middle Frontal Gyrus | 3 | 46 | 14 | 54 |
| 8 | | | | | |
| | L Superior Frontal Gyrus | 2.9 | -14 | 30 | 56 |
| 8 | | | | | |
| | R Middle Frontal Gyrus | 2.96 | 38 | 12 | 58 |
| 8 | | | | | |
| | R Postcentral Gyrus | 2.82 | 28 | -32 | 68 |
| 8 | | | | | |
| | R Postcentral Gyrus | 2.86 | 24 | -32 | 70 |

**Supplementary Table S20. [$UT_{onset}$ > $US_{onset}$].** Descriptive statistics for clusters and local maxima showing greater activation for Uncertain-Threat relative to Uncertain-Safety anticipation for the OSP Onset regressor (FDR *q*<0.05, whole-brain corrected).

| *mm³* | Label | *t* | *x* | *y* | *z* |
|---|---|---|---|---|---|
| 286,080 | | | | | |
| | L Frontal Pole | 6.60 | -6 | 68 | -10 |
| | R Frontal Pole | 6.10 | 2 | 66 | -10 |
| | L Cingulate Gyrus, anterior division | 3.96 | -4 | 44 | 4 |
| | R Superior Frontal Gyrus | 7.45 | 4 | 44 | 44 |
| | L Frontal Medial Cortex | 4.41 | -4 | 38 | -22 |
| | R Frontal Medial Cortex | 5.02 | 6 | 36 | -18 |
| | R Inferior Frontal Gyrus, pars triangularis | 6.74 | 56 | 34 | 10 |
| | L Inferior Frontal Gyrus, pars triangularis | 3.19 | -40 | 26 | 20 |
| | L Frontal Operculum Cortex | 4.67 | -34 | 24 | 6 |
| | R Frontal Operculum Cortex | 4.17 | 36 | 24 | 10 |
| | R Insular Cortex | 7.37 | 32 | 22 | -6 |
| | L Insular Cortex | 3.86 | -36 | 18 | -6 |
| | L Frontal Orbital Cortex | 7.73 | -24 | 16 | -20 |
| | L Temporal Pole | 6.00 | -36 | 16 | -32 |
| | L Subcallosal Cortex | 4.78 | -4 | 16 | -8 |
| | R Frontal Orbital Cortex | 9.05 | 22 | 16 | -20 |
| | L Paracingulate Gyrus | 5.48 | -4 | 12 | 52 |
| | L Inferior Frontal Gyrus, pars opercularis | 5.22 | -50 | 12 | 28 |
| | R Subcallosal Cortex | 5.26 | 12 | 12 | -16 |
| | L Caudate | 4.87 | -6 | 10 | 0 |
| | L Middle Frontal Gyrus | 4.70 | -44 | 10 | 36 |
| | R Inferior Frontal Gyrus, pars opercularis | 7.41 | 44 | 10 | 24 |
| | R Paracingulate Gyrus | 4.69 | 6 | 10 | 52 |
| | R Caudate | 5.44 | 10 | 8 | 8 |
| | R Temporal Pole | 8.97 | 54 | 6 | -18 |
| | R Juxtapositional Lobule Cortex (formerly Supplementary Motor Cortex) | 7.88 | 6 | 6 | 66 |
| | R Middle Frontal Gyrus | 7.84 | 42 | 6 | 38 |
| | L Juxtapositional Lobule Cortex (formerly Supplementary Motor Cortex) | 6.72 | -4 | 4 | 66 |
| | L Superior Frontal Gyrus | 5.46 | -12 | 2 | 70 |
| | R Precentral Gyrus | 6.98 | 50 | 2 | 50 |
| | L Precentral Gyrus | 6.82 | -46 | 0 | 48 |
| | L Amygdala | 4.25 | -24 | 0 | -26 |
| | L Inferior Temporal Gyrus, anterior division | 4.25 | -42 | 0 | -38 |
| | R Superior Temporal Gyrus, anterior division | 6.24 | 58 | 0 | -10 |
| | R Inferior Temporal Gyrus, anterior division | 5.06 | 40 | 0 | -40 |

| | | | | | |
|---|---|---|---|---|---|
| | R Temporal Fusiform Cortex, anterior division | 3.98 | 36 | 0 | -44 |
| | L Middle Temporal Gyrus, anterior division | 4.34 | -52 | -2 | -18 |
| | L Temporal Fusiform Cortex, anterior division | 4.93 | -34 | -4 | -40 |
| | R Amygdala | 4.92 | 16 | -6 | -16 |
| | L Parahippocampal Gyrus, anterior division | 5.27 | -30 | -10 | -34 |
| | L Superior Temporal Gyrus, anterior division | 2.77 | -54 | -10 | -8 |
| | R Middle Temporal Gyrus, posterior division | 6.90 | 58 | -10 | -12 |
| | R Temporal Fusiform Cortex, posterior division | 5.53 | 34 | -10 | -42 |
| | R Parahippocampal Gyrus, anterior division | 4.84 | 30 | -10 | -32 |
| | R Inferior Temporal Gyrus, posterior division | 3.83 | 44 | -24 | -22 |
| | R Parahippocampal Gyrus, posterior division | 4.31 | 32 | -26 | -22 |
| | R Superior Temporal Gyrus, posterior division | 7.55 | 50 | -28 | 0 |
| | L Parahippocampal Gyrus, posterior division | 3.41 | -28 | -30 | -24 |
| | R Supramarginal Gyrus, anterior division | 2.90 | 54 | -32 | 48 |
| | R Postcentral Gyrus | 2.81 | 6 | -32 | 80 |
| | L Thalamus | 3.23 | -16 | -34 | 8 |
| | R Hippocampus | 8.40 | 28 | -34 | -4 |
| | R Thalamus | 4.50 | 14 | -34 | 4 |
| | L Hippocampus | 7.24 | -24 | -38 | 0 |
| | R Supramarginal Gyrus, posterior division | 4.45 | 42 | -40 | 40 |
| | L Temporal Fusiform Cortex, posterior division | 4.18 | -40 | -42 | -20 |
| | L Inferior Temporal Gyrus, posterior division | 3.76 | -46 | -42 | -18 |
| | R Cingulate Gyrus, posterior division | 6.25 | 10 | -42 | 4 |
| | L Cingulate Gyrus, posterior division | 5.86 | -6 | -44 | 10 |
| | R Superior Parietal Lobule | 5.97 | 34 | -44 | 48 |
| | L Inferior Temporal Gyrus, temporooccipital part | 5.08 | -52 | -50 | -20 |
| | R Inferior Temporal Gyrus, temporooccipital part | 9.31 | 50 | -50 | -22 |
| | R Precuneus Cortex | 6.87 | 2 | -54 | 60 |
| | R Angular Gyrus | 5.53 | 38 | -54 | 50 |
| | R Temporal Occipital Fusiform Cortex | 6.64 | 46 | -58 | -18 |
| | L Temporal Occipital Fusiform Cortex | 5.00 | -42 | -60 | -18 |

| | | | | | |
|---|---|---|---|---|---|
| | L Middle Temporal Gyrus, temporooccipital part | 4.68 | -50 | -60 | -2 |
| | R Middle Temporal Gyrus, temporooccipital part | 4.89 | 54 | -60 | -2 |
| | L Lateral Occipital Cortex, inferior division | 6.21 | -50 | -64 | -18 |
| | L Intracalcarine Cortex | 3.96 | -14 | -64 | 10 |
| | R Lateral Occipital Cortex, superior division | 7.48 | 32 | -64 | 32 |
| | L Lateral Occipital Cortex, superior division | 7.65 | -8 | -70 | 58 |
| | L Precuneus Cortex | 8.79 | -2 | -72 | 54 |
| | L Occipital Fusiform Gyrus | 9.52 | -20 | -82 | -14 |
| | R Lateral Occipital Cortex, inferior division | 7.82 | 46 | -82 | -8 |
| | L Cuneal Cortex | 3.64 | -4 | -84 | 18 |
| | L Lingual Gyrus | 9.21 | -8 | -86 | -12 |
| | R Occipital Fusiform Gyrus | 11.24 | 22 | -86 | -10 |
| | R Lingual Gyrus | 9.50 | 4 | -86 | -6 |
| | R Cuneal Cortex | 2.95 | 0 | -86 | 24 |
| | L Occipital Pole | 10.14 | -16 | -94 | -12 |
| | R Occipital Pole | 10.51 | 12 | -94 | -8 |
| 3,432 | | | | | |
| | L Supramarginal Gyrus, anterior division | 3.70 | -44 | -34 | 38 |
| | L Superior Parietal Lobule | 4.61 | -40 | -44 | 46 |
| | L Supramarginal Gyrus, posterior division | 5.08 | -36 | -48 | 40 |
| | L Lateral Occipital Cortex, superior division | 3.04 | -32 | -62 | 46 |
| 2,760 | | | | | |
| | R Precentral Gyrus | 5.07 | 40 | -22 | 66 |
| | R Postcentral Gyrus | 3.26 | 44 | -22 | 56 |
| 1,232 | | | | | |
| | L Thalamus | 3.20 | -2 | -22 | -2 |
| | R Thalamus | 5.55 | 4 | -22 | -2 |
| | R Brain-Stem | 4.99 | 6 | -30 | -6 |
| 1,056 | | | | | |
| | R Brain-Stem | 5.80 | 0 | -18 | -22 |
| | L Brain-Stem | 3.60 | -6 | -34 | -24 |
| 760 | | | | | |
| | L Cingulate Gyrus, anterior division | 4.81 | -4 | 6 | 28 |
| | R Cingulate Gyrus, anterior division | 4.58 | 0 | -4 | 36 |
| 704 | | | | | |
| | L Superior Temporal Gyrus, posterior division | 4.31 | -54 | -20 | -4 |
| 192 | | | | | |
| | R Cingulate Gyrus, anterior division | 4.14 | 4 | 12 | 26 |
| 168 | | | | | |
| | R Cingulate Gyrus, anterior division | 4.44 | 6 | 24 | 20 |

| | | | | | |
|---|---|---|---|---|---|
| 160 | | | | | |
| | R Brain-Stem | 3.32 | 4 | -20 | -40 |
| 136 | | | | | |
| | L Cingulate Gyrus, posterior division | 3.79 | -2 | -20 | 32 |
| 128 | | | | | |
| | R Supramarginal Gyrus, anterior division | 2.89 | 60 | -22 | 46 |
| 120 | | | | | |
| | L Frontal Pole | 2.86 | -26 | 66 | 8 |
| 112 | | | | | |
| | L Brain-Stem | 3.51 | -2 | -30 | -36 |
| | R Brain-Stem | 3.41 | 0 | -32 | -38 |
| 104 | | | | | |
| | R Cingulate Gyrus, posterior division | 3.23 | 6 | -24 | 28 |
| 104 | | | | | |
| | L Cingulate Gyrus, posterior division | 3.81 | -10 | -42 | 36 |
| 104 | | | | | |
| | L Precuneus Cortex | 2.57 | -12 | -44 | 40 |
| 96 | | | | | |
| | R Brain-Stem | 3.08 | 10 | -20 | -28 |
| 96 | | | | | |
| | L Brain-Stem | 4.81 | -8 | -28 | -12 |
| 96 | | | | | |
| | L Supramarginal Gyrus, anterior division | 3.01 | -66 | -36 | 34 |
| 88 | | | | | |
| | R Brain-Stem | 3.57 | 2 | -30 | -52 |
| 80 | | | | | |
| | L Angular Gyrus | 3.09 | -50 | -56 | 14 |
| 64 | | | | | |
| | R Brain-Stem | 3.19 | 2 | -22 | -34 |
| 64 | | | | | |
| | L Amygdala | 4.09 | -28 | 2 | -20 |
| 64 | | | | | |
| | R Amygdala | 3.43 | 30 | 2 | -18 |
| 64 | | | | | |
| | R Thalamus | 2.71 | 10 | -6 | 4 |
| 64 | | | | | |
| | R Supramarginal Gyrus, anterior division | 3.27 | 54 | -24 | 42 |
| 56 | | | | | |
| | R Brain-Stem | 2.67 | 10 | -32 | -32 |
| 56 | | | | | |
| | L Parahippocampal Gyrus, posterior division | 3.10 | -24 | -40 | -16 |
| 56 | | | | | |
| | L Temporal Fusiform Cortex, posterior division | 2.87 | -32 | -40 | -14 |
| 56 | | | | | |

| | | | | | |
|---|---|---|---|---|---|
| | L Inferior Frontal Gyrus, pars opercularis | 3.79 | -48 | 12 | 10 |
| 48 | | | | | |
| | L Inferior Temporal Gyrus, posterior division | 3.38 | -46 | -24 | -28 |
| 48 | | | | | |
| | R Frontal Orbital Cortex | 2.68 | 40 | 30 | -14 |
| 48 | | | | | |
| | L Middle Temporal Gyrus, posterior division | 2.92 | -62 | -22 | -10 |
| 48 | | | | | |
| | L Postcentral Gyrus | 3.01 | -64 | -22 | 36 |
| 48 | | | | | |
| | R Paracingulate Gyrus | 2.97 | 12 | 14 | 36 |
| 40 | | | | | |
| | L Brain-Stem | 2.69 | -4 | -38 | -40 |
| 40 | | | | | |
| | L Middle Frontal Gyrus | 2.79 | -38 | 34 | 20 |
| 40 | | | | | |
| | L Cingulate Gyrus, anterior division | 3.16 | -4 | 26 | 18 |
| 40 | | | | | |
| | L Cingulate Gyrus, anterior division | 3.05 | -2 | -14 | 30 |
| 32 | | | | | |
| | L Brain-Stem | 3.13 | -8 | -44 | -42 |
| 32 | | | | | |
| | R Brain-Stem | 2.46 | 0 | -16 | -38 |
| 32 | | | | | |
| | L Inferior Temporal Gyrus, posterior division | 3.11 | -60 | -24 | -26 |
| 32 | | | | | |
| | L Insular Cortex | 2.51 | -38 | -8 | -10 |
| 32 | | | | | |
| | L Inferior Frontal Gyrus, pars opercularis | 3.05 | -50 | 16 | 4 |
| 32 | | | | | |
| | L Supramarginal Gyrus, anterior division | 2.43 | -56 | -36 | 34 |
| 32 | | | | | |
| | L Precentral Gyrus | 3.05 | -30 | -12 | 66 |
| 24 | | | | | |
| | L Insular Cortex | 2.50 | -38 | 10 | -14 |
| 24 | | | | | |
| | L Lingual Gyrus | 3.14 | -26 | -42 | -8 |
| 24 | | | | | |
| | L Putamen | 3.31 | -18 | 18 | -8 |
| 24 | | | | | |
| | L Middle Temporal Gyrus, posterior division | 3.06 | -60 | -28 | -4 |
| 24 | | | | | |

| | | | | | |
|---|---|---|---|---|---|
| | R Pallidum | 3.51 | 16 | 6 | 2 |
| 24 | | | | | |
| | L Frontal Operculum Cortex | 2.36 | -40 | 12 | 6 |
| 24 | | | | | |
| | L Middle Temporal Gyrus, temporooccipital part | 2.47 | -52 | -58 | 8 |
| 24 | | | | | |
| | L Precuneus Cortex | 2.60 | -2 | -56 | 10 |
| 24 | | | | | |
| | L Paracingulate Gyrus | 2.47 | -10 | 10 | 38 |
| 24 | | | | | |
| | L Middle Frontal Gyrus | 2.99 | -26 | 30 | 40 |
| 24 | | | | | |
| | R Postcentral Gyrus | 3.00 | 48 | -28 | 56 |
| 16 | | | | | |
| | R Brain-Stem | 2.68 | 10 | -44 | -38 |
| 16 | | | | | |
| | R Brain-Stem | 2.70 | 12 | -28 | -38 |
| 16 | | | | | |
| | L Inferior Temporal Gyrus, posterior division | 2.71 | -60 | -36 | -28 |
| 16 | | | | | |
| | L Subcallosal Cortex | 3.16 | -4 | 18 | -24 |
| 16 | | | | | |
| | R Parahippocampal Gyrus, posterior division | 2.85 | 18 | -28 | -20 |
| 16 | | | | | |
| | L Planum Polare | 2.76 | -40 | -4 | -16 |
| 16 | | | | | |
| | L Insular Cortex | 2.69 | -32 | 14 | 6 |
| 16 | | | | | |
| | L Thalamus | 2.45 | -10 | -10 | 8 |
| 16 | | | | | |
| | R Thalamus | 2.61 | 8 | -30 | 10 |
| 16 | | | | | |
| | R Cingulate Gyrus, anterior division | 2.48 | 4 | 38 | 10 |
| 16 | | | | | |
| | L Lateral Occipital Cortex, superior division | 2.65 | -38 | -70 | 16 |
| 16 | | | | | |
| | L Thalamus | 2.57 | -10 | -26 | 16 |
| 16 | | | | | |
| | L Inferior Frontal Gyrus, pars opercularis | 2.50 | -52 | 14 | 16 |
| 16 | | | | | |
| | L Thalamus | 2.51 | -10 | -20 | 18 |
| 16 | | | | | |

| | | | | | |
|---|---|---|---|---|---|
| | L Middle Frontal Gyrus | 2.36 | -44 | 30 | 22 |
| 16 | | | | | |
| | R Cingulate Gyrus, anterior division | 2.66 | 8 | 16 | 34 |
| 16 | | | | | |
| | L Precentral Gyrus | 2.81 | -42 | -8 | 48 |
| 16 | | | | | |
| | R Precentral Gyrus | 2.58 | 6 | -22 | 78 |
| 8 | | | | | |
| | R Brain-Stem | 2.48 | 2 | -42 | -62 |
| 8 | | | | | |
| | R Brain-Stem | 2.66 | 0 | -36 | -44 |
| 8 | | | | | |
| | L Inferior Temporal Gyrus, anterior division | 2.75 | -54 | -6 | -42 |
| 8 | | | | | |
| | L Brain-Stem | 2.84 | -16 | -30 | -40 |
| 8 | | | | | |
| | L Brain-Stem | 2.47 | -10 | -42 | -38 |
| 8 | | | | | |
| | L Temporal Fusiform Cortex, posterior division | 2.31 | -38 | -20 | -34 |
| 8 | | | | | |
| | L Temporal Pole | 2.58 | -54 | 12 | -32 |
| 8 | | | | | |
| | R Brain-Stem | 2.50 | 6 | -42 | -30 |
| 8 | | | | | |
| | L Temporal Fusiform Cortex, posterior division | 2.48 | -40 | -32 | -30 |
| 8 | | | | | |
| | L Temporal Fusiform Cortex, posterior division | 2.44 | -38 | -28 | -30 |
| 8 | | | | | |
| | L Inferior Temporal Gyrus, posterior division | 2.55 | -52 | -24 | -28 |
| 8 | | | | | |
| | L Temporal Pole | 2.48 | -38 | 8 | -28 |
| 8 | | | | | |
| | R Frontal Orbital Cortex | 2.31 | 10 | 22 | -26 |
| 8 | | | | | |
| | L Para hippocampal Gyrus, posterior division | 2.43 | -22 | -28 | -24 |
| 8 | | | | | |
| | L Parahippocampal Gyrus, anterior division | 2.33 | -20 | -20 | -24 |
| 8 | | | | | |
| | L Frontal Orbital Cortex | 2.33 | -36 | 20 | -24 |

| | | | | | |
|---|---|---|---|---|---|
| 8 | | | | | |
| | L Parahippocampal Gyrus, posterior division | 2.75 | -16 | -28 | -20 |
| 8 | | | | | |
| | L Brain-Stem | 2.60 | -2 | -32 | -18 |
| 8 | | | | | |
| | R Temporal Fusiform Cortex, posterior division | 2.77 | 40 | -30 | -18 |
| 8 | | | | | |
| | L Planum Polare | 2.74 | -40 | 0 | -18 |
| 8 | | | | | |
| | L Parahippocampal Gyrus, posterior division | 2.37 | -20 | -28 | -14 |
| 8 | | | | | |
| | R Insular Cortex | 2.42 | 40 | 14 | -14 |
| 8 | | | | | |
| | L Middle Temporal Gyrus, posterior division | 2.35 | -56 | -16 | -10 |
| 8 | | | | | |
| | L Pallidum | 2.37 | -14 | 0 | -6 |
| 8 | | | | | |
| | L Frontal Orbital Cortex | 2.41 | -44 | 34 | -6 |
| 8 | | | | | |
| | L Paracingulate Gyrus | 2.65 | -2 | 46 | -6 |
| 8 | | | | | |
| | L Cingulate Gyrus, anterior division | 2.40 | -8 | 38 | -2 |
| 8 | | | | | |
| | R Frontal Operculum Cortex | 2.54 | 48 | 12 | 4 |
| 8 | | | | | |
| | R Lateral Occipital Cortex, inferior division | 2.64 | 42 | -74 | 6 |
| 8 | | | | | |
| | L Caudate | 2.66 | -8 | 16 | 6 |
| 8 | | | | | |
| | R Cingulate Gyrus, anterior division | 2.42 | 6 | 40 | 6 |
| 8 | | | | | |
| | R Frontal Pole | 2.35 | 10 | 56 | 8 |
| 8 | | | | | |
| | R Thalamus | 2.55 | 4 | -4 | 10 |
| 8 | | | | | |
| | L Supramarginal Gyrus, posterior division | 2.44 | -50 | -48 | 14 |
| 8 | | | | | |
| | L Thalamus | 2.44 | -2 | -14 | 14 |
| 8 | | | | | |
| | L Frontal Pole | 2.46 | -8 | 62 | 14 |
| 8 | | | | | |
| | L Thalamus | 2.44 | -6 | -16 | 18 |

| | | | | | |
|---|---|---|---|---|---|
| 8 | | | | | |
| | L Lateral Occipital Cortex, superior division | 2.55 | -30 | -80 | 20 |
| 8 | | | | | |
| | L Cingulate Gyrus, posterior division | 2.33 | -4 | -42 | 22 |
| 8 | | | | | |
| | R Precuneus Cortex | 2.46 | 8 | -54 | 30 |
| 8 | | | | | |
| | R Supramarginal Gyrus, posterior division | 2.40 | 66 | -44 | 32 |
| 8 | | | | | |
| | L Supramarginal Gyrus, anterior division | 2.44 | -54 | -24 | 34 |
| 8 | | | | | |
| | R Paracingulate Gyrus | 2.39 | 12 | 20 | 34 |
| 8 | | | | | |
| | L Middle Frontal Gyrus | 2.41 | -34 | 34 | 34 |
| 8 | | | | | |
| | L Middle Frontal Gyrus | 2.30 | -36 | 36 | 36 |
| 8 | | | | | |
| | R Superior Frontal Gyrus | 2.30 | 20 | -2 | 58 |
| 8 | | | | | |
| | R Postcentral Gyrus | 2.41 | 38 | -32 | 60 |
| 8 | | | | | |
| | L Superior Frontal Gyrus | 2.43 | -4 | 24 | 62 |

**Supplementary Table S21. [$CT_{onset} > CS_{onset}$].** Descriptive statistics for clusters and local maxima showing greater activation for Certain-Threat relative to Certain-Safety anticipation for the OSP Onset regressor (FDR *q*<0.05, whole-brain corrected).

| ***mm³*** | **Label** | ***t*** | ***x*** | ***y*** | ***z*** |
|---|---|---|---|---|---|
| 33,112 | | | | | |
| | R Inferior Temporal Gyrus, temporooccipital part | 5.58 | 50 | -54 | -18 |
| | R Middle Temporal Gyrus, temporooccipital part | 4.37 | 58 | -56 | -8 |
| | R Temporal Occipital Fusiform Cortex | 3.47 | 44 | -56 | -18 |
| | R Lateral Occipital Cortex, inferior division | 6.50 | 48 | -74 | -8 |
| | L Lingual Gyrus | 6.55 | -4 | -86 | -8 |
| | L Occipital Fusiform Gyrus | 3.54 | -24 | -86 | -18 |
| | R Lingual Gyrus | 6.75 | 8 | -88 | -8 |
| | R Occipital Pole | 8.52 | 14 | -102 | 12 |
| | L Occipital Pole | 9.29 | -12 | -104 | 2 |
| 3,440 | | | | | |
| | L Supramarginal Gyrus, posterior division | 5.87 | -36 | -50 | 42 |
| | L Angular Gyrus | 3.75 | -52 | -56 | 34 |
| | L Lateral Occipital Cortex, superior division | 4.63 | -48 | -64 | 46 |
| 3,384 | | | | | |
| | R Frontal Pole | 3.68 | 10 | 60 | 24 |
| | R Superior Frontal Gyrus | 6.03 | 4 | 48 | 40 |
| | L Paracingulate Gyrus | 4.13 | -4 | 36 | 38 |
| | R Paracingulate Gyrus | 4.13 | 6 | 36 | 40 |
| | L Superior Frontal Gyrus | 4.69 | -2 | 34 | 48 |
| 2,888 | | | | | |
| | L Frontal Pole | 5.43 | -30 | 66 | 2 |
| 2,520 | | | | | |
| | L Precuneus Cortex | 5.13 | -2 | -72 | 52 |
| | R Precuneus Cortex | 4.43 | 0 | -74 | 40 |
| 1,728 | | | | | |
| | R Lateral Occipital Cortex, superior division | 5.36 | 32 | -70 | 34 |
| 1,072 | | | | | |
| | R Supramarginal Gyrus, posterior division | 4.09 | 38 | -40 | 38 |
| | R Superior Parietal Lobule | 5.66 | 38 | -50 | 48 |
| 1,032 | | | | | |
| | R Frontal Pole | 5.18 | 50 | 42 | 12 |
| | R Middle Frontal Gyrus | 4.00 | 46 | 32 | 20 |
| 800 | | | | | |
| | R Cingulate Gyrus, posterior division | 4.09 | 0 | -34 | 40 |
| | L Cingulate Gyrus, posterior division | 3.24 | -2 | -40 | 38 |
| | R Precuneus Cortex | 3.35 | 4 | -42 | 46 |
| 696 | | | | | |
| | L Frontal Pole | 4.09 | -46 | 40 | 16 |

| | | | | | |
|---|---|---|---|---|---|
| | L Middle Frontal Gyrus | 3.13 | -44 | 38 | 30 |
| 632 | | | | | |
| | L Middle Frontal Gyrus | 4.61 | -38 | 34 | 38 |
| 552 | | | | | |
| | R Inferior Frontal Gyrus, pars opercularis | 3.92 | 50 | 12 | 26 |
| | R Precentral Gyrus | 4.15 | 48 | 6 | 24 |
| 440 | | | | | |
| | R Frontal Pole | 3.98 | 32 | 36 | -12 |
| | R Frontal Orbital Cortex | 4.37 | 26 | 30 | -16 |
| 440 | | | | | |
| | R Frontal Pole | 4.19 | 32 | 54 | 8 |
| 416 | | | | | |
| | R Frontal Pole | 4.03 | 42 | 50 | 6 |
| 312 | | | | | |
| | L Paracingulate Gyrus | 4.07 | -8 | 44 | 16 |
| 272 | | | | | |
| | R Amygdala | 4.34 | 20 | -4 | -16 |
| | R Amygdala | 4.51 | 28 | -6 | -14 |
| | R Amygdala | 3.76 | 22 | -10 | -12 |
| 240 | | | | | |
| | L Paracingulate Gyrus | 4.36 | -6 | 40 | 24 |
| 216 | | | | | |
| | R Brain-Stem | 3.93 | 14 | -26 | -18 |
| 208 | | | | | |
| | R Cingulate Gyrus, posterior division | 3.80 | 0 | -34 | 30 |
| 184 | | | | | |
| | L Cingulate Gyrus, posterior division | 3.31 | -2 | -24 | 32 |
| | R Cingulate Gyrus, posterior division | 3.46 | 2 | -24 | 34 |
| 176 | | | | | |
| | L Frontal Orbital Cortex | 4.03 | -22 | 30 | -22 |
| 144 | | | | | |
| | R Precuneus Cortex | 5.00 | 16 | -60 | 28 |
| 136 | | | | | |
| | L Middle Temporal Gyrus, posterior division | 3.95 | -70 | -30 | -10 |
| 136 | | | | | |
| | R Middle Frontal Gyrus | 3.25 | 46 | 30 | 40 |
| 128 | | | | | |
| | R Frontal Pole | 4.01 | 8 | 72 | -4 |
| 120 | | | | | |
| | L Brain-Stem | 4.61 | -8 | -26 | -20 |
| 120 | | | | | |
| | R Brain-Stem | 3.39 | 0 | -36 | -4 |
| 120 | | | | | |
| | L Cingulate Gyrus, posterior division | 4.02 | -10 | -28 | 40 |
| 112 | | | | | |
| | L Frontal Pole | 2.96 | -22 | 50 | -14 |

| | | | | | |
|---|---|---|---|---|---|
| 104 | | | | | |
| | R Hippocampus | 4.44 | 26 | -38 | 0 |
| 104 | | | | | |
| | L Lingual Gyrus | 4.48 | -20 | -54 | -2 |
| 88 | | | | | |
| | R Frontal Pole | 4.25 | 46 | 54 | -4 |
| 80 | | | | | |
| | L Temporal Pole | 3.71 | -36 | 4 | -18 |
| 80 | | | | | |
| | L Amygdala | 4.08 | -20 | -10 | -14 |
| | L Hippocampus/Amygdala | 3.15 | -22 | -12 | -16 |
| 80 | | | | | |
| | R Superior Temporal Gyrus, posterior division | 3.50 | 52 | -18 | -6 |
| 72 | | | | | |
| | R Inferior Temporal Gyrus, temporooccipital part | 3.67 | 58 | -42 | -22 |
| 72 | | | | | |
| | L Insular Cortex | 3.20 | -38 | -6 | -8 |
| | L Planum Polare | 4.35 | -42 | -10 | -10 |
| 72 | | | | | |
| | R Frontal Medial Cortex | 3.47 | 2 | 48 | -10 |
| 72 | | | | | |
| | L Planum Polare | 4.54 | -52 | 2 | -6 |
| 72 | | | | | |
| | R Caudate | 4.08 | 8 | 12 | 10 |
| 72 | | | | | |
| | L Frontal Pole | 3.60 | -44 | 48 | 16 |
| 64 | | | | | |
| | R Parahippocampal Gyrus, anterior division | 3.04 | 18 | -2 | -38 |
| 64 | | | | | |
| | R Temporal Pole | 3.67 | 40 | 10 | -26 |
| 64 | | | | | |
| | R Hippocampus | 3.29 | 34 | -14 | -16 |
| 64 | | | | | |
| | R Frontal Pole | 3.87 | 6 | 66 | -16 |
| 64 | | | | | |
| | R Hippocampus | 3.46 | 28 | -24 | -12 |
| 64 | | | | | |
| | R Insular Cortex | 3.59 | 38 | -2 | -10 |
| 64 | | | | | |
| | R Insular Cortex | 3.73 | 38 | -6 | 4 |
| 56 | | | | | |
| | R Brain-Stem | 3.58 | 18 | -34 | -42 |
| 56 | | | | | |
| | L Subcallosal Cortex | 3.24 | -4 | 10 | -8 |

| | | | | | |
|---|---|---|---|---|---|
| 56 | | | | | |
| | L Paracingulate Gyrus | 3.35 | -10 | 48 | 8 |
| 56 | | | | | |
| | R Middle Frontal Gyrus | 3.43 | 30 | 30 | 36 |
| 48 | | | | | |
| | L Temporal Pole | 3.52 | -54 | 2 | -34 |
| 48 | | | | | |
| | L Frontal Medial Cortex | 3.65 | -8 | 34 | -26 |
| 48 | | | | | |
| | L Amygdala | 3.31 | -30 | -8 | -16 |
| | L Amygdala | 3.32 | -28 | -10 | -14 |
| 48 | | | | | |
| | L Frontal Pole | 3.59 | -50 | 40 | -14 |
| 48 | | | | | |
| | L Insular Cortex | 3.78 | -40 | 0 | -10 |
| 48 | | | | | |
| | R Insular Cortex | 3.51 | 34 | 22 | -4 |
| 48 | | | | | |
| | R Frontal Pole | 3.69 | 40 | 62 | 6 |
| 48 | | | | | |
| | R Insular Cortex | 3.20 | 36 | -12 | 16 |
| 48 | | | | | |
| | R Precuneus Cortex | 3.27 | 2 | -52 | 18 |
| | R Cingulate Gyrus, posterior division | 2.95 | 2 | -52 | 24 |
| 48 | | | | | |
| | R Precentral Gyrus | 3.37 | 44 | 4 | 34 |
| 48 | | | | | |
| | R Supramarginal Gyrus, anterior division | 3.14 | 54 | -32 | 48 |
| | R Supramarginal Gyrus, posterior division | 3.21 | 56 | -36 | 54 |
| 40 | | | | | |
| | L Temporal Pole | 3.32 | -30 | 4 | -28 |
| 40 | | | | | |
| | R Frontal Orbital Cortex | 3.32 | 10 | 28 | -26 |
| 40 | | | | | |
| | R Frontal Orbital Cortex | 3.16 | 20 | 28 | -16 |
| 40 | | | | | |
| | R Frontal Orbital Cortex | 3.36 | 26 | 18 | -18 |
| 40 | | | | | |
| | R Paracingulate Gyrus | 3.20 | 6 | 42 | -6 |
| 40 | | | | | |
| | R Lingual Gyrus | 3.33 | 20 | -52 | 0 |
| 40 | | | | | |
| | L Hippocampus | 3.31 | -18 | -38 | 0 |
| 40 | | | | | |
| | R Thalamus | 3.26 | 18 | -36 | 2 |
| 40 | | | | | |
| | R Frontal Pole | 3.10 | 28 | 66 | 0 |

| | | | | | |
|---|---|---|---|---|---|
| 40 | | | | | |
| | R Lateral Occipital Cortex, superior division | 3.71 | 42 | -70 | 26 |
| 40 | | | | | |
| | L Lateral Occipital Cortex, superior division | 3.27 | -44 | -64 | 54 |
| 32 | | | | | |
| | L Brain-Stem | 3.22 | -4 | -42 | -50 |
| 32 | | | | | |
| | L Temporal Fusiform Cortex, anterior division | 3.95 | -30 | -8 | -44 |
| 32 | | | | | |
| | R Brain-Stem | 3.49 | 12 | -28 | -34 |
| 32 | | | | | |
| | R Temporal Pole | 3.21 | 34 | 18 | -34 |
| 32 | | | | | |
| | L Inferior Temporal Gyrus, posterior division | 3.58 | -52 | -18 | -32 |
| 32 | | | | | |
| | L Inferior Temporal Gyrus, temporooccipital part | 3.05 | -62 | -50 | -18 |
| 32 | | | | | |
| | L Frontal Medial Cortex | 3.29 | -4 | 36 | -18 |
| 32 | | | | | |
| | R Thalamus | 3.62 | 6 | -26 | 2 |
| 32 | | | | | |
| | R Frontal Pole | 3.23 | 50 | 46 | 4 |
| 32 | | | | | |
| | L Planum Temporale | 3.12 | -56 | -20 | 4 |
| 32 | | | | | |
| | L Cingulate Gyrus, posterior division | 3.64 | -6 | -46 | 10 |
| 32 | | | | | |
| | R Frontal Pole | 2.89 | 2 | 68 | 10 |
| 32 | | | | | |
| | L Frontal Pole | 3.25 | -40 | 58 | 12 |
| 32 | | | | | |
| | L Occipital Pole | 3.43 | -2 | -96 | 24 |
| 32 | | | | | |
| | R Occipital Pole | 3.58 | 2 | -92 | 26 |
| 32 | | | | | |
| | R Supramarginal Gyrus, posterior division | 3.51 | 60 | -42 | 36 |
| 32 | | | | | |
| | L Lateral Occipital Cortex, superior division | 3.28 | -34 | -62 | 40 |
| 32 | | | | | |
| | R Middle Frontal Gyrus | 3.49 | 34 | 6 | 56 |
| 32 | | | | | |

| | | | | | |
|---|---|---|---|---|---|
| | R Lateral Occipital Cortex, superior division | 3.42 | 36 | -64 | 60 |
| 24 | | | | | |
| | R Brain-Stem | 3.44 | 0 | -44 | -64 |
| 24 | | | | | |
| | R Temporal Fusiform Cortex, anterior division | 2.97 | 30 | -6 | -42 |
| 24 | | | | | |
| | R Temporal Pole | 3.36 | 54 | 12 | -34 |
| 24 | | | | | |
| | L Hippocampus | 3.16 | -30 | -10 | -24 |
| 24 | | | | | |
| | R Temporal Pole | 3.37 | 28 | 6 | -24 |
| 24 | | | | | |
| | L Temporal Pole | 3.57 | -44 | 10 | -22 |
| 24 | | | | | |
| | R Frontal Orbital Cortex | 2.95 | 26 | 28 | -24 |
| 24 | | | | | |
| | R Brain-Stem | 3.05 | 0 | -20 | -22 |
| 24 | | | | | |
| | L Hippocampus | 3.32 | -20 | -12 | -22 |
| 24 | | | | | |
| | R Frontal Medial Cortex | 3.18 | 0 | 48 | -20 |
| 24 | | | | | |
| | R Frontal Orbital Cortex | 3.03 | 34 | 30 | -18 |
| 24 | | | | | |
| | L Brain-Stem | 3.17 | -4 | -34 | -12 |
| 24 | | | | | |
| | R Frontal Pole | 3.60 | 26 | 42 | -14 |
| 24 | | | | | |
| | L Hippocampus | 3.12 | -28 | -36 | -8 |
| 24 | | | | | |
| | R Middle Temporal Gyrus, posterior division | 3.05 | 68 | -16 | -6 |
| 24 | | | | | |
| | R Planum Polare | 3.48 | 48 | -8 | -6 |
| 24 | | | | | |
| | R Frontal Pole | 3.46 | 28 | 56 | 0 |
| 24 | | | | | |
| | R Insular Cortex | 2.89 | 40 | -14 | 4 |
| 24 | | | | | |
| | R Thalamus | 3.04 | 6 | -16 | 12 |
| 24 | | | | | |
| | L Central Opercular Cortex | 3.23 | -58 | -8 | 12 |
| 24 | | | | | |
| | R Angular Gyrus | 3.10 | 52 | -44 | 32 |
| 24 | | | | | |

| | | | | | |
|---|---|---|---|---|---|
| | R Cingulate Gyrus, anterior division | 3.25 | 2 | -4 | 34 |
| 24 | | | | | |
| | R Middle Frontal Gyrus | 3.24 | 28 | 32 | 42 |
| 24 | | | | | |
| | R Cingulate Gyrus, posterior division | 3.45 | 6 | -22 | 42 |
| 16 | | | | | |
| | R Brain-Stem | 3.04 | 0 | -34 | -42 |
| 16 | | | | | |
| | R Temporal Pole | 3.03 | 42 | 8 | -38 |
| 16 | | | | | |
| | L Temporal Pole | 2.98 | -30 | 12 | -40 |
| 16 | | | | | |
| | L Inferior Temporal Gyrus, posterior division | 3.15 | -64 | -20 | -28 |
| 16 | | | | | |
| | R Temporal Pole | 2.95 | 30 | 10 | -30 |
| 16 | | | | | |
| | L Brain-Stem | 2.96 | -2 | -28 | -26 |
| 16 | | | | | |
| | L Frontal Pole | 3.05 | -8 | 44 | -24 |
| 16 | | | | | |
| | R Hippocampus | 2.96 | 28 | -18 | -20 |
| 16 | | | | | |
| | R Frontal Orbital Cortex | 2.93 | 14 | 22 | -18 |
| 16 | | | | | |
| | L Amygdala | 3.19 | -18 | -6 | -16 |
| 16 | | | | | |
| | R Insular Cortex | 2.94 | 36 | 6 | -16 |
| 16 | | | | | |
| | R Frontal Medial Cortex | 2.87 | 0 | 38 | -16 |
| 16 | | | | | |
| | L Frontal Pole | 3.01 | -24 | 42 | -16 |
| 16 | | | | | |
| | L Frontal Pole | 3.40 | -32 | 42 | -16 |
| 16 | | | | | |
| | L Hippocampus | 3.00 | -34 | -22 | -14 |
| 16 | | | | | |
| | L Hippocampus | 3.00 | -28 | -28 | -12 |
| 16 | | | | | |
| | L Frontal Pole | 3.10 | -22 | 44 | -12 |
| 16 | | | | | |
| | L Frontal Pole | 3.17 | -14 | 64 | -4 |
| 16 | | | | | |
| | R Superior Temporal Gyrus, posterior division | 2.92 | 62 | -22 | -2 |
| 16 | | | | | |
| | R Frontal Pole | 3.18 | 40 | 44 | -2 |

| | | | | | |
|---|---|---|---|---|---|
| 16 | | | | | |
| | R Planum Temporale | 3.03 | 58 | -12 | 6 |
| 16 | | | | | |
| | R Thalamus | 2.90 | 4 | -6 | 10 |
| 16 | | | | | |
| | L Insular Cortex | 3.66 | -38 | -6 | 12 |
| 16 | | | | | |
| | L Thalamus | 4.50 | -4 | -8 | 12 |
| 16 | | | | | |
| | R Central Opercular Cortex | 3.77 | 54 | -10 | 14 |
| 16 | | | | | |
| | R Paracingulate Gyrus | 3.01 | 10 | 46 | 14 |
| 16 | | | | | |
| | R Frontal Pole | 3.18 | 28 | 54 | 14 |
| 16 | | | | | |
| | L Frontal Pole | 2.99 | -22 | 54 | 16 |
| 16 | | | | | |
| | R Occipital Pole | 3.02 | 4 | -98 | 18 |
| 16 | | | | | |
| | L Precuneus Cortex | 2.96 | -16 | -64 | 28 |
| 16 | | | | | |
| | R Cingulate Gyrus, anterior division | 3.05 | 0 | -12 | 30 |
| 16 | | | | | |
| | L Paracingulate Gyrus | 3.31 | -8 | 34 | 30 |
| 16 | | | | | |
| | R Cuneal Cortex | 3.64 | 2 | -88 | 34 |
| 16 | | | | | |
| | L Precuneus Cortex | 3.02 | -6 | -70 | 34 |
| 16 | | | | | |
| | R Middle Frontal Gyrus | 3.94 | 38 | 14 | 36 |
| 16 | | | | | |
| | R Precentral Gyrus | 3.30 | 52 | -4 | 42 |
| 16 | | | | | |
| | R Angular Gyrus | 2.94 | 54 | -56 | 50 |
| 8 | | | | | |
| | L Brain-Stem | 2.92 | -6 | -48 | -50 |
| 8 | | | | | |
| | R Brain-Stem | 2.88 | 10 | -42 | -46 |
| 8 | | | | | |
| | L Temporal Pole | 2.93 | -36 | 14 | -44 |
| 8 | | | | | |
| | R Temporal Pole | 3.03 | 28 | 10 | -42 |
| 8 | | | | | |
| | L Brain-Stem | 2.90 | -4 | -20 | -36 |
| 8 | | | | | |
| | L Parahippocampal Gyrus, anterior division | 2.95 | -22 | -8 | -36 |

| | | | | | |
|---|---|---|---|---|---|
| 8 | | | | | |
| | L Parahippocampal Gyrus, anterior division | 2.90 | -22 | 2 | -36 |
| 8 | | | | | |
| | R Temporal Pole | 2.88 | 42 | 20 | -36 |
| 8 | | | | | |
| | L Parahippocampal Gyrus, anterior division | 3.05 | -24 | -4 | -34 |
| 8 | | | | | |
| | L Parahippocampal Gyrus, anterior division | 2.89 | -28 | -4 | -34 |
| 8 | | | | | |
| | R Parahippocampal Gyrus, anterior division | 2.87 | 26 | -2 | -34 |
| 8 | | | | | |
| | R Brain-Stem | 2.85 | 0 | -16 | -32 |
| 8 | | | | | |
| | R Temporal Pole | 2.99 | 28 | 16 | -32 |
| 8 | | | | | |
| | R Temporal Pole | 3.06 | 42 | 12 | -30 |
| 8 | | | | | |
| | R Brain-Stem | 2.91 | 18 | -22 | -28 |
| 8 | | | | | |
| | R Amygdala | 2.93 | 26 | 2 | -28 |
| | R Parahippocampal Gyrus, anterior division | 2.93 | 26 | 2 | -28 |
| 8 | | | | | |
| | R Parahippocampal Gyrus, posterior division | 2.86 | 22 | -24 | -26 |
| 8 | | | | | |
| | R Temporal Fusiform Cortex, posterior division | 2.90 | 38 | -14 | -26 |
| 8 | | | | | |
| | R Temporal Pole | 3.04 | 34 | 6 | -26 |
| 8 | | | | | |
| | R Subcallosal Cortex | 3.10 | 2 | 30 | -24 |
| 8 | | | | | |
| | L Frontal Pole | 2.89 | -6 | 60 | -24 |
| 8 | | | | | |
| | L Middle Temporal Gyrus, posterior division | 3.01 | -64 | -14 | -22 |
| 8 | | | | | |
| | L Frontal Orbital Cortex | 2.89 | -12 | 28 | -22 |
| 8 | | | | | |
| | L Frontal Medial Cortex | 2.88 | -10 | 34 | -22 |
| 8 | | | | | |

| | | | | | |
|---|---|---|---|---|---|
| | L Inferior Temporal Gyrus, posterior division | 3.30 | -56 | -42 | -20 |
| 8 | | | | | |
| | L Amygdala | 2.94 | -32 | 2 | -20 |
| 8 | | | | | |
| | R Amygdala | 3.01 | 26 | 0 | -18 |
| 8 | | | | | |
| | L Frontal Orbital Cortex | 2.94 | -30 | 14 | -18 |
| 8 | | | | | |
| | R Temporal Pole | 2.98 | 52 | 12 | -16 |
| 8 | | | | | |
| | L Hippocampus | 3.15 | -34 | -26 | -14 |
| 8 | | | | | |
| | L Frontal Pole | 2.92 | -28 | 58 | -14 |
| 8 | | | | | |
| | R Middle Temporal Gyrus, posterior division | 2.89 | 64 | -26 | -12 |
| 8 | | | | | |
| | R Frontal Pole | 2.95 | 38 | 56 | -10 |
| 8 | | | | | |
| | R Brain-Stem | 3.20 | 10 | -26 | -8 |
| 8 | | | | | |
| | L Middle Temporal Gyrus, posterior division | 2.86 | -70 | -24 | -6 |
| 8 | | | | | |
| | R Pallidum | 2.86 | 18 | 2 | -6 |
| 8 | | | | | |
| | R Planum Polare | 3.26 | 44 | -16 | -4 |
| 8 | | | | | |
| | R Superior Temporal Gyrus, anterior division | 2.96 | 64 | 0 | -4 |
| 8 | | | | | |
| | R Putamen | 2.90 | 28 | 14 | -2 |
| 8 | | | | | |
| | R Frontal Pole | 2.85 | 22 | 64 | -2 |
| 8 | | | | | |
| | L Hippocampus | 3.17 | -24 | -36 | 0 |
| 8 | | | | | |
| | L Thalamus | 2.98 | -2 | -4 | 0 |
| 8 | | | | | |
| | R Superior Temporal Gyrus, anterior division | 2.93 | 64 | 0 | 0 |
| 8 | | | | | |
| | R Insular Cortex | 2.92 | 34 | 22 | 0 |
| 8 | | | | | |
| | L Insular Cortex | 2.85 | -28 | 26 | 0 |
| 8 | | | | | |

| | | | | | |
|---|---|---|---|---|---|
| | R Frontal Pole | 2.92 | 22 | 58 | 0 |
| 8 | | | | | |
| | R Frontal Pole | 2.85 | 22 | 70 | 0 |
| 8 | | | | | |
| | L Frontal Pole | 2.93 | -22 | 68 | 4 |
| 8 | | | | | |
| | R Cingulate Gyrus, posterior division | 2.94 | 8 | -42 | 6 |
| 8 | | | | | |
| | R Thalamus | 2.95 | 10 | -24 | 6 |
| 8 | | | | | |
| | L Thalamus | 2.97 | -10 | -32 | 10 |
| 8 | | | | | |
| | L Planum Temporale | 3.19 | -44 | -30 | 10 |
| 8 | | | | | |
| | R Frontal Pole | 2.86 | 36 | 60 | 10 |
| 8 | | | | | |
| | R Cingulate Gyrus, posterior division | 2.94 | 6 | -44 | 12 |
| 8 | | | | | |
| | L Central Opercular Cortex | 2.97 | -52 | -14 | 12 |
| 8 | | | | | |
| | L Planum Temporale | 2.88 | -48 | -36 | 14 |
| 8 | | | | | |
| | R Thalamus | 2.95 | 16 | -22 | 14 |
| 8 | | | | | |
| | R Cingulate Gyrus, posterior division | 2.92 | 4 | -48 | 16 |
| 8 | | | | | |
| | R Central Opercular Cortex | 2.86 | 42 | -12 | 16 |
| 8 | | | | | |
| | R Frontal Pole | 2.96 | 2 | 66 | 18 |
| 8 | | | | | |
| | L Cuneal Cortex | 3.37 | -2 | -86 | 20 |
| 8 | | | | | |
| | R Frontal Pole | 3.19 | 2 | 64 | 26 |
| 8 | | | | | |
| | R Cingulate Gyrus, posterior division | 3.09 | 8 | -34 | 28 |
| 8 | | | | | |
| | L Supramarginal Gyrus, anterior division | 2.86 | -56 | -32 | 28 |
| 8 | | | | | |
| | R Cingulate Gyrus, anterior division | 2.87 | 4 | 8 | 28 |
| 8 | | | | | |
| | R Middle Frontal Gyrus | 2.86 | 56 | 20 | 34 |
| 8 | | | | | |
| | L Cingulate Gyrus, posterior division | 2.90 | -8 | -42 | 40 |
| 8 | | | | | |
| | L Middle Frontal Gyrus | 3.46 | -44 | 22 | 40 |
| 8 | | | | | |
| | R Superior Parietal Lobule | 2.92 | 30 | -42 | 42 |

| | | | | | |
|---|---|---|---|---|---|
| 8 | | | | | |
| | R Precuneus Cortex | 3.16 | 12 | -48 | 46 |
| 8 | | | | | |
| | R Supramarginal Gyrus, posterior division | 2.91 | 48 | -36 | 46 |
| 8 | | | | | |
| | R Superior Frontal Gyrus | 2.96 | 22 | 10 | 54 |
| 8 | | | | | |
| | R Precuneus Cortex | 2.88 | 4 | -50 | 70 |

**Supplementary Table S22. [$UT_{onset}$ > $US_{onset}$] ∩ [$CT_{onset}$ > $CS_{onset}$].** Descriptive statistics for clusters and local maxima showing greater activation for Uncertain-Threat relative to Uncertain-Safety anticipation for the OSP Onset regressor AND greater activation for Certain-Threat relative to Certain-Safety anticipation for the OSP Onset regressor (FDR $q$<0.05, whole-brain corrected).

| *mm³* | Label | *t* | *x* | *y* | *z* |
|---|---|---|---|---|---|
| 28,256 | | | | | |
| | R Inferior Temporal Gyrus, temporooccipital part | 5.58 | 50 | -54 | -18 |
| | R Temporal Occipital Fusiform Cortex | 3.47 | 44 | -56 | -18 |
| | R Middle Temporal Gyrus, temporooccipital part | 3.52 | 50 | -58 | 6 |
| | R Occipital Fusiform Gyrus | 3.85 | 40 | -70 | -18 |
| | R Lateral Occipital Cortex, inferior division | 6.03 | 48 | -74 | -8 |
| | L Lingual Gyrus | 6.55 | -4 | -86 | -8 |
| | L Occipital Fusiform Gyrus | 3.54 | -24 | -86 | -18 |
| | R Lingual Gyrus | 6.75 | 8 | -88 | -8 |
| | R Occipital Pole | 7.87 | 14 | -100 | 12 |
| | L Occipital Pole | 8.68 | -12 | -102 | 2 |
| 2,624 | | | | | |
| | R Superior Frontal Gyrus | 6.03 | 4 | 48 | 40 |
| | R Paracingulate Gyrus | 5.32 | 6 | 44 | 34 |
| | L Superior Frontal Gyrus | 4.08 | -6 | 38 | 36 |
| | L Paracingulate Gyrus | 4.13 | -4 | 36 | 38 |
| 2,472 | | | | | |
| | L Precuneus Cortex | 5.13 | -2 | -72 | 52 |
| | R Precuneus Cortex | 3.71 | 0 | -76 | 42 |
| 1,664 | | | | | |
| | R Lateral Occipital Cortex, superior division | 5.36 | 32 | -70 | 34 |
| 1,112 | | | | | |
| | L Supramarginal Gyrus, posterior division | 4.55 | -40 | -46 | 42 |
| 952 | | | | | |
| | R Supramarginal Gyrus, posterior division | 3.65 | 38 | -38 | 38 |
| | R Superior Parietal Lobule | 4.78 | 40 | -50 | 48 |
| 648 | | | | | |
| | R Frontal Pole | 4.32 | 54 | 38 | 16 |
| | R Middle Frontal Gyrus | 4 | 46 | 32 | 20 |
| 576 | | | | | |
| | R Cingulate Gyrus, posterior division | 3.39 | 8 | -40 | 34 |
| | R Precuneus Cortex | 3.63 | 6 | -46 | 40 |
| 552 | | | | | |
| | R Inferior Frontal Gyrus, pars opercular is | 3.92 | 50 | 12 | 26 |
| | R Precentral Gyrus | 4.15 | 48 | 6 | 24 |

| | | | | | |
|---|---|---|---|---|---|
| 352 | | | | | |
| | R Frontal Pole | 3.41 | 30 | 36 | -12 |
| | R Frontal Orbital Cortex | 4.01 | 26 | 30 | -16 |
| 168 | | | | | |
| | R Middle Temporal Gyrus, temporooccipital part | 3.11 | 62 | -42 | -12 |
| 160 | | | | | |
| | L Paracingulate Gyrus | 3.39 | -6 | 44 | 16 |
| 136 | | | | | |
| | R Precuneus Cortex | 4.26 | 16 | -60 | 28 |
| 104 | | | | | |
| | L Lingual Gyrus | 4.48 | -20 | -54 | -2 |
| 96 | | | | | |
| | R Hippocampus | 4.44 | 26 | -38 | 0 |
| 88 | | | | | |
| | R Frontal Pole | 3.67 | 4 | 72 | -6 |
| 80 | | | | | |
| | L Frontal Orbital Cortex | 3.95 | -24 | 30 | -20 |
| 80 | | | | | |
| | R Brain-Stem | 2.93 | 0 | -36 | -6 |
| 80 | | | | | |
| | L Cingulate Gyrus, posterior division | 3.63 | -6 | -28 | 42 |
| 72 | | | | | |
| | R Amygdala | 3.68 | 18 | -6 | -16 |
| 64 | | | | | |
| | R Frontal Pole | 3.49 | 6 | 68 | -16 |
| 64 | | | | | |
| | R Hippocampus | 3.46 | 28 | -24 | -12 |
| 56 | | | | | |
| | L Subcallosal Cortex | 3.24 | -4 | 10 | -8 |
| 56 | | | | | |
| | L Paracingulate Gyrus | 3.35 | -10 | 48 | 8 |
| 56 | | | | | |
| | L Paracingulate Gyrus | 3.02 | -8 | 42 | 24 |
| 48 | | | | | |
| | R Frontal Medial Cortex | 3.38 | 2 | 50 | -12 |
| 48 | | | | | |
| | R Insular Cortex | 3.51 | 34 | 22 | -4 |
| 48 | | | | | |
| | R Caudate | 3.21 | 10 | 12 | 8 |
| 48 | | | | | |
| | R Cingulate Gyrus, posterior division | 3.04 | 2 | -52 | 20 |

| | | | | | |
|---|---|---|---|---|---|
| 48 | | | | | |
| | R Cingulate Gyrus, posterior division | 3.08 | 0 | -40 | 26 |
| 48 | | | | | |
| | R Precentral Gyrus | 3.37 | 44 | 4 | 34 |
| 48 | | | | | |
| | R Cingulate Gyrus, posterior division | 3.23 | 4 | -32 | 34 |
| 40 | | | | | |
| | R Frontal Orbital Cortex | 3.32 | 10 | 28 | -26 |
| 40 | | | | | |
| | R Right Hippocampus | 3.28 | 34 | -14 | -18 |
| 40 | | | | | |
| | R Lingual Gyrus | 3.33 | 20 | -52 | 0 |
| 40 | | | | | |
| | L Left Hippocampus | 3.31 | -18 | -38 | 0 |
| 40 | | | | | |
| | R Lateral Occipital Cortex, superior division | 3.71 | 42 | -70 | 26 |
| 32 | | | | | |
| | L Temporal Fusiform Cortex, anterior division | 3.95 | -30 | -8 | -44 |
| 32 | | | | | |
| | R Temporal Pole | 3.21 | 34 | 18 | -34 |
| 32 | | | | | |
| | R Temporal Pole | 3.43 | 40 | 10 | -24 |
| 32 | | | | | |
| | L Brain-Stem | 2.79 | -6 | -24 | -22 |
| 32 | | | | | |
| | R Frontal Orbital Cortex | 2.9 | 20 | 26 | -20 |
| 32 | | | | | |
| | R Amygdala | 3.33 | 28 | -4 | -14 |
| 32 | | | | | |
| | R Thalamus | 3.26 | 18 | -36 | 2 |
| 32 | | | | | |
| | R Occipital Pole | 3.03 | 2 | -92 | 26 |
| 32 | | | | | |
| | L Cingulate Gyrus, posterior division | 3.02 | -2 | -22 | 32 |
| 24 | | | | | |
| | R Temporal Fusiform Cortex, anterior division | 2.97 | 30 | -6 | -42 |
| 24 | | | | | |
| | R Inferior Temporal Gyrus, temporooccipital part | 3.06 | 56 | -42 | -22 |
| 24 | | | | | |

| | | | | | |
|---|---|---|---|---|---|
| | R Brain-Stem | 3.05 | 0 | -20 | -22 |
| 24 | | | | | |
| | R Frontal Orbital Cortex | 3.15 | 24 | 18 | -18 |
| 24 | | | | | |
| | L Hippocampus | 3 | -28 | -36 | -6 |
| 24 | | | | | |
| | L Cingulate Gyrus, posterior division | 3.64 | -6 | -46 | 10 |
| 24 | | | | | |
| | L Paracingulate Gyrus | 2.71 | -4 | 40 | 28 |
| 24 | | | | | |
| | R Cingulate Gyrus, anterior division | 3.21 | 2 | -4 | 34 |
| 16 | | | | | |
| | R Temporal Pole | 3.21 | 52 | 12 | -32 |
| 16 | | | | | |
| | L Temporal Pole | 3.18 | -30 | 4 | -28 |
| 16 | | | | | |
| | R Temporal Pole | 2.95 | 30 | 10 | -30 |
| 16 | | | | | |
| | R Hippocampus | 2.96 | 28 | -18 | -20 |
| 16 | | | | | |
| | R Frontal Medial Cortex | 2.8 | 0 | 50 | -20 |
| 16 | | | | | |
| | R Frontal Orbital Cortex | 2.93 | 14 | 22 | -18 |
| 16 | | | | | |
| | R Frontal Orbital Cortex | 2.98 | 36 | 32 | -18 |
| 16 | | | | | |
| | R Brain-Stem | 3.1 | 6 | -30 | -14 |
| 16 | | | | | |
| | L Amygdala | 2.46 | -30 | -10 | -16 |
| 16 | | | | | |
| | R Frontal Medial Cortex | 2.82 | 0 | 38 | -16 |
| 16 | | | | | |
| | L Hippocampus | 3 | -28 | -28 | -12 |
| 16 | | | | | |
| | L Frontal Pole | 2.46 | -28 | 66 | 8 |
| 16 | | | | | |
| | R Paracingulate Gyrus | 3.01 | 10 | 46 | 14 |
| 16 | | | | | |
| | R Occipital Pole | 3.02 | 4 | -98 | 18 |
| 16 | | | | | |
| | L Precuneus Cortex | 2.96 | -16 | -64 | 28 |
| 16 | | | | | |

| | | | | | |
|---|---|---|---|---|---|
| | L Cingulate Gyrus, anterior division | 2.88 | -4 | 0 | 28 |
| 16 | | | | | |
| | R Cuneal Cortex | 3.44 | 2 | -88 | 34 |
| 16 | | | | | |
| | L Precuneus Cortex | 3.02 | -6 | -70 | 34 |
| 16 | | | | | |
| | R Cingulate Gyrus, posterior division | 2.89 | 6 | -22 | 42 |
| 8 | | | | | |
| | R Temporal Pole | 2.48 | 28 | 10 | -42 |
| 8 | | | | | |
| | L Temporal Pole | 2.42 | -32 | 14 | -40 |
| 8 | | | | | |
| | R Para hippocampal Gyrus, anterior division | 2.91 | 28 | -10 | -38 |
| 8 | | | | | |
| | R Temporal Pole | 3.03 | 42 | 8 | -38 |
| 8 | | | | | |
| | L Parahippocampal Gyrus, anterior division | 2.74 | -22 | -8 | -36 |
| 8 | | | | | |
| | R Temporal Pole | 2.71 | 42 | 20 | -36 |
| 8 | | | | | |
| | R Parahippocampal Gyrus, anterior division | 2.89 | 22 | -4 | -34 |
| 8 | | | | | |
| | L Parahippocampal Gyrus, anterior division | 2.8 | -24 | -4 | -34 |
| 8 | | | | | |
| | L Parahippocampal Gyrus, anterior division | 2.89 | -28 | -4 | -34 |
| 8 | | | | | |
| | R Parahippocampal Gyrus, anterior division | 2.87 | 26 | -2 | -34 |
| 8 | | | | | |
| | R Temporal Pole | 2.99 | 28 | 16 | -32 |
| 8 | | | | | |
| | R Temporal Pole | 3.06 | 42 | 12 | -30 |
| 8 | | | | | |
| | R Temporal Occipital Fusiform Cortex | 3.26 | 44 | -42 | -26 |
| 8 | | | | | |
| | R Frontal Orbital Cortex | 2.49 | 26 | 28 | -24 |
| 8 | | | | | |
| | L Frontal Medial Cortex | 2.51 | -10 | 34 | -22 |
| 8 | | | | | |

| | | | | | |
|---|---|---|---|---|---|
| | L Inferior Temporal Gyrus, posterior division | 2.77 | -56 | -42 | -20 |
| 8 | | | | | |
| | L Frontal Pole | 2.37 | -16 | 44 | -20 |
| 8 | | | | | |
| | L Frontal Pole | 2.56 | -18 | 48 | -20 |
| 8 | | | | | |
| | R Lateral Occipital Cortex, inferior division | 3.04 | 48 | -64 | -18 |
| 8 | | | | | |
| | L Frontal Orbital Cortex | 2.94 | -30 | 14 | -18 |
| 8 | | | | | |
| | L Amygdala | 2.64 | -18 | -6 | -16 |
| 8 | | | | | |
| | R Temporal Pole | 2.98 | 52 | 12 | -16 |
| 8 | | | | | |
| | L Hippocampus | 2.36 | -34 | -22 | -12 |
| 8 | | | | | |
| | R Superior Temporal Gyrus, posterior division | 2.89 | 60 | -22 | -2 |
| 8 | | | | | |
| | L Thalamus | 2.94 | -2 | -22 | -2 |
| 8 | | | | | |
| | L Hippocampus | 3.17 | -24 | -36 | 0 |
| 8 | | | | | |
| | R Insular Cortex | 2.92 | 34 | 22 | 0 |
| 8 | | | | | |
| | R Cingulate Gyrus, posterior division | 2.94 | 8 | -42 | 6 |
| 8 | | | | | |
| | L Thalamus | 2.58 | -10 | -32 | 10 |
| 8 | | | | | |
| | R Cingulate Gyrus, posterior division | 2.94 | 6 | -44 | 12 |
| 8 | | | | | |
| | R Cingulate Gyrus, posterior division | 2.46 | 4 | -48 | 16 |
| 8 | | | | | |
| | L Cuneal Cortex | 2.87 | -2 | -86 | 20 |
| 8 | | | | | |
| | R Frontal Pole | 2.89 | 2 | 64 | 26 |
| 8 | | | | | |
| | R Cingulate Gyrus, anterior division | 2.41 | 4 | 8 | 28 |
| 8 | | | | | |
| | L Cingulate Gyrus, anterior division | 2.98 | -2 | -12 | 30 |
| 8 | | | | | |
| | R Middle Frontal Gyrus | 2.99 | 38 | 14 | 34 |

| | | | | | |
|---|---|---|---|---|---|
| 8 | | | | | |
| | R Middle Frontal Gyrus | 2.86 | 56 | 20 | 34 |
| 8 | | | | | |
| | R Superior Parietal Lobule | 2.92 | 30 | -42 | 42 |
| 8 | | | | | |
| | R Cingulate Gyrus, posterior division | 2.75 | 4 | -26 | 42 |
| 8 | | | | | |
| | R Precuneus Cortex | 2.75 | 12 | -48 | 46 |
| 8 | | | | | |
| | R Supramarginal Gyrus, posterior division | 2.91 | 48 | -36 | 46 |
| 8 | | | | | |
| | R Supramarginal Gyrus, anterior division | 2.9 | 54 | -32 | 48 |
| 8 | | | | | |
| | R Superior Frontal Gyrus | 2.77 | 22 | 10 | 54 |
| 8 | | | | | |
| | R Lateral Occipital Cortex, superior division | 2.66 | 36 | -64 | 58 |
| 8 | | | | | |
| | R Precuneus Cortex | 2.88 | 4 | -50 | 70 |

**Supplementary Table S23. [$US_{onset}$ > $UT_{onset}$].** Descriptive statistics for clusters and local maxima showing greater activation for Uncertain-Safety relative to Uncertain-Threat anticipation for the OSP Onset regressor (FDR $q$<0.05, whole-brain corrected).

| *mm³* | **Label** | ***t*** | ***x*** | ***y*** | ***z*** |
|---|---|---|---|---|---|
| 27,264 | | | | | |
| | L Lingual Gyrus | 5.6 | -16 | -52 | -10 |
| | L Temporal Occipital Fusiform Cortex | 5.58 | -26 | -56 | -8 |
| | R Temporal Occipital Fusiform Cortex | 4.1 | 24 | -58 | -16 |
| | R Lingual Gyrus | 7.76 | 18 | -62 | -8 |
| | R Supracalcarine Cortex | 3.98 | 18 | -64 | 16 |
| | R Cuneal Cortex | 6.13 | 10 | -72 | 24 |
| | R Intracalcarine Cortex | 10.58 | 14 | -74 | 12 |
| | L Intracalcarine Cortex | 10.46 | -12 | -78 | 10 |
| 20,304 | | | | | |
| | R Temporal Pole | 3.36 | 46 | 10 | -12 |
| | R Insular Cortex | 5.99 | 42 | 8 | -2 |
| | R Precentral Gyrus | 2.84 | 60 | 6 | 12 |
| | R Superior Temporal Gyrus, anterior division | 5.55 | 62 | 4 | 0 |
| | R Central Opercular Cortex | 5.89 | 60 | -2 | 6 |
| | R Planum Polare | 4.93 | 48 | -2 | -8 |
| | R Right Amygdala | 4.3 | 24 | -6 | -10 |
| | R Right Putamen | 8.61 | 32 | -14 | 0 |
| | R Heschls Gyrus (includes H1 and H2) | 5.93 | 38 | -22 | 14 |
| | R Planum Temporale | 5.9 | 64 | -22 | 12 |
| | R Superior Temporal Gyrus, posterior division | 4.96 | 66 | -26 | 10 |
| | R Parietal Operculum Cortex | 5.71 | 44 | -32 | 20 |
| 20,240 | | | | | |
| | L Frontal Pole | 4.26 | -34 | 44 | -16 |
| | L Left Accumbens | 3.01 | -10 | 6 | -12 |
| | L Insular Cortex | 5.05 | -38 | 2 | -2 |
| | L Precentral Gyrus | 4.52 | -54 | 0 | 14 |
| | L Left Amygdala | 8.32 | -26 | -2 | -12 |
| | L Planum Polare | 5.52 | -48 | -8 | 2 |
| | L Left Pallidum | 3.93 | -20 | -8 | -6 |
| | L Central Opercular Cortex | 5.77 | -40 | -12 | 18 |
| | L Left Putamen | 7.88 | -30 | -16 | 2 |
| | L Heschls Gyrus (includes H1 and H2) | 5.99 | -42 | -18 | 10 |
| | L Planum Temporale | 5.56 | -60 | -28 | 14 |
| | L Parietal Operculum Cortex | 5.84 | -34 | -30 | 18 |
| 11,504 | | | | | |
| | R Frontal Pole | 6.27 | 32 | 58 | 24 |
| | R Superior Frontal Gyrus | 5.78 | 18 | 30 | 58 |
| | R Middle Frontal Gyrus | 3.86 | 30 | 30 | 50 |

| | | | | | |
|---|---|---|---|---|---|
| 6,072 | | | | | |
| | R Cingulate Gyrus, anterior division | 4.21 | 8 | -4 | 42 |
| | R Juxtapositional Lobule Cortex (formerly Supplementary Motor Cortex) | 4.14 | 8 | -10 | 48 |
| | L Juxtapositional Lobule Cortex (formerly Supplementary Motor Cortex) | 4.07 | -8 | -12 | 48 |
| | R Precentral Gyrus | 5.32 | 2 | -18 | 60 |
| | L Precentral Gyrus | 4.66 | -6 | -22 | 60 |
| | R Postcentral Gyrus | 4.44 | 6 | -38 | 62 |
| 4,192 | | | | | |
| | L Frontal Pole | 5.33 | -18 | 42 | 50 |
| | L Superior Frontal Gyrus | 5.07 | -18 | 32 | 58 |
| | L Middle Frontal Gyrus | 5.25 | -34 | 26 | 52 |
| 4,136 | | | | | |
| | L Postcentral Gyrus | 4.82 | -26 | -30 | 68 |
| | L Precentral Gyrus | 7.08 | -20 | -30 | 60 |
| | L Superior Parietal Lobule | 3.09 | -32 | -40 | 60 |
| 3,120 | | | | | |
| | R Precentral Gyrus | 6.75 | 24 | -26 | 58 |
| | R Postcentral Gyrus | 4.78 | 24 | -34 | 62 |
| 2,528 | | | | | |
| | L Angular Gyrus | 5.31 | -46 | -58 | 34 |
| | L Lateral Occipital Cortex, superior division | 6.15 | -56 | -68 | 32 |
| 1,360 | | | | | |
| | L Precentral Gyrus | 6.23 | -36 | -18 | 44 |
| | L Postcentral Gyrus | 5.84 | -40 | -18 | 48 |
| 1,312 | | | | | |
| | R Angular Gyrus | 3.43 | 50 | -56 | 42 |
| | R Lateral Occipital Cortex, superior division | 5.67 | 56 | -60 | 36 |
| 1,056 | | | | | |
| | R Right Caudate | 5.1 | 14 | 20 | 12 |
| | R Right Thalamus | 4.78 | 14 | -12 | 16 |
| 920 | | | | | |
| | R Brain-Stem | 3.22 | 18 | -30 | -30 |
| 600 | | | | | |
| | L Left Caudate | 4.39 | -16 | -4 | 24 |
| | L Left Thalamus | 4.65 | -14 | -20 | 2 |
| 416 | | | | | |
| | L Frontal Pole | 3.67 | -10 | 70 | 16 |
| 392 | | | | | |
| | R Frontal Medial Cortex | 5.75 | 4 | 38 | -30 |
| 392 | | | | | |
| | L Postcentral Gyrus | 3.7 | -56 | -16 | 42 |

| | | | | | |
|---|---|---|---|---|---|
| 384 | | | | | |
| | R Right Thalamus | 4.58 | 16 | -24 | 0 |
| 288 | | | | | |
| | L Postcentral Gyrus | 3.33 | -52 | -8 | 26 |
| 288 | | | | | |
| | R Precentral Gyrus | 3.71 | 40 | -16 | 40 |
| 256 | | | | | |
| | L Subcallosal Cortex | 4.03 | -2 | 26 | -2 |
| | R Cingulate Gyrus, anterior division | 4.51 | 0 | 32 | 0 |
| 248 | | | | | |
| | R Precentral Gyrus | 3.43 | 56 | -2 | 26 |
| | R Postcentral Gyrus | 3.03 | 62 | -6 | 26 |
| 208 | | | | | |
| | R Brain-Stem | 4.94 | 2 | -46 | -54 |
| 144 | | | | | |
| | L Frontal Pole | 3.64 | -30 | 52 | 4 |
| 136 | | | | | |
| | L Lateral Occipital Cortex, superior division | 3.45 | -46 | -68 | 22 |
| 128 | | | | | |
| | R Paracingulate Gyrus | 3.28 | 4 | 30 | 34 |
| 120 | | | | | |
| | L Precuneus Cortex | 3.79 | -6 | -60 | 32 |
| 120 | | | | | |
| | R Juxtapositional Lobule Cortex (formerly Supplementary Motor Cortex) | 3.76 | 0 | 4 | 46 |
| 112 | | | | | |
| | L Frontal Pole | 4.44 | -40 | 46 | -6 |
| 112 | | | | | |
| | R Postcentral Gyrus | 3.66 | 54 | -14 | 38 |
| 112 | | | | | |
| | L Postcentral Gyrus | 3.58 | -12 | -44 | 60 |
| 96 | | | | | |
| | L Frontal Pole | 3.34 | -36 | 46 | 8 |
| 80 | | | | | |
| | R Paracingulate Gyrus | 3.56 | 0 | 34 | 28 |
| 80 | | | | | |
| | L Middle Frontal Gyrus | 3.55 | -42 | 32 | 40 |
| 72 | | | | | |
| | R Postcentral Gyrus | 3.72 | 66 | -16 | 26 |
| 64 | | | | | |
| | L Frontal Medial Cortex | 3.24 | -4 | 42 | -28 |
| 56 | | | | | |
| | L Frontal Pole | 3.34 | -34 | 58 | -10 |
| 56 | | | | | |

| | | | | | |
|---|---|---|---|---|---|
| | R Postcentral Gyrus | 3.22 | 62 | -6 | 16 |
| 48 | | | | | |
| | L Middle Temporal Gyrus, posterior division | 3.44 | -60 | -12 | -22 |
| 48 | | | | | |
| | L Frontal Pole | 3.1 | -22 | 58 | -6 |
| 48 | | | | | |
| | R Parietal Operculum Cortex | 3.36 | 60 | -34 | 28 |
| 40 | | | | | |
| | R Frontal Pole | 3.11 | 32 | 54 | -14 |
| 40 | | | | | |
| | L Left Caudate | 3.57 | -14 | 20 | 10 |
| 32 | | | | | |
| | L Middle Temporal Gyrus, posterior division | 3.36 | -66 | -28 | -18 |
| 32 | | | | | |
| | R Right Hippocampus | 3.43 | 18 | -16 | -18 |
| 32 | | | | | |
| | L Middle Temporal Gyrus, posterior division | 3.66 | -56 | -22 | -16 |
| 32 | | | | | |
| | R Lateral Occipital Cortex, inferior division | 2.83 | 42 | -64 | 10 |
| 24 | | | | | |
| | L Brain-Stem | 2.97 | -10 | -40 | -46 |
| 24 | | | | | |
| | L Frontal Medial Cortex | 3.09 | -4 | 46 | -24 |
| 24 | | | | | |
| | R Temporal Pole | 3.13 | 38 | 14 | -20 |
| 24 | | | | | |
| | L Temporal Pole | 2.87 | -42 | 12 | -20 |
| 24 | | | | | |
| | R Frontal Pole | 3.41 | 18 | 60 | -12 |
| 24 | | | | | |
| | R Angular Gyrus | 2.9 | 44 | -56 | 28 |
| 24 | | | | | |
| | L Frontal Pole | 3.14 | -28 | 42 | 28 |
| 24 | | | | | |
| | L Precentral Gyrus | 2.78 | -54 | 2 | 28 |
| 24 | | | | | |
| | R Precentral Gyrus | 3.16 | 44 | -6 | 50 |
| 24 | | | | | |
| | L Superior Parietal Lobule | 2.85 | -14 | -52 | 68 |
| 16 | | | | | |
| | L Inferior Temporal Gyrus, anterior division | 3.37 | -52 | -10 | -38 |

| | | | | | |
|---|---|---|---|---|---|
| 16 | | | | | |
| | L Middle Temporal Gyrus, anterior division | 2.97 | -62 | -4 | -26 |
| 16 | | | | | |
| | L Temporal Occipital Fusiform Cortex | 3.22 | -32 | -54 | -20 |
| 16 | | | | | |
| | L Middle Temporal Gyrus, posterior division | 2.7 | -60 | -32 | -16 |
| 16 | | | | | |
| | L Middle Temporal Gyrus, posterior division | 2.91 | -68 | -30 | -14 |
| 16 | | | | | |
| | R Insular Cortex | 3.1 | 34 | 14 | -8 |
| 16 | | | | | |
| | R Frontal Pole | 2.76 | 22 | 56 | -4 |
| 16 | | | | | |
| | L Frontal Pole | 2.71 | -32 | 56 | -4 |
| 16 | | | | | |
| | R Cingulate Gyrus, anterior division | 3.21 | 8 | 40 | 0 |
| 16 | | | | | |
| | R Left Thalamus | 2.88 | 0 | -8 | 0 |
| 16 | | | | | |
| | R Frontal Pole | 2.75 | 40 | 46 | 0 |
| 16 | | | | | |
| | L Cingulate Gyrus, anterior division | 3.84 | -8 | 38 | 10 |
| 16 | | | | | |
| | L Left Thalamus | 3.63 | -12 | -4 | 14 |
| 16 | | | | | |
| | R Precentral Gyrus | 2.9 | 56 | 4 | 12 |
| 16 | | | | | |
| | R Right Thalamus | 2.82 | 20 | -20 | 14 |
| 16 | | | | | |
| | L Left Thalamus | 3.26 | -16 | -20 | 18 |
| 16 | | | | | |
| | L Cuneal Cortex | 2.68 | -18 | -72 | 20 |
| 16 | | | | | |
| | R Cingulate Gyrus, anterior division | 3.38 | 12 | 30 | 22 |
| 16 | | | | | |
| | R Paracingulate Gyrus | 2.9 | 2 | 12 | 42 |
| 16 | | | | | |
| | L Precentral Gyrus | 2.85 | -30 | -20 | 74 |
| 8 | | | | | |
| | L Brain-Stem | 2.87 | -6 | -48 | -56 |
| 8 | | | | | |
| | L Brain-Stem | 2.72 | -14 | -28 | -34 |

| | | | | | |
|---|---|---|---|---|---|
| 8 | | | | | |
| | L Temporal Pole | 2.85 | -20 | 4 | -32 |
| 8 | | | | | |
| | L Inferior Temporal Gyrus, posterior division | 2.71 | -56 | -20 | -28 |
| 8 | | | | | |
| | R Inferior Temporal Gyrus, posterior division | 2.67 | 64 | -22 | -26 |
| 8 | | | | | |
| | L Brain-Stem | 2.81 | -14 | -22 | -22 |
| 8 | | | | | |
| | L Left Hippocampus | 3.46 | -12 | -12 | -18 |
| 8 | | | | | |
| | R Frontal Pole | 3.06 | 26 | 46 | -16 |
| 8 | | | | | |
| | R Periaqueductal gray | 2.69 | 0 | -32 | -12 |
| 8 | | | | | |
| | L Middle Temporal Gyrus, anterior division | 2.88 | -62 | -4 | -10 |
| 8 | | | | | |
| | R Right Hippocampus | 5.06 | 24 | -26 | -8 |
| 8 | | | | | |
| | R Accumbens | 2.8 | 12 | 18 | -8 |
| 8 | | | | | |
| | R Cingulate Gyrus, anterior division | 2.75 | 4 | 38 | -6 |
| 8 | | | | | |
| | L Left Thalamus | 3.01 | -2 | -12 | -4 |
| 8 | | | | | |
| | L Left Pallidum | 2.71 | -12 | 2 | 0 |
| 8 | | | | | |
| | L Left Caudate | 2.81 | -10 | 18 | 2 |
| 8 | | | | | |
| | L Lateral Occipital Cortex, inferior division | 3.05 | -42 | -76 | 10 |
| 8 | | | | | |
| | R Right Thalamus | 2.97 | 16 | -22 | 10 |
| 8 | | | | | |
| | L Frontal Pole | 2.76 | -28 | 48 | 12 |
| 8 | | | | | |
| | R Frontal Pole | 3.2 | 18 | 62 | 12 |
| 8 | | | | | |
| | R Cingulate Gyrus, anterior division | 2.71 | 2 | 30 | 14 |
| 8 | | | | | |
| | R Frontal Pole | 2.71 | 14 | 66 | 14 |
| 8 | | | | | |
| | R Right Thalamus | 3 | 18 | -24 | 16 |

| | | | | | |
|---|---|---|---|---|---|
| 8 | | | | | |
| | R Occipital Pole | 2.86 | 6 | -88 | 18 |
| 8 | | | | | |
| | L Lateral Occipital Cortex, superior division | 2.71 | -22 | -86 | 18 |
| 8 | | | | | |
| | L Planum Temporale | 3 | -56 | -40 | 18 |
| 8 | | | | | |
| | L Central Opercular Cortex | 2.86 | -40 | -4 | 18 |
| 8 | | | | | |
| | L Cingulate Gyrus, anterior division | 2.76 | -8 | 32 | 18 |
| 8 | | | | | |
| | R Frontal Pole | 2.78 | 6 | 70 | 18 |
| 8 | | | | | |
| | L Precentral Gyrus | 2.76 | -52 | 2 | 24 |
| 8 | | | | | |
| | L Precentral Gyrus | 2.94 | -62 | -2 | 26 |
| 8 | | | | | |
| | L Frontal Pole | 2.79 | -8 | 64 | 30 |
| 8 | | | | | |
| | R Supramarginal Gyrus, anterior division | 2.81 | 56 | -20 | 32 |
| 8 | | | | | |
| | L Angular Gyrus | 2.73 | -50 | -52 | 34 |
| 8 | | | | | |
| | R Postcentral Gyrus | 2.68 | 42 | -16 | 34 |
| 8 | | | | | |
| | L Precentral Gyrus | 2.72 | -14 | -30 | 44 |
| 8 | | | | | |
| | R Precentral Gyrus | 2.73 | 50 | -12 | 48 |
| 8 | | | | | |
| | R Middle Frontal Gyrus | 2.7 | 34 | 24 | 50 |
| 8 | | | | | |
| | R Middle Frontal Gyrus | 2.85 | 38 | 22 | 52 |
| 8 | | | | | |
| | R Postcentral Gyrus | 2.69 | 32 | -30 | 56 |
| 8 | | | | | |
| | L Superior Parietal Lobule | 2.7 | -20 | -48 | 62 |
| 8 | | | | | |
| | L Postcentral Gyrus | 2.73 | -44 | -38 | 62 |
| 8 | | | | | |
| | L Precentral Gyrus | 2.68 | -2 | -26 | 70 |

**Supplementary Table S24. [$CS_{onset}$ > $CT_{onset}$].** Descriptive statistics for clusters and local maxima showing greater activation for Certain-Safety relative to Certain-Threat anticipation for the OSP Onset regressor (FDR *q*<0.05, whole-brain corrected).

| ***mm³*** | **Label** | ***t*** | ***x*** | ***y*** | ***z*** |
|---|---|---|---|---|---|
| 52,856 | | | | | |
| | L Cingulate Gyrus, anterior division | 3.71 | -4 | 12 | 40 |
| | R Paracingulate Gyrus | 4.07 | 2 | 12 | 50 |
| | L Juxtapositional Lobule Cortex (formerly Supplementary Motor Cortex) | 6.58 | -4 | 6 | 56 |
| | L Middle Frontal Gyrus | 7.51 | -38 | -2 | 62 |
| | L Superior Frontal Gyrus | 7.55 | -6 | -8 | 74 |
| | R Juxtapositional Lobule Cortex (formerly Supplementary Motor Cortex) | 7.77 | 4 | -8 | 72 |
| | R Superior Frontal Gyrus | 5.45 | 14 | -8 | 74 |
| | L Precentral Gyrus | 9.30 | -18 | -24 | 76 |
| | R Precentral Gyrus | 8.61 | 20 | -26 | 74 |
| | R Postcentral Gyrus | 6.57 | 24 | -34 | 72 |
| | L Postcentral Gyrus | 8.72 | -20 | -36 | 74 |
| | L Superior Parietal Lobule | 8.12 | -22 | -46 | 70 |
| | R Superior Parietal Lobule | 5.52 | 22 | -48 | 70 |
| | R Lateral Occipital Cortex, superior division | 4.95 | 26 | -58 | 60 |
| | L Lateral Occipital Cortex, superior division | 4.92 | -24 | -66 | 56 |
| 21,776 | | | | | |
| | R Parahippocampal Gyrus, posterior division | 3.42 | 20 | -28 | -18 |
| | L Temporal Fusiform Cortex, posterior division | 4.94 | -28 | -42 | -14 |
| | R Lingual Gyrus | 8.44 | 24 | -56 | -8 |
| | L Lingual Gyrus | 8.20 | -28 | -58 | -6 |
| | L Precuneus Cortex | 3.53 | -6 | -62 | 14 |
| | R Occipital Fusiform Gyrus | 5.76 | 26 | -70 | -8 |
| | R Intracalcarine Cortex | 12.11 | 14 | -72 | 12 |
| | R Cuneal Cortex | 7.16 | 8 | -72 | 22 |
| | L Occipital Fusiform Gyrus | 5.91 | -28 | -74 | -8 |
| | L Intracalcarine Cortex | 12.44 | -12 | -76 | 8 |
| | L Cuneal Cortex | 4.29 | -6 | -78 | 22 |
| 6,392 | | | | | |
| | L Superior Temporal Gyrus, posterior division | 2.91 | -62 | -32 | 2 |
| | L Middle Temporal Gyrus, temporooccipital part | 5.17 | -56 | -48 | 6 |
| | L Supramarginal Gyrus, posterior division | 4.15 | -50 | -48 | 12 |
| | L Angular Gyrus | 6.36 | -60 | -52 | 12 |
| | L Lateral Occipital Cortex, inferior division | 3.71 | -44 | -68 | 12 |

| | | | | | |
|---|---|---|---|---|---|
| | L Lateral Occipital Cortex, superior division | 4.93 | -54 | -70 | 16 |
| | L Occipital Pole | 4.41 | -38 | -92 | 12 |
| 1,344 | | | | | |
| | R Accumbens | 4.57 | 10 | 20 | -4 |
| | R Putamen | 4.90 | 18 | 16 | -6 |
| | R Amygdala | 4.19 | 24 | 0 | -12 |
| 1,200 | | | | | |
| | R Postcentral Gyrus | 4.60 | 58 | -16 | 40 |
| | R Precentral Gyrus | 4.50 | 48 | -10 | 58 |
| 1,160 | | | | | |
| | L Putamen | 4.35 | -28 | -2 | -2 |
| | L Pallidum | 3.84 | -20 | 2 | 0 |
| 856 | | | | | |
| | L Accumbens | 5.16 | -10 | 10 | -12 |
| 832 | | | | | |
| | R Angular Gyrus | 4.84 | 62 | -48 | 14 |
| | R Middle Temporal Gyrus, temporooccipital part | 3.83 | 54 | -50 | 10 |
| | R Supramarginal Gyrus, posterior division | 2.89 | 64 | -42 | 20 |
| 800 | | | | | |
| | R Postcentral Gyrus | 5.53 | 12 | -42 | 60 |
| | R Precuneus Cortex | 5.26 | 8 | -44 | 64 |
| | R Superior Parietal Lobule | 3.03 | 10 | -48 | 70 |
| 680 | | | | | |
| | L Lateral Occipital Cortex, superior division | 5.13 | -32 | -84 | 22 |
| | L Occipital Pole | 3.23 | -24 | -92 | 26 |
| 672 | | | | | |
| | R Superior Frontal Gyrus | 4.48 | 18 | 28 | 60 |
| 416 | | | | | |
| | L Cingulate Gyrus, anterior division | 3.47 | -2 | 18 | 34 |
| | R Paracingulate Gyrus | 3.99 | 4 | 20 | 38 |
| | R Cingulate Gyrus, anterior division | 3.87 | 8 | 14 | 38 |
| 384 | | | | | |
| | L Middle Temporal Gyrus, anterior division | 4.37 | -56 | -2 | -16 |
| | L Superior Temporal Gyrus, anterior division | 4.08 | -60 | 0 | -12 |
| 368 | | | | | |
| | R Putamen | 5.06 | 32 | -14 | -2 |
| 264 | | | | | |
| | R Angular Gyrus | 3.96 | 48 | -58 | 22 |
| | R Lateral Occipital Cortex, superior division | 3.18 | 56 | -62 | 28 |

| | | | | | |
|---|---|---|---|---|---|
| 216 | | | | | |
| | R Postcentral Gyrus | 3.39 | 54 | -22 | 50 |
| 144 | | | | | |
| | L Insular Cortex | 3.74 | -42 | 6 | 0 |
| | L Central Opercular Cortex | 3.62 | -48 | 6 | 2 |
| 128 | | | | | |
| | R Precentral Gyrus | 3.39 | 64 | 6 | 26 |
| 112 | | | | | |
| | R Putamen | 4.47 | 26 | -4 | 6 |
| 104 | | | | | |
| | R Putamen | 3.63 | 28 | -4 | -6 |
| 104 | | | | | |
| | L Parietal Operculum Cortex | 3.70 | -40 | -28 | 20 |
| 88 | | | | | |
| | R Middle Temporal Gyrus, temporooccipital part | 4.11 | 58 | -42 | 6 |
| 88 | | | | | |
| | R Supramarginal Gyrus, anterior division | 3.32 | 62 | -20 | 26 |
| 80 | | | | | |
| | L Superior Temporal Gyrus, posterior division | 3.59 | -56 | -22 | -4 |
| 80 | | | | | |
| | R Lateral Occipital Cortex, superior division | 3.05 | 58 | -62 | 20 |
| 80 | | | | | |
| | R Superior Frontal Gyrus | 3.35 | 10 | 8 | 70 |
| 72 | | | | | |
| | L Central Opercular Cortex | 3.30 | -58 | -20 | 18 |
| 72 | | | | | |
| | L Frontal Pole | 3.25 | -12 | 42 | 52 |
| 64 | | | | | |
| | L Temporal Occipital Fusiform Cortex | 3.65 | -42 | -58 | -14 |
| 64 | | | | | |
| | L Hippocampus | 3.94 | -22 | -28 | -8 |
| 64 | | | | | |
| | L Precentral Gyrus | 3.25 | -56 | 2 | 28 |
| 56 | | | | | |
| | L Precentral Gyrus | 3.60 | -10 | -22 | 46 |
| 48 | | | | | |
| | L Temporal Fusiform Cortex, posterior division | 3.13 | -40 | -24 | -18 |
| 48 | | | | | |
| | L Central Opercular Cortex | 3.24 | -48 | -22 | 20 |
| 48 | | | | | |
| | R Cingulate Gyrus, anterior division | 3.22 | 4 | 34 | 22 |

| | | | | | |
|---|---|---|---|---|---|
| 48 | | | | | |
| | L Postcentral Gyrus | 3.23 | -54 | -20 | 30 |
| 40 | | | | | |
| | L Temporal Occipital Fusiform Cortex | 3.09 | -30 | -46 | -18 |
| 40 | | | | | |
| | L Thalamus | 3.18 | -10 | -16 | 4 |
| 40 | | | | | |
| | R Frontal Pole | 3.43 | 18 | 46 | 46 |
| 40 | | | | | |
| | R Lateral Occipital Cortex, superior division | 3.51 | 14 | -62 | 62 |
| 32 | | | | | |
| | L Temporal Fusiform Cortex, posterior division | 3.30 | -34 | -24 | -26 |
| 32 | | | | | |
| | L Thalamus | 2.96 | -16 | -24 | 12 |
| 32 | | | | | |
| | L Central Opercular Cortex | 2.97 | -50 | -22 | 14 |
| | L Parietal Operculum Cortex | 2.94 | -48 | -24 | 16 |
| 32 | | | | | |
| | R Cingulate Gyrus, anterior division | 3.20 | 4 | 30 | 18 |
| 24 | | | | | |
| | L Inferior Temporal Gyrus, anterior division | 3.45 | -44 | -6 | -46 |
| 24 | | | | | |
| | L Brain-Stem | 3.84 | -12 | -34 | -36 |
| 24 | | | | | |
| | L Lateral Occipital Cortex, inferior division | 3.11 | -46 | -66 | -12 |
| 24 | | | | | |
| | R Temporal Occipital Fusiform Cortex | 2.99 | 32 | -42 | -10 |
| 24 | | | | | |
| | R Hippocampus | 5.79 | 24 | -26 | -8 |
| 24 | | | | | |
| | R Insular Cortex | 2.96 | 44 | 6 | 0 |
| 24 | | | | | |
| | R Caudate | 3.25 | 18 | 12 | 18 |
| 24 | | | | | |
| | L Lateral Occipital Cortex, superior division | 2.83 | -18 | -84 | 20 |
| 24 | | | | | |
| | R Lateral Occipital Cortex, superior division | 3.12 | 24 | -84 | 20 |
| 24 | | | | | |
| | R Occipital Pole | 2.77 | 12 | -90 | 28 |
| 24 | | | | | |

| | | | | | |
|---|---|---|---|---|---|
| | R Middle Frontal Gyrus | 3.32 | 30 | 24 | 48 |
| 24 | | | | | |
| | R Precentral Gyrus | 3.18 | 44 | 0 | 52 |
| 24 | | | | | |
| | L Middle Frontal Gyrus | 3.05 | -30 | 22 | 56 |
| 16 | | | | | |
| | L Brain-Stem | 2.99 | -20 | -28 | -38 |
| 16 | | | | | |
| | L Temporal Pole | 3.39 | -50 | 12 | -20 |
| 16 | | | | | |
| | L Parahippocampal Gyrus, posterior division | 3.21 | -20 | -34 | -20 |
| 16 | | | | | |
| | L Middle Temporal Gyrus, anterior division | 2.82 | -60 | -8 | -10 |
| 16 | | | | | |
| | L Putamen | 3.04 | -26 | 10 | -4 |
| 16 | | | | | |
| | R Inferior Frontal Gyrus, pars triangularis | 3.13 | 52 | 28 | -4 |
| 16 | | | | | |
| | R Precuneus Cortex | 3.00 | 18 | -50 | 6 |
| 16 | | | | | |
| | R Caudate | 2.91 | 16 | 20 | 8 |
| 16 | | | | | |
| | R Thalamus | 3.63 | 14 | -16 | 20 |
| 16 | | | | | |
| | R Supramarginal Gyrus, posterior division | 2.80 | 60 | -44 | 24 |
| 16 | | | | | |
| | R Lateral Occipital Cortex, superior division | 3.09 | 56 | -70 | 24 |
| 16 | | | | | |
| | L Supramarginal Gyrus, anterior division | 3.00 | -62 | -40 | 28 |
| 16 | | | | | |
| | R Paracingulate Gyrus | 3.56 | 2 | 28 | 32 |
| 16 | | | | | |
| | L Middle Frontal Gyrus | 2.96 | -40 | 10 | 38 |
| 16 | | | | | |
| | L Lateral Occipital Cortex, superior division | 3.15 | -24 | -74 | 40 |
| 16 | | | | | |
| | R Precentral Gyrus | 3.07 | 42 | -6 | 48 |
| 16 | | | | | |
| | L Middle Frontal Gyrus | 2.88 | -28 | 28 | 50 |
| 8 | | | | | |
| | R Brain-Stem | 2.79 | 12 | -34 | -40 |

| | | | | | |
|---|---|---|---|---|---|
| 8 | | | | | |
| | R Inferior Temporal Gyrus, posterior division | 2.74 | 50 | -14 | -32 |
| 8 | | | | | |
| | L Temporal Pole | 3.08 | -24 | 8 | -32 |
| 8 | | | | | |
| | R Temporal Pole | 2.94 | 56 | 14 | -26 |
| 8 | | | | | |
| | L Temporal Fusiform Cortex, posterior division | 3.01 | -34 | -44 | -24 |
| 8 | | | | | |
| | R Temporal Fusiform Cortex, posterior division | 3.01 | 34 | -34 | -20 |
| 8 | | | | | |
| | L Temporal Fusiform Cortex, posterior division | 2.81 | -40 | -36 | -18 |
| 8 | | | | | |
| | L Temporal Pole | 3.05 | -50 | 16 | -18 |
| 8 | | | | | |
| | L Temporal Fusiform Cortex, posterior division | 3.11 | -32 | -38 | -16 |
| 8 | | | | | |
| | L Frontal Orbital Cortex | 2.76 | -34 | 20 | -16 |
| 8 | | | | | |
| | L Temporal Pole | 2.78 | -58 | 8 | -14 |
| 8 | | | | | |
| | L Hippocampus | 2.88 | -24 | -32 | -12 |
| 8 | | | | | |
| | L Frontal Orbital Cortex | 3.29 | -30 | 32 | -12 |
| 8 | | | | | |
| | L Pallidum | 2.90 | -22 | -4 | -4 |
| 8 | | | | | |
| | R Pallidum | 2.86 | 24 | -6 | -2 |
| 8 | | | | | |
| | L Caudate | 3.11 | -10 | 20 | -2 |
| 8 | | | | | |
| | L Putamen | 2.73 | -28 | 8 | 2 |
| 8 | | | | | |
| | L Insular Cortex | 2.82 | -38 | 2 | 4 |
| 8 | | | | | |
| | L Precentral Gyrus | 2.84 | -52 | 8 | 6 |
| 8 | | | | | |
| | R Inferior Frontal Gyrus, pars opercularis | 2.83 | 58 | 12 | 6 |
| 8 | | | | | |
| | L Lateral Occipital Cortex, inferior division | 2.83 | -50 | -68 | 8 |

| | | | | | |
|---|---|---|---|---|---|
| 8 | | | | | |
| | R Supramarginal Gyrus, posterior division | 2.89 | 66 | -40 | 8 |
| 8 | | | | | |
| | R Caudate | 2.79 | 18 | 10 | 12 |
| 8 | | | | | |
| | R Angular Gyrus | 2.86 | 52 | -52 | 16 |
| 8 | | | | | |
| | L Lateral Occipital Cortex, superior division | 2.74 | -22 | -82 | 18 |
| 8 | | | | | |
| | R Lateral Occipital Cortex, superior division | 2.74 | 52 | -64 | 18 |
| 8 | | | | | |
| | R Supramarginal Gyrus, posterior division | 2.76 | 46 | -44 | 18 |
| 8 | | | | | |
| | R Caudate | 3.08 | 18 | -20 | 22 |
| 8 | | | | | |
| | R Frontal Pole | 2.98 | 34 | 56 | 28 |
| 8 | | | | | |
| | L Cingulate Gyrus, anterior division | 2.85 | -8 | 16 | 34 |
| 8 | | | | | |
| | R Cingulate Gyrus, anterior division | 2.75 | 6 | 4 | 42 |
| 8 | | | | | |
| | R Precentral Gyrus | 2.82 | 60 | -6 | 44 |
| 8 | | | | | |
| | R Frontal Pole | 2.88 | 16 | 52 | 44 |
| 8 | | | | | |
| | L Lateral Occipital Cortex, superior division | 2.73 | -20 | -82 | 48 |
| 8 | | | | | |
| | L Precuneus Cortex | 3.40 | -10 | -48 | 48 |
| 8 | | | | | |
| | R Superior Frontal Gyrus | 2.73 | 20 | 30 | 48 |
| 8 | | | | | |
| | L Lateral Occipital Cortex, superior division | 2.74 | -24 | -70 | 50 |
| 8 | | | | | |
| | L Precentral Gyrus | 2.93 | -44 | -8 | 50 |
| 8 | | | | | |
| | R Superior Frontal Gyrus | 2.81 | 22 | 22 | 54 |
| 8 | | | | | |
| | R Superior Frontal Gyrus | 2.85 | 16 | 20 | 56 |
| 8 | | | | | |
| | R Precentral Gyrus | 3.08 | 38 | -14 | 64 |
| 8 | | | | | |

| | | | | | |
|---|---|---|---|---|---|
| | L Lateral Occipital Cortex, superior division | 3.10 | -14 | -70 | 66 |
| 8 | | | | | |
| | R Lateral Occipital Cortex, superior division | 3.00 | 18 | -64 | 66 |
| 8 | | | | | |
| | R Precentral Gyrus | 2.89 | 28 | -20 | 66 |
| 8 | | | | | |
| | R Superior Frontal Gyrus | 2.79 | 4 | 14 | 68 |

**Supplementary Table S25. [$CT_{CB3}$ > $CT_{CB2}$].** Descriptive statistics for clusters and local maxima showing greater activation for Convolved Block 3 relative to Convolved Block 2 for Certain-Threat anticipation (FDR $q<0.05$, whole-brain corrected).

| ***mm³*** | **Label** | ***t*** | ***x*** | ***y*** | ***z*** |
|---|---|---|---|---|---|
| 153,424 | | | | | |
| | L Paracingulate Gyrus | 4.45 | -4 | 28 | 30 |
| | R Frontal Orbital Cortex | 6.93 | 40 | 28 | 0 |
| | R Frontal Operculum Cortex | 8.48 | 34 | 24 | 8 |
| | L Frontal Orbital Cortex | 4.75 | -36 | 22 | -6 |
| | L Frontal Operculum Cortex | 9.83 | -34 | 20 | 8 |
| | R Insular Cortex | 5.92 | 38 | 20 | -8 |
| | R Cingulate Gyrus, anterior division | 4.90 | 0 | 16 | 26 |
| | L Insular Cortex | 8.72 | -36 | 14 | 4 |
| | L Cingulate Gyrus, anterior division | 8.11 | -8 | 14 | 36 |
| | R Inferior Frontal Gyrus, pars opercularis | 4.93 | 60 | 14 | 8 |
| | L Inferior Frontal Gyrus, pars opercularis | 6.35 | -54 | 12 | 0 |
| | R Paracingulate Gyrus | 8.54 | 0 | 12 | 46 |
| | L Caudate | 7.63 | -6 | 10 | -2 |
| | L Precentral Gyrus | 8.53 | -60 | 8 | 12 |
| | R Caudate | 8.23 | 8 | 8 | 2 |
| | L Superior Frontal Gyrus | 5.66 | -14 | 6 | 70 |
| | R Putamen | 11.18 | 22 | 6 | 0 |
| | R Juxtapositional Lobule Cortex (formerly Supplementary Motor Cortex) | 11.46 | 10 | 4 | 44 |
| | R Central Opercular Cortex | 7.57 | 52 | 4 | 2 |
| | L Putamen | 9.71 | -22 | 2 | 2 |
| | L Central Opercular Cortex | 7.38 | -50 | 2 | 2 |
| | R Thalamus | 8.53 | 12 | -4 | 14 |
| | R Amygdala | 4.90 | 30 | -6 | -18 |
| | L Juxtapositional Lobule Cortex (formerly Supplementary Motor Cortex) | 10.43 | -4 | -8 | 66 |
| | L Pallidum | 5.91 | -26 | -8 | -4 |
| | R Amygdala | 6.56 | 28 | -10 | -12 |
| | L Amygdala | 6.25 | -26 | -10 | -12 |
| | R Superior Frontal Gyrus | 9.96 | 14 | -10 | 70 |
| | R Amygdala | 6.56 | 28 | -10 | -12 |
| | R Planum Polare | 3.96 | 40 | -10 | -10 |
| | L Postcentral Gyrus | 5.22 | -36 | -18 | 42 |
| | R Heschls Gyrus (includes H1 and H2) | 7.36 | 52 | -18 | 8 |
| | L Thalamus | 6.75 | -4 | -22 | 0 |
| | R Postcentral Gyrus | 13.03 | 36 | -24 | 56 |
| | R Precentral Gyrus | 12.97 | 30 | -24 | 52 |
| | R Planum Temporale | 6.80 | 54 | -24 | 14 |
| | R Cingulate Gyrus, posterior division | 5.52 | 12 | -24 | 42 |
| | L Cingulate Gyrus, posterior division | 6.07 | -10 | -26 | 40 |

| | | | | | |
|---|---|---|---|---|---|
| | R Brain-Stem | 5.63 | 10 | -26 | -18 |
| | R Parietal Operculum Cortex | 8.64 | 56 | -28 | 22 |
| | R Supramarginal Gyrus, anterior division | 5.39 | 58 | -32 | 34 |
| | R Superior Temporal Gyrus, posterior division | 4.63 | 68 | -32 | 12 |
| | L Brain-Stem | 6.79 | -6 | -34 | -8 |
| | R Supramarginal Gyrus, posterior division | 7.56 | 66 | -38 | 34 |
| | R Superior Parietal Lobule | 8.25 | 24 | -42 | 60 |
| | R Angular Gyrus | 7.07 | 58 | -46 | 30 |
| 15,840 | | | | | |
| | L Central Opercular Cortex | 3.01 | -44 | -18 | 14 |
| | L Insular Cortex | 2.81 | -38 | -20 | 2 |
| | L Postcentral Gyrus | 6.18 | -62 | -22 | 26 |
| | L Heschls Gyrus (includes H1 and H2) | 6.47 | -46 | -26 | 6 |
| | L Supramarginal Gyrus, anterior division | 7.82 | -62 | -34 | 24 |
| | L Parietal Operculum Cortex | 6.41 | -46 | -36 | 22 |
| | L Planum Temporale | 6.12 | -48 | -38 | 18 |
| | L Supramarginal Gyrus, posterior division | 5.84 | -64 | -42 | 36 |
| | L Angular Gyrus | 4.21 | -46 | -56 | 46 |
| | L Lateral Occipital Cortex, superior division | 4.16 | -50 | -64 | 48 |
| 12,984 | | | | | |
| | R Occipital Fusiform Gyrus | 3.45 | 30 | -80 | -14 |
| | R Lingual Gyrus | 3.99 | 8 | -88 | -8 |
| | R Lateral Occipital Cortex, inferior division | 4.19 | 40 | -90 | -2 |
| | R Occipital Pole | 11.33 | 26 | -94 | 10 |
| 6,888 | | | | | |
| | L Frontal Pole | 8.02 | -36 | 48 | 30 |
| | L Middle Frontal Gyrus | 6.27 | -38 | 36 | 40 |
| 5,208 | | | | | |
| | R Frontal Pole | 6.87 | 36 | 46 | 32 |
| 1,504 | | | | | |
| | L Occipital Fusiform Gyrus | 5.18 | -22 | -84 | -10 |
| | L Lingual Gyrus | 6.05 | -8 | -86 | -8 |
| | L Occipital Pole | 3.96 | -20 | -96 | -12 |
| 1,056 | | | | | |
| | L Precuneus Cortex | 6.24 | -10 | -70 | 40 |
| 808 | | | | | |
| | L Frontal Pole | 4.58 | -24 | 68 | -10 |
| 808 | | | | | |
| | R Precuneus Cortex | 5.12 | 14 | -70 | 40 |
| 640 | | | | | |
| | R Middle Temporal Gyrus, posterior division | 4.33 | 60 | -22 | -10 |
| 600 | | | | | |

| | | | | | |
|---|---|---|---|---|---|
| | R Temporal Occipital Fusiform Cortex | 4.71 | 34 | -60 | -20 |
| 584 | | | | | |
| | R Frontal Pole | 4.21 | 24 | 68 | -6 |
| 424 | | | | | |
| | L Lateral Occipital Cortex, inferior division | 3.95 | -50 | -82 | -4 |
| 288 | | | | | |
| | L Precentral Gyrus | 4.99 | -8 | -24 | 58 |
| 280 | | | | | |
| | R Brain-Stem | 4.01 | 8 | -24 | -34 |
| 280 | | | | | |
| | L Frontal Orbital Cortex | 4.19 | -28 | 36 | -12 |
| 232 | | | | | |
| | R Middle Temporal Gyrus, anterior division | 3.72 | 58 | -4 | -34 |
| | R Inferior Temporal Gyrus, anterior division | 3.06 | 50 | -6 | -38 |
| 200 | | | | | |
| | R Temporal Pole | 3.73 | 48 | 12 | -38 |
| 176 | | | | | |
| | L Planum Polare | 4.77 | -44 | -10 | -10 |
| 120 | | | | | |
| | R Inferior Temporal Gyrus, temporooccipital part | 2.77 | 54 | -46 | -26 |
| 112 | | | | | |
| | L Middle Temporal Gyrus, posterior division | 3.49 | -60 | -26 | -12 |
| 112 | | | | | |
| | R Angular Gyrus | 3.21 | 44 | -52 | 58 |
| 104 | | | | | |
| | R Temporal Fusiform Cortex, posterior division | 4.02 | 34 | -16 | -32 |
| 96 | | | | | |
| | L Precuneus Cortex | 3.01 | -8 | -54 | 60 |
| 96 | | | | | |
| | R Precuneus Cortex | 3.13 | 2 | -44 | 58 |
| 80 | | | | | |
| | L Inferior Temporal Gyrus, temporooccipital part | 3.30 | -60 | -50 | -24 |
| 80 | | | | | |
| | L Supramarginal Gyrus, posterior division | 3.31 | -36 | -48 | 40 |
| 80 | | | | | |
| | R Precuneus Cortex | 3.03 | 2 | -44 | 42 |
| 72 | | | | | |
| | L Inferior Temporal Gyrus, anterior division | 3.36 | -46 | -10 | -38 |

| | | | | | |
|---|---|---|---|---|---|
| 72 | | | | | |
| | L Angular Gyrus | 3.84 | -36 | -56 | 38 |
| 72 | | | | | |
| | L Precentral Gyrus | 3.42 | -6 | -26 | 50 |
| 64 | | | | | |
| | L Postcentral Gyrus | 3.28 | -20 | -40 | 66 |
| 56 | | | | | |
| | L Inferior Temporal Gyrus, posterior division | 2.79 | -54 | -16 | -38 |
| 56 | | | | | |
| | R Frontal Orbital Cortex | 3.23 | 34 | 26 | -16 |
| 56 | | | | | |
| | R Paracingulate Gyrus | 2.44 | 6 | 44 | 34 |
| | R Superior Frontal Gyrus | 3.32 | 6 | 40 | 38 |
| 48 | | | | | |
| | L Brain-Stem | 3.24 | -4 | -40 | -56 |
| 48 | | | | | |
| | L Inferior Temporal Gyrus, posterior division | 3.23 | -50 | -40 | -22 |
| 48 | | | | | |
| | L Insular Cortex | 2.69 | -40 | -16 | -4 |
| | L Planum Polare | 2.65 | -46 | -16 | -4 |
| 48 | | | | | |
| | L Middle Temporal Gyrus, temporooccipital part | 2.87 | -58 | -60 | 8 |
| 48 | | | | | |
| | L Precuneus Cortex | 2.96 | -2 | -64 | 56 |
| 48 | | | | | |
| | L Postcentral Gyrus | 3.36 | -20 | -46 | 70 |
| 40 | | | | | |
| | L Middle Temporal Gyrus, posterior division | 2.80 | -70 | -28 | -12 |
| 40 | | | | | |
| | L Occipital Pole | 3.87 | -12 | -94 | 0 |
| 40 | | | | | |
| | R Cingulate Gyrus, anterior division | 2.85 | 2 | 34 | 4 |
| 40 | | | | | |
| | R Middle Temporal Gyrus, temporooccipital part | 2.80 | 52 | -48 | 6 |
| 40 | | | | | |
| | R Precuneus Cortex | 3.02 | 2 | -50 | 50 |
| 32 | | | | | |
| | L Brain-Stem | 3.03 | -10 | -38 | -38 |
| 32 | | | | | |
| | L Temporal Pole | 2.76 | -52 | 4 | -34 |

| | | | | | |
|---|---|---|---|---|---|
| 32 | | | | | |
| | R Inferior Temporal Gyrus, temporooccipital part | 2.71 | 58 | -50 | -20 |
| 32 | | | | | |
| | L Middle Temporal Gyrus, posterior division | 3.09 | -58 | -34 | -12 |
| 32 | | | | | |
| | L Thalamus | 2.99 | -18 | -28 | 8 |
| 32 | | | | | |
| | R Cingulate Gyrus, anterior division | 3.04 | 2 | -4 | 34 |
| 32 | | | | | |
| | L Middle Frontal Gyrus | 3.06 | -38 | 22 | 42 |
| 32 | | | | | |
| | R Superior Parietal Lobule | 2.88 | 38 | -54 | 64 |
| 32 | | | | | |
| | R Precuneus Cortex | 2.80 | 6 | -48 | 62 |
| 24 | | | | | |
| | R Parahippocampal Gyrus, anterior division | 3.28 | 22 | -10 | -32 |
| 24 | | | | | |
| | R Inferior Temporal Gyrus, posterior division | 2.74 | 54 | -12 | -30 |
| 24 | | | | | |
| | R Brain-Stem | 2.75 | 14 | -20 | -28 |
| 24 | | | | | |
| | L Inferior Temporal Gyrus, posterior division | 2.96 | -56 | -42 | -24 |
| 24 | | | | | |
| | R Temporal Fusiform Cortex, posterior division | 3.46 | 42 | -32 | -24 |
| 24 | | | | | |
| | L Frontal Orbital Cortex | 3.42 | -22 | 28 | -22 |
| 24 | | | | | |
| | L Frontal Pole | 2.99 | -20 | 54 | -14 |
| 24 | | | | | |
| | R Middle Temporal Gyrus, posterior division | 2.83 | 66 | -32 | -14 |
| 24 | | | | | |
| | R Hippocampus | 2.60 | 36 | -20 | -12 |
| 24 | | | | | |
| | L Lateral Occipital Cortex, inferior division | 3.23 | -40 | -74 | 2 |
| 24 | | | | | |
| | L Middle Temporal Gyrus, temporooccipital part | 2.54 | -56 | -54 | 8 |
| 24 | | | | | |

| | | | | | |
|---|---|---|---|---|---|
| | L Putamen | 3.15 | -30 | -14 | 8 |
| 24 | | | | | |
| | L Middle Frontal Gyrus | 2.73 | -42 | 20 | 36 |
| 24 | | | | | |
| | R Precuneus Cortex | 2.68 | 10 | -70 | 52 |
| 24 | | | | | |
| | R Precuneus Cortex | 2.66 | 10 | -58 | 58 |
| 16 | | | | | |
| | R Brain-Stem | 2.46 | 6 | -44 | -46 |
| 16 | | | | | |
| | L Temporal Pole | 2.93 | -38 | 8 | -44 |
| 16 | | | | | |
| | R Brain-Stem | 2.86 | 6 | -28 | -42 |
| 16 | | | | | |
| | R Inferior Temporal Gyrus, anterior division | 2.67 | 48 | 0 | -34 |
| 16 | | | | | |
| | L Hippocampus | 2.74 | -28 | -8 | -24 |
| 16 | | | | | |
| | R Inferior Temporal Gyrus, posterior division | 2.66 | 50 | -32 | -22 |
| 16 | | | | | |
| | L Inferior Temporal Gyrus, posterior division | 2.55 | -56 | -34 | -18 |
| 16 | | | | | |
| | R Middle Temporal Gyrus, posterior division | 2.71 | 70 | -22 | -10 |
| 16 | | | | | |
| | L Occipital Pole | 2.59 | -12 | -102 | -4 |
| 16 | | | | | |
| | L Occipital Pole | 3.04 | -20 | -106 | -2 |
| 16 | | | | | |
| | R Lateral Occipital Cortex, inferior division | 2.69 | 58 | -66 | 4 |
| 16 | | | | | |
| | L Middle Temporal Gyrus, temporooccipital part | 2.59 | -54 | -60 | 4 |
| 16 | | | | | |
| | L Frontal Pole | 2.73 | -44 | 36 | 4 |
| 16 | | | | | |
| | R Frontal Pole | 2.48 | 34 | 56 | 8 |
| 16 | | | | | |
| | L Frontal Pole | 2.51 | -36 | 60 | 12 |
| 16 | | | | | |
| | R Frontal Pole | 2.60 | 24 | 50 | 14 |
| 16 | | | | | |

| | | | | | |
|---|---|---|---|---|---|
| | R Frontal Pole | 2.64 | 38 | 42 | 18 |
| 16 | | | | | |
| | R Middle Frontal Gyrus | 2.89 | 42 | 32 | 28 |
| 16 | | | | | |
| | L Precuneus Cortex | 2.61 | -10 | -46 | 46 |
| 8 | | | | | |
| | L Temporal Fusiform Cortex, anterior division | 2.65 | -36 | 0 | -46 |
| 8 | | | | | |
| | R Inferior Temporal Gyrus, anterior division | 2.81 | 46 | 4 | -46 |
| 8 | | | | | |
| | R Temporal Fusiform Cortex, posterior division | 2.43 | 30 | -10 | -44 |
| 8 | | | | | |
| | R Brain-Stem | 2.58 | 12 | -24 | -42 |
| 8 | | | | | |
| | L Temporal Pole | 2.44 | -46 | 4 | -42 |
| 8 | | | | | |
| | L Brain-Stem | 2.82 | -4 | -20 | -36 |
| 8 | | | | | |
| | R Parahippocampal Gyrus, anterior division | 2.45 | 22 | -8 | -36 |
| 8 | | | | | |
| | L Inferior Temporal Gyrus, anterior division | 2.67 | -56 | -6 | -36 |
| 8 | | | | | |
| | R Temporal Pole | 2.50 | 54 | 10 | -36 |
| 8 | | | | | |
| | L Brain-Stem | 2.90 | -18 | -24 | -34 |
| 8 | | | | | |
| | R Inferior Temporal Gyrus, posterior division | 2.50 | 58 | -34 | -28 |
| 8 | | | | | |
| | L Inferior Temporal Gyrus, posterior division | 2.60 | -54 | -32 | -28 |
| 8 | | | | | |
| | L Inferior Temporal Gyrus, posterior division | 2.77 | -58 | -28 | -20 |
| 8 | | | | | |
| | L Middle Temporal Gyrus, posterior division | 2.49 | -60 | -20 | -20 |
| 8 | | | | | |
| | L Frontal Orbital Cortex | 2.69 | -14 | 32 | -20 |
| 8 | | | | | |
| | L Frontal Orbital Cortex | 2.69 | -20 | 20 | -18 |

| | | | | | |
|---|---|---|---|---|---|
| 8 | | | | | |
| | R Frontal Pole | 2.89 | 26 | 58 | -18 |
| 8 | | | | | |
| | L Frontal Pole | 2.85 | -38 | 56 | -16 |
| 8 | | | | | |
| | R Middle Temporal Gyrus, posterior division | 2.61 | 62 | -34 | -14 |
| 8 | | | | | |
| | L Amygdala | 2.43 | -18 | -10 | -14 |
| 8 | | | | | |
| | L Hippocampus | 2.67 | -34 | -20 | -12 |
| 8 | | | | | |
| | R Lateral Occipital Cortex, inferior division | 2.50 | 40 | -78 | -10 |
| 8 | | | | | |
| | L Occipital Pole | 2.45 | -28 | -92 | -2 |
| 8 | | | | | |
| | L Thalamus | 2.60 | -12 | -26 | 0 |
| 8 | | | | | |
| | R Pallidum | 2.58 | 22 | -14 | 0 |
| 8 | | | | | |
| | L Hippocampus | 2.49 | -24 | -40 | 4 |
| 8 | | | | | |
| | R Thalamus | 2.60 | 22 | -28 | 8 |
| 8 | | | | | |
| | L Caudate | 2.55 | -18 | 14 | 10 |
| 8 | | | | | |
| | R Central Opercular Cortex | 2.51 | 44 | -4 | 14 |
| 8 | | | | | |
| | L Frontal Pole | 2.73 | -34 | 58 | 22 |
| 8 | | | | | |
| | L Paracingulate Gyrus | 2.47 | -4 | 36 | 28 |
| 8 | | | | | |
| | R Postcentral Gyrus | 2.62 | 54 | -14 | 32 |
| 8 | | | | | |
| | L Paracingulate Gyrus | 2.43 | -2 | 34 | 32 |
| 8 | | | | | |
| | L Middle Frontal Gyrus | 2.56 | -40 | 24 | 34 |
| 8 | | | | | |
| | R Middle Frontal Gyrus | 2.68 | 38 | 26 | 34 |
| 8 | | | | | |
| | R Superior Frontal Gyrus | 2.74 | 6 | 36 | 50 |
| 8 | | | | | |
| | R Superior Frontal Gyrus | 2.46 | 18 | 16 | 66 |

**Supplementary Table S26. [$UT_{CB3}$ > $UT_{CB2}$].** Descriptive statistics for clusters and local maxima showing greater activation for Convolved Block 3 relative to Convolved Block 2 of Uncertain-Threat anticipation (FDR *q*<0.05, whole-brain corrected).

| ***mm³*** | **Label** | ***t*** | ***x*** | ***y*** | ***z*** |
|---|---|---|---|---|---|
| 63,520 | | | | | |
| | L Paracingulate Gyrus | 3.17 | -4 | 48 | 14 |
| | R Paracingulate Gyrus | 4.97 | 0 | 38 | 28 |
| | L Cingulate Gyrus, anterior division | 5.71 | -6 | 10 | 40 |
| | R Cingulate Gyrus, anterior division | 8.59 | 2 | 8 | 44 |
| | L Juxtapositional Lobule Cortex (formerly Supplementary Motor Cortex) | 5.12 | -8 | 2 | 46 |
| | L Superior Frontal Gyrus | 5.75 | -20 | -2 | 70 |
| | R Middle Frontal Gyrus | 4.40 | 36 | -4 | 56 |
| | R Superior Frontal Gyrus | 7.95 | 22 | -8 | 70 |
| | R Juxtapositional Lobule Cortex (formerly Supplementary Motor Cortex) | 8.41 | 4 | -12 | 48 |
| | L Cingulate Gyrus, posterior division | 7.57 | -12 | -22 | 40 |
| | R Cingulate Gyrus, posterior division | 9.91 | 12 | -24 | 40 |
| | R Precentral Gyrus | 7.75 | 4 | -24 | 50 |
| | R Postcentral Gyrus | 5.77 | 28 | -36 | 68 |
| | R Superior Parietal Lobule | 6.47 | 26 | -42 | 66 |
| | L Postcentral Gyrus | 2.85 | -10 | -46 | 66 |
| | L Precuneus Cortex | 5.31 | -4 | -48 | 62 |
| | R Precuneus Cortex | 6.04 | 8 | -48 | 58 |
| | L Superior Parietal Lobule | 4.12 | -16 | -54 | 60 |
| 42,320 | | | | | |
| | R Frontal Pole | 3.98 | 56 | 36 | -4 |
| | R Frontal Orbital Cortex | 4.28 | 46 | 30 | -4 |
| | R Inferior Frontal Gyrus, pars triangularis | 4.97 | 48 | 24 | 6 |
| | R Frontal Operculum Cortex | 6.50 | 42 | 18 | 4 |
| | R Inferior Frontal Gyrus, pars opercularis | 4.60 | 60 | 14 | 14 |
| | R Temporal Pole | 7.70 | 50 | 10 | -4 |
| | R Precentral Gyrus | 6.11 | 64 | 8 | 10 |
| | R Planum Polare | 8.70 | 58 | 2 | 2 |
| | R Central Opercular Cortex | 7.46 | 58 | 0 | 8 |
| | R Pallidum | 2.97 | 22 | 0 | -2 |
| | R Putamen | 5.87 | 32 | -6 | -4 |
| | R Insular Cortex | 6.26 | 42 | -8 | -6 |
| | R Middle Temporal Gyrus, posterior division | 3.81 | 58 | -22 | -10 |
| | R Planum Temporale | 6.38 | 56 | -24 | 14 |
| | R Heschls Gyrus (includes H1 and H2) | 5.34 | 46 | -24 | 14 |
| | R Superior Temporal Gyrus, posterior division | 6.17 | 70 | -26 | 6 |
| | R Parietal Operculum Cortex | 10.23 | 52 | -30 | 24 |

| | | | | | |
|---|---|---|---|---|---|
| | R Supramarginal Gyrus, anterior division | 7.18 | 56 | -32 | 34 |
| | R Supramarginal Gyrus, posterior division | 6.69 | 58 | -44 | 32 |
| | R Angular Gyrus | 3.77 | 46 | -50 | 40 |
| 26,904 | | | | | |
| | L Frontal Orbital Cortex | 4.34 | -38 | 24 | -6 |
| | L Frontal Operculum Cortex | 5.70 | -46 | 16 | -4 |
| | L Inferior Frontal Gyrus, pars opercularis | 5.74 | -56 | 12 | 0 |
| | L Precentral Gyrus | 6.23 | -60 | 8 | 6 |
| | L Central Opercular Cortex | 7.21 | -52 | 2 | 2 |
| | L Heschls Gyrus (includes H1 and H2) | 5.06 | -50 | -16 | 8 |
| | L Planum Polare | 5.01 | -40 | -20 | 0 |
| | L Postcentral Gyrus | 5.48 | -64 | -22 | 18 |
| | L Insular Cortex | 4.56 | -38 | -22 | 2 |
| | L Planum Temporale | 6.20 | -58 | -24 | 12 |
| | L Parietal Operculum Cortex | 6.78 | -62 | -32 | 20 |
| | L Supramarginal Gyrus, anterior division | 6.91 | -66 | -34 | 34 |
| | L Supramarginal Gyrus, posterior division | 5.89 | -58 | -42 | 24 |
| | L Angular Gyrus | 3.07 | -58 | -54 | 40 |
| 16,584 | | | | | |
| | R Middle Temporal Gyrus, temporooccipital part | 5.14 | 50 | -56 | 4 |
| | R Inferior Temporal Gyrus, temporooccipital part | 5.26 | 46 | -58 | -14 |
| | R Occipital Fusiform Gyrus | 5.23 | 38 | -68 | -14 |
| | R Lateral Occipital Cortex, inferior division | 9.79 | 46 | -76 | -10 |
| | R Lateral Occipital Cortex, superior division | 3.51 | 30 | -86 | 12 |
| 6,576 | | | | | |
| | L Lateral Occipital Cortex, inferior division | 6.37 | -36 | -90 | -8 |
| | L Occipital Pole | 6.53 | -38 | -92 | -10 |
| 1,640 | | | | | |
| | L Precentral Gyrus | 5.22 | -48 | -6 | 54 |
| 1,512 | | | | | |
| | R Caudate | 4.45 | 14 | 4 | 22 |
| | R Thalamus | 5.29 | 10 | -8 | 6 |
| 736 | | | | | |
| | R Frontal Pole | 4.83 | 10 | 62 | 28 |
| | R Superior Frontal Gyrus | 3.43 | 6 | 56 | 20 |
| 632 | | | | | |
| | R Frontal Pole | 3.71 | 34 | 40 | 40 |
| | R Middle Frontal Gyrus | 2.89 | 32 | 34 | 36 |
| 552 | | | | | |
| | R Superior Frontal Gyrus | 3.82 | 2 | 24 | 54 |
| 512 | | | | | |
| | L Frontal Pole | 4.77 | -36 | 40 | 38 |

| | | | | | |
|---|---|---|---|---|---|
| 456 | | | | | |
| | R Brain-Stem | 4.31 | 12 | -26 | -34 |
| 320 | | | | | |
| | L Thalamus | 4.79 | -12 | -24 | 18 |
| 248 | | | | | |
| | L Putamen | 3.49 | -24 | 0 | -2 |
| 208 | | | | | |
| | L Frontal Pole | 3.40 | -38 | 44 | 28 |
| 200 | | | | | |
| | L Putamen | 3.24 | -30 | -14 | -2 |
| 144 | | | | | |
| | R Caudate | 3.92 | 12 | 12 | 18 |
| 120 | | | | | |
| | R Inferior Temporal Gyrus, posterior division | 3.45 | 52 | -36 | -18 |
| | R Inferior Temporal Gyrus, temporooccipital part | 3.37 | 54 | -42 | -14 |
| 104 | | | | | |
| | R Brain-Stem | 3.01 | 2 | -16 | -36 |
| | L Brain-Stem | 2.91 | -2 | -18 | -38 |
| 104 | | | | | |
| | L Insular Cortex | 3.48 | -36 | -6 | -2 |
| | L Putamen | 2.92 | -32 | -8 | -4 |
| 96 | | | | | |
| | R Temporal Occipital Fusiform Cortex | 3.33 | 46 | -46 | -22 |
| 88 | | | | | |
| | R Frontal Orbital Cortex | 3.26 | 24 | 18 | -24 |
| 80 | | | | | |
| | R Brain-Stem | 4.24 | 12 | -16 | -32 |
| 80 | | | | | |
| | L Caudate | 3.56 | -12 | 22 | -4 |
| 80 | | | | | |
| | L Thalamus | 3.27 | -12 | -8 | 8 |
| 72 | | | | | |
| | L Frontal Orbital Cortex | 3.20 | -28 | 20 | -12 |
| 72 | | | | | |
| | L Middle Temporal Gyrus, temporooccipital part | 3.32 | -54 | -58 | 10 |
| 72 | | | | | |
| | R Frontal Pole | 2.87 | 26 | 54 | 28 |
| 72 | | | | | |
| | R Middle Frontal Gyrus | 3.12 | 34 | 28 | 40 |
| 64 | | | | | |
| | L Thalamus | 3.05 | -6 | -24 | 10 |
| 48 | | | | | |

| | | | | | |
|---|---|---|---|---|---|
| | R Temporal Pole | 3.42 | 40 | 24 | -36 |
| 48 | | | | | |
| | R Middle Temporal Gyrus, posterior division | 3.12 | 62 | -12 | -30 |
| | R Inferior Temporal Gyrus, posterior division | 2.94 | 60 | -16 | -30 |
| 48 | | | | | |
| | R Subcallosal Cortex | 3.39 | 4 | 30 | -8 |
| 40 | | | | | |
| | L Inferior Temporal Gyrus, posterior division | 3.44 | -50 | -20 | -30 |
| 40 | | | | | |
| | R Inferior Temporal Gyrus, temporooccipital part | 2.81 | 58 | -52 | -20 |
| 40 | | | | | |
| | L Parahippocampal Gyrus, posterior division | 3.01 | -16 | -30 | -14 |
| 40 | | | | | |
| | L Caudate | 3.23 | -16 | 16 | 14 |
| 40 | | | | | |
| | R Superior Frontal Gyrus | 2.98 | 6 | 38 | 50 |
| 32 | | | | | |
| | L Brain-Stem | 2.90 | -10 | -28 | -32 |
| 32 | | | | | |
| | R Inferior Temporal Gyrus, posterior division | 3.01 | 62 | -24 | -28 |
| 32 | | | | | |
| | L Hippocampus | 3.22 | -28 | -20 | -18 |
| 32 | | | | | |
| | R Inferior Temporal Gyrus, temporooccipital part | 3.66 | 62 | -50 | -18 |
| 32 | | | | | |
| | L Caudate | 2.79 | -14 | 4 | 22 |
| 32 | | | | | |
| | L Frontal Pole | 2.99 | -4 | 62 | 26 |
| 32 | | | | | |
| | R Frontal Pole | 2.94 | 38 | 52 | 28 |
| 32 | | | | | |
| | R Middle Frontal Gyrus | 3.33 | 40 | 16 | 44 |
| 24 | | | | | |
| | L Brain-Stem | 2.63 | -8 | -44 | -38 |
| 24 | | | | | |
| | R Inferior Temporal Gyrus, posterior division | 2.93 | 44 | -22 | -30 |
| 24 | | | | | |

| | | | | | |
|---|---|---|---|---|---|
| | R Inferior Temporal Gyrus, posterior division | 3.01 | 46 | -36 | -18 |
| 24 | | | | | |
| | R Middle Temporal Gyrus, temporooccipital part | 2.70 | 62 | -46 | 2 |
| 24 | | | | | |
| | R Cingulate Gyrus, anterior division | 3.64 | 12 | 42 | 4 |
| 24 | | | | | |
| | L Caudate | 2.85 | -8 | 16 | 8 |
| 24 | | | | | |
| | L Caudate | 3.88 | -14 | -12 | 22 |
| 24 | | | | | |
| | R Frontal Pole | 2.91 | 16 | 56 | 22 |
| 24 | | | | | |
| | R Cingulate Gyrus, posterior division | 2.83 | 0 | -18 | 28 |
| 24 | | | | | |
| | R Frontal Pole | 2.83 | 28 | 52 | 38 |
| 24 | | | | | |
| | L Superior Frontal Gyrus | 2.69 | -18 | 12 | 58 |
| 24 | | | | | |
| | L Superior Frontal Gyrus | 2.75 | -6 | 14 | 66 |
| 16 | | | | | |
| | R Brain-Stem | 2.86 | 14 | -28 | -40 |
| 16 | | | | | |
| | R Brain-Stem | 2.66 | 6 | -24 | -38 |
| 16 | | | | | |
| | L Temporal Pole | 2.77 | -28 | 22 | -36 |
| 16 | | | | | |
| | R Brain-Stem | 2.58 | 6 | -26 | -24 |
| 16 | | | | | |
| | L Subcallosal Cortex | 2.89 | -8 | 16 | -22 |
| 16 | | | | | |
| | R Inferior Temporal Gyrus, posterior division | 2.59 | 44 | -32 | -22 |
| 16 | | | | | |
| | R Frontal Orbital Cortex | 2.82 | 34 | 30 | -20 |
| 16 | | | | | |
| | R Amygdala | 2.80 | 22 | 2 | -20 |
| 16 | | | | | |
| | R Hippocampus | 2.81 | 38 | -22 | -14 |
| 16 | | | | | |
| | R Frontal Pole | 2.61 | 26 | 58 | -10 |
| 16 | | | | | |
| | R Frontal Pole | 2.67 | 32 | 52 | -10 |
| 16 | | | | | |

| | | | | | |
|---|---|---|---|---|---|
| | L Subcallosal Cortex | 2.97 | -2 | 26 | -6 |
| 16 | | | | | |
| | L Hippocampus | 3.04 | -32 | -36 | -4 |
| 16 | | | | | |
| | R Pallidum | 2.69 | 16 | 6 | 2 |
| 16 | | | | | |
| | L Caudate | 3.21 | -18 | 22 | 4 |
| 16 | | | | | |
| | L Caudate | 2.65 | -8 | 0 | 10 |
| 16 | | | | | |
| | R Caudate | 2.67 | 16 | 18 | 14 |
| 16 | | | | | |
| | L Caudate | 3.13 | -16 | -18 | 22 |
| 16 | | | | | |
| | L Paracingulate Gyrus | 2.68 | -10 | 44 | 24 |
| 16 | | | | | |
| | R Precentral Gyrus | 2.94 | 60 | 4 | 26 |
| 16 | | | | | |
| | R Cingulate Gyrus, anterior division | 2.70 | 2 | -10 | 30 |
| 8 | | | | | |
| | R Brain-Stem | 2.53 | 4 | -26 | -34 |
| 8 | | | | | |
| | L Parahippocampal Gyrus, anterior division | 3.13 | -22 | -14 | -34 |
| 8 | | | | | |
| | L Brain-Stem | 3.08 | -6 | -34 | -28 |
| 8 | | | | | |
| | L Hippocampus/Amygdala | 2.55 | -28 | -6 | -24 |
| 8 | | | | | |
| | R Temporal Pole | 2.57 | 46 | 10 | -24 |
| 8 | | | | | |
| | L Frontal Orbital Cortex | 2.85 | -14 | 14 | -24 |
| 8 | | | | | |
| | R Temporal Pole | 2.76 | 50 | 18 | -18 |
| 8 | | | | | |
| | L Frontal Orbital Cortex | 2.87 | -36 | 22 | -18 |
| 8 | | | | | |
| | R Temporal Occipital Fusiform Cortex | 2.59 | 38 | -48 | -16 |
| 8 | | | | | |
| | L Frontal Orbital Cortex | 2.92 | -28 | 36 | -16 |
| 8 | | | | | |
| | R Brain-Stem | 2.62 | 4 | -38 | -12 |
| 8 | | | | | |
| | L Frontal Orbital Cortex | 2.55 | -28 | 36 | -10 |

| | | | | | |
|---|---|---|---|---|---|
| 8 | | | | | |
| | R Middle Temporal Gyrus, posterior division | 2.82 | 48 | -28 | -6 |
| 8 | | | | | |
| | L Cingulate Gyrus, anterior division | 2.66 | -2 | 36 | -6 |
| 8 | | | | | |
| | L Lingual Gyrus | 2.58 | -4 | -86 | -4 |
| 8 | | | | | |
| | L Pallidum | 2.55 | -12 | -2 | -4 |
| 8 | | | | | |
| | R Caudate | 2.95 | 14 | 24 | -4 |
| 8 | | | | | |
| | R Lingual Gyrus | 2.61 | 22 | -46 | -2 |
| 8 | | | | | |
| | R Accumbens | 2.74 | 8 | 20 | -2 |
| 8 | | | | | |
| | R Putamen | 2.63 | 28 | 6 | 0 |
| 8 | | | | | |
| | L Frontal Pole | 2.54 | -50 | 38 | 0 |
| 8 | | | | | |
| | L Thalamus | 2.57 | -12 | -10 | 2 |
| 8 | | | | | |
| | R Middle Temporal Gyrus, temporooccipital part | 2.59 | 52 | -44 | 4 |
| 8 | | | | | |
| | L Frontal Pole | 2.59 | -44 | 36 | 4 |
| 8 | | | | | |
| | R Thalamus | 2.93 | 0 | -8 | 6 |
| 8 | | | | | |
| | R Caudate | 2.55 | 16 | 14 | 6 |
| 8 | | | | | |
| | R Thalamus | 2.54 | 8 | -22 | 10 |
| 8 | | | | | |
| | L Thalamus | 2.54 | -2 | -20 | 14 |
| 8 | | | | | |
| | R Paracingulate Gyrus | 2.56 | 10 | 46 | 20 |
| 8 | | | | | |
| | R Caudate | 2.76 | 14 | -10 | 22 |
| 8 | | | | | |
| | R Inferior Frontal Gyrus, pars opercularis | 2.87 | 46 | 14 | 22 |
| 8 | | | | | |
| | R Caudate | 2.64 | 16 | -14 | 24 |
| 8 | | | | | |
| | R Paracingulate Gyrus | 2.57 | 2 | 48 | 24 |
| 8 | | | | | |

| | | | | | |
|---|---|---|---|---|---|
| | L Frontal Pole | 2.75 | -2 | 58 | 24 |
| 8 | | | | | |
| | R Superior Frontal Gyrus | 2.58 | 6 | 44 | 38 |
| 8 | | | | | |
| | R Frontal Pole | 2.77 | 8 | 54 | 38 |
| 8 | | | | | |
| | L Precentral Gyrus | 2.67 | -46 | -4 | 40 |
| 8 | | | | | |
| | R Superior Frontal Gyrus | 2.58 | 12 | 16 | 60 |
| 8 | | | | | |
| | L Superior Frontal Gyrus | 2.53 | -22 | 14 | 62 |
| 8 | | | | | |
| | R Superior Frontal Gyrus | 2.60 | 6 | 28 | 64 |
| 8 | | | | | |
| | R Precentral Gyrus | 2.77 | 14 | -20 | 70 |
| 8 | | | | | |
| | L Superior Frontal Gyrus | 2.71 | -10 | 8 | 70 |

**Supplementary Table S27. [$CT_{CB3}$ - $CT_{CB2}$] > [$UT_{CB3}$ - $UT_{CB2}$].** Descriptive statistics for clusters and local maxima showing greater increases in activation from Convolved Block 2 to Convolved Block 3 of Certain-Threat anticipation relative to increases in activation from Convolved Block 2 to Convolved Block 3 of Uncertain-Threat anticipation (FDR $q$<0.05, whole-brain corrected).

| *mm³* | Label | *t* | *x* | *y* | *z* |
|---|---|---|---|---|---|
| 29,952 | | | | | |
| | L Insular Cortex | 7.11 | -32 | 18 | 8 |
| | L Frontal Operculum Cortex | 6.36 | -38 | 16 | 2 |
| | L Cingulate Gyrus, anterior division | 6.02 | -8 | 14 | 36 |
| | R Paracingulate Gyrus | 4.56 | 6 | 14 | 50 |
| | L Paracingulate Gyrus | 5.58 | -4 | 12 | 46 |
| | L Inferior Frontal Gyrus, pars opercularis | 3.44 | -52 | 10 | 4 |
| | R Cingulate Gyrus, anterior division | 5.25 | 10 | 10 | 40 |
| | L Precentral Gyrus | 6.34 | -58 | 8 | 14 |
| | L Superior Frontal Gyrus | 3.65 | -14 | 8 | 68 |
| | L Juxtapositional Lobule Cortex (formerly Supplementary Motor Cortex) | 6.40 | -6 | 4 | 56 |
| | R Juxtapositional Lobule Cortex (formerly Supplementary Motor Cortex) | 6.56 | 6 | 4 | 58 |
| | R Superior Frontal Gyrus | 3.76 | 14 | 2 | 68 |
| | L Postcentral Gyrus | 5.09 | -46 | -10 | 30 |
| | R Postcentral Gyrus | 8.36 | 36 | -26 | 56 |
| | R Precentral Gyrus | 5.66 | 26 | -26 | 66 |
| 10,416 | | | | | |
| | R Putamen | 10.29 | 22 | 8 | 0 |
| | R Caudate | 6.61 | 8 | 8 | 2 |
| | R Accumbens | 4.24 | 8 | 6 | -8 |
| | R Thalamus | 5.16 | 6 | -4 | -2 |
| | R Amygdala | 7.18 | 28 | -10 | -12 |
| | R Pallidum | 3.00 | 24 | -10 | 6 |
| 9,240 | | | | | |
| | R Occipital Fusiform Gyrus | 3.10 | 22 | -82 | -12 |
| | R Lingual Gyrus | 6.34 | 10 | -86 | -8 |
| | R Occipital Pole | 10.61 | 16 | -96 | 2 |
| 9,008 | | | | | |
| | L Caudate | 6.96 | -8 | 8 | 0 |
| | L Putamen | 7.88 | -22 | 0 | 2 |
| | L Amygdala | 7.77 | -26 | -8 | -12 |
| | L Pallidum | 4.87 | -24 | -10 | 0 |
| 6,376 | | | | | |
| | L Inferior Temporal Gyrus, temporooccipital part | 3.28 | -48 | -46 | -28 |
| | L Temporal Occipital Fusiform Cortex | 3.59 | -30 | -54 | -20 |
| | L Occipital Fusiform Gyrus | 3.75 | -24 | -70 | -18 |
| 3,904 | | | | | |

| | | | | | |
|---|---|---|---|---|---|
| | L Thalamus | 5.49 | -4 | -22 | 0 |
| | R Brain-Stem | 5.18 | 10 | -26 | -20 |
| | L Brain-Stem | 6.36 | -6 | -34 | -8 |
| 3,408 | | | | | |
| | R Temporal Occipital Fusiform Cortex | 4.90 | 34 | -60 | -20 |
| | R Occipital Fusiform Gyrus | 3.70 | 30 | -66 | -18 |
| 3,224 | | | | | |
| | L Frontal Pole | 5.57 | -36 | 48 | 30 |
| | L Middle Frontal Gyrus | 4.32 | -40 | 36 | 30 |
| 2,152 | | | | | |
| | L Occipital Fusiform Gyrus | 5.31 | -22 | -84 | -10 |
| | L Lingual Gyrus | 5.54 | -8 | -86 | -8 |
| | L Occipital Pole | 5.78 | -12 | -92 | -10 |
| 2,048 | | | | | |
| | R Frontal Operculum Cortex | 5.69 | 40 | 24 | 4 |
| | R Frontal Orbital Cortex | 4.55 | 36 | 24 | -4 |
| | R Central Opercular Cortex | 2.89 | 40 | 8 | 6 |
| 1,336 | | | | | |
| | R Thalamus | 6.41 | 4 | -22 | -2 |
| 1,312 | | | | | |
| | L Precuneus Cortex | 5.74 | -8 | -74 | 42 |
| | L Lateral Occipital Cortex, superior division | 3.73 | -8 | -74 | 54 |
| | L Angular Gyrus | 4.05 | -36 | -56 | 38 |
| | L Superior Parietal Lobule | 3.86 | -36 | -56 | 52 |
| | L Lateral Occipital Cortex, superior division | 3.88 | -34 | -60 | 50 |
| 1,144 | | | | | |
| | L Frontal Pole | 4.96 | -16 | 68 | -6 |
| 1,112 | | | | | |
| | R Precuneus Cortex | 6.02 | 12 | -70 | 40 |
| 872 | | | | | |
| | R Frontal Pole | 4.48 | 36 | 46 | 32 |
| 760 | | | | | |
| | R Frontal Pole | 4.69 | 20 | 70 | -8 |
| 640 | | | | | |
| | L Angular Gyrus | 4.62 | -46 | -56 | 46 |
| | L Lateral Occipital Cortex, superior division | 3.91 | -48 | -60 | 40 |
| 632 | | | | | |
| | R Precentral Gyrus | 4.22 | 64 | 8 | 16 |
| 400 | | | | | |
| | L Heschls Gyrus (includes H1 and H2) | 4.82 | -46 | -26 | 6 |
| | L Planum Temporale | 3.77 | -50 | -30 | 8 |
| 400 | | | | | |

| | | | | | |
|---|---|---|---|---|---|
| | R Precentral Gyrus | 3.99 | 48 | -8 | 32 |
| | R Postcentral Gyrus | 4.26 | 54 | -12 | 38 |
| 400 | | | | | |
| | L Supramarginal Gyrus, posterior division | 3.82 | -54 | -48 | 50 |
| 328 | | | | | |
| | R Brain-Stem | 4.01 | 8 | -40 | -52 |
| | L Brain-Stem | 4.02 | -4 | -46 | -58 |
| 312 | | | | | |
| | L Parietal Operculum Cortex | 4.10 | -56 | -32 | 26 |
| | L Supramarginal Gyrus, anterior division | 3.80 | -62 | -34 | 24 |
| 296 | | | | | |
| | L Brain-Stem | 5.09 | -4 | -40 | -48 |
| 232 | | | | | |
| | R Heschls Gyrus (includes H1 and H2) | 4.41 | 50 | -18 | 8 |
| | R Planum Temporale | 3.46 | 60 | -18 | 6 |
| 224 | | | | | |
| | L Postcentral Gyrus | 3.94 | -60 | -22 | 30 |
| | L Supramarginal Gyrus, anterior division | 3.44 | -62 | -24 | 26 |
| 192 | | | | | |
| | R Brain-Stem | 4.62 | 2 | -36 | -38 |
| 192 | | | | | |
| | R Occipital Fusiform Gyrus | 3.07 | 24 | -72 | -16 |
| 192 | | | | | |
| | R Superior Parietal Lobule | 4.03 | 26 | -42 | 62 |
| 168 | | | | | |
| | L Cingulate Gyrus, posterior division | 3.77 | -4 | -22 | 28 |
| 136 | | | | | |
| | L Inferior Temporal Gyrus, anterior division | 4.26 | -48 | -8 | -38 |
| 136 | | | | | |
| | L Supramarginal Gyrus, posterior division | 3.64 | -60 | -50 | 42 |
| 120 | | | | | |
| | L Precuneus Cortex | 3.27 | -4 | -64 | 58 |
| | R Precuneus Cortex | 3.24 | 0 | -64 | 58 |
| 112 | | | | | |
| | R Cingulate Gyrus, posterior division | 4.12 | 4 | -22 | 30 |
| 112 | | | | | |
| | L Precentral Gyrus | 4.95 | -8 | -24 | 58 |
| 112 | | | | | |
| | L Precentral Gyrus | 4.46 | -6 | -22 | 66 |
| 104 | | | | | |
| | L Superior Frontal Gyrus | 3.81 | -22 | 2 | 60 |
| 80 | | | | | |
| | L Insular Cortex | 3.48 | -28 | 20 | -6 |

| | | | | | |
|---|---|---|---|---|---|
| 80 | | | | | |
| | R Frontal Pole | 3.45 | 30 | 40 | 36 |
| 80 | | | | | |
| | L Supramarginal Gyrus, anterior division | 3.42 | -38 | -38 | 38 |
| 72 | | | | | |
| | R Central Opercular Cortex | 3.87 | 52 | 4 | 4 |
| 72 | | | | | |
| | L Middle Frontal Gyrus | 3.89 | -42 | 20 | 36 |
| 72 | | | | | |
| | R Precuneus Cortex | 3.24 | 0 | -46 | 42 |
| 56 | | | | | |
| | R Middle Temporal Gyrus, anterior division | 3.24 | 56 | -4 | -34 |
| 56 | | | | | |
| | L Brain-Stem | 3.72 | -4 | -14 | -24 |
| 56 | | | | | |
| | L Putamen | 3.74 | -30 | -18 | -2 |
| 56 | | | | | |
| | R Precentral Gyrus | 3.77 | 6 | -22 | 66 |
| 48 | | | | | |
| | L Frontal Pole | 4.03 | -20 | 52 | -14 |
| 48 | | | | | |
| | R Frontal Pole | 3.26 | 2 | 70 | -4 |
| 48 | | | | | |
| | L Postcentral Gyrus | 2.95 | -58 | -10 | 18 |
| 48 | | | | | |
| | L Cingulate Gyrus, posterior division | 4.17 | -6 | -38 | 26 |
| 48 | | | | | |
| | R Precentral Gyrus | 3.17 | 8 | -20 | 50 |
| 48 | | | | | |
| | L Precuneus Cortex | 2.91 | -2 | -58 | 66 |
| 40 | | | | | |
| | L Occipital Pole | 3.55 | -12 | -94 | 0 |
| 40 | | | | | |
| | R Parietal Operculum Cortex | 2.97 | 56 | -28 | 22 |
| 40 | | | | | |
| | R Supramarginal Gyrus, anterior division | 3.63 | 60 | -22 | 24 |
| 40 | | | | | |
| | L Juxtapositional Lobule Cortex (formerly Supplementary Motor Cortex) | 3.32 | -6 | -10 | 48 |
| 32 | | | | | |
| | R Temporal Fusiform Cortex, posterior division | 2.99 | 34 | -14 | -32 |
| 32 | | | | | |

| | | | | | |
|---|---|---|---|---|---|
| | L Inferior Temporal Gyrus, posterior division | 3.51 | -64 | -22 | -28 |
| 32 | | | | | |
| | L Inferior Temporal Gyrus, posterior division | 3.12 | -56 | -44 | -24 |
| | L Inferior Temporal Gyrus, temporooccipital part | 3.17 | -54 | -46 | -26 |
| 32 | | | | | |
| | L Middle Temporal Gyrus, posterior division | 3.00 | -58 | -26 | -12 |
| 32 | | | | | |
| | L Frontal Pole | 3.59 | -2 | 70 | -2 |
| 32 | | | | | |
| | R Putamen | 3.34 | 30 | -12 | 6 |
| 32 | | | | | |
| | L Planum Temporale | 3.21 | -60 | -28 | 10 |
| 32 | | | | | |
| | R Frontal Pole | 3.06 | 40 | 56 | 16 |
| 32 | | | | | |
| | L Central Opercular Cortex | 3.09 | -44 | -22 | 16 |
| 32 | | | | | |
| | R Frontal Pole | 3.55 | 32 | 36 | 26 |
| 32 | | | | | |
| | R Supramarginal Gyrus, posterior division | 3.24 | 66 | -38 | 34 |
| 32 | | | | | |
| | L Postcentral Gyrus | 3.44 | -34 | -28 | 50 |
| 24 | | | | | |
| | L Brain-Stem | 3.21 | -2 | -32 | -44 |
| 24 | | | | | |
| | R Middle Temporal Gyrus, anterior division | 2.98 | 50 | 0 | -34 |
| 24 | | | | | |
| | L Inferior Temporal Gyrus, posterior division | 3.02 | -50 | -34 | -28 |
| 24 | | | | | |
| | R Inferior Temporal Gyrus, temporooccipital part | 2.85 | 54 | -46 | -26 |
| 24 | | | | | |
| | L Middle Temporal Gyrus, posterior division | 2.91 | -66 | -24 | -18 |
| 24 | | | | | |
| | L Middle Temporal Gyrus, posterior division | 3.13 | -70 | -28 | -12 |
| 24 | | | | | |
| | R Planum Temporale | 3.30 | 58 | -24 | 8 |
| 24 | | | | | |

| | | | | | |
|---|---|---|---|---|---|
| | L Central Opercular Cortex | 3.26 | -52 | -2 | 8 |
| 24 | | | | | |
| | L Precentral Gyrus | 3.28 | -58 | 0 | 10 |
| 24 | | | | | |
| | L Central Opercular Cortex | 2.84 | -38 | 2 | 10 |
| 24 | | | | | |
| | L Supramarginal Gyrus, posterior division | 2.95 | -52 | -40 | 52 |
| 24 | | | | | |
| | R Precentral Gyrus | 3.16 | 32 | -6 | 54 |
| 24 | | | | | |
| | R Lateral Occipital Cortex, superior division | 2.88 | 10 | -72 | 62 |
| 16 | | | | | |
| | R Brain-Stem | 3.00 | 16 | -22 | -36 |
| 16 | | | | | |
| | R Brain-Stem | 2.86 | 10 | -38 | -28 |
| 16 | | | | | |
| | R Brain-Stem | 2.76 | 4 | -28 | -24 |
| 16 | | | | | |
| | L Inferior Temporal Gyrus, temporooccipital part | 2.97 | -58 | -50 | -22 |
| 16 | | | | | |
| | L Inferior Temporal Gyrus, posterior division | 2.89 | -50 | -40 | -22 |
| 16 | | | | | |
| | R Temporal Pole | 2.79 | 32 | 6 | -22 |
| 16 | | | | | |
| | L Middle Temporal Gyrus, posterior division | 3.29 | -70 | -20 | -14 |
| 16 | | | | | |
| | L Frontal Pole | 3.01 | -28 | 40 | -14 |
| 16 | | | | | |
| | L Planum Polare | 3.44 | -44 | -10 | -10 |
| 16 | | | | | |
| | R Insular Cortex | 3.00 | 38 | -8 | -10 |
| 16 | | | | | |
| | L Insular Cortex | 2.83 | -36 | 20 | -6 |
| 16 | | | | | |
| | L Occipital Pole | 3.14 | -8 | -98 | 6 |
| 16 | | | | | |
| | R Central Opercular Cortex | 3.25 | 44 | 6 | 8 |
| 16 | | | | | |
| | R Frontal Pole | 2.94 | 38 | 40 | 18 |
| 16 | | | | | |
| | R Caudate | 2.89 | 18 | 0 | 22 |

| | | | | | |
|---|---|---|---|---|---|
| 16 | | | | | |
| | R Angular Gyrus | 3.12 | 58 | -48 | 30 |
| 16 | | | | | |
| | R Cingulate Gyrus, anterior division | 3.27 | 4 | 0 | 32 |
| 16 | | | | | |
| | L Frontal Pole | 2.90 | -28 | 54 | 32 |
| 16 | | | | | |
| | R Frontal Pole | 3.01 | 24 | 48 | 36 |
| 16 | | | | | |
| | L Lateral Occipital Cortex, superior division | 2.88 | -56 | -64 | 38 |
| 16 | | | | | |
| | L Cingulate Gyrus, anterior division | 3.05 | -4 | -2 | 40 |
| 16 | | | | | |
| | R Postcentral Gyrus | 3.01 | 50 | -22 | 48 |
| 16 | | | | | |
| | L Postcentral Gyrus | 3.14 | -18 | -40 | 64 |
| 8 | | | | | |
| | L Temporal Fusiform Cortex, anterior division | 2.86 | -38 | -4 | -48 |
| 8 | | | | | |
| | R Brain-Stem | 3.03 | 4 | -46 | -46 |
| 8 | | | | | |
| | R Inferior Temporal Gyrus, anterior division | 2.95 | 50 | -2 | -46 |
| 8 | | | | | |
| | R Inferior Temporal Gyrus, anterior division | 3.04 | 46 | 4 | -46 |
| 8 | | | | | |
| | R Brain-Stem | 3.11 | 2 | -26 | -44 |
| 8 | | | | | |
| | R Inferior Temporal Gyrus, anterior division | 2.85 | 54 | -2 | -42 |
| 8 | | | | | |
| | L Brain-Stem | 2.87 | -4 | -24 | -40 |
| 8 | | | | | |
| | R Temporal Fusiform Cortex, posterior division | 2.80 | 36 | -18 | -38 |
| 8 | | | | | |
| | L Inferior Temporal Gyrus, posterior division | 2.74 | -54 | -16 | -36 |
| 8 | | | | | |
| | L Inferior Temporal Gyrus, anterior division | 2.74 | -56 | -6 | -36 |
| 8 | | | | | |

| | | | | | |
|---|---|---|---|---|---|
| | L Temporal Pole | 2.78 | -36 | 10 | -36 |
| 8 | | | | | |
| | L Parahippocampal Gyrus, anterior division | 2.86 | -24 | -20 | -32 |
| 8 | | | | | |
| | L Brain-Stem | 2.82 | -6 | -32 | -30 |
| 8 | | | | | |
| | L Hippocampus | 2.99 | -30 | -10 | -26 |
| 8 | | | | | |
| | L Brain-Stem | 2.99 | -4 | -30 | -24 |
| 8 | | | | | |
| | R Subcallosal Cortex | 3.48 | 0 | 30 | -24 |
| 8 | | | | | |
| | L Amygdala | 2.73 | -32 | -2 | -22 |
| 8 | | | | | |
| | L Frontal Orbital Cortex | 2.72 | -12 | 22 | -22 |
| 8 | | | | | |
| | R Occipital Fusiform Gyrus | 2.81 | 24 | -90 | -18 |
| 8 | | | | | |
| | L Frontal Pole | 3.20 | -38 | 56 | -16 |
| 8 | | | | | |
| | R Frontal Pole | 2.91 | 4 | 66 | -16 |
| 8 | | | | | |
| | R Occipital Fusiform Gyrus | 2.73 | 30 | -78 | -12 |
| 8 | | | | | |
| | L Planum Polare | 2.75 | -46 | -16 | -4 |
| 8 | | | | | |
| | R Frontal Pole | 2.72 | 10 | 72 | 0 |
| 8 | | | | | |
| | R Inferior Frontal Gyrus, pars opercularis | 2.89 | 54 | 10 | 4 |
| 8 | | | | | |
| | R Thalamus | 2.77 | 14 | -12 | 6 |
| 8 | | | | | |
| | L Thalamus | 2.77 | -10 | -10 | 6 |
| 8 | | | | | |
| | L Heschls Gyrus (includes H1 and H2) | 2.91 | -34 | -26 | 10 |
| 8 | | | | | |
| | R Heschls Gyrus (includes H1 and H2) | 2.84 | 38 | -24 | 10 |
| 8 | | | | | |
| | R Frontal Pole | 2.75 | 42 | 52 | 12 |
| 8 | | | | | |
| | L Planum Temporale | 2.74 | -58 | -32 | 14 |
| 8 | | | | | |
| | L Thalamus | 2.84 | -6 | -26 | 14 |

| | | | | | |
|---|---|---|---|---|---|
| 8 | | | | | |
| | L Caudate | 2.98 | -16 | 10 | 14 |
| 8 | | | | | |
| | R Frontal Pole | 2.72 | 24 | 50 | 14 |
| 8 | | | | | |
| | R Frontal Pole | 2.84 | 30 | 52 | 16 |
| 8 | | | | | |
| | L Planum Temporale | 3.08 | -48 | -38 | 18 |
| 8 | | | | | |
| | R Caudate | 2.78 | 12 | 8 | 20 |
| 8 | | | | | |
| | L Frontal Pole | 2.83 | -24 | 44 | 20 |
| 8 | | | | | |
| | L Postcentral Gyrus | 2.77 | -54 | -20 | 26 |
| 8 | | | | | |
| | R Precentral Gyrus | 2.77 | 60 | 4 | 32 |
| 8 | | | | | |
| | R Middle Frontal Gyrus | 2.73 | 32 | 32 | 32 |
| 8 | | | | | |
| | R Precuneus Cortex | 2.73 | 14 | -52 | 36 |
| 8 | | | | | |
| | R Precuneus Cortex | 2.71 | 6 | -46 | 40 |
| 8 | | | | | |
| | R Cingulate Gyrus, posterior division | 2.78 | 8 | -42 | 40 |
| 8 | | | | | |
| | L Supramarginal Gyrus, posterior division | 2.75 | -54 | -48 | 42 |
| 8 | | | | | |
| | R Postcentral Gyrus | 2.80 | 36 | -30 | 44 |
| 8 | | | | | |
| | R Lateral Occipital Cortex, superior division | 2.75 | 54 | -60 | 46 |
| 8 | | | | | |
| | R Superior Parietal Lobule | 2.84 | 40 | -48 | 50 |
| 8 | | | | | |
| | R Juxtapositional Lobule Cortex (formerly Supplementary Motor Cortex) | 2.83 | 10 | -10 | 50 |
| 8 | | | | | |
| | L Lateral Occipital Cortex, superior division | 2.71 | -26 | -80 | 52 |
| 8 | | | | | |
| | R Postcentral Gyrus | 2.93 | 20 | -42 | 66 |
| 8 | | | | | |
| | R Postcentral Gyrus | 2.98 | 4 | -34 | 68 |
| 8 | | | | | |
| | L Precentral Gyrus | 2.93 | -24 | -20 | 68 |

**Supplementary Table S28. [$CT_{CB2} > CT_{CB3}$].** Descriptive statistics for clusters and local maxima showing greater activation for Convolved Block 2 relative to Convolved Block 3 of Certain-Threat anticipation (FDR *q*<0.05, whole-brain corrected).

| ***mm³*** | **Label** | ***t*** | ***x*** | ***y*** | ***z*** |
|---|---|---|---|---|---|
| 132,784 | | | | | |
| | R Amygdala | 3.59 | 12 | -8 | -16 |
| | L Parahippocampal Gyrus, anterior division | 3.30 | -20 | -20 | -22 |
| | R Hippocampus | 4.54 | 22 | -20 | -16 |
| | L Postcentral Gyrus | 8.31 | -38 | -24 | 68 |
| | R Inferior Temporal Gyrus, posterior division | 3.15 | 44 | -24 | -24 |
| | L Hippocampus | 5.59 | -20 | -28 | -12 |
| | L Precentral Gyrus | 4.15 | -10 | -28 | 80 |
| | L Thalamus | 4.75 | -8 | -34 | 2 |
| | R Thalamus | 4.57 | 14 | -34 | -2 |
| | L Parahippocampal Gyrus, posterior division | 6.27 | -26 | -36 | -16 |
| | R Parahippocampal Gyrus, posterior division | 8.46 | 30 | -36 | -14 |
| | R Temporal Fusiform Cortex, posterior division | 7.52 | 30 | -38 | -22 |
| | R Lingual Gyrus | 9.30 | 32 | -40 | -10 |
| | L Temporal Fusiform Cortex, posterior division | 6.80 | -34 | -44 | -20 |
| | L Cingulate Gyrus, posterior division | 6.29 | -2 | -46 | 12 |
| | R Cingulate Gyrus, posterior division | 6.76 | 8 | -46 | 4 |
| | L Temporal Occipital Fusiform Cortex | 7.49 | -28 | -52 | -10 |
| | R Temporal Occipital Fusiform Cortex | 6.88 | 24 | -52 | -12 |
| | R Superior Parietal Lobule | 3.33 | 30 | -54 | 60 |
| | L Lingual Gyrus | 8.27 | -22 | -56 | 2 |
| | R Angular Gyrus | 3.62 | 52 | -56 | 28 |
| | L Superior Parietal Lobule | 5.44 | -24 | -58 | 62 |
| | L Middle Temporal Gyrus, temporooccipital part | 4.01 | -56 | -58 | -8 |
| | L Inferior Temporal Gyrus, temporooccipital part | 6.81 | -42 | -62 | -8 |
| | L Precuneus Cortex | 6.00 | -10 | -62 | 12 |
| | L Intracalcarine Cortex | 7.40 | -18 | -64 | 6 |
| | L Lateral Occipital Cortex, inferior division | 7.13 | -44 | -64 | -10 |
| | R Precuneus Cortex | 7.67 | 10 | -64 | 18 |
| | R Occipital Fusiform Gyrus | 5.62 | 22 | -66 | -12 |
| | L Occipital Fusiform Gyrus | 6.10 | -36 | -74 | -14 |
| | R Lateral Occipital Cortex, inferior division | 6.04 | 50 | -76 | 12 |

| | | | | | |
|---|---|---|---|---|---|
| | R Intracalcarine Cortex | 7.68 | 14 | -78 | 8 |
| | L Cuneal Cortex | 8.06 | -2 | -84 | 36 |
| | R Lateral Occipital Cortex, superior division | 6.25 | 34 | -84 | 32 |
| | R Cuneal Cortex | 8.62 | 6 | -86 | 38 |
| | R Supracalcarine Cortex | 7.29 | 2 | -86 | 12 |
| | L Lateral Occipital Cortex, superior division | 6.94 | -14 | -88 | 40 |
| | L Occipital Pole | 8.80 | -6 | -90 | 34 |
| | R Occipital Pole | 7.13 | 10 | -92 | 32 |
| 10,656 | | | | | |
| | L Frontal Orbital Cortex | 3.70 | -40 | 22 | -16 |
| | L Temporal Pole | 5.97 | -36 | 18 | -26 |
| | L Superior Temporal Gyrus, anterior division | 5.39 | -58 | 0 | -14 |
| | L Planum Polare | 2.80 | -56 | -2 | -2 |
| | L Middle Temporal Gyrus, anterior division | 5.01 | -54 | -4 | -18 |
| | L Superior Temporal Gyrus, posterior division | 4.04 | -56 | -24 | -4 |
| | L Middle Temporal Gyrus, posterior division | 3.78 | -68 | -42 | 2 |
| | L Middle Temporal Gyrus, temporooccipital part | 3.05 | -62 | -44 | 2 |
| 8,272 | | | | | |
| | R Frontal Pole | 6.56 | 50 | 36 | 18 |
| | R Middle Frontal Gyrus | 5.73 | 50 | 32 | 22 |
| | R Frontal Orbital Cortex | 3.48 | 48 | 28 | -8 |
| | R Inferior Frontal Gyrus, pars opercularis | 6.47 | 48 | 14 | 30 |
| 6,528 | | | | | |
| | R Frontal Orbital Cortex | 3.13 | 30 | 24 | -20 |
| | R Temporal Pole | 6.29 | 48 | 20 | -22 |
| | R Superior Temporal Gyrus, anterior division | 5.36 | 60 | 2 | -10 |
| | R Middle Temporal Gyrus, anterior division | 4.48 | 60 | -2 | -16 |
| | R Middle Temporal Gyrus, posterior division | 4.12 | 68 | -10 | -10 |
| | R Planum Temporale | 3.56 | 66 | -10 | 6 |
| 5,536 | | | | | |
| | R Frontal Pole | 4.81 | 12 | 46 | 50 |
| | R Superior Frontal Gyrus | 6.44 | 22 | 30 | 48 |
| | R Middle Frontal Gyrus | 4.77 | 36 | 16 | 56 |
| 4,904 | | | | | |
| | L Frontal Pole | 4.64 | -48 | 44 | -12 |
| | L Frontal Orbital Cortex | 4.33 | -48 | 34 | -8 |

| | | | | | |
|---|---|---|---|---|---|
| | L Frontal Operculum Cortex | 2.68 | -46 | 28 | -2 |
| | L Inferior Frontal Gyrus, pars triangularis | 3.70 | -50 | 26 | 24 |
| | L Middle Frontal Gyrus | 4.20 | -48 | 22 | 28 |
| | L Inferior Frontal Gyrus, pars opercularis | 4.20 | -46 | 8 | 28 |
| 3,768 | | | | | |
| | L Frontal Pole | 5.14 | -8 | 66 | 22 |
| | R Superior Frontal Gyrus | 2.91 | 0 | 42 | 42 |
| | L Superior Frontal Gyrus | 3.40 | -16 | 36 | 56 |
| 2,352 | | | | | |
| | L Frontal Pole | 2.97 | -2 | 60 | -14 |
| | R Frontal Pole | 4.66 | 8 | 60 | 0 |
| | R Frontal Medial Cortex | 3.71 | 10 | 54 | -8 |
| | L Frontal Medial Cortex | 4.45 | -8 | 46 | -10 |
| | L Paracingulate Gyrus | 4.19 | -10 | 46 | -6 |
| | R Paracingulate Gyrus | 2.99 | 6 | 42 | -8 |
| 1,880 | | | | | |
| | R Postcentral Gyrus | 4.77 | 58 | -18 | 38 |
| | R Supramarginal Gyrus, anterior division | 4.60 | 54 | -26 | 46 |
| 1,048 | | | | | |
| | L Superior Frontal Gyrus | 4.04 | -22 | 28 | 50 |
| | L Middle Frontal Gyrus | 3.08 | -28 | 18 | 56 |
| 664 | | | | | |
| | R Frontal Pole | 4.77 | 10 | 68 | 18 |
| 568 | | | | | |
| | L Temporal Pole | 3.18 | -24 | 4 | -26 |
| | L Amygdala | 3.35 | -22 | 2 | -24 |
| | L Amygdala | 3.61 | -28 | 0 | -28 |
| | L Parahippocampal Gyrus, anterior division | 4.16 | -18 | -12 | -28 |
| | L Hippocampus | 3.06 | -18 | -16 | -22 |
| 504 | | | | | |
| | R Precentral Gyrus | 4.73 | 6 | -24 | 80 |
| 472 | | | | | |
| | L Temporal Fusiform Cortex, posterior division | 5.13 | -40 | -14 | -28 |
| 456 | | | | | |
| | R Brain-Stem | 3.90 | 10 | -40 | -58 |
| 400 | | | | | |
| | R Supramarginal Gyrus, posterior division | 4.15 | 42 | -40 | 40 |
| 352 | | | | | |
| | R Middle Temporal Gyrus, temporooccipital part | 4.60 | 58 | -48 | -8 |
| 312 | | | | | |
| | R Subcallosal Cortex | 4.16 | 0 | 10 | -8 |
| 304 | | | | | |

| | | | | | |
|---|---|---|---|---|---|
| | L Postcentral Gyrus | 3.85 | -64 | -14 | 34 |
| 288 | | | | | |
| | L Brain-Stem | 3.81 | -8 | -36 | -60 |
| 240 | | | | | |
| | R Insular Cortex | 4.74 | 38 | -6 | 12 |
| 232 | | | | | |
| | R Paracingulate Gyrus | 3.70 | 6 | 46 | 22 |
| 192 | | | | | |
| | L Superior Frontal Gyrus | 4.30 | -4 | 18 | 62 |
| 176 | | | | | |
| | L Postcentral Gyrus | 3.59 | -60 | -26 | 42 |
| 168 | | | | | |
| | R Parahippocampal Gyrus, anterior division | 3.99 | 18 | -8 | -28 |
| 160 | | | | | |
| | R Middle Temporal Gyrus, posterior division | 4.23 | 48 | -36 | -2 |
| 160 | | | | | |
| | L Angular Gyrus | 3.89 | -44 | -58 | 24 |
| 152 | | | | | |
| | R Brain-Stem | 3.69 | 0 | -38 | -22 |
| 144 | | | | | |
| | R Brain-Stem | 4.20 | 2 | -46 | -52 |
| 104 | | | | | |
| | R Temporal Pole | 3.08 | 26 | 12 | -44 |
| 104 | | | | | |
| | L Frontal Pole | 3.60 | -20 | 60 | -8 |
| 96 | | | | | |
| | R Subcallosal Cortex | 4.24 | 4 | 30 | -20 |
| 96 | | | | | |
| | R Occipital Fusiform Gyrus | 3.44 | 38 | -64 | -12 |
| 96 | | | | | |
| | L Middle Frontal Gyrus | 3.47 | -40 | 12 | 46 |
| 96 | | | | | |
| | R Postcentral Gyrus | 3.04 | 6 | -42 | 72 |
| 88 | | | | | |
| | R Brain-Stem | 3.32 | 2 | -36 | -66 |
| 88 | | | | | |
| | R Planum Polare | 3.95 | 50 | 0 | -6 |
| 80 | | | | | |
| | R Brain-Stem | 3.67 | 16 | -32 | -28 |
| 80 | | | | | |
| | L Insular Cortex | 3.61 | -36 | -8 | 12 |
| 80 | | | | | |
| | L Angular Gyrus | 3.25 | -60 | -56 | 22 |

| | | | | | |
|---|---|---|---|---|---|
| 80 | | | | | |
| | R Superior Frontal Gyrus | 3.12 | 24 | 8 | 52 |
| 72 | | | | | |
| | R Middle Temporal Gyrus, temporooccipital part | 3.05 | 68 | -40 | -10 |
| 72 | | | | | |
| | L Insular Cortex | 2.93 | -40 | -6 | 2 |
| 72 | | | | | |
| | R Postcentral Gyrus | 3.16 | 20 | -30 | 78 |
| 64 | | | | | |
| | R Frontal Pole | 3.34 | 12 | 60 | -22 |
| 64 | | | | | |
| | R Paracingulate Gyrus | 2.86 | 10 | 44 | -4 |
| 64 | | | | | |
| | L Central Opercular Cortex | 3.03 | -56 | -14 | 12 |
| 64 | | | | | |
| | R Postcentral Gyrus | 3.75 | 68 | -10 | 24 |
| 64 | | | | | |
| | L Cingulate Gyrus, posterior division | 3.17 | -4 | -42 | 36 |
| 56 | | | | | |
| | R Frontal Orbital Cortex | 3.59 | 14 | 16 | -22 |
| 56 | | | | | |
| | L Hippocampus | 3.86 | -14 | -12 | -18 |
| 56 | | | | | |
| | L Subcallosal Cortex | 3.08 | -2 | 28 | -12 |
| 56 | | | | | |
| | L Frontal Pole | 3.89 | -46 | 44 | 8 |
| 56 | | | | | |
| | L Cingulate Gyrus, anterior division | 2.81 | -8 | 44 | 8 |
| 48 | | | | | |
| | L Brain-Stem | 3.49 | -4 | -42 | -26 |
| 48 | | | | | |
| | R Temporal Fusiform Cortex, posterior division | 3.16 | 42 | -12 | -26 |
| 48 | | | | | |
| | R Frontal Orbital Cortex | 2.90 | 16 | 8 | -22 |
| 48 | | | | | |
| | R Cingulate Gyrus, anterior division | 2.76 | 4 | 42 | 6 |
| 48 | | | | | |
| | L Paracingulate Gyrus | 2.68 | -2 | 50 | 10 |
| 48 | | | | | |
| | R Cingulate Gyrus, posterior division | 3.41 | 6 | -34 | 38 |
| 40 | | | | | |
| | L Frontal Orbital Cortex | 2.77 | -22 | 14 | -26 |
| 40 | | | | | |

| | | | | | |
|---|---|---|---|---|---|
| | L Frontal Pole | 2.82 | -8 | 58 | -24 |
| 40 | | | | | |
| | R Middle Temporal Gyrus, posterior division | 2.90 | 62 | -12 | -22 |
| 40 | | | | | |
| | R Paracingulate Gyrus | 3.25 | 10 | 48 | 8 |
| 40 | | | | | |
| | R Central Opercular Cortex | 3.04 | 64 | -12 | 12 |
| 40 | | | | | |
| | R Cingulate Gyrus, posterior division | 3.09 | 2 | -48 | 30 |
| 40 | | | | | |
| | L Frontal Pole | 2.67 | -18 | 42 | 36 |
| 32 | | | | | |
| | L Temporal Fusiform Cortex, posterior division | 3.22 | -40 | -16 | -36 |
| 32 | | | | | |
| | L Frontal Medial Cortex | 3.50 | -8 | 40 | -22 |
| 32 | | | | | |
| | L Planum Polare | 2.72 | -42 | 2 | -18 |
| 32 | | | | | |
| | R Temporal Pole | 3.87 | 48 | 8 | -12 |
| 32 | | | | | |
| | R Subcallosal Cortex | 3.11 | 4 | 20 | -10 |
| 24 | | | | | |
| | L Temporal Fusiform Cortex, anterior division | 3.09 | -38 | -8 | -44 |
| 24 | | | | | |
| | L Frontal Medial Cortex | 2.53 | -6 | 48 | -24 |
| 24 | | | | | |
| | L Subcallosal Cortex | 2.99 | -10 | 28 | -20 |
| 24 | | | | | |
| | R Planum Polare | 3.09 | 44 | 2 | -12 |
| 24 | | | | | |
| | R Frontal Pole | 3.02 | 30 | 62 | -6 |
| 24 | | | | | |
| | R Cingulate Gyrus, anterior division | 3.06 | 4 | 38 | -2 |
| 24 | | | | | |
| | R Inferior Frontal Gyrus, pars opercularis | 2.80 | 60 | 20 | 14 |
| 24 | | | | | |
| | L Lateral Occipital Cortex, superior division | 3.09 | -56 | -70 | 16 |
| 24 | | | | | |
| | R Frontal Pole | 3.03 | 4 | 64 | 30 |
| 24 | | | | | |
| | R Middle Frontal Gyrus | 3.65 | 52 | 26 | 38 |

| | | | | | |
|---|---|---|---|---|---|
| 24 | | | | | |
| | L Postcentral Gyrus | 2.97 | -40 | -28 | 40 |
| 24 | | | | | |
| | L Precuneus Cortex | 2.85 | -10 | -54 | 46 |
| 24 | | | | | |
| | L Middle Frontal Gyrus | 3.05 | -38 | 10 | 54 |
| 16 | | | | | |
| | L Temporal Fusiform Cortex, posterior division | 2.97 | -32 | -12 | -40 |
| 16 | | | | | |
| | L Inferior Temporal Gyrus, posterior division | 2.96 | -46 | -18 | -34 |
| 16 | | | | | |
| | L Inferior Temporal Gyrus, posterior division | 2.53 | -44 | -32 | -24 |
| 16 | | | | | |
| | L Inferior Temporal Gyrus, posterior division | 3.06 | -48 | -30 | -24 |
| 16 | | | | | |
| | L Brain-Stem | 3.33 | -14 | -22 | -22 |
| 16 | | | | | |
| | R Subcallosal Cortex | 2.92 | 8 | 12 | -18 |
| 16 | | | | | |
| | L Inferior Temporal Gyrus, temporooccipital part | 2.74 | -60 | -60 | -16 |
| 16 | | | | | |
| | R Middle Temporal Gyrus, posterior division | 2.65 | 48 | -16 | -14 |
| 16 | | | | | |
| | R Frontal Pole | 2.63 | 20 | 58 | -6 |
| 16 | | | | | |
| | R Middle Temporal Gyrus, temporooccipital part | 2.77 | 70 | -42 | -4 |
| 16 | | | | | |
| | L Frontal Pole | 2.67 | -42 | 58 | -4 |
| 16 | | | | | |
| | L Frontal Pole | 2.73 | -38 | 54 | -2 |
| 16 | | | | | |
| | L Planum Polare | 2.80 | -50 | -8 | 0 |
| 16 | | | | | |
| | R Thalamus | 2.81 | 16 | -30 | 6 |
| 16 | | | | | |
| | L Insular Cortex | 2.73 | -42 | -10 | 8 |
| 16 | | | | | |
| | L Frontal Pole | 2.79 | -8 | 62 | 8 |

| | | | | | |
|---|---|---|---|---|---|
| 16 | | | | | |
| | L Central Opercular Cortex | 3.21 | -44 | -12 | 16 |
| 16 | | | | | |
| | R Frontal Pole | 2.69 | 22 | 58 | 18 |
| 16 | | | | | |
| | R Frontal Pole | 2.61 | 4 | 60 | 20 |
| 16 | | | | | |
| | L Inferior Frontal Gyrus, pars opercularis | 2.93 | -36 | 14 | 26 |
| 16 | | | | | |
| | L Middle Frontal Gyrus | 3.47 | -34 | 2 | 50 |
| 16 | | | | | |
| | L Middle Frontal Gyrus | 2.73 | -30 | 16 | 58 |
| 16 | | | | | |
| | L Superior Parietal Lobule | 2.62 | -18 | -56 | 70 |
| 8 | | | | | |
| | R Brain-Stem | 2.93 | 4 | -46 | -66 |
| 8 | | | | | |
| | L Inferior Temporal Gyrus, anterior division | 2.53 | -52 | -6 | -44 |
| 8 | | | | | |
| | L Temporal Pole | 2.59 | -28 | 8 | -34 |
| 8 | | | | | |
| | L Temporal Fusiform Cortex, anterior division | 2.67 | -36 | -8 | -32 |
| 8 | | | | | |
| | L Temporal Pole | 2.67 | -24 | 10 | -32 |
| 8 | | | | | |
| | L Frontal Pole | 2.50 | -12 | 40 | -28 |
| 8 | | | | | |
| | R Inferior Temporal Gyrus, posterior division | 2.56 | 52 | -34 | -26 |
| 8 | | | | | |
| | R Middle Temporal Gyrus, posterior division | 2.48 | 62 | -16 | -26 |
| 8 | | | | | |
| | R Hippocampus | 2.67 | 24 | -10 | -24 |
| 8 | | | | | |
| | R Frontal Medial Cortex | 2.48 | 0 | 50 | -24 |
| 8 | | | | | |
| | R Hippocampus | 2.63 | 32 | -14 | -22 |
| 8 | | | | | |
| | L Hippocampus | 2.51 | -16 | -8 | -22 |
| 8 | | | | | |
| | L Frontal Orbital Cortex | 2.55 | -28 | 34 | -22 |
| 8 | | | | | |

| | | | | | |
|---|---|---|---|---|---|
| | R Planum Polare | 2.74 | 42 | 2 | -20 |
| 8 | | | | | |
| | L Frontal Orbital Cortex | 2.57 | -24 | 8 | -20 |
| 8 | | | | | |
| | L Subcallosal Cortex | 2.66 | -4 | 24 | -18 |
| 8 | | | | | |
| | L Frontal Pole | 2.51 | -26 | 54 | -18 |
| 8 | | | | | |
| | R Frontal Pole | 2.50 | 14 | 56 | -18 |
| 8 | | | | | |
| | L Amygdala | 2.82 | -14 | -4 | -16 |
| 8 | | | | | |
| | R Subcallosal Cortex | 2.69 | 12 | 14 | -16 |
| 8 | | | | | |
| | R Subcallosal Cortex | 3.01 | 4 | 26 | -16 |
| 8 | | | | | |
| | R Frontal Pole | 2.48 | 18 | 50 | -16 |
| 8 | | | | | |
| | R Inferior Temporal Gyrus, temporooccipital part | 2.52 | 62 | -54 | -14 |
| 8 | | | | | |
| | L Frontal Pole | 3.13 | -12 | 60 | -14 |
| 8 | | | | | |
| | L Insular Cortex | 2.86 | -40 | 2 | -12 |
| 8 | | | | | |
| | R Cingulate Gyrus, anterior division | 2.56 | 6 | 38 | -6 |
| 8 | | | | | |
| | L Middle Temporal Gyrus, temporooccipital part | 2.54 | -66 | -50 | -4 |
| 8 | | | | | |
| | L Pallidum | 2.84 | -24 | -16 | -4 |
| 8 | | | | | |
| | R Subcallosal Cortex | 2.84 | 2 | 24 | -4 |
| 8 | | | | | |
| | L Subcallosal Cortex | 2.55 | -2 | 16 | -2 |
| 8 | | | | | |
| | L Frontal Pole | 2.58 | -26 | 54 | -2 |
| 8 | | | | | |
| | L Frontal Pole | 2.50 | -6 | 60 | -2 |
| 8 | | | | | |
| | R Superior Temporal Gyrus, posterior division | 2.60 | 66 | -18 | 0 |
| 8 | | | | | |
| | R Insular Cortex | 2.54 | 44 | 0 | 0 |
| 8 | | | | | |

| | | | | | |
|---|---|---|---|---|---|
| | R Frontal Pole | 2.75 | 26 | 62 | 4 |
| 8 | | | | | |
| | R Paracingulate Gyrus | 2.81 | 6 | 54 | 8 |
| 8 | | | | | |
| | R Cingulate Gyrus, anterior division | 2.51 | 12 | 42 | 12 |
| 8 | | | | | |
| | R Central Opercular Cortex | 2.61 | 48 | -8 | 16 |
| 8 | | | | | |
| | R Angular Gyrus | 2.62 | 48 | -52 | 20 |
| 8 | | | | | |
| | L Central Opercular Cortex | 2.86 | -40 | -18 | 20 |
| 8 | | | | | |
| | L Inferior Frontal Gyrus, pars opercularis | 2.59 | -44 | 14 | 22 |
| 8 | | | | | |
| | L Lateral Occipital Cortex, superior division | 2.62 | -50 | -62 | 26 |
| 8 | | | | | |
| | R Middle Frontal Gyrus | 2.51 | 50 | 22 | 32 |
| 8 | | | | | |
| | R Frontal Pole | 2.64 | 10 | 56 | 40 |
| 8 | | | | | |
| | R Precuneus Cortex | 2.55 | 0 | -56 | 42 |
| 8 | | | | | |
| | R Middle Frontal Gyrus | 2.49 | 40 | 10 | 44 |
| 8 | | | | | |
| | R Superior Parietal Lobule | 2.58 | 28 | -52 | 52 |
| 8 | | | | | |
| | L Superior Frontal Gyrus | 2.88 | -22 | -2 | 52 |
| 8 | | | | | |
| | L Superior Frontal Gyrus | 2.65 | -16 | 18 | 58 |
| 8 | | | | | |
| | R Superior Frontal Gyrus | 2.74 | 12 | 20 | 60 |
| 8 | | | | | |
| | L Precentral Gyrus | 2.72 | -28 | -20 | 66 |
| 8 | | | | | |
| | L Postcentral Gyrus | 2.58 | -12 | -34 | 78 |

**Supplementary Table S29. [$UT_{CB2}$ > $UT_{CB3}$].** Descriptive statistics for clusters and local maxima showing greater activation for Convolved Block 2 relative to Convolved Block 3 of Uncertain-Threat anticipation (FDR $q$<0.05, whole-brain corrected).

| $mm^3$ | Label | *t* | *x* | *y* | *z* |
|---|---|---|---|---|---|
| 50,152 | | | | | |
| | R Temporal Fusiform Cortex, posterior division | 4.42 | 32 | -34 | -20 |
| | R Parahippocampal Gyrus, posterior division | 3.46 | 14 | -34 | -12 |
| | L Temporal Fusiform Cortex, posterior division | 4.67 | -28 | -36 | -22 |
| | R Temporal Occipital Fusiform Cortex | 6.47 | 32 | -50 | -20 |
| | L Temporal Occipital Fusiform Cortex | 5.00 | -32 | -52 | -18 |
| | L Inferior Temporal Gyrus, temporooccipital part | 3.55 | -52 | -52 | -26 |
| | R Lingual Gyrus | 5.27 | 16 | -70 | -12 |
| | L Occipital Fusiform Gyrus | 5.56 | -18 | -74 | -14 |
| | R Occipital Fusiform Gyrus | 6.36 | 22 | -80 | -18 |
| | L Occipital Pole | 5.53 | -12 | -92 | -12 |
| | R Occipital Pole | 2.95 | 6 | -92 | -4 |
| 30,432 | | | | | |
| | L Precentral Gyrus | 4.50 | -30 | -26 | 54 |
| | L Postcentral Gyrus | 5.28 | -38 | -28 | 50 |
| | L Supramarginal Gyrus, anterior division | 4.50 | -46 | -40 | 44 |
| | L Superior Parietal Lobule | 4.33 | -38 | -40 | 44 |
| | L Supramarginal Gyrus, posterior division | 4.79 | -42 | -44 | 44 |
| | R Cingulate Gyrus, posterior division | 3.62 | 0 | -48 | 14 |
| | L Precuneus Cortex | 4.53 | -4 | -60 | 38 |
| | L Lingual Gyrus | 4.42 | -10 | -62 | 0 |
| | L Lateral Occipital Cortex, superior division | 5.95 | -24 | -64 | 48 |
| | R Intracalcarine Cortex | 6.81 | 8 | -66 | 16 |
| | R Precuneus Cortex | 5.27 | 10 | -66 | 22 |
| | R Lingual Gyrus | 3.25 | 10 | -66 | -4 |
| | R Cuneal Cortex | 5.06 | 6 | -70 | 22 |
| | R Supracalcarine Cortex | 6.02 | 0 | -72 | 16 |
| | L Intracalcarine Cortex | 6.06 | -12 | -74 | 12 |
| | L Cuneal Cortex | 3.26 | -2 | -80 | 34 |
| 2,288 | | | | | |
| | L Frontal Pole | 4.51 | -10 | 72 | -6 |
| | R Frontal Pole | 5.55 | 8 | 72 | 2 |
| 1,704 | | | | | |
| | R Lateral Occipital Cortex, superior division | 5.76 | 30 | -64 | 40 |

| | | | | | |
|---|---|---|---|---|---|
| 1,680 | | | | | |
| | R Frontal Pole | 3.93 | 18 | 38 | 50 |
| | R Superior Frontal Gyrus | 5.12 | 24 | 32 | 50 |
| | R Middle Frontal Gyrus | 4.29 | 32 | 16 | 58 |
| 1,216 | | | | | |
| | R Middle Frontal Gyrus | 4.36 | 46 | 34 | 32 |
| 1,008 | | | | | |
| | L Middle Frontal Gyrus | 4.55 | -44 | 16 | 34 |
| | L Precentral Gyrus | 3.42 | -42 | 4 | 32 |
| 760 | | | | | |
| | L Middle Temporal Gyrus, posterior division | 3.81 | -64 | -40 | 2 |
| 712 | | | | | |
| | L Middle Frontal Gyrus | 4.22 | -46 | 32 | 20 |
| | L Inferior Frontal Gyrus, pars triangularis | 3.95 | -48 | 26 | 20 |
| 648 | | | | | |
| | L Lateral Occipital Cortex, superior division | 4.03 | -38 | -86 | 20 |
| 608 | | | | | |
| | R Postcentral Gyrus | 4.28 | 58 | -8 | 26 |
| 552 | | | | | |
| | R Lateral Occipital Cortex, superior division | 3.62 | 36 | -80 | 20 |
| 544 | | | | | |
| | R Brain-Stem | 4.03 | 0 | -42 | -58 |
| | L Brain-Stem | 4.18 | -2 | -48 | -62 |
| 488 | | | | | |
| | R Angular Gyrus | 4.02 | 46 | -56 | 32 |
| 488 | | | | | |
| | R Precentral Gyrus | 3.15 | 0 | -28 | 70 |
| | R Postcentral Gyrus | 4.26 | 4 | -34 | 66 |
| 448 | | | | | |
| | R Accumbens | 4.84 | 8 | 6 | -6 |
| 440 | | | | | |
| | L Postcentral Gyrus | 4.30 | -62 | -8 | 30 |
| 384 | | | | | |
| | R Amygdala | 3.35 | 14 | -8 | -16 |
| | R Hippocampus | 5.15 | 18 | -16 | -16 |
| 376 | | | | | |
| | L Postcentral Gyrus | 4.31 | -6 | -36 | 74 |
| 296 | | | | | |
| | L Brain-Stem | 3.88 | -6 | -40 | -46 |
| | R Brain-Stem | 3.81 | 0 | -42 | -42 |
| 288 | | | | | |

| | | | | | |
|---|---|---|---|---|---|
| | R Hippocampus | 4.34 | 28 | -32 | -6 |
| | R Thalamus | 3.67 | 18 | -34 | -2 |
| 264 | | | | | |
| | L Thalamus | 4.59 | -22 | -34 | -2 |
| 256 | | | | | |
| | L Thalamus | 4.18 | -16 | -24 | 4 |
| 248 | | | | | |
| | L Insular Cortex | 4.23 | -36 | -10 | 14 |
| 232 | | | | | |
| | L Lateral Occipital Cortex, inferior division | 3.66 | -48 | -62 | -8 |
| 192 | | | | | |
| | L Amygdala | 4.82 | -26 | -8 | -12 |
| 176 | | | | | |
| | R Amygdala | 3.26 | 26 | -6 | -12 |
| | R Putamen | 3.61 | 32 | -10 | -10 |
| | R Amygdala | 3.44 | 26 | -10 | -10 |
| 168 | | | | | |
| | L Hippocampus | 4.32 | -16 | -16 | -18 |
| 160 | | | | | |
| | L Postcentral Gyrus | 3.53 | -48 | -24 | 38 |
| 152 | | | | | |
| | L Temporal Pole | 3.57 | -58 | 6 | -12 |
| 136 | | | | | |
| | L Parahippocampal Gyrus, posterior division | 3.64 | -24 | -42 | -12 |
| | L Temporal Occipital Fusiform Cortex | 3.19 | -24 | -48 | -10 |
| 128 | | | | | |
| | L Temporal Fusiform Cortex, posterior division | 3.77 | -38 | -32 | -28 |
| 128 | | | | | |
| | L Frontal Pole | 3.85 | -4 | 68 | 4 |
| 112 | | | | | |
| | L Superior Temporal Gyrus, anterior division | 3.43 | -56 | -8 | -8 |
| 112 | | | | | |
| | R Superior Frontal Gyrus | 3.19 | 22 | 18 | 52 |
| | R Middle Frontal Gyrus | 3.04 | 28 | 18 | 46 |
| 112 | | | | | |
| | R Lateral Occipital Cortex, superior division | 3.57 | 32 | -64 | 52 |
| 104 | | | | | |
| | L Brain-Stem | 4.42 | -18 | -34 | -42 |
| 104 | | | | | |
| | L Temporal Pole | 3.31 | -36 | 6 | -22 |
| 104 | | | | | |

| | | | | | |
|---|---|---|---|---|---|
| | R Angular Gyrus | 3.37 | 52 | -58 | 26 |
| 96 | | | | | |
| | L Middle Temporal Gyrus, posterior division | 2.88 | -66 | -12 | -10 |
| 88 | | | | | |
| | R Frontal Pole | 3.21 | 2 | 56 | -22 |
| 88 | | | | | |
| | L Brain-Stem | 4.12 | -14 | -26 | -16 |
| 88 | | | | | |
| | L Postcentral Gyrus | 3.94 | -6 | -42 | 68 |
| 80 | | | | | |
| | R Brain-Stem | 3.49 | 6 | -34 | -50 |
| 80 | | | | | |
| | L Inferior Temporal Gyrus, anterior division | 3.30 | -42 | -6 | -40 |
| 80 | | | | | |
| | L Temporal Fusiform Cortex, posterior division | 3.82 | -32 | -22 | -32 |
| 72 | | | | | |
| | L Lateral Occipital Cortex, superior division | 3.31 | -38 | -78 | 20 |
| 72 | | | | | |
| | R Frontal Pole | 3.35 | 20 | 48 | 36 |
| 72 | | | | | |
| | R Supramarginal Gyrus, posterior division | 3.13 | 50 | -40 | 50 |
| 64 | | | | | |
| | L Frontal Pole | 3.63 | -4 | 64 | -22 |
| 64 | | | | | |
| | R Cingulate Gyrus, posterior division | 3.69 | 6 | -46 | 4 |
| 64 | | | | | |
| | L Frontal Pole | 3.24 | -16 | 66 | 16 |
| 64 | | | | | |
| | L Occipital Pole | 3.04 | -12 | -94 | 30 |
| 56 | | | | | |
| | L Thalamus | 3.33 | -12 | -36 | 6 |
| 56 | | | | | |
| | R Central Opercular Cortex | 4.41 | 38 | -6 | 18 |
| 56 | | | | | |
| | R Middle Frontal Gyrus | 2.89 | 30 | 8 | 50 |
| 48 | | | | | |
| | R Temporal Pole | 2.79 | 32 | 14 | -30 |
| 48 | | | | | |
| | L Frontal Pole | 3.33 | -24 | 58 | 0 |
| 48 | | | | | |
| | R Occipital Pole | 3.42 | 4 | -98 | 6 |

| | | | | | |
|---|---|---|---|---|---|
| 48 | | | | | |
| | L Lateral Occipital Cortex, superior division | 3.57 | -46 | -68 | 26 |
| 48 | | | | | |
| | L Superior Frontal Gyrus | 3.44 | -6 | 46 | 42 |
| 40 | | | | | |
| | R Brain-Stem | 4.27 | 6 | -36 | -64 |
| 40 | | | | | |
| | R Brain-Stem | 3.12 | 10 | -36 | -44 |
| 40 | | | | | |
| | L Brain-Stem | 3.44 | -18 | -26 | -38 |
| 40 | | | | | |
| | L Middle Temporal Gyrus, anterior division | 3.59 | -58 | -2 | -18 |
| 40 | | | | | |
| | R Superior Temporal Gyrus, anterior division | 2.98 | 54 | -2 | -18 |
| 40 | | | | | |
| | L Accumbens | 2.94 | -10 | 8 | -10 |
| | L Putamen | 3.02 | -14 | 6 | -8 |
| 40 | | | | | |
| | R Lingual Gyrus | 3.56 | 12 | -56 | -4 |
| 40 | | | | | |
| | R Insular Cortex | 2.98 | 38 | -4 | 10 |
| 40 | | | | | |
| | R Occipital Pole | 3.55 | 2 | -94 | 18 |
| 40 | | | | | |
| | L Cingulate Gyrus, posterior division | 3.28 | -4 | -40 | 30 |
| 40 | | | | | |
| | L Cingulate Gyrus, anterior division | 3.32 | -4 | 2 | 30 |
| 40 | | | | | |
| | R Lateral Occipital Cortex, superior division | 3.42 | 48 | -72 | 42 |
| 32 | | | | | |
| | L Inferior Temporal Gyrus, anterior division | 3.11 | -50 | -10 | -36 |
| 32 | | | | | |
| | R Brain-Stem | 3.12 | 20 | -34 | -32 |
| 32 | | | | | |
| | L Parahippocampal Gyrus, anterior division | 3.02 | -18 | 0 | -32 |
| 32 | | | | | |
| | L Amygdala | 2.94 | -14 | -2 | -20 |
| 32 | | | | | |

| | | | | | |
|---|---|---|---|---|---|
| | L Superior Temporal Gyrus, anterior division | 3.06 | -50 | -6 | -16 |
| 32 | | | | | |
| | L Amygdala | 3.40 | -26 | 0 | -18 |
| 32 | | | | | |
| | L Frontal Pole | 2.91 | -4 | 62 | -12 |
| 32 | | | | | |
| | L Lingual Gyrus | 2.83 | -28 | -44 | -6 |
| 32 | | | | | |
| | R Lingual Gyrus | 3.89 | 10 | -72 | -2 |
| 32 | | | | | |
| | L Angular Gyrus | 3.13 | -42 | -58 | 30 |
| 32 | | | | | |
| | R Lateral Occipital Cortex, superior division | 2.99 | 54 | -68 | 34 |
| 32 | | | | | |
| | R Cingulate Gyrus, posterior division | 2.98 | 10 | -42 | 38 |
| 32 | | | | | |
| | L Postcentral Gyrus | 2.93 | -58 | -18 | 50 |
| 32 | | | | | |
| | L Superior Frontal Gyrus | 3.63 | -22 | -6 | 56 |
| 24 | | | | | |
| | L Brain-Stem | 2.87 | -6 | -38 | -64 |
| 24 | | | | | |
| | L Temporal Pole | 3.67 | -22 | 6 | -40 |
| 24 | | | | | |
| | L Parahippocampal Gyrus, anterior division | 3.70 | -18 | -8 | -36 |
| 24 | | | | | |
| | L Inferior Temporal Gyrus, posterior division | 3.01 | -50 | -36 | -24 |
| 24 | | | | | |
| | L Middle Temporal Gyrus, anterior division | 2.86 | -56 | -2 | -24 |
| 24 | | | | | |
| | L Brain-Stem | 2.95 | -4 | -36 | -20 |
| 24 | | | | | |
| | L Frontal Pole | 3.11 | -22 | 40 | -22 |
| 24 | | | | | |
| | R Frontal Pole | 2.79 | 12 | 48 | -22 |
| 24 | | | | | |
| | R Middle Temporal Gyrus, posterior division | 3.24 | 54 | -10 | -18 |
| 24 | | | | | |
| | L Occipital Pole | 3.51 | -6 | -100 | 6 |

| | | | | | |
|---|---|---|---|---|---|
| 24 | | | | | |
| | L Frontal Pole | 3.11 | -20 | 70 | 8 |
| 24 | | | | | |
| | R Frontal Pole | 2.99 | 38 | 38 | 16 |
| 24 | | | | | |
| | L Occipital Pole | 3.11 | -6 | -90 | 18 |
| 24 | | | | | |
| | L Cuneal Cortex | 3.20 | -8 | -84 | 34 |
| 24 | | | | | |
| | L Precuneus Cortex | 3.61 | -6 | -74 | 42 |
| 24 | | | | | |
| | R Superior Frontal Gyrus | 3.31 | 20 | 24 | 42 |
| 24 | | | | | |
| | L Frontal Pole | 3.17 | -14 | 42 | 44 |
| 16 | | | | | |
| | L Temporal Fusiform Cortex, anterior division | 3.28 | -38 | -2 | -48 |
| 16 | | | | | |
| | R Temporal Fusiform Cortex, anterior division | 2.78 | 34 | -8 | -46 |
| 16 | | | | | |
| | L Temporal Fusiform Cortex, anterior division | 2.72 | -28 | -8 | -42 |
| 16 | | | | | |
| | L Brain-Stem | 2.91 | -6 | -24 | -40 |
| 16 | | | | | |
| | L Parahippocampal Gyrus, anterior division | 2.81 | -24 | 0 | -40 |
| 16 | | | | | |
| | L Temporal Pole | 3.09 | -42 | 2 | -40 |
| 16 | | | | | |
| | L Temporal Pole | 2.94 | -38 | 14 | -40 |
| 16 | | | | | |
| | L Temporal Pole | 3.28 | -36 | 10 | -36 |
| 16 | | | | | |
| | L Temporal Fusiform Cortex, posterior division | 2.87 | -38 | -14 | -26 |
| 16 | | | | | |
| | R Subcallosal Cortex | 2.81 | 8 | 22 | -24 |
| 16 | | | | | |
| | L Temporal Fusiform Cortex, posterior division | 2.99 | -40 | -18 | -22 |
| 16 | | | | | |
| | L Inferior Temporal Gyrus, temporooccipital part | 3.00 | -48 | -56 | -20 |

| | | | | | |
|---|---|---|---|---|---|
| 16 | | | | | |
| | L Amygdala | 3.94 | -32 | -4 | -20 |
| 16 | | | | | |
| | R Temporal Pole | 3.43 | 30 | 6 | -20 |
| 16 | | | | | |
| | L Frontal Medial Cortex | 3.79 | -4 | 52 | -20 |
| 16 | | | | | |
| | L Subcallosal Cortex | 3.51 | -4 | 18 | -16 |
| 16 | | | | | |
| | L Occipital Fusiform Gyrus | 2.97 | -28 | -80 | -14 |
| 16 | | | | | |
| | L Inferior Temporal Gyrus, temporooccipital part | 3.01 | -46 | -58 | -14 |
| 16 | | | | | |
| | R Hippocampus | 2.94 | 30 | -18 | -12 |
| 16 | | | | | |
| | L Hippocampus | 3.54 | -28 | -18 | -12 |
| 16 | | | | | |
| | R Occipital Pole | 2.89 | 4 | -98 | -2 |
| 16 | | | | | |
| | R Occipital Fusiform Gyrus | 3.32 | 28 | -72 | -4 |
| 16 | | | | | |
| | L Frontal Pole | 2.91 | -34 | 60 | 2 |
| 16 | | | | | |
| | R Frontal Pole | 3.12 | 42 | 48 | 2 |
| 16 | | | | | |
| | L Central Opercular Cortex | 3.32 | -44 | -10 | 16 |
| 16 | | | | | |
| | L Cingulate Gyrus, posterior division | 3.06 | -6 | -46 | 20 |
| 16 | | | | | |
| | L Angular Gyrus | 3.40 | -50 | -52 | 24 |
| 16 | | | | | |
| | L Precentral Gyrus | 2.93 | -52 | -4 | 24 |
| 16 | | | | | |
| | L Middle Frontal Gyrus | 3.14 | -38 | 28 | 24 |
| 16 | | | | | |
| | R Cuneal Cortex | 2.86 | 16 | -72 | 28 |
| 16 | | | | | |
| | L Postcentral Gyrus | 3.05 | -58 | -18 | 36 |
| 16 | | | | | |
| | R Frontal Pole | 2.84 | 16 | 54 | 38 |
| 16 | | | | | |
| | R Superior Parietal Lobule | 2.84 | 38 | -48 | 46 |
| 16 | | | | | |

| | | | | | |
|---|---|---|---|---|---|
| | R Lateral Occipital Cortex, superior division | 2.74 | 38 | -64 | 54 |
| 16 | | | | | |
| | R Lateral Occipital Cortex, superior division | 2.80 | 18 | -68 | 58 |
| 16 | | | | | |
| | L Middle Frontal Gyrus | 2.99 | -30 | 20 | 56 |
| 8 | | | | | |
| | R Brain-Stem | 2.83 | 10 | -38 | -58 |
| 8 | | | | | |
| | L Brain-Stem | 2.75 | -2 | -30 | -54 |
| 8 | | | | | |
| | R Temporal Pole | 2.91 | 30 | 4 | -48 |
| 8 | | | | | |
| | R Temporal Pole | 2.87 | 44 | 14 | -36 |
| 8 | | | | | |
| | R Temporal Pole | 2.78 | 38 | 18 | -36 |
| 8 | | | | | |
| | R Brain-Stem | 2.71 | 8 | -40 | -28 |
| 8 | | | | | |
| | R Frontal Medial Cortex | 2.77 | 2 | 32 | -24 |
| 8 | | | | | |
| | L Brain-Stem | 2.93 | -4 | -14 | -22 |
| 8 | | | | | |
| | L Temporal Pole | 3.05 | -44 | 12 | -22 |
| 8 | | | | | |
| | L Frontal Pole | 2.74 | -10 | 64 | -20 |
| 8 | | | | | |
| | L Hippocampus | 2.92 | -12 | -12 | -18 |
| 8 | | | | | |
| | L Frontal Pole | 2.70 | -14 | 66 | -18 |
| 8 | | | | | |
| | L Parahippocampal Gyrus, posterior division | 2.95 | -20 | -30 | -16 |
| 8 | | | | | |
| | L Frontal Pole | 2.71 | -6 | 56 | -16 |
| 8 | | | | | |
| | L Lateral Occipital Cortex, inferior division | 2.76 | -56 | -64 | -14 |
| 8 | | | | | |
| | L Brain-Stem | 2.78 | -2 | -28 | -14 |
| 8 | | | | | |
| | L Frontal Pole | 2.73 | -20 | 50 | -14 |
| 8 | | | | | |
| | R Frontal Pole | 3.20 | 22 | 66 | -14 |

| | | | | | |
|---|---|---|---|---|---|
| 8 | | | | | |
| | L Frontal Pole | 2.85 | -18 | 68 | -12 |
| 8 | | | | | |
| | R Brain-Stem | 2.68 | 10 | -28 | -8 |
| 8 | | | | | |
| | R Frontal Pole | 3.27 | 32 | 36 | -8 |
| 8 | | | | | |
| | L Frontal Pole | 2.68 | -10 | 64 | -8 |
| 8 | | | | | |
| | L Lingual Gyrus | 3.01 | -22 | -60 | -6 |
| 8 | | | | | |
| | L Inferior Temporal Gyrus, temporooccipital part | 2.74 | -42 | -54 | -6 |
| 8 | | | | | |
| | R Subcallosal Cortex | 3.49 | 4 | 22 | -6 |
| 8 | | | | | |
| | L Frontal Pole | 2.69 | -40 | 46 | -6 |
| 8 | | | | | |
| | L Middle Temporal Gyrus, temporooccipital part | 2.71 | -68 | -48 | -4 |
| 8 | | | | | |
| | R Pallidum | 3.45 | 20 | -8 | -4 |
| 8 | | | | | |
| | L Frontal Pole | 2.95 | -20 | 66 | 4 |
| 8 | | | | | |
| | R Postcentral Gyrus | 2.77 | 66 | -8 | 14 |
| 8 | | | | | |
| | L Frontal Pole | 2.69 | -8 | 60 | 18 |
| 8 | | | | | |
| | R Lateral Occipital Cortex, superior division | 2.72 | 46 | -62 | 24 |
| 8 | | | | | |
| | L Angular Gyrus | 2.72 | -40 | -56 | 24 |
| 8 | | | | | |
| | L Cingulate Gyrus, posterior division | 2.71 | -14 | -46 | 32 |
| 8 | | | | | |
| | R Cingulate Gyrus, anterior division | 2.81 | 6 | -4 | 32 |
| 8 | | | | | |
| | R Cingulate Gyrus, anterior division | 2.71 | 4 | 0 | 32 |
| 8 | | | | | |
| | L Lateral Occipital Cortex, superior division | 2.70 | -44 | -62 | 36 |
| 8 | | | | | |
| | L Middle Frontal Gyrus | 2.80 | -50 | 26 | 36 |
| 8 | | | | | |

| | | | | | |
|---|---|---|---|---|---|
| | R Frontal Pole | 2.69 | 14 | 52 | 36 |
| 8 | | | | | |
| | L Frontal Pole | 2.73 | -18 | 40 | 38 |
| 8 | | | | | |
| | L Cuneal Cortex | 2.80 | -6 | -86 | 40 |
| 8 | | | | | |
| | L Superior Frontal Gyrus | 2.84 | -20 | 26 | 40 |
| 8 | | | | | |
| | R Lateral Occipital Cortex, superior division | 2.80 | 38 | -78 | 42 |
| 8 | | | | | |
| | L Lateral Occipital Cortex, superior division | 2.69 | -34 | -74 | 42 |
| 8 | | | | | |
| | L Frontal Pole | 2.83 | -24 | 40 | 42 |
| 8 | | | | | |
| | L Precuneus Cortex | 2.70 | -8 | -70 | 44 |
| 8 | | | | | |
| | R Lateral Occipital Cortex, superior division | 2.81 | 38 | -70 | 46 |
| 8 | | | | | |
| | L Precuneus Cortex | 2.91 | -6 | -56 | 46 |
| 8 | | | | | |
| | L Paracingulate Gyrus | 2.99 | -8 | 16 | 46 |
| 8 | | | | | |
| | L Lateral Occipital Cortex, superior division | 2.78 | -18 | -82 | 48 |
| 8 | | | | | |
| | R Lateral Occipital Cortex, superior division | 2.86 | 38 | -74 | 50 |
| 8 | | | | | |
| | L Superior Parietal Lobule | 2.71 | -28 | -50 | 54 |
| 8 | | | | | |
| | R Superior Frontal Gyrus | 3.37 | 16 | 28 | 54 |
| 8 | | | | | |
| | L Lateral Occipital Cortex, superior division | 2.86 | -14 | -68 | 62 |
| 8 | | | | | |
| | R Postcentral Gyrus | 2.69 | 14 | -36 | 72 |
| 8 | | | | | |
| | R Postcentral Gyrus | 2.74 | 8 | -38 | 74 |
| 8 | | | | | |
| | R Postcentral Gyrus | 2.91 | 10 | -34 | 74 |
| 8 | | | | | |
| | R Precentral Gyrus | 2.78 | 6 | -28 | 78 |

**Supplementary Table S30. [$UT_{CB3}$ - $UT_{CB2}$] > [$CT_{CB3}$ - $CT_{CB2}$].** Descriptive statistics for clusters and local maxima showing greater increases in activation from Convolved Block 2 to Convolved Block 3 of Uncertain-Threat anticipation relative to increases in activation from Convolved Block 2 to Convolved Block 3 of Certain-Threat anticipation (FDR *q*<0.05, whole-brain corrected).

| mm³ | Label | t | x | y | z |
|---|---|---|---|---|---|
| 57,120 | | | | | |
| | R Hippocampus | 4.13 | 30 | -22 | -16 |
| | R Parahippocampal Gyrus, posterior division | 5.53 | 26 | -28 | -22 |
| | L Parahippocampal Gyrus, posterior division | 4.89 | -26 | -32 | -20 |
| | R Temporal Fusiform Cortex, posterior division | 4.76 | 26 | -34 | -20 |
| | R Thalamus | 4.07 | 8 | -34 | 6 |
| | L Temporal Fusiform Cortex, posterior division | 3.11 | -32 | -36 | -22 |
| | L Cingulate Gyrus, posterior division | 4.25 | -6 | -46 | 4 |
| | R Cingulate Gyrus, posterior division | 4.45 | 8 | -46 | 4 |
| | L Temporal Occipital Fusiform Cortex | 5.79 | -30 | -48 | -8 |
| | L Inferior Temporal Gyrus, temporooccipital part | 4.42 | -42 | -48 | -12 |
| | R Lingual Gyrus | 7.75 | 22 | -50 | 0 |
| | R Temporal Occipital Fusiform Cortex | 4.24 | 24 | -52 | -12 |
| | L Precuneus Cortex | 4.77 | -2 | -54 | 10 |
| | L Lingual Gyrus | 7.06 | -22 | -56 | 2 |
| | R Inferior Temporal Gyrus, temporooccipital part | 3.79 | 46 | -56 | -14 |
| | R Precuneus Cortex | 5.22 | 20 | -58 | 14 |
| | R Occipital Fusiform Gyrus | 5.19 | 38 | -64 | -12 |
| | R Lateral Occipital Cortex, superior division | 5.89 | 32 | -72 | 32 |
| | L Occipital Fusiform Gyrus | 5.11 | -36 | -74 | -14 |
| | L Lateral Occipital Cortex, superior division | 5.56 | -26 | -76 | 30 |
| | R Lateral Occipital Cortex, inferior division | 5.79 | 48 | -78 | 8 |
| | R Intracalcarine Cortex | 5.04 | 14 | -78 | 8 |
| | R Cuneal Cortex | 6.57 | 6 | -82 | 40 |
| | L Cuneal Cortex | 5.79 | -4 | -86 | 36 |
| | L Intracalcarine Cortex | 5.07 | -8 | -86 | 0 |
| | R Supracalcarine Cortex | 5.43 | 2 | -86 | 12 |
| | L Lateral Occipital Cortex, inferior division | 7.31 | -38 | -88 | -10 |
| | R Occipital Pole | 6.01 | 8 | -88 | 34 |
| | L Occipital Pole | 6.68 | -6 | -94 | 30 |
| 5,936 | | | | | |
| | R Temporal Pole | 6.00 | 46 | 20 | -20 |
| | R Planum Polare | 5.43 | 50 | 4 | -4 |

| | | | | | |
|---|---|---|---|---|---|
| | R Superior Temporal Gyrus, anterior division | 4.65 | 58 | 4 | -8 |
| | R Middle Temporal Gyrus, anterior division | 3.61 | 60 | -2 | -16 |
| | R Insular Cortex | 3.06 | 44 | -4 | -8 |
| | R Planum Temporale | 4.19 | 66 | -10 | 6 |
| | R Central Opercular Cortex | 3.64 | 64 | -12 | 12 |
| | R Middle Temporal Gyrus, posterior division | 3.53 | 68 | -14 | -12 |
| | R Superior Temporal Gyrus, posterior division | 3.71 | 66 | -18 | 0 |
| 2,216 | | | | | |
| | L Frontal Orbital Cortex | 4.45 | -40 | 22 | -16 |
| | L Temporal Pole | 5.95 | -36 | 18 | -24 |
| 2,088 | | | | | |
| | R Inferior Frontal Gyrus, pars triangularis | 3.88 | 58 | 28 | 10 |
| | R Inferior Frontal Gyrus, pars opercularis | 5.52 | 52 | 12 | 22 |
| | R Middle Frontal Gyrus | 3.04 | 48 | 12 | 42 |
| | R Precentral Gyrus | 3.78 | 44 | 8 | 34 |
| 2,056 | | | | | |
| | L Superior Temporal Gyrus, anterior division | 3.16 | -60 | 0 | -6 |
| | L Planum Polare | 4.67 | -52 | -8 | 2 |
| | L Superior Temporal Gyrus, posterior division | 4.46 | -66 | -14 | 0 |
| | L Heschls Gyrus (includes H1 and H2) | 4.07 | -52 | -14 | 8 |
| | L Central Opercular Cortex | 4.45 | -62 | -20 | 12 |
| 1,800 | | | | | |
| | R Frontal Pole | 5.23 | 12 | 62 | 28 |
| | R Superior Frontal Gyrus | 3.64 | 6 | 54 | 20 |
| | L Paracingulate Gyrus | 2.99 | -4 | 52 | 8 |
| | R Paracingulate Gyrus | 3.86 | 10 | 48 | 10 |
| | R Cingulate Gyrus, anterior division | 3.89 | 4 | 38 | 8 |
| 1,592 | | | | | |
| | R Postcentral Gyrus | 5.13 | 66 | -14 | 34 |
| | R Supramarginal Gyrus, anterior division | 4.28 | 56 | -22 | 40 |
| 1,440 | | | | | |
| | R Frontal Pole | 4.27 | 48 | 38 | -12 |
| | R Frontal Orbital Cortex | 4.82 | 48 | 28 | -8 |
| 1,432 | | | | | |
| | R Frontal Pole | 4.12 | 50 | 36 | 18 |
| | R Middle Frontal Gyrus | 3.24 | 50 | 30 | 24 |
| 1,080 | | | | | |
| | L Inferior Frontal Gyrus, pars triangularis | 3.88 | -52 | 36 | 6 |
| | L Frontal Orbital Cortex | 4.64 | -48 | 34 | -8 |

| | | | | | |
|---|---|---|---|---|---|
| | L Frontal Operculum Cortex | 4.01 | -46 | 24 | 0 |
| 1,024 | | | | | |
| | L Precentral Gyrus | 2.88 | -42 | -16 | 66 |
| | L Postcentral Gyrus | 4.12 | -46 | -20 | 60 |
| 616 | | | | | |
| | L Frontal Pole | 4.60 | -8 | 66 | 24 |
| | L Superior Frontal Gyrus | 2.90 | -2 | 56 | 26 |
| 504 | | | | | |
| | R Superior Frontal Gyrus | 4.00 | 20 | 28 | 48 |
| | R Middle Frontal Gyrus | 3.88 | 26 | 26 | 48 |
| 432 | | | | | |
| | L Paracingulate Gyrus | 4.20 | -12 | 46 | -4 |
| | L Frontal Medial Cortex | 3.84 | -8 | 46 | -10 |
| 432 | | | | | |
| | L Postcentral Gyrus | 3.57 | -12 | -42 | 54 |
| | L Precuneus Cortex | 4.50 | -6 | -46 | 64 |
| 400 | | | | | |
| | R Frontal Pole | 4.05 | 10 | 46 | 50 |
| 368 | | | | | |
| | L Superior Temporal Gyrus, posterior division | 3.67 | -52 | -36 | 2 |
| 344 | | | | | |
| | L Parahippocampal Gyrus, anterior division | 3.26 | -18 | -22 | -20 |
| | L Hippocampus | 4.61 | -20 | -28 | -12 |
| | L Parahippocampal Gyrus, posterior division | 3.49 | -16 | -30 | -14 |
| 256 | | | | | |
| | L Temporal Fusiform Cortex, posterior division | 3.90 | -34 | -44 | -20 |
| | L Temporal Occipital Fusiform Cortex | 3.39 | -42 | -48 | -18 |
| 256 | | | | | |
| | L Frontal Pole | 3.79 | -4 | 58 | 34 |
| 256 | | | | | |
| | R Postcentral Gyrus | 3.54 | 20 | -30 | 78 |
| 232 | | | | | |
| | L Frontal Pole | 3.33 | -46 | 44 | -12 |
| 216 | | | | | |
| | L Superior Frontal Gyrus | 4.62 | -4 | 16 | 62 |
| 192 | | | | | |
| | L Temporal Pole | 4.21 | -48 | 10 | -10 |
| 184 | | | | | |
| | R Middle Temporal Gyrus, temporooccipital part | 4.47 | 56 | -48 | -8 |
| 184 | | | | | |

| | | | | | |
|---|---|---|---|---|---|
| | L Postcentral Gyrus | 3.09 | -18 | -30 | 78 |
| 176 | | | | | |
| | R Temporal Occipital Fusiform Cortex | 3.54 | 38 | -44 | -20 |
| 160 | | | | | |
| | L Superior Parietal Lobule | 3.20 | -26 | -56 | 64 |
| 144 | | | | | |
| | R Middle Temporal Gyrus, temporooccipital part | 4.17 | 58 | -58 | 12 |
| 144 | | | | | |
| | R Lateral Occipital Cortex, superior division | 3.66 | 18 | -58 | 60 |
| 136 | | | | | |
| | R Frontal Pole | 3.78 | 10 | 68 | 18 |
| 128 | | | | | |
| | R Middle Temporal Gyrus, posterior division | 4.09 | 48 | -36 | -2 |
| 112 | | | | | |
| | L Paracingulate Gyrus | 3.38 | -6 | 48 | 14 |
| 104 | | | | | |
| | L Occipital Fusiform Gyrus | 4.14 | -28 | -74 | -14 |
| 104 | | | | | |
| | R Cuneal Cortex | 3.58 | 14 | -76 | 30 |
| 96 | | | | | |
| | R Parahippocampal Gyrus, anterior division | 3.55 | 18 | -8 | -28 |
| 96 | | | | | |
| | L Frontal Pole | 3.73 | -12 | 68 | 12 |
| 96 | | | | | |
| | R Precuneus Cortex | 3.35 | 4 | -62 | 20 |
| 88 | | | | | |
| | L Temporal Pole | 3.29 | -54 | 6 | -20 |
| 80 | | | | | |
| | R Temporal Pole | 3.27 | 44 | 12 | -28 |
| 80 | | | | | |
| | L Temporal Fusiform Cortex, posterior division | 3.74 | -42 | -38 | -18 |
| 80 | | | | | |
| | L Superior Temporal Gyrus, posterior division | 3.49 | -62 | -30 | 4 |
| 72 | | | | | |
| | R Frontal Pole | 3.36 | 32 | 40 | -12 |
| 71 | | | | | |
| | L Middle Frontal Gyrus | 3.46 | -40 | 12 | 46 |
| 56 | | | | | |
| | L Temporal Pole | 3.01 | -26 | 4 | -28 |

| | | | | | |
|---|---|---|---|---|---|
| | L Amygdala | 3.28 | -24 | 2 | -26 |
| | L Parahippocampal Gyrus, anterior division | 3.21 | -24 | 2 | -30 |
| 56 | | | | | |
| | L Superior Temporal Gyrus, anterior division | 3.02 | -52 | -2 | -16 |
| | L Middle Temporal Gyrus, anterior division | 3.10 | -54 | -4 | -18 |
| 56 | | | | | |
| | L Superior Temporal Gyrus, anterior division | 3.91 | -58 | 0 | -14 |
| 56 | | | | | |
| | R Middle Temporal Gyrus, temporooccipital part | 3.08 | 68 | -40 | -10 |
| 56 | | | | | |
| | L Lateral Occipital Cortex, inferior division | 3.91 | -56 | -66 | -6 |
| 56 | | | | | |
| | R Central Opercular Cortex | 3.24 | 40 | -6 | 12 |
| 48 | | | | | |
| | L Planum Polare | 2.91 | -48 | 0 | -6 |
| 48 | | | | | |
| | R Middle Frontal Gyrus | 2.86 | 42 | 12 | 44 |
| 48 | | | | | |
| | R Postcentral Gyrus | 3.44 | 8 | -44 | 68 |
| 48 | | | | | |
| | R Precentral Gyrus | 3.93 | 6 | -24 | 80 |
| 40 | | | | | |
| | L Temporal Pole | 3.30 | -30 | 4 | -36 |
| | L Parahippocampal Gyrus, anterior division | 3.17 | -30 | 0 | -34 |
| 40 | | | | | |
| | L Temporal Fusiform Cortex, posterior division | 3.91 | -40 | -14 | -28 |
| 40 | | | | | |
| | L Parahippocampal Gyrus, anterior division | 3.45 | -20 | -6 | -30 |
| 40 | | | | | |
| | R Subcallosal Cortex | 3.66 | 4 | 30 | -20 |
| 40 | | | | | |
| | L Inferior Temporal Gyrus, temporooccipital part | 3.17 | -50 | -56 | -14 |
| 40 | | | | | |
| | R Inferior Frontal Gyrus, pars triangularis | 2.94 | 52 | 28 | 4 |
| 40 | | | | | |
| | R Cingulate Gyrus, anterior division | 2.89 | 0 | 34 | 14 |
| 40 | | | | | |

| | | | | | |
|---|---|---|---|---|---|
| | L Inferior Frontal Gyrus, pars opercularis | 3.39 | -46 | 8 | 28 |
| 40 | | | | | |
| | R Superior Frontal Gyrus | 3.60 | 0 | 42 | 42 |
| 40 | | | | | |
| | L Superior Frontal Gyrus | 3.14 | -22 | 28 | 52 |
| 40 | | | | | |
| | R Superior Parietal Lobule | 3.14 | 26 | -56 | 60 |
| 32 | | | | | |
| | R Temporal Pole | 3.12 | 28 | 14 | -42 |
| 32 | | | | | |
| | R Amygdala | 3.04 | 24 | 4 | -20 |
| 32 | | | | | |
| | R Planum Polare | 3.76 | 42 | -2 | -16 |
| 32 | | | | | |
| | R Insular Cortex | 3.72 | 38 | 6 | -16 |
| 32 | | | | | |
| | L Insular Cortex | 3.89 | -40 | 2 | -12 |
| 32 | | | | | |
| | L Subcallosal Cortex | 2.99 | -2 | 16 | -6 |
| 32 | | | | | |
| | R Frontal Pole | 3.33 | 30 | 62 | -6 |
| 32 | | | | | |
| | L Superior Temporal Gyrus, posterior division | 3.23 | -68 | -24 | 2 |
| 32 | | | | | |
| | L Thalamus | 3.46 | -6 | -34 | 2 |
| 32 | | | | | |
| | R Central Opercular Cortex | 2.86 | 56 | -12 | 14 |
| 32 | | | | | |
| | R Parietal Operculum Cortex | 3.65 | 36 | -28 | 20 |
| 32 | | | | | |
| | R Superior Frontal Gyrus | 3.00 | 2 | 44 | 48 |
| 32 | | | | | |
| | L Lateral Occipital Cortex, superior division | 2.89 | -16 | -64 | 58 |
| 32 | | | | | |
| | R Superior Frontal Gyrus | 3.06 | 26 | 14 | 58 |
| 24 | | | | | |
| | R Brain-Stem | 3.22 | 2 | -36 | -66 |
| 24 | | | | | |
| | R Brain-Stem | 3.31 | 8 | -30 | -34 |
| 24 | | | | | |
| | L Parahippocampal Gyrus, anterior division | 3.59 | -18 | -12 | -28 |
| 24 | | | | | |

| | | | | | |
|---|---|---|---|---|---|
| | L Frontal Orbital Cortex | 3.14 | -24 | 14 | -24 |
| 24 | | | | | |
| | L Temporal Pole | 3.25 | -52 | 12 | -20 |
| 24 | | | | | |
| | R Frontal Orbital Cortex | 3.46 | 14 | 16 | -22 |
| 24 | | | | | |
| | L Temporal Occipital Fusiform Cortex | 3.10 | -34 | -60 | -16 |
| 24 | | | | | |
| | R Subcallosal Cortex | 3.09 | 2 | 12 | -10 |
| 24 | | | | | |
| | R Subcallosal Cortex | 3.07 | 4 | 30 | -8 |
| 24 | | | | | |
| | R Paracingulate Gyrus | 3.25 | 10 | 46 | -6 |
| 24 | | | | | |
| | R Superior Temporal Gyrus, posterior division | 3.02 | 60 | -28 | 2 |
| 24 | | | | | |
| | R Paracingulate Gyrus | 3.40 | 10 | 44 | 0 |
| 24 | | | | | |
| | L Paracingulate Gyrus | 3.38 | -4 | 44 | 2 |
| 24 | | | | | |
| | R Central Opercular Cortex | 2.93 | 60 | -8 | 10 |
| 24 | | | | | |
| | R Lateral Occipital Cortex, superior division | 3.31 | 36 | -82 | 12 |
| 24 | | | | | |
| | R Lateral Occipital Cortex, superior division | 3.01 | 28 | -80 | 18 |
| 24 | | | | | |
| | L Frontal Pole | 2.92 | -6 | 48 | 50 |
| 16 | | | | | |
| | L Brain-Stem | 2.84 | -2 | -32 | -60 |
| 16 | | | | | |
| | L Parahippocampal Gyrus, anterior division | 3.44 | -22 | -14 | -34 |
| 16 | | | | | |
| | R Brain-Stem | 2.87 | 16 | -32 | -28 |
| 16 | | | | | |
| | L Amygdala | 3.59 | -28 | 0 | -28 |
| 16 | | | | | |
| | L Temporal Fusiform Cortex, posterior division | 3.24 | -34 | -24 | -24 |
| 16 | | | | | |
| | R Inferior Temporal Gyrus, posterior division | 3.01 | 44 | -24 | -22 |

| | | | | | |
|---|---|---|---|---|---|
| 16 | | | | | |
| | R Frontal Medial Cortex | 3.09 | 8 | 48 | -20 |
| 16 | | | | | |
| | R Frontal Orbital Cortex | 2.89 | 22 | 16 | -18 |
| 16 | | | | | |
| | R Frontal Pole | 3.03 | 6 | 56 | -10 |
| 16 | | | | | |
| | L Frontal Pole | 2.89 | -40 | 58 | -10 |
| 16 | | | | | |
| | R Hippocampus | 2.89 | 24 | -28 | -8 |
| 16 | | | | | |
| | L Frontal Pole | 2.89 | -54 | 40 | -6 |
| 16 | | | | | |
| | R Frontal Pole | 3.49 | 4 | 62 | -4 |
| 16 | | | | | |
| | R Lateral Occipital Cortex, inferior division | 3.10 | 56 | -62 | -4 |
| 16 | | | | | |
| | L Middle Temporal Gyrus, posterior division | 3.07 | -62 | -44 | -2 |
| 16 | | | | | |
| | R Insular Cortex | 2.90 | 42 | -2 | 2 |
| 16 | | | | | |
| | R Frontal Pole | 3.19 | 42 | 38 | 6 |
| 16 | | | | | |
| | L Frontal Pole | 3.11 | -46 | 44 | 8 |
| 16 | | | | | |
| | L Inferior Frontal Gyrus, pars opercularis | 2.87 | -56 | 22 | 16 |
| 16 | | | | | |
| | R Cingulate Gyrus, anterior division | 3.30 | 2 | 26 | 16 |
| 16 | | | | | |
| | L Angular Gyrus | 3.05 | -60 | -56 | 22 |
| 16 | | | | | |
| | R Middle Frontal Gyrus | 3.46 | 44 | 28 | 22 |
| 16 | | | | | |
| | L Lateral Occipital Cortex, superior division | 2.92 | -50 | -70 | 24 |
| 16 | | | | | |
| | L Frontal Pole | 2.87 | -10 | 60 | 36 |
| 16 | | | | | |
| | L Postcentral Gyrus | 3.39 | -60 | -26 | 42 |
| 16 | | | | | |
| | R Middle Frontal Gyrus | 3.17 | 36 | 16 | 56 |
| 16 | | | | | |
| | L Superior Frontal Gyrus | 3.42 | -18 | 22 | 56 |
| 16 | | | | | |

| | | | | | |
|---|---|---|---|---|---|
| | R Postcentral Gyrus | 3.11 | 10 | -44 | 60 |
| 16 | | | | | |
| | R Superior Frontal Gyrus | 3.00 | 20 | 22 | 60 |
| 16 | | | | | |
| | R Superior Parietal Lobule | 3.31 | 24 | -52 | 68 |
| 8 | | | | | |
| | L Brain-Stem | 3.06 | -8 | -36 | -60 |
| 8 | | | | | |
| | L Temporal Pole | 3.01 | -24 | 10 | -32 |
| 8 | | | | | |
| | L Brain-Stem | 2.91 | -14 | -22 | -24 |
| 8 | | | | | |
| | L Temporal Pole | 3.20 | -48 | 4 | -24 |
| 8 | | | | | |
| | L Hippocampus | 2.84 | -16 | -8 | -22 |
| 8 | | | | | |
| | L Frontal Medial Cortex | 3.06 | -8 | 38 | -22 |
| 8 | | | | | |
| | L Frontal Medial Cortex | 3.09 | -2 | 42 | -22 |
| 8 | | | | | |
| | R Temporal Pole | 3.32 | 42 | 6 | -20 |
| 8 | | | | | |
| | L Frontal Orbital Cortex | 3.07 | -24 | 8 | -20 |
| 8 | | | | | |
| | R Frontal Orbital Cortex | 2.84 | 32 | 26 | -20 |
| 8 | | | | | |
| | L Occipital Fusiform Gyrus | 2.75 | -20 | -82 | -18 |
| 8 | | | | | |
| | L Temporal Occipital Fusiform Cortex | 2.82 | -34 | -54 | -18 |
| 8 | | | | | |
| | R Temporal Fusiform Cortex, posterior division | 2.92 | 40 | -26 | -18 |
| 8 | | | | | |
| | R Frontal Medial Cortex | 2.83 | 0 | 40 | -14 |
| 8 | | | | | |
| | L Frontal Pole | 2.96 | -12 | 60 | -14 |
| 8 | | | | | |
| | L Subcallosal Cortex | 2.82 | -2 | 28 | -12 |
| 8 | | | | | |
| | L Occipital Fusiform Gyrus | 2.88 | -30 | -80 | -10 |
| 8 | | | | | |
| | L Inferior Temporal Gyrus, temporooccipital part | 2.84 | -56 | -58 | -10 |
| 8 | | | | | |
| | R Subcallosal Cortex | 3.16 | 6 | 20 | -10 |

| | | | | | |
|---|---|---|---|---|---|
| 8 | | | | | |
| | R Frontal Medial Cortex | 2.81 | 8 | 52 | -10 |
| 8 | | | | | |
| | L Middle Temporal Gyrus, temporooccipital part | 2.85 | -58 | -54 | -8 |
| 8 | | | | | |
| | R Frontal Medial Cortex | 2.94 | 10 | 54 | -8 |
| 8 | | | | | |
| | L Frontal Pole | 3.13 | -44 | 56 | -8 |
| 8 | | | | | |
| | R Middle Temporal Gyrus, temporooccipital part | 2.83 | 70 | -42 | -4 |
| 8 | | | | | |
| | R Inferior Frontal Gyrus, pars triangularis | 3.09 | 52 | 30 | -4 |
| 8 | | | | | |
| | L Cingulate Gyrus, anterior division | 2.84 | -2 | 36 | -4 |
| 8 | | | | | |
| | R Frontal Pole | 3.00 | 8 | 58 | -4 |
| 8 | | | | | |
| | R Frontal Pole | 2.99 | 36 | 60 | -4 |
| 8 | | | | | |
| | R Thalamus | 3.14 | 14 | -34 | -2 |
| 8 | | | | | |
| | R Lingual Gyrus | 2.82 | 4 | -64 | 0 |
| 8 | | | | | |
| | L Middle Temporal Gyrus, temporooccipital part | 2.80 | -60 | -48 | 0 |
| 8 | | | | | |
| | L Caudate | 3.09 | -18 | 22 | 6 |
| 8 | | | | | |
| | R Precuneus Cortex | 2.81 | 4 | -54 | 10 |
| 8 | | | | | |
| | L Lateral Occipital Cortex, superior division | 2.79 | -54 | -70 | 16 |
| 8 | | | | | |
| | R Central Opercular Cortex | 2.93 | 50 | -8 | 16 |
| 8 | | | | | |
| | R Angular Gyrus | 2.85 | 56 | -54 | 18 |
| 8 | | | | | |
| | R Angular Gyrus | 2.85 | 54 | -56 | 20 |
| 8 | | | | | |
| | L Angular Gyrus | 2.82 | -62 | -54 | 20 |
| 8 | | | | | |
| | L Central Opercular Cortex | 2.86 | -40 | -18 | 20 |
| 8 | | | | | |

| | | | | | |
|---|---|---|---|---|---|
| | R Superior Frontal Gyrus | 2.88 | 6 | 54 | 28 |
| 8 | | | | | |
| | L Middle Frontal Gyrus | 2.85 | -50 | 14 | 30 |
| 8 | | | | | |
| | R Precentral Gyrus | 2.76 | 48 | 4 | 32 |
| 8 | | | | | |
| | R Supramarginal Gyrus, anterior division | 2.75 | 66 | -26 | 38 |
| 8 | | | | | |
| | R Frontal Pole | 2.90 | 8 | 56 | 38 |
| 8 | | | | | |
| | R Frontal Pole | 2.85 | 10 | 58 | 40 |
| 8 | | | | | |
| | L Middle Frontal Gyrus | 2.78 | -30 | 22 | 46 |
| 8 | | | | | |
| | L Frontal Pole | 2.75 | -18 | 46 | 46 |
| 8 | | | | | |
| | R Juxtapositional Lobule Cortex (formerly Supplementary Motor Cortex) | 2.76 | 0 | -12 | 48 |
| 8 | | | | | |
| | R Middle Frontal Gyrus | 2.79 | 36 | 12 | 48 |
| 8 | | | | | |
| | L Lateral Occipital Cortex, superior division | 2.82 | -20 | -76 | 50 |
| 8 | | | | | |
| | L Superior Frontal Gyrus | 2.86 | -22 | -2 | 52 |
| 8 | | | | | |
| | R Superior Parietal Lobule | 3.10 | 34 | -54 | 56 |
| 8 | | | | | |
| | L Superior Parietal Lobule | 2.99 | -16 | -56 | 60 |
| 8 | | | | | |
| | R Postcentral Gyrus | 2.92 | 6 | -32 | 80 |
| 8 | | | | | |
| | L Precentral Gyrus | 2.81 | -10 | -28 | 80 |
| 8 | | | | | |
| | L Precentral Gyrus | 2.88 | -8 | -20 | 80 |

**Supplementary Table S31. [$CT_{CB3}$ - $CS_{CB3}$] > [$UT_{CB2}$ - $US_{CB2}$].** Descriptive statistics for clusters and local maxima showing greater increases in activation during Convolved Block 3 of Certain-Threat anticipation compared to Convolved Block 2 of Uncertain-Threat anticipation, each relative to their respective reference conditions (FDR *q*<0.05, whole-brain corrected).

| | | **mm³** | ***t*** | ***x*** | ***y*** | ***z*** |
|---|---|---|---|---|---|---|
| Cluster 1 | L Postcentral Gyrus | 26448 | 7.99 | -44 | -14 | 34 |
| | L Precentral Gyrus | | 7.49 | -48 | -14 | 40 |
| | L Juxtapositional Lobule Cortex (formerly Supplementary Motor Cortex) | | 5.53 | -8 | -12 | 48 |
| | L Precuneus Cortex | | 5.25 | -6 | -72 | 46 |
| | L Lateral Occipital Cortex, superior division | | 4.61 | -6 | -74 | 58 |
| | L Cuneal Cortex | | 4.26 | -10 | -76 | 26 |
| | L Central Opercular Cortex | | 3.32 | -60 | -8 | 8 |
| | R Precentral Gyrus | | 7.30 | 6 | -24 | 64 |
| | R Postcentral Gyrus | | 5.21 | 40 | -22 | 52 |
| | R Cingulate Gyrus, posterior division | | 4.21 | 10 | -34 | 46 |
| | R Precuneus Cortex | | 3.80 | 12 | -50 | 38 |
| | R Juxtapositional Lobule Cortex (formerly Supplementary Motor Cortex) | | 3.68 | 6 | -14 | 62 |
| Cluster 2 | R Amygdala | 14800 | 3.56 | 28 | 0 | -16 |
| | R Amygdala | | 3.46 | 26 | -4 | -12 |
| | R Amygdala | | 3.32 | 22 | 2 | -18 |
| | R Amygdala | | 2.91 | 16 | -4 | -16 |
| | L Amygdala | | 4.09 | -26 | -2 | -12 |
| | L Amygdala | | 3.81 | -20 | -2 | -18 |
| | L Amygdala | | 3.71 | -22 | 0 | -16 |
| | L Central Opercular Cortex | | 7.34 | -38 | -12 | 18 |
| | L Caudate | | 6.91 | -14 | 22 | -2 |
| | L Accumbens | | 5.78 | -6 | 10 | -4 |
| | L Insular Cortex | | 5.68 | -38 | 4 | -16 |
| | L Planum Polare | | 4.99 | -42 | -10 | -8 |

| | | | | | | |
|---|---|---|---|---|---|---|
| | L Putamen | | 4.74 | -22 | 14 | -6 |
| | L Temporal Pole | | 4.53 | -50 | 8 | -6 |
| | L Amygdala | | 4.09 | -26 | -2 | -12 |
| | R Insular Cortex | | 6.19 | 40 | 0 | -12 |
| | R Putamen | | 5.66 | 22 | 10 | -4 |
| | R Accumbens | | 5.64 | 14 | 16 | -6 |
| | R Caudate | | 5.42 | 14 | 20 | -2 |
| | R Frontal Orbital Cortex | | 4.81 | 28 | 10 | -22 |
| | R Pallidum | | 4.54 | 16 | 6 | 0 |
| | R Temporal Pole | | 4.31 | 34 | 8 | -22 |
| | R Amygdala | | 3.56 | 28 | 0 | -16 |
| | R BST | | 4.52 | 8 | 4 | 2 |
| Cluster 3 | R Precuneus Cortex | 5728 | 5.44 | 16 | -66 | 22 |
| | R Cuneal Cortex | | 5.01 | 12 | -70 | 26 |
| | R Lateral Occipital Cortex, superior division | | 4.12 | 18 | -74 | 48 |
| | R Intracalcarine Cortex | | 3.94 | 14 | -78 | 4 |
| | R Lingual Gyrus | | 3.90 | 12 | -50 | 2 |
| Cluster 3 | R Cingulate Gyrus, posterior division | 5728 | 2.96 | 16 | -50 | 4 |
| Cluster 4 | L Thalamus | 4128 | 5.81 | -2 | -20 | 2 |
| | L Brain-Stem | | 5.55 | -4 | -36 | -6 |
| | R Thalamus | | 5.26 | 4 | -14 | 6 |
| | R Brain-Stem | | 3.55 | 2 | -32 | -2 |
| Cluster 5 | R Precentral Gyrus | 4096 | 6.29 | 44 | -10 | 34 |
| | R Postcentral Gyrus | | 4.51 | 68 | -8 | 20 |
| Cluster 7 | R Superior Temporal Gyrus, anterior division | 1248 | 5.17 | 58 | -4 | -12 |
| | R Middle Temporal Gyrus, posterior division | | 3.20 | 58 | -10 | -12 |
| Cluster 8 | L Lateral Occipital Cortex, superior division | 1120 | 4.91 | -40 | -70 | 26 |
| Cluster 9 | L Intracalcarine Cortex | 832 | 4.64 | -14 | -70 | 8 |

| | | | | | | |
|---|---|---|---|---|---|---|
| | L Lingual Gyrus | | 4.44 | -10 | -62 | 4 |
| Cluster 10 | R Insular Cortex | 816 | 5.03 | 40 | -4 | 8 |
| | R Central Opercular Cortex | | 4.58 | 42 | -12 | 16 |
| Cluster 11 | L Pallidum | 760 | 4.78 | -20 | -2 | 0 |
| | L Putamen | | 4.00 | -20 | 8 | 2 |
| Cluster 12 | L Parietal Operculum Cortex | 728 | 5.14 | -44 | -34 | 22 |
| | L Planum Temporale | | 4.66 | -48 | -38 | 18 |
| | L Supramarginal Gyrus, posterior division | | 4.38 | -56 | -42 | 22 |
| Cluster 13 | R Amygdala | 592 | 4.16 | 22 | -14 | -12 |
| | R Amygdala | | 4.16 | 22 | -14 | -12 |
| | R Hippocampus | | 3.87 | 26 | -20 | -14 |
| Cluster 15 | L Middle Frontal Gyrus | 568 | 4.27 | -26 | 36 | 32 |
| | L Frontal Pole | | 3.74 | -34 | 40 | 36 |
| Cluster 20 | L Angular Gyrus | 408 | 4.16 | -50 | -52 | 16 |
| Cluster 21 | R Lingual Gyrus | 384 | 4.35 | 24 | -54 | 4 |
| Cluster 21 | R Intracalcarine Cortex | 384 | 3.58 | 22 | -64 | 6 |
| Cluster 22 | R Frontal Pole | 376 | 4.62 | 32 | 34 | -14 |
| Cluster 23 | L Middle Frontal Gyrus | 352 | 4.80 | -46 | 30 | 24 |
| Cluster 24 | L Middle Temporal Gyrus, temporooccipital part | 328 | 4.12 | -60 | -60 | 2 |
| Cluster 25 | L Frontal Orbital Cortex | 312 | 4.22 | -28 | 36 | -12 |
| | L Frontal Pole | | 3.76 | -24 | 38 | -14 |
| Cluster 30 | L Middle Frontal Gyrus | 208 | 3.83 | -44 | 18 | 32 |
| Cluster 31 | L Brain-Stem | 200 | 3.32 | -4 | -38 | -20 |
| Cluster 32 | R Brain-Stem | 200 | 4.68 | 0 | -20 | -20 |
| Cluster 33 | R Hippocampus | 192 | 4.58 | 24 | -38 | 2 |
| Cluster 36 | R Cingulate Gyrus, anterior division | 184 | 4.16 | 0 | -2 | 32 |
| Cluster 37 | R Frontal Pole | 176 | 4.20 | 2 | 56 | -16 |
| | R Frontal Medial Cortex | | 3.62 | 2 | 52 | -18 |
| Cluster 38 | L Thalamus | 176 | 4.92 | -22 | -36 | 0 |

| | | | | | | |
|---|---|---|---|---|---|---|
| Cluster 42 | L Occipital Fusiform Gyrus | 168 | 3.98 | -20 | -86 | -8 |
| | L Occipital Pole | | 3.84 | -16 | -92 | -10 |
| Cluster 43 | R Frontal Medial Cortex | 160 | 4.22 | 4 | 40 | -26 |
| Cluster 45 | L Lingual Gyrus | 152 | 3.03 | -14 | -58 | 0 |
| Cluster 46 | L Lateral Occipital Cortex, superior division | 152 | 3.45 | -26 | -80 | 50 |
| Cluster 47 | R Precuneus Cortex | 152 | 3.68 | 8 | -64 | 56 |
| Cluster 49 | R Brain-Stem | 144 | 3.97 | 8 | -24 | -32 |
| Cluster 50 | L Superior Frontal Gyrus | 144 | 3.91 | -20 | -10 | 58 |
| Cluster 51 | R Temporal Fusiform Cortex, posterior division | 136 | 5.36 | 34 | -16 | -32 |
| Cluster 54 | R Brain-Stem | 128 | 4.01 | 6 | -40 | -52 |
| Cluster 56 | L Hippocampus | 128 | 4.51 | -24 | -12 | -26 |
| | L Amygdala | | 5.56 | -18 | -12 | -14 |
| | L Amygdala | | 5.56 | -18 | -12 | -14 |
| | L Hippocampus | | 3.15 | -22 | -14 | -18 |
| Cluster 58 | R Temporal Fusiform Cortex, posterior division | 120 | 3.14 | 42 | -16 | -38 |
| | R Inferior Temporal Gyrus, posterior division | | 3.03 | 48 | -10 | -38 |
| | R Inferior Temporal Gyrus, anterior division | | 3.01 | 50 | -6 | -38 |
| Cluster 60 | L Hippocampus | 120 | 3.55 | -26 | -34 | -8 |
| Cluster 61 | R Superior Temporal Gyrus, posterior division | 120 | 4.28 | 68 | -20 | 6 |
| Cluster 62 | R Frontal Pole | 120 | 3.42 | 30 | 40 | 30 |
| Cluster 63 | L Precentral Gyrus | 120 | 4.80 | -34 | -14 | 68 |
| Cluster 64 | L Lingual Gyrus | 112 | 4.37 | -18 | -52 | -2 |
| Cluster 65 | L Insular Cortex | 112 | 3.83 | -38 | -16 | -4 |
| Cluster 66 | R Insular Cortex | 112 | 4.46 | 36 | -20 | 4 |
| Cluster 67 | L Frontal Pole | 112 | 3.62 | -40 | 40 | 14 |
| Cluster 68 | L Middle Temporal Gyrus, posterior division | 104 | 3.85 | -62 | -22 | -12 |

| | | | | | | |
|---|---|---|---|---|---|---|
| Cluster 70 | R Cingulate Gyrus, anterior division | 104 | 3.62 | 4 | 34 | 10 |
| Cluster 71 | L Supramarginal Gyrus, anterior division | 104 | 3.98 | -64 | -38 | 24 |
| Cluster 72 | L Supramarginal Gyrus, anterior division | 104 | 3.43 | -56 | -34 | 36 |
| Cluster 73 | L Cingulate Gyrus, posterior division | 104 | 3.44 | -10 | -32 | 38 |
| Cluster 74 | L Thalamus | 96 | 4.80 | -14 | -24 | 0 |
| Cluster 75 | L Lateral Occipital Cortex, superior division | 96 | 3.65 | -32 | -60 | 40 |
| Cluster 78 | L Planum Temporale | 88 | 3.60 | -54 | -36 | 12 |
| Cluster 79 | R Lateral Occipital Cortex, superior division | 88 | 3.67 | 44 | -60 | 20 |
| Cluster 80 | R Lateral Occipital Cortex, superior division | 88 | 3.20 | 42 | -70 | 34 |
| Cluster 81 | L Supramarginal Gyrus, posterior division | 88 | 3.21 | -36 | -48 | 40 |
| Cluster 82 | L Lateral Occipital Cortex, superior division | 88 | 3.52 | -30 | -64 | 42 |
| Cluster 86 | L Occipital Fusiform Gyrus | 80 | 3.86 | -26 | -88 | -12 |
| Cluster 87 | L Central Opercular Cortex | 80 | 3.34 | -52 | -12 | 14 |
| Cluster 89 | L Cingulate Gyrus, posterior division | 80 | 2.84 | -2 | -20 | 28 |
| Cluster 90 | L Superior Frontal Gyrus | 80 | 3.55 | -20 | 0 | 54 |
| Cluster 92 | R Brain-Stem | 72 | 3.38 | 0 | -30 | -50 |
| Cluster 93 | R Brain-Stem | 72 | 3.67 | 14 | -20 | -28 |
| Cluster 95 | L Putamen | 72 | 3.68 | -26 | -8 | -6 |
| Cluster 99 | R Parahippocampal Gyrus, anterior division | 64 | 3.59 | 24 | -6 | -34 |
| Cluster 101 | L Inferior Temporal Gyrus, posterior division | 64 | 3.80 | -60 | -42 | -24 |
| Cluster 102 | L Hippocampus | 64 | 3.96 | -34 | -16 | -18 |
| Cluster 104 | L Frontal Pole | 64 | 4.03 | -12 | 68 | -8 |
| Cluster 105 | R Frontal Pole | 64 | 3.79 | 2 | 70 | 2 |
| Cluster 107 | R Parietal Operculum Cortex | 64 | 3.11 | 38 | -26 | 18 |
| Cluster 108 | L Cingulate Gyrus, anterior division | 64 | 3.54 | -8 | 36 | 22 |

| | | | | | | |
|---|---|---|---|---|---|---|
| Cluster 109 | L Frontal Pole | 64 | 3.15 | -38 | 50 | 22 |
| Cluster 110 | L Occipital Pole | 64 | 3.72 | -34 | -90 | 26 |
| Cluster 113 | R Inferior Temporal Gyrus, temporooccipital part | 56 | 3.50 | 54 | -42 | -26 |
| Cluster 115 | R Frontal Pole | 56 | 3.23 | 2 | 66 | -10 |
| Cluster 116 | R Planum Polare | 56 | 3.65 | 50 | 4 | -8 |
| Cluster 117 | R Occipital Pole | 56 | 3.60 | 28 | -94 | -4 |
| Cluster 118 | R Frontal Pole | 56 | 3.11 | 2 | 58 | -4 |
| Cluster 119 | R Thalamus | 56 | 4.79 | 18 | -26 | 12 |
| Cluster 120 | L Supramarginal Gyrus, posterior division | 56 | 3.35 | -58 | -50 | 28 |
| Cluster 121 | R Cingulate Gyrus, posterior division | 56 | 3.51 | 6 | -38 | 34 |
| Cluster 122 | R Lateral Occipital Cortex, superior division | 56 | 3.27 | 34 | -68 | 44 |
| Cluster 126 | L Temporal Pole | 48 | 3.56 | -42 | 14 | -22 |
| Cluster 127 | R Middle Temporal Gyrus, posterior division | 48 | 3.74 | 62 | -34 | -14 |
| Cluster 128 | L Occipital Pole | 48 | 3.47 | -28 | -100 | -12 |
| Cluster 129 | R Lingual Gyrus | 48 | 3.52 | 20 | -46 | -4 |
| Cluster 130 | L Paracingulate Gyrus | 48 | 3.40 | -6 | 22 | 34 |
| Cluster 131 | L Frontal Pole | 48 | 2.99 | -26 | 44 | 44 |
| Cluster 132 | L Lateral Occipital Cortex, superior division | 48 | 4.02 | -26 | -60 | 46 |
| Cluster 133 | R Middle Frontal Gyrus | 48 | 3.10 | 32 | 0 | 54 |
| Cluster 134 | L Juxtapositional Lobule Cortex (formerly Supplementary Motor Cortex) | 48 | 4.09 | -4 | 2 | 64 |
| Cluster 143 | L Brain-Stem | 40 | 3.09 | -10 | -32 | -28 |
| Cluster 145 | R Hippocampus | 40 | 3.79 | 36 | -20 | -14 |
| Cluster 147 | R Frontal Pole | 40 | 4.07 | 14 | 62 | -10 |
| Cluster 148 | R Paracingulate Gyrus | 40 | 3.05 | 6 | 48 | 2 |
| Cluster 150 | R Hippocampus | 40 | 3.79 | 12 | -38 | 4 |
| Cluster 151 | R Central Opercular Cortex | 40 | 3.90 | 52 | -8 | 12 |

| | | | | | | |
|---|---|---|---|---|---|---|
| Cluster 152 | L Cingulate Gyrus, anterior division | 40 | 3.23 | -2 | 38 | 18 |
| Cluster 153 | R Cuneal Cortex | 40 | 3.15 | 4 | -80 | 20 |
| Cluster 155 | L Inferior Frontal Gyrus, pars triangularis | 40 | 3.31 | -44 | 24 | 22 |
| Cluster 156 | L Lateral Occipital Cortex, superior division | 40 | 3.33 | -38 | -76 | 34 |
| Cluster 157 | L Lateral Occipital Cortex, superior division | 40 | 2.92 | -28 | -82 | 34 |
| Cluster 158 | L Cingulate Gyrus, posterior division | 40 | 3.40 | -6 | -38 | 34 |
| Cluster 161 | L Brain-Stem | 32 | 3.45 | -6 | -46 | -48 |
| Cluster 164 | L Brain-Stem | 32 | 3.33 | -18 | -26 | -34 |
| Cluster 165 | R Middle Temporal Gyrus, posterior division | 32 | 3.25 | 62 | -10 | -28 |
| Cluster 169 | L Inferior Temporal Gyrus, temporooccipital part | 32 | 3.33 | -64 | -54 | -14 |
| Cluster 170 | R Frontal Medial Cortex | 32 | 3.25 | 2 | 50 | -12 |
| Cluster 171 | L Frontal Pole | 32 | 3.65 | -14 | 62 | -10 |
| Cluster 172 | L Insular Cortex | 32 | 3.44 | -30 | 22 | 0 |
| Cluster 173 | L Planum Temporale | 32 | 2.99 | -60 | -30 | 10 |
| Cluster 174 | R Frontal Pole | 32 | 2.99 | 46 | 44 | 14 |
| Cluster 175 | L Precuneus Cortex | 32 | 2.95 | -20 | -62 | 20 |
| Cluster 176 | L Occipital Pole | 32 | 3.34 | -12 | -102 | 22 |
| Cluster 177 | R Cingulate Gyrus, posterior division | 32 | 2.96 | 2 | -36 | 26 |
| Cluster 179 | R Cingulate Gyrus, posterior division | 32 | 3.38 | 0 | -52 | 28 |
| Cluster 180 | R Frontal Pole | 32 | 3.17 | 24 | 46 | 26 |
| Cluster 181 | R Precentral Gyrus | 32 | 3.87 | 58 | 12 | 30 |
| Cluster 183 | L Lateral Occipital Cortex, superior division | 32 | 3.02 | -42 | -74 | 42 |
| Cluster 185 | L Precuneus Cortex | 32 | 3.16 | -4 | -62 | 58 |
| Cluster 189 | R Parahippocampal Gyrus, anterior division | 24 | 3.05 | 26 | -10 | -38 |
| Cluster 193 | L Temporal Pole | 24 | 3.49 | -34 | 10 | -30 |
| Cluster 195 | R Inferior Temporal Gyrus, posterior division | 24 | 3.65 | 50 | -34 | -22 |

| | | | | | | |
|---|---|---|---|---|---|---|
| Cluster 197 | R Brain-Stem | 24 | 3.63 | 8 | -24 | -22 |
| Cluster 198 | L Lateral Occipital Cortex, inferior division | 24 | 3.34 | -50 | -76 | -16 |
| Cluster 199 | R Frontal Orbital Cortex | 24 | 3.01 | 26 | 30 | -14 |
| Cluster 200 | R Middle Temporal Gyrus, posterior division | 24 | 3.03 | 58 | -22 | -10 |
| Cluster 203 | R Brain-Stem | 24 | 3.54 | 8 | -34 | -8 |
| Cluster 204 | L Thalamus | 24 | 3.21 | -2 | -6 | -2 |
| Cluster 206 | R Insular Cortex | 24 | 3.22 | 34 | 18 | 0 |
| Cluster 207 | R Middle Temporal Gyrus, temporooccipital part | 24 | 3.03 | 66 | -46 | 2 |
| Cluster 209 | R Superior Temporal Gyrus, posterior division | 24 | 3.07 | 66 | -34 | 4 |
| Cluster 210 | R Cingulate Gyrus, posterior division | 24 | 3.46 | 6 | -42 | 8 |
| Cluster 211 | R Occipital Pole | 24 | 2.93 | 0 | -92 | 14 |
| Cluster 212 | L Lateral Occipital Cortex, superior division | 24 | 2.92 | -36 | -82 | 14 |
| Cluster 213 | L Caudate | 24 | 3.04 | -14 | 2 | 14 |
| Cluster 214 | R Angular Gyrus | 24 | 3.32 | 60 | -56 | 18 |
| Cluster 215 | L Planum Temporale | 24 | 2.92 | -64 | -34 | 18 |
| Cluster 216 | R Cingulate Gyrus, anterior division | 24 | 3.35 | 4 | 12 | 26 |
| Cluster 217 | L Cingulate Gyrus, anterior division | 24 | 3.31 | -2 | 14 | 24 |
| Cluster 218 | L Paracingulate Gyrus | 24 | 3.00 | -2 | 44 | 24 |
| Cluster 220 | R Lateral Occipital Cortex, superior division | 24 | 2.95 | 54 | -66 | 28 |
| Cluster 221 | L Cingulate Gyrus, anterior division | 24 | 3.00 | -6 | 20 | 30 |
| Cluster 222 | L Middle Frontal Gyrus | 24 | 3.32 | -54 | 18 | 30 |
| Cluster 224 | L Middle Frontal Gyrus | 24 | 3.05 | -26 | 24 | 36 |
| Cluster 225 | L Postcentral Gyrus | 24 | 3.28 | -34 | -38 | 42 |
| Cluster 226 | R Lateral Occipital Cortex, superior division | 24 | 3.12 | 14 | -72 | 58 |
| Cluster 227 | R Postcentral Gyrus | 24 | 3.14 | 38 | -30 | 60 |
| Cluster 235 | R Brain-Stem | 16 | 2.97 | 2 | -36 | -28 |

| | | | | | | |
|---|---|---|---|---|---|---|
| Cluster 237 | L Middle Temporal Gyrus, anterior division | 16 | 2.99 | -56 | -8 | -26 |
| Cluster 238 | R Temporal Fusiform Cortex, posterior division | 16 | 3.01 | 42 | -16 | -24 |
| Cluster 239 | L Subcallosal Cortex | 16 | 3.08 | -2 | 30 | -24 |
| Cluster 241 | L Temporal Occipital Fusiform Cortex | 16 | 2.93 | -40 | -48 | -18 |
| Cluster 242 | L Inferior Temporal Gyrus, posterior division | 16 | 2.99 | -60 | -30 | -20 |
| Cluster 244 | L Subcallosal Cortex | 16 | 3.21 | -2 | 22 | -14 |
| Cluster 245 | R Inferior Temporal Gyrus, temporooccipital part | 16 | 3.18 | 62 | -58 | -12 |
| Cluster 246 | R Temporal Occipital Fusiform Cortex | 16 | 3.80 | 32 | -44 | -12 |
| Cluster 247 | L Temporal Pole | 16 | 3.23 | -48 | 8 | -12 |
| Cluster 248 | L Frontal Pole | 16 | 2.98 | -26 | 64 | -12 |
| Cluster 249 | R Lingual Gyrus | 16 | 2.86 | 26 | -46 | -8 |
| Cluster 250 | R Middle Temporal Gyrus, posterior division | 16 | 3.08 | 50 | -20 | -8 |
| Cluster 251 | L Lingual Gyrus | 16 | 3.43 | -14 | -78 | -6 |
| Cluster 252 | L Superior Temporal Gyrus, anterior division | 16 | 2.87 | -52 | -10 | -6 |
| Cluster 253 | R Planum Polare | 16 | 3.18 | 44 | -16 | -4 |
| Cluster 254 | R Insular Cortex | 16 | 3.18 | 40 | 14 | -2 |
| Cluster 255 | L Subcallosal Cortex | 16 | 3.59 | -2 | 30 | -2 |
| Cluster 256 | L Frontal Pole | 16 | 3.29 | -14 | 70 | -2 |
| Cluster 257 | L Cingulate Gyrus, posterior division | 16 | 3.04 | -12 | -46 | 2 |
| Cluster 258 | L Hippocampus | 16 | 2.80 | -12 | -40 | 2 |
| Cluster 259 | R Thalamus | 16 | 3.13 | 20 | -20 | 2 |
| Cluster 260 | L Thalamus | 16 | 2.89 | -16 | -18 | 2 |
| Cluster 261 | L Paracingulate Gyrus | 16 | 3.14 | -4 | 48 | 4 |
| Cluster 262 | L Caudate | 16 | 2.93 | -8 | 12 | 8 |
| Cluster 263 | R Caudate | 16 | 3.07 | 10 | 18 | 6 |
| Cluster 264 | R Insular Cortex | 16 | 2.88 | 34 | 0 | 10 |
| Cluster 265 | L Planum Temporale | 16 | 2.96 | -50 | -32 | 10 |

| | | | | | | |
|---|---|---|---|---|---|---|
| Cluster 267 | L Parietal Operculum Cortex | 16 | 3.31 | -44 | -28 | 14 |
| Cluster 268 | L Occipital Pole | 16 | 3.22 | -12 | -98 | 16 |
| Cluster 269 | R Lateral Occipital Cortex, superior division | 16 | 3.03 | 44 | -70 | 18 |
| Cluster 270 | R Planum Temporale | 16 | 3.11 | 60 | -30 | 18 |
| Cluster 271 | R Precuneus Cortex | 16 | 2.80 | 0 | -64 | 20 |
| Cluster 273 | R Cingulate Gyrus, anterior division | 16 | 3.32 | 4 | 24 | 22 |
| Cluster 274 | R Lateral Occipital Cortex, superior division | 16 | 3.00 | 46 | -70 | 26 |
| Cluster 276 | R Precuneus Cortex | 16 | 3.06 | 4 | -64 | 26 |
| Cluster 277 | R Cingulate Gyrus, posterior division | 16 | 3.15 | 0 | -42 | 26 |
| Cluster 278 | R Cingulate Gyrus, posterior division | 16 | 3.11 | 6 | -24 | 28 |
| Cluster 279 | R Lateral Occipital Cortex, superior division | 16 | 2.93 | 22 | -86 | 30 |
| Cluster 280 | R Cingulate Gyrus, anterior division | 16 | 3.20 | 4 | 24 | 32 |
| Cluster 281 | L Cingulate Gyrus, posterior division | 16 | 3.20 | -10 | -38 | 36 |
| Cluster 282 | R Middle Frontal Gyrus | 16 | 3.18 | 38 | 26 | 34 |
| Cluster 283 | R Middle Frontal Gyrus | 16 | 3.07 | 26 | 30 | 34 |
| Cluster 284 | L Lateral Occipital Cortex, superior division | 16 | 2.87 | -40 | -62 | 38 |
| Cluster 285 | L Lateral Occipital Cortex, superior division | 16 | 2.98 | -22 | -62 | 42 |
| Cluster 286 | L Angular Gyrus | 16 | 2.95 | -38 | -58 | 40 |
| Cluster 287 | R Lateral Occipital Cortex, superior division | 16 | 2.84 | 32 | -80 | 48 |
| Cluster 288 | L Paracingulate Gyrus | 16 | 2.79 | -6 | 10 | 52 |
| Cluster 289 | L Juxtapositional Lobule Cortex (formerly Supplementary Motor Cortex) | 16 | 2.99 | -8 | 4 | 52 |
| Cluster 306 | L Brain-Stem | 8 | 2.98 | -2 | -26 | -42 |
| Cluster 307 | R Brain-Stem | 8 | 2.85 | 6 | -28 | -40 |
| Cluster 309 | L Brain-Stem | 8 | 3.11 | -4 | -20 | -36 |
| Cluster 310 | R Brain-Stem | 8 | 2.84 | 2 | -16 | -36 |
| Cluster 313 | L Brain-Stem | 8 | 2.80 | -6 | -42 | -34 |

| | | | | | | |
|---|---|---|---|---|---|---|
| Cluster 319 | R Brain-Stem | 8 | 2.88 | 0 | -32 | -32 |
| Cluster 320 | L Brain-Stem | 8 | 2.87 | -10 | -26 | -32 |
| Cluster 321 | L Temporal Fusiform Cortex, posterior division | 8 | 2.79 | -40 | -20 | -32 |
| Cluster 329 | L Brain-Stem | 8 | 2.82 | -2 | -32 | -28 |
| Cluster 330 | R Inferior Temporal Gyrus, posterior division | 8 | 3.05 | 58 | -20 | -28 |
| Cluster 331 | L Temporal Fusiform Cortex, posterior division | 8 | 2.77 | -42 | -18 | -26 |
| Cluster 333 | L Brain-Stem | 8 | 3.05 | -12 | -26 | -24 |
| Cluster 334 | L Parahippocampal Gyrus, anterior division | 8 | 2.94 | -12 | -4 | -24 |
| Cluster 335 | R Temporal Pole | 8 | 2.85 | 54 | 18 | -24 |
| Cluster 340 | L Brain-Stem | 8 | 2.80 | -8 | -24 | -22 |
| Cluster 341 | L Middle Temporal Gyrus, posterior division | 8 | 3.14 | -64 | -16 | -22 |
| Cluster 342 | L Frontal Orbital Cortex | 8 | 3.32 | -22 | 28 | -22 |
| Cluster 343 | L Inferior Temporal Gyrus, temporooccipital part | 8 | 2.77 | -44 | -52 | -20 |
| Cluster 344 | L Inferior Temporal Gyrus, posterior division | 8 | 3.00 | -64 | -34 | -20 |
| Cluster 345 | L Hippocampus | 8 | 2.88 | -16 | -10 | -20 |
| Cluster 348 | L Inferior Temporal Gyrus, posterior division | 8 | 3.23 | -54 | -34 | -18 |
| Cluster 349 | R Temporal Pole | 8 | 2.85 | 58 | 12 | -18 |
| Cluster 350 | L Occipital Pole | 8 | 2.76 | -32 | -92 | -16 |
| Cluster 352 | R Brain-Stem | 8 | 2.90 | 14 | -24 | -16 |
| Cluster 353 | R Planum Polare | 8 | 2.81 | 46 | 2 | -16 |
| Cluster 354 | L Lingual Gyrus | 8 | 2.89 | -12 | -78 | -14 |
| Cluster 355 | R Parahippocampal Gyrus, posterior division | 8 | 3.02 | 18 | -30 | -14 |
| Cluster 358 | L Middle Temporal Gyrus, posterior division | 8 | 2.86 | -60 | -26 | -12 |
| Cluster 359 | L Superior Temporal Gyrus, anterior division | 8 | 2.91 | -54 | -4 | -10 |

| | | | | | | |
|---|---|---|---|---|---|---|
| Cluster 361 | L Occipital Fusiform Gyrus | 8 | 3.04 | -20 | -76 | -8 |
| Cluster 362 | L Lingual Gyrus | 8 | 2.95 | -22 | -50 | -8 |
| Cluster 363 | R Occipital Pole | 8 | 2.86 | 26 | -98 | -6 |
| Cluster 364 | L Superior Temporal Gyrus, posterior division | 8 | 2.77 | -62 | -22 | -4 |
| Cluster 366 | R Lingual Gyrus | 8 | 2.78 | 10 | -60 | -2 |
| Cluster 367 | R Occipital Pole | 8 | 3.07 | 30 | -92 | 0 |
| Cluster 369 | R Frontal Pole | 8 | 3.14 | 22 | 68 | 0 |
| Cluster 370 | L Frontal Pole | 8 | 2.84 | -6 | 70 | 2 |
| Cluster 371 | L Central Opercular Cortex | 8 | 3.16 | -42 | 2 | 4 |
| Cluster 372 | R Inferior Frontal Gyrus, pars triangularis | 8 | 2.82 | 50 | 32 | 4 |
| Cluster 374 | R Middle Temporal Gyrus, temporooccipital part | 8 | 3.07 | 52 | -46 | 6 |
| Cluster 376 | L Insular Cortex | 8 | 2.90 | -36 | -20 | 6 |
| Cluster 377 | L Thalamus | 8 | 2.89 | -10 | -10 | 6 |
| Cluster 378 | R Thalamus | 8 | 3.10 | 22 | -28 | 8 |
| Cluster 379 | L Insular Cortex | 8 | 2.82 | -34 | 10 | 8 |
| Cluster 380 | R Frontal Pole | 8 | 2.81 | 22 | 66 | 8 |
| Cluster 381 | R Frontal Pole | 8 | 2.81 | 2 | 68 | 8 |
| Cluster 382 | L Middle Temporal Gyrus, temporooccipital part | 8 | 2.80 | -46 | -54 | 10 |
| Cluster 384 | R Thalamus | 8 | 3.05 | 14 | -20 | 14 |
| Cluster 385 | L Cingulate Gyrus, anterior division | 8 | 2.88 | -4 | 40 | 14 |
| Cluster 387 | L Lateral Occipital Cortex, superior division | 8 | 2.80 | -52 | -74 | 16 |
| Cluster 388 | L Angular Gyrus | 8 | 2.93 | -60 | -54 | 16 |
| Cluster 389 | R Planum Temporale | 8 | 2.92 | 56 | -30 | 16 |
| Cluster 391 | L Lateral Occipital Cortex, superior division | 8 | 2.96 | -26 | -88 | 18 |
| Cluster 392 | L Frontal Pole | 8 | 2.90 | -34 | 42 | 18 |
| Cluster 393 | R Lateral Occipital Cortex, superior division | 8 | 2.85 | 50 | -68 | 20 |

| | | | | | | |
|---|---|---|---|---|---|---|
| Cluster 394 | L Caudate | 8 | 2.96 | -14 | -10 | 20 |
| Cluster 395 | L Inferior Frontal Gyrus, pars opercularis | 8 | 2.93 | -44 | 18 | 20 |
| Cluster 396 | R Frontal Pole | 8 | 2.78 | 28 | 50 | 20 |
| Cluster 397 | L Frontal Pole | 8 | 2.82 | -22 | 58 | 20 |
| Cluster 398 | L Occipital Pole | 8 | 2.83 | -30 | -96 | 22 |
| Cluster 401 | L Lateral Occipital Cortex, superior division | 8 | 3.24 | -32 | -76 | 26 |
| Cluster 403 | R Cingulate Gyrus, posterior division | 8 | 2.81 | 8 | -50 | 26 |
| Cluster 404 | L Supramarginal Gyrus, posterior division | 8 | 3.02 | -64 | -46 | 26 |
| Cluster 405 | L Cingulate Gyrus, posterior division | 8 | 3.00 | -2 | -30 | 26 |
| Cluster 406 | L Cingulate Gyrus, anterior division | 8 | 2.93 | -4 | 28 | 26 |
| Cluster 407 | R Frontal Pole | 8 | 2.82 | 10 | 58 | 26 |
| Cluster 408 | L Frontal Pole | 8 | 3.01 | -24 | 62 | 26 |
| Cluster 409 | R Lateral Occipital Cortex, superior division | 8 | 2.89 | 44 | -64 | 28 |
| Cluster 410 | R Supramarginal Gyrus, anterior division | 8 | 3.15 | 58 | -24 | 30 |
| Cluster 411 | L Precentral Gyrus | 8 | 2.90 | -38 | 4 | 30 |
| Cluster 412 | R Lateral Occipital Cortex, superior division | 8 | 3.21 | 26 | -78 | 32 |
| Cluster 413 | R Frontal Pole | 8 | 2.81 | 16 | 48 | 32 |
| Cluster 414 | R Precuneus Cortex | 8 | 2.77 | 4 | -68 | 34 |
| Cluster 415 | L Precentral Gyrus | 8 | 2.84 | -54 | 10 | 34 |
| Cluster 416 | L Middle Frontal Gyrus | 8 | 2.87 | -38 | 34 | 34 |
| Cluster 417 | L Postcentral Gyrus | 8 | 2.82 | -50 | -20 | 36 |
| Cluster 418 | R Frontal Pole | 8 | 2.82 | 30 | 38 | 36 |
| Cluster 419 | L Lateral Occipital Cortex, superior division | 8 | 2.81 | -12 | -84 | 38 |
| Cluster 420 | L Lateral Occipital Cortex, superior division | 8 | 2.93 | -28 | -72 | 38 |
| Cluster 423 | L Lateral Occipital Cortex, superior division | 8 | 2.82 | -32 | -84 | 40 |

| | | | | | | |
|---|---|---|---|---|---|---|
| Cluster 424 | L Lateral Occipital Cortex, superior division | 8 | 2.86 | -32 | -78 | 42 |
| Cluster 425 | L Cingulate Gyrus, anterior division | 8 | 2.83 | -6 | -2 | 42 |
| Cluster 426 | R Angular Gyrus | 8 | 2.93 | 46 | -48 | 44 |
| Cluster 428 | L Precentral Gyrus | 8 | 2.94 | -12 | -22 | 46 |
| Cluster 429 | L Postcentral Gyrus | 8 | 3.16 | -40 | -32 | 48 |
| Cluster 432 | R Superior Frontal Gyrus | 8 | 2.93 | 6 | 38 | 50 |
| Cluster 433 | L Middle Frontal Gyrus | 8 | 2.92 | -28 | 22 | 52 |
| Cluster 434 | R Lateral Occipital Cortex, superior division | 8 | 2.86 | 16 | -60 | 56 |
| Cluster 435 | L Precuneus Cortex | 8 | 2.78 | -8 | -58 | 56 |
| Cluster 438 | R Precuneus Cortex | 8 | 2.88 | 2 | -64 | 58 |
| Cluster 439 | R Postcentral Gyrus | 8 | 3.03 | 46 | -28 | 60 |
| Cluster 440 | R Postcentral Gyrus | 8 | 3.01 | 14 | -40 | 62 |
| Cluster 441 | L Postcentral Gyrus | 8 | 3.13 | -44 | -32 | 62 |
| Cluster 442 | L Precentral Gyrus | 8 | 2.84 | -40 | -14 | 64 |
| Cluster 443 | L Lateral Occipital Cortex, superior division | 8 | 2.87 | -10 | -64 | 68 |
| Cluster 444 | | | | | | |

**Supplementary Table S32. [$UT_{CB2}$ - $US_{CB2}$]> [$CT_{CB3}$ - $CS_{CB3}$].** Descriptive statistics for clusters and local maxima showing greater increases in activation during Convolved Block 2 of Uncertain-Threat anticipation compared to Convolved Block 3 of Certain-Threat anticipation, each relative to their respective reference conditions (FDR $q$<0.05, whole-brain corrected).

| | | **mm³** | ***t*** | ***x*** | ***y*** | ***z*** |
|---|---|---|---|---|---|---|
| Cluster 1 | R Superior Parietal Lobule | 736 | 6.08 | 24 | -44 | 70 |
| | R Postcentral Gyrus | | 5.04 | 28 | -38 | 70 |
| Cluster 2 | L Superior Parietal Lobule | 624 | 5.13 | -18 | -52 | 66 |
| Cluster 3 | R Superior Frontal Gyrus | 576 | 5.11 | 16 | -10 | 74 |
| | R Precentral Gyrus | | 4.98 | 30 | -8 | 70 |
| Cluster 4 | L Superior Frontal Gyrus | 304 | 5.30 | -8 | -8 | 76 |
| Cluster 5 | R Precentral Gyrus | 200 | 4.92 | 40 | -2 | 52 |
| Cluster 7 | R Occipital Pole | 120 | 4.47 | 16 | -90 | 0 |
| Cluster 8 | L Occipital Pole | 88 | 4.88 | -4 | -98 | 0 |
| Cluster 9 | L Occipital Pole | 64 | 4.32 | -8 | -102 | -6 |
| Cluster 10 | R Superior Frontal Gyrus | 56 | 4.84 | 10 | 32 | 58 |
| Cluster 11 | R Superior Parietal Lobule | 56 | 4.90 | 32 | -54 | 60 |
| Cluster 12 | L Temporal Pole | 40 | 4.42 | -38 | 22 | -36 |
| Cluster 13 | L Superior Frontal Gyrus | 32 | 4.49 | -6 | 20 | 68 |
| Cluster 14 | L Temporal Pole | 24 | 4.81 | -30 | 12 | -34 |
| Cluster 16 | R Brain-Stem | 16 | 4.09 | 4 | -36 | -64 |
| Cluster 17 | L Superior Frontal Gyrus | 16 | 4.64 | -6 | 38 | 56 |
| Cluster 18 | L Temporal Fusiform Cortex, anterior division | 8 | 3.90 | -36 | -8 | -44 |
| Cluster 19 | L Temporal Occipital Fusiform Cortex | 8 | 3.80 | -32 | -56 | -18 |
| Cluster 20 | L Lingual Gyrus | 8 | 3.83 | -2 | -90 | -12 |
| Cluster 21 | L Lingual Gyrus | 8 | 4.00 | -2 | -78 | -10 |
| Cluster 22 | R Occipital Pole | 8 | 3.86 | 10 | -96 | -4 |
| Cluster 23 | L Occipital Pole | 8 | 3.87 | -14 | -96 | -4 |
| Cluster 25 | R Frontal Operculum Cortex | 8 | 4.01 | 42 | 24 | 6 |
| Cluster 26 | R Superior Frontal Gyrus | 8 | 3.90 | 20 | 22 | 46 |
| Cluster 27 | R Superior Frontal Gyrus | 8 | 3.76 | 4 | 32 | 54 |

| | | | | | | |
|---|---|---|---|---|---|---|
| Cluster 28 | R Superior Frontal Gyrus | 8 | 3.95 | 16 | 22 | 60 |
| Cluster 29 | L Superior Frontal Gyrus | 8 | 3.82 | -2 | 16 | 62 |
| Cluster 30 | R Superior Parietal Lobule | 8 | 3.82 | 16 | -54 | 66 |
| Cluster 31 | R Juxtapositional Lobule Cortex (formerly Supplementary Motor Cortex) | 8 | 3.77 | 4 | -4 | 70 |
| Cluster 32 | L Postcentral Gyrus | 8 | 3.90 | -22 | -38 | 74 |
